# Mapping Evidence Gap Between NMN and NR for Metabolic Outcomes: A Systematic Review, Transitivity Assessment, and Indirect Comparison Meta-Analysis

**DOI:** 10.64898/2026.04.07.716917

**Authors:** Alexander Tai Nguyen, Bryan Nguyen

## Abstract

**Background:** Nicotinamide mononucleotide (NMN) and nicotinamide riboside (NR) are NAD+ precursor supplements widely marketed for metabolic health benefits. Despite billions of dollars in annual sales, no head-to-head randomized controlled trial (RCT) has compared their effects on metabolic endpoints, and no systematic characterization of why reliable comparison is currently impossible has been published.

**Objective:** To characterize the structural heterogeneity of the NMN and NR trial evidence bases across population, dose, duration, and biomarker dimensions; to formally assess transitivity; and to estimate indirect NMN versus NR effects where methodologically feasible using the Bucher indirect comparison method.

**Methods:** Five databases (PubMed, Embase, Scopus, Web of Science, Cochrane CENTRAL) were searched from January 2018 to May 2025. Eligible studies were RCTs of oral NMN or NR versus placebo in adults reporting metabolic outcomes. A formal transitivity assessment was conducted comparing effect modifier distributions across NMN and NR trial arms. Random-effects pairwise meta-analyses were conducted for each precursor versus placebo, and Bucher indirect comparisons estimated NMN versus NR effects through the common placebo node. Risk of bias was assessed using RoB 2 and certainty of evidence using the GRADE/CINeMA framework.

**Results:** Fifteen studies (5 NMN, 10 NR; 740 participants) were included. The NMN and NR trial evidence bases were systematically asymmetric across every major effect modifier: NR was dosed 1.9 to 9.2 times higher than NMN on a molar basis; NMN trials were conducted predominantly in East Asian populations while NR trials were predominantly Western; and available NAD+ pharmacodynamic measures used incompatible assay matrices precluding indirect comparison. Across 14 metabolically comparable outcomes, no indirect comparison reached statistical significance and all were rated Very Low certainty by GRADE/CINeMA, consistent with the structural limitations of the evidence base. Leave-one-out sensitivity analyses showed zero pairwise significance changes and one indirect significance change (triglycerides upon exclusion of Conze 2019).

**Conclusion:** Current evidence is structurally insufficient to support reliable indirect comparison of NMN and NR for metabolic outcomes. The barriers are quantifiable and modifiable: future head-to-head trials should use equimolar dosing (approximately 1,150 mg NMN is molar-equivalent to 1,000 mg NR), harmonized whole-blood NAD+ assays reported in μmol/L, minimum 24 weeks duration, and enrollment of metabolically at-risk populations to generate interpretable comparative evidence.

**Registration:** PROSPERO 2026 CRD420261330487; registered prior to data screening.

## 1. Introduction

Nicotinamide adenine dinucleotide (NAD+) is an essential coenzyme present in every living cell, where it participates in more than 500 enzymatic reactions spanning glycolysis, the tricarboxylic acid cycle, fatty acid oxidation, and mitochondrial electron transport [31]. In addition to its role as a redox shuttle, NAD+ serves as a consumed substrate for three major families of signaling enzymes: the sirtuins (SIRT1 through SIRT7), which catalyze protein deacetylation and ADP-ribosylation reactions that regulate gene expression, mitochondrial biogenesis, and circadian rhythms [33]; poly(ADP-ribose) polymerases (PARPs), which coordinate DNA damage repair and genomic stability; and the cyclic ADP-ribose hydrolase CD38, a major NAD+ consumer whose expression increases with age and chronic inflammation [32]. Together, these NAD+-dependent processes position the molecule at the intersection of energy metabolism, stress responses, and aging.

A substantial body of evidence indicates that intracellular NAD+ levels decline progressively with age across multiple tissues, including skeletal muscle, liver, adipose tissue, and brain [31,32]. In murine models, this decline has been causally linked to insulin resistance, hepatic steatosis, endothelial dysfunction, neuroinflammation, and reduced exercise capacity [28,29]. Post-mortem human studies and cross-sectional analyses of circulating NAD+ metabolites support a similar trend in humans, although the magnitude and tissue specificity of the decline remain less well characterized [31]. The concept of restoring NAD+ to youthful levels as a therapeutic strategy for metabolic and age-related diseases has therefore gained considerable scientific and commercial attention.

Two NAD+ precursor supplements have emerged as the leading candidates for oral NAD+ augmentation in humans: nicotinamide mononucleotide (NMN) and nicotinamide riboside (NR) [1]. NMN is a nucleotide composed of nicotinamide, ribose, and a phosphate group, and it is a direct biosynthetic intermediate in the NAD+ salvage pathway. Its conversion to NAD+ requires a single adenylylation step catalyzed by nicotinamide mononucleotide adenylyltransferases (NMNAT1, NMNAT2, or NMNAT3), depending on the subcellular compartment [1]. NR, by contrast, is a nucleoside (nicotinamide plus ribose without the phosphate group) that enters the salvage pathway one step earlier. NR must first be phosphorylated by nicotinamide riboside kinases (NRK1 or NRK2) to form NMN before undergoing the same NMNAT-catalyzed adenylylation to NAD+ [1]. This biochemical difference has led to speculation that NMN may be more efficient at raising NAD+ levels because it bypasses the rate-limiting NRK phosphorylation step.

The pharmacokinetics of oral NAD+ precursor delivery add further complexity. NR is readily absorbed in the small intestine, but a substantial fraction is cleaved to nicotinamide by purine nucleoside phosphorylase before reaching the systemic circulation, meaning that only a portion of ingested NR enters cells intact [30]. NMN absorption was initially thought to require extracellular dephosphorylation to NR followed by re-phosphorylation inside the cell, but the identification of a putative NMN-specific transporter, Slc12a8, in murine gut epithelium suggests that direct cellular uptake of intact NMN may occur [27]. Whether Slc12a8-mediated transport operates efficiently in humans and at physiological concentrations remains debated. Additionally, differences in supplement formulation, capsule stability, and first-pass hepatic metabolism may influence the bioavailability of each precursor independently of their intrinsic biochemistry.

The clinical evidence base for NMN and NR has expanded rapidly since the first human RCTs were published in 2018 [7]. As of May 2025, at least five NMN [2,3,4,5,6] and ten NR [7,8,9,10,11,12,13,14,15,16] RCTs have reported metabolic outcomes, including fasting blood glucose, HbA1c, HOMA-IR, fasting insulin, lipid panel (total cholesterol, LDL, HDL, triglycerides), hepatic transaminases (ALT, AST), blood pressure (systolic and diastolic), body mass index, body weight, and circulating NAD+ concentrations. However, most trials are small (median sample size approximately 40 participants), of short duration (8 to 12 weeks), and compare a single precursor against placebo rather than against each other. The NMN trials have predominantly enrolled Asian populations (Japan, India) at doses of 250 to 300 mg per day, while NR trials have typically studied Western populations (Europe, North America) at higher doses of 500 to 2,000 mg per day [2–16]. No head-to-head RCT has directly compared NMN with NR on any clinical endpoint.

Several recent systematic reviews have synthesized the pairwise evidence for NMN or NR versus placebo individually. Zheng et al. (2024) conducted a meta-analysis of nine NMN RCTs and reported no significant effects on most cardiometabolic outcomes [24]. Alegre and Pastore (2024) reviewed both precursors narratively and concluded that evidence of metabolic benefit was limited for both [25]. Nascimento and Nogueira-de-Almeida (2024) focused specifically on NR in cardiometabolic disease and found insufficient evidence to recommend clinical use [26]. Importantly, none of these reviews attempted to compare NMN against NR, because no closed network of direct evidence exists. However, network meta-analysis (NMA) offers a framework for indirect comparisons between treatments that share a common comparator. When the evidence network is star-shaped, with two active treatments connected only through a shared placebo arm, the Bucher indirect comparison method provides a valid approach to estimating relative treatment effects, provided that the transitivity assumption holds [20].

The transitivity assumption requires that the distribution of effect modifiers is similar across the trials being compared. In the context of NMN versus NR, potential effect modifiers include participant age, sex, baseline metabolic status, ethnicity, supplement dose, and intervention duration. The known demographic and dosing differences between NMN and NR trial populations represent the principal threat to transitivity and must be carefully considered when interpreting indirect comparisons.

The objective of this systematic review was to: (1) characterize the structural heterogeneity of the NMN and NR trial evidence bases across population, dose, duration, and biomarker dimensions through a formal transitivity assessment; (2) conduct pairwise meta-analyses for each precursor versus placebo; (3) estimate indirect NMN versus NR comparisons via the Bucher method [20] for outcomes where indirect comparison was methodologically feasible; and (4) evaluate the certainty of evidence using the GRADE/CINeMA framework [21]. To our knowledge, this is the first study to formally characterize the evidence gap between NMN and NR and the first indirect comparison meta-analysis to systematically demonstrate why that gap cannot yet be reliably closed with existing trial data.

## 2. Methods

This systematic review and indirect comparison meta-analysis was conducted and reported in accordance with the Preferred Reporting Items for Systematic Reviews and Meta-Analyses (PRISMA) 2020 guidelines [17] and the PRISMA extension statement for network meta-analyses [18]. This review was registered in PROSPERO prior to data screening (March 2026) (registration ID: CRD420261330487). The manuscript title was subsequently revised to better reflect the study’s primary contribution as an evidence gap characterization; the registered and submitted titles differ accordingly.

### 2.1 Eligibility Criteria

Studies were eligible if they met the following PICO criteria: (P) adult participants (≥18 years) of any health status; (I) oral supplementation with NMN or NR at any dose; (C) placebo comparator; (O) at least one quantitative metabolic outcome (glycemic markers, lipid profile, hepatic enzymes, blood pressure, anthropometric measures, or blood NAD+ levels). Only RCTs (parallel or crossover design) were included. Non-randomized studies, animal or in vitro studies, reviews, protocols, combination supplements, and pediatric populations were excluded.

### 2.2 Information Sources and Search Strategy

Five electronic databases were searched from January 2018 to May 2025 (the 2018 start date was selected to coincide with the earliest registered clinical trials of commercially available NAD+ precursor formulations): PubMed (n = 286), Embase (n = 375), Scopus (n = 398), Web of Science (n = 319), and Cochrane CENTRAL (n = 309), yielding 1,687 records. The search strategy combined terms for the intervention (“nicotinamide mononucleotide” OR “NMN” OR “nicotinamide riboside” OR “NR” OR “NAD+ precursor”) with study design filters (“randomized” OR “clinical trial” OR “placebo”). The complete search strategy is provided in Supplementary Table S1. No language restrictions were applied.

### 2.3 Selection Process

After removal of 562 duplicate records, 1,125 unique records were screened in three stages: (1) title and abstract screening identified 874 clearly irrelevant records (no NMN/NR intervention, not an RCT, reviews or protocols, animal or in vitro studies, pediatric or type 1 diabetes populations); (2) 251 records were assessed at full text, of which 217 were excluded (133 upon full-text review confirming ineligibility that was apparent from title/abstract criteria, 65 upon re-screening of initially uncertain records, and 19 after detailed full-text assessment for reasons including non-extractable data, combination interventions, or out-of-scope populations); (3) 34 reports meeting inclusion criteria were further assessed, with 19 excluded from qualitative synthesis (trial registrations only, no extractable data, combination supplements, or duplicate publications). Fifteen studies were included in the qualitative synthesis (5 NMN, 10 NR), of which 8 (4 NMN, 4 NR) contributed data to the quantitative NMA. The selection process is detailed in the PRISMA flow diagram (Figure 1).

**Figure 1.**
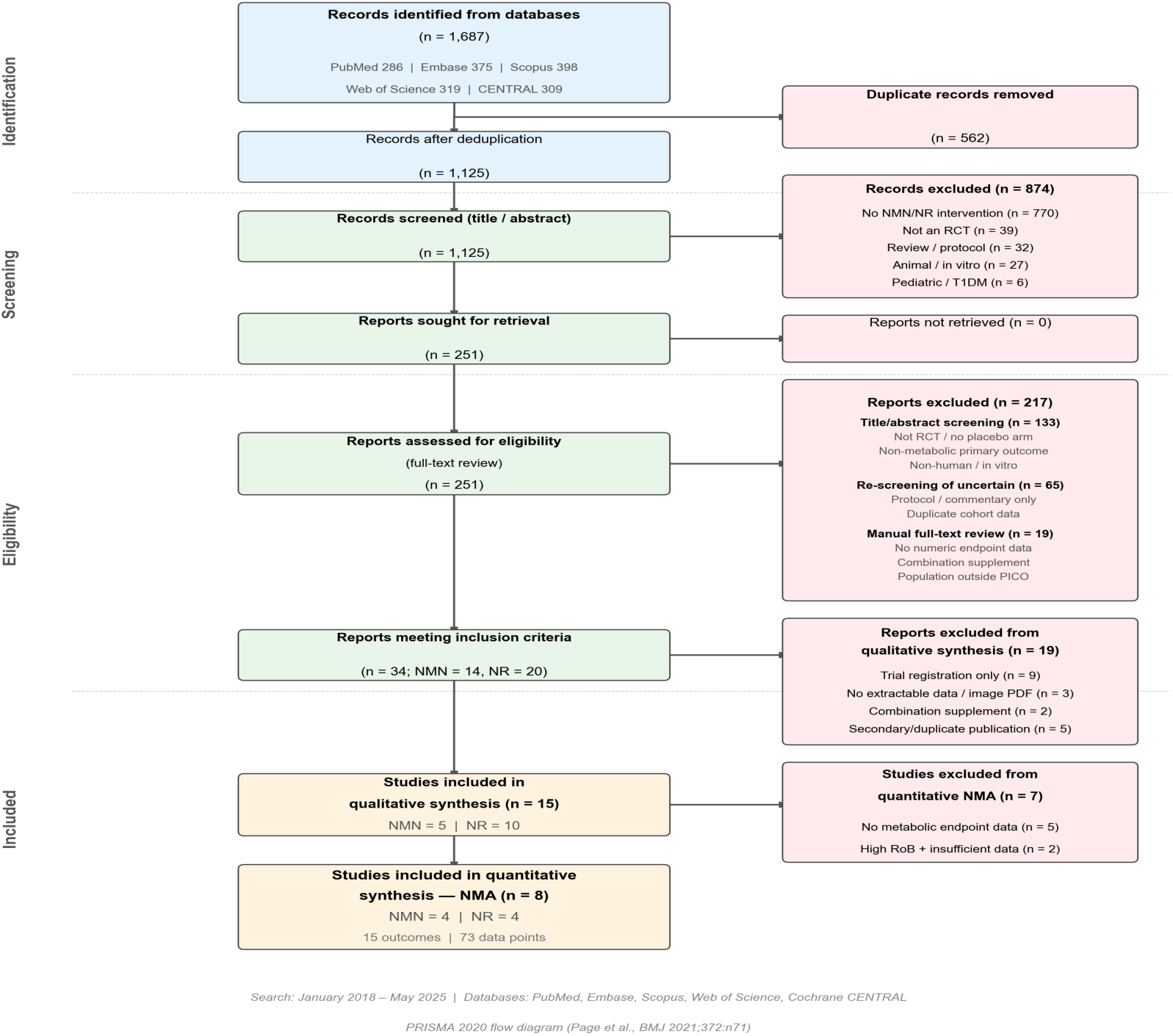
PRISMA 2020 flow diagram showing study identification, screening, eligibility, and inclusion.

### 2.4 Data Extraction

Data were extracted into a standardized form by two independent reviewers. For each study, we recorded: first author, year, country, design, population, sample size (randomized, intervention arm, control arm), dose (mg/day), intervention duration (weeks), participant demographics (age, sex, BMI), health status, trial registration, and funding source. For each outcome, we extracted mean change from baseline and standard deviation (SD) for intervention and control groups. When means and SDs were not directly reported, we applied standard conversions: standard error (SE) was converted to SD as SD = SE × √n; medians and interquartile ranges were converted using the method of Wan et al [22]. All data points were independently verified against original publications and supplementary materials. Inter-rater agreement was substantial to almost perfect across all phases: title/abstract screening (κ = 0.92), full-text screening (κ = 0.84), RoB 2 domain-level assessment (κ = 0.75), and data extraction (93.3% agreement; Supplementary Table S9).

### 2.5 Risk of Bias Assessment

Risk of bias was assessed at the study level using the revised Cochrane Risk of Bias tool (RoB 2) across five domains: randomization process, deviations from intended interventions, missing outcome data, measurement of the outcome, and selection of the reported result [19]. Each domain was rated as Low, Some Concerns, or High, with an overall judgment following RoB 2 algorithms. Results are presented as a traffic-light plot and summary bar chart (Figure 2). Detailed domain-level assessments with justifications are provided in Supplementary Table S4.

**Figure 2A.**
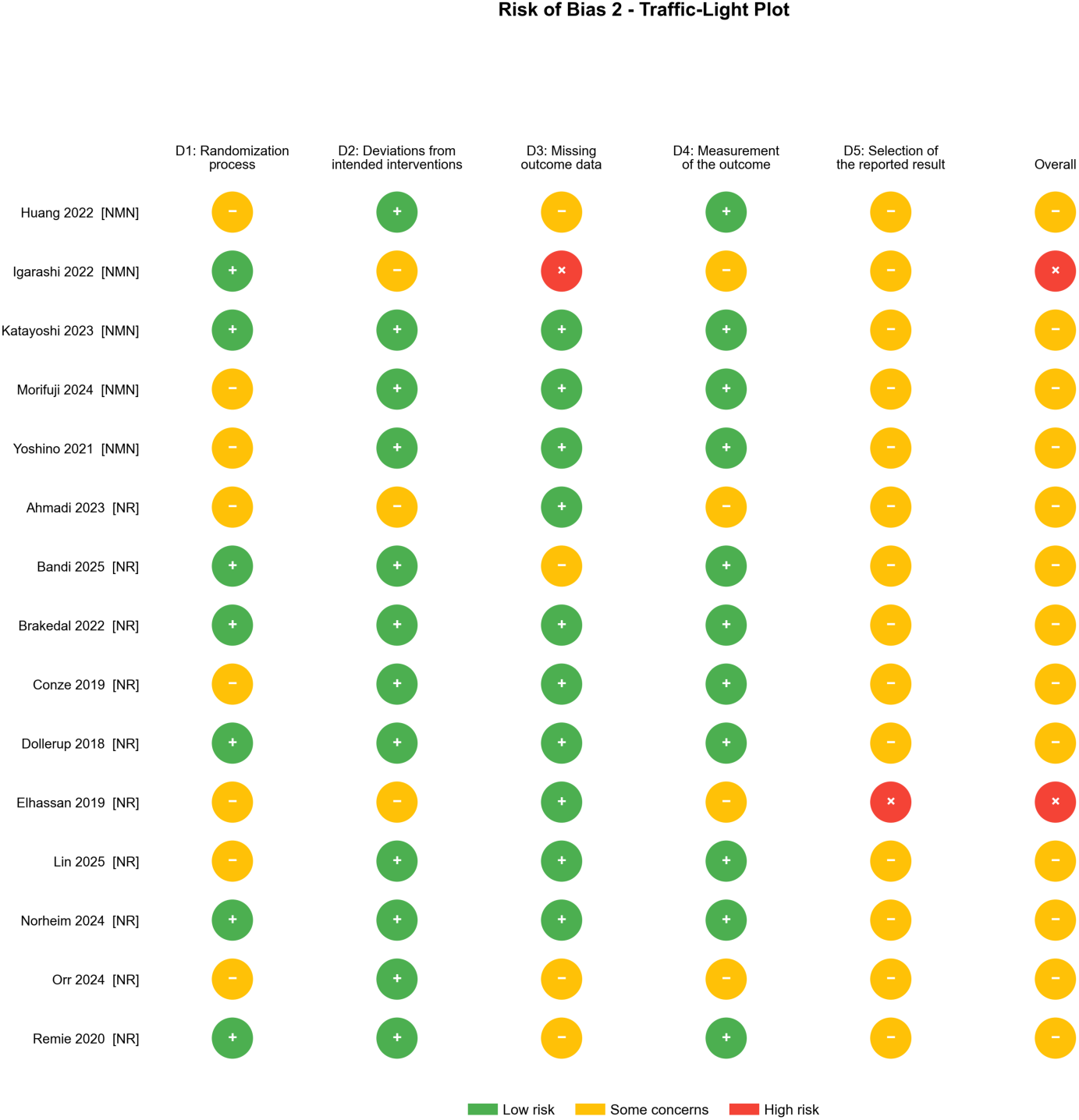
Risk of bias traffic-light plot showing domain-level RoB 2 judgments for each included study.

**Figure 2B.**
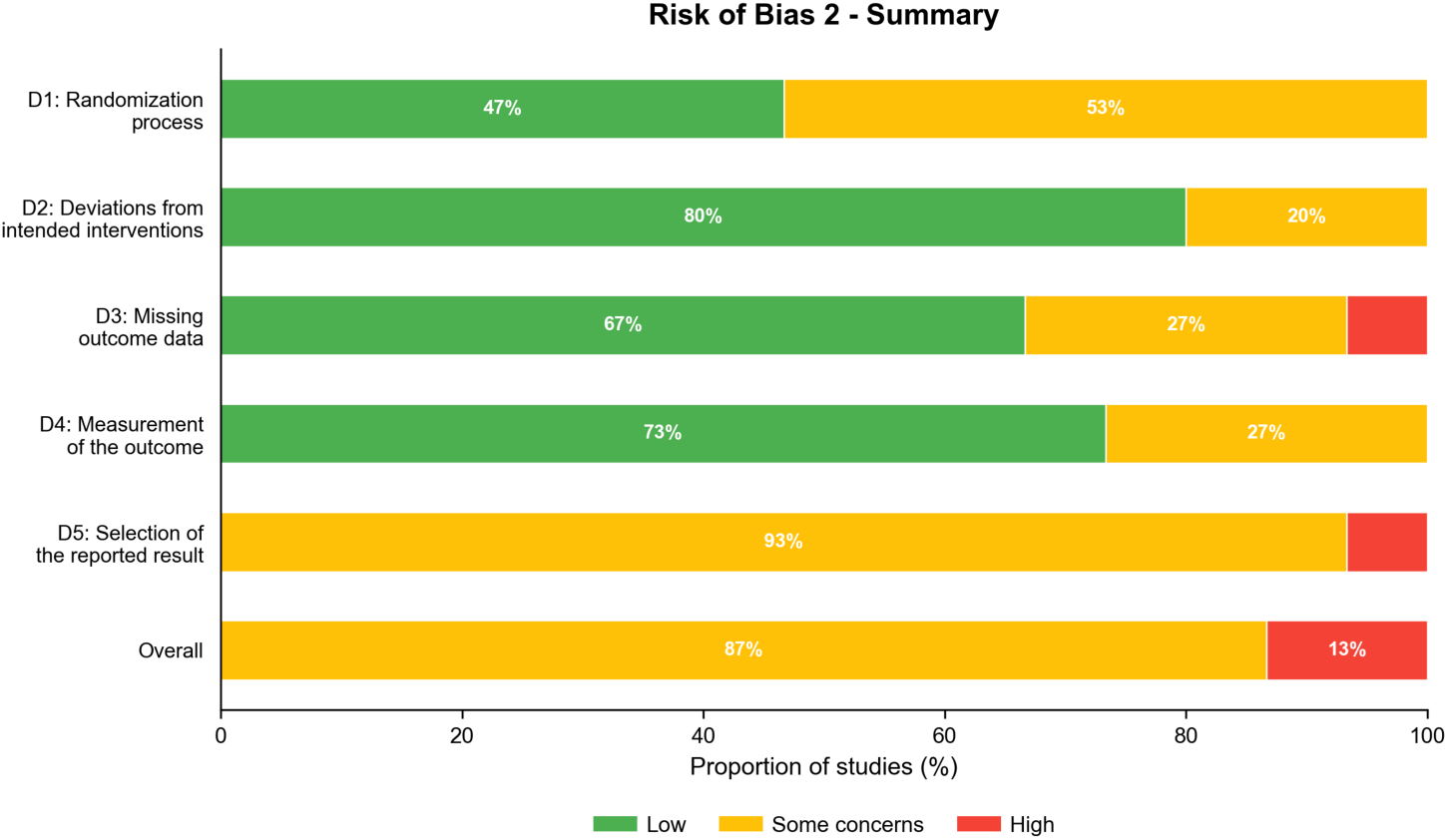
Risk of bias summary bar chart showing the distribution of Low, Some Concerns, and High ratings across all studies for each domain.

### 2.6 Data Synthesis and Statistical Analysis

#### Pairwise meta-analyses

For each precursor-placebo comparison, random-effects meta-analyses were conducted using the inverse-variance method with DerSimonian-Laird estimation of between-study variance (τ²) [23]. The summary effect measure was the mean difference (MD) with 95% confidence intervals (CIs). Statistical heterogeneity was quantified using the I² statistic, with thresholds of 25%, 50%, and 75% indicating low, moderate, and high heterogeneity, respectively.

#### Network meta-analysis

Because no closed loops of evidence existed (NMN and NR trials each compared only against placebo), the network geometry was star-shaped. Indirect comparisons of NMN versus NR were estimated using the Bucher method [20] (equivalent to a contrast-based network meta-analysis on a three-node network without closed loops): MD(NMN vs NR) = MD(NMN vs Placebo) minus MD(NR vs Placebo), with variance equal to the sum of variances of the two pairwise estimates. Indirect comparison was performed only for outcomes with compatible measurement definitions and units across NMN and NR arms. Outcomes with incompatible measurement context or unit scaling were excluded from indirect estimation.

#### Sensitivity analyses

Three predefined sensitivity analyses were conducted: (1) leave-one-out (LOO) analysis for each pairwise comparison, systematically omitting one study at a time and recording changes in pooled estimates and statistical significance; (2) LOO analysis for each indirect comparison, propagating LOO pairwise results through the Bucher formula; (3) exclusion of studies rated High risk of bias on the overall RoB 2 assessment.

#### Certainty of evidence

The certainty of evidence for each indirect comparison was evaluated using the GRADE framework adapted for NMA (CINeMA: Confidence in Network Meta-Analysis) [21]. Six domains were assessed: within-study bias, reporting bias, indirectness, imprecision, heterogeneity, and incoherence. Each domain was rated as No Concerns, Some Concerns, or Major Concerns, and certainty was downgraded accordingly from a starting level of High.

#### Software

All primary analyses were implemented in Python 3 using scipy (v1.14) for statistical computations (normal distribution CIs, Z-tests), numpy for numerical operations, and matplotlib for figure generation. Results were validated using R (v4.5) with the metafor package (v4.8) using restricted maximum likelihood (REML) estimation, confirming concordance of pairwise and indirect estimates. Multiplicity adjustment was performed using Bonferroni and Benjamini-Hochberg corrections across all tested indirectly comparable outcomes (n = 14; NAD+ excluded as non-comparable). The complete analytical pipeline is available as open-source code.

#### Publication bias

Formal assessment of publication bias using funnel plots or Egger’s regression test was conducted for all pairwise comparisons with two or more contributing studies; funnel plots and Egger’s regression test results are presented as exploratory visualizations in Supplementary Table S10 and Supplementary Figures. Formal interpretation was not performed because the number of studies per comparison was insufficient (k ≤ 4 for all outcomes; conventional guidance recommends k ≥ 10 for reliable interpretation). Accordingly, reporting bias was rated as Some Concerns for all outcomes in the GRADE/CINeMA assessment, based on the predominance of industry funding and the limited number of available studies.

## 3. Results

### 3.1 Study Selection

The database search identified 1,687 records, of which 562 were duplicates, leaving 1,125 unique records for title and abstract screening. After excluding 874 irrelevant records, 251 full-text reports were assessed for eligibility. Of these, 217 were excluded (133 upon full-text review confirming ineligibility apparent from title/abstract criteria, 65 upon re-screening of initially uncertain records, and 19 after detailed full-text assessment for non-extractable data, combination supplements, or populations outside the PICO framework). Thirty-four reports met the inclusion criteria, from which 19 were excluded from the qualitative synthesis (9 trial registrations only, 3 without extractable data, 2 combination supplements, and 5 secondary or duplicate publications). Fifteen studies (5 NMN, 10 NR) were included in the qualitative synthesis, and 8 studies (4 NMN, 4 NR) contributed 73 data points spanning 21 metabolic outcomes (15 reported by both precursors) to quantitative analyses, of which 14 were eligible for indirect comparison (Figure 1; Supplementary Figure S6).

### 3.2 Study Characteristics

Table 1 summarizes the characteristics of the 15 included studies. The five NMN trials (Yoshino 2021 [2], Huang 2022 [3], Igarashi 2022 [4], Katayoshi 2023 [5], Morifuji 2024 [6]) were conducted in the USA (n = 1), India (n = 1), and Japan (n = 3), with doses of 250 to 300 mg/day and durations of 8.6 to 12 weeks. NMN participants were predominantly from Asian populations with a mean BMI ranging from 22.5 to 35.0 kg/m². The ten NR trials (Dollerup 2018 [7], Conze 2019 [8], Elhassan 2019 [9], Remie 2020 [10], Brakedal 2022 [11], Ahmadi 2023 [12], Norheim 2024 [13], Orr 2024 [14], Lin 2025 [15], Bandi 2025 [16]) were conducted in Denmark (n = 2), Canada, the UK, the Netherlands, Norway, the USA (n = 3), and India (n = 1), with doses of 500 to 2,000 mg/day and durations of 3 to 12 weeks. NR cohorts were predominantly Western populations. Total randomized participants across all 15 studies numbered 740. Study populations ranged from healthy volunteers to individuals with obesity, prediabetes, chronic kidney disease, Parkinson’s disease, COPD, and mild cognitive impairment.

**Table 1.** Characteristics of included studies (n = 15).

| Study | Precursor | Country | N | Dose | Duration | Population | RoB 2 |
| --- | --- | --- | --- | --- | --- | --- | --- |

|  |  |  |  | (mg/d) | (wk) |  | Overall |
| --- | --- | --- | --- | --- | --- | --- | --- |
| Yoshino 2021 | NMN | USA | 25 | 250 | 10 | Prediabetic/insulin resistant | Some concerns |
| Huang 2022 | NMN | India | 62 | 300 | 8.6 | Healthy overweight | Some concerns |
| Igarashi 2022 | NMN | Japan | 42 | 250 | 12 | Healthy older men | High |
| Katayoshi 2023 | NMN | Japan | 34 | 250 | 12 | Healthy | Some concerns |
| Morifuji 2024 | NMN | Japan | 59 | 250 | 12 | Healthy older adults | Some concerns |
| Dollerup 2018 | NR | Denmark | 40 | 2000 | 12 | Obese insulin-resistant | Some concerns |
| Conze 2019 | NR | Canada | 140 | 1000 | 8 | Healthy overweight | Some concerns |
| Elhassan 2019 | NR | UK | 12 | 1000 | 3 | Healthy aged men | High |
| Remie 2020 | NR | Netherlands | 15 | 1000 | 6 | Healthy overweight/obese | Some concerns |
| Brakedal 2022 | NR | Norway | 30 | 1000 | 4.3 | Parkinson's disease treatment-naive | Some concerns |
| Ahmadi 2023 | NR | USA | 25 | 1000 | 6 | CKD (eGFR ~37) | Some concerns |
| Norheim 2024 | NR | Denmark | 40 | 2000 | 6 | COPD (ex-smokers) | Some concerns |
| Orr 2024 | NR | USA | 22 | 1000 | 10 | MCI (mild cognitive impairment) | Some concerns |
| Lin 2025 | NR | USA | 54 | 1000 | 6 | Hypertension | Some concerns |
| Bandi 2025 | NR | India | 140 | 500 | 8.6 | Healthy aging | Some concerns |

Abbreviations: NMN, nicotinamide mononucleotide; NR, nicotinamide riboside; RoB, risk of bias; wk, weeks.

Several systematic differences between NMN and NR trial populations are important for interpreting the indirect comparisons. NMN trials used a narrow dose range (250 to 300 mg/day), whereas NR doses varied nearly fourfold (500 to 2,000 mg/day), making dose-equivalence assumptions uncertain. The geographic and ethnic heterogeneity, with NMN trials concentrated in East Asia and NR trials predominantly in Northern Europe and North America, introduces potential confounding through differences in dietary NAD+ intake, genetic variation in NAD+ metabolism enzymes, and baseline cardiometabolic risk profiles. These population-level differences represent the principal threat to the transitivity assumption underlying the indirect comparisons.

To quantify the dosing disparity, we compared molar intakes using the molecular weights of NMN (334.2 g/mol) and NR chloride (290.7 g/mol, the formulation used in most NR trials). At 250 mg/day, NMN provides approximately 0.75 mmol/day; at 300 mg/day, 0.90 mmol/day. NR at 500 mg/day provides 1.72 mmol/day, at 1,000 mg/day 3.44 mmol/day, and at 2,000 mg/day 6.88 mmol/day. Thus, NR molar intake ranged from 1.9 to 9.2 times that of NMN across the included trials. This substantial molar dose asymmetry must be considered when interpreting comparative efficacy. Because NAD+ measures were not harmonizable for a valid indirect contrast, cross-precursor superiority in NAD+ elevation cannot be inferred from this dataset.

### 3.3 Transitivity Assessment

Transitivity requires that the distribution of effect modifiers be sufficiently similar across NMN and NR trial arms for the placebo node to serve as a valid common comparator. Table 2 presents a formal comparison of key effect modifier distributions across the two arms.

**Table 2.** Transitivity assessment: comparison of effect modifier distributions across NMN and NR trial arms.

| Effect Modifier | NMN Trials (n=5) | NR Trials (n=10) | Standardized Difference |
| --- | --- | --- | --- |
| Mean age (years) | 59.3 | 63.2 | Large |
| Female sex (%) | 46.8 | 39.7 | Moderate |
| Mean BMI (kg/m<sup>2</sup>) | 26.4 | 28.9 | Moderate |
| Molar dose (mmol/day), mean | 0.78 | 3.96 | Very large |
| Duration (weeks), mean | 10.9 | 7.0 | Moderate |
| Metabolic population (%) | 20 | 20 | None |
| Disease-specific population (%) | 0 | 40 | Large |
| Geographic origin | East Asia (60%),<br>South Asia (20%),<br>North America (20%) | Northern Europe (50%), North America (40%),<br>South Asia (10%) | Large |

The NMN and NR trial arms differed substantially on every major effect modifier. The most consequential asymmetry was molar dose: NR trials delivered a mean of 3.96 mmol/day of precursor versus 0.78 mmol/day for NMN, a 5.1-fold difference on average, ranging from 1.9-to 9.2-fold across individual trial pairs. Geographic and ethnic separation was systematic, with NMN trials concentrated in East Asia (Japan, India) and NR trials predominantly in Northern Europe and North America, introducing potential confounding from differences in dietary NAD+ precursor intake, genetic variation in salvage pathway enzymes (NAMPT, NRK1, CD38), and baseline cardiometabolic risk profiles. Additionally, 40% of NR trial participants were drawn from disease-specific populations (Parkinson’s disease, chronic kidney disease, COPD, mild cognitive impairment) with no analogous disease-specific representation in the NMN arm, creating clinical heterogeneity that further threatens transitivity.

The NAD+ pharmacodynamic endpoint, the most direct marker of precursor bioavailability, could not be validly compared indirectly because the NMN-contributing study (Huang 2022) reported blood cellular NAD+/NADH in pmol/mL while the NR-contributing study (Bandi 2025) reported blood NAD+ in μM, representing incompatible matrices and scales. This biomarker incompatibility is itself evidence of the absence of measurement standards in this field.

Collectively, these asymmetries indicate that the transitivity assumption underlying indirect comparison is substantially threatened in this evidence base. The Very Low certainty ratings observed across all 14 indirect comparisons are the expected analytical consequence of this structural inadequacy, not merely a reflection of small sample sizes.

### 3.4 Risk of Bias

All 15 included studies were assessed using RoB 2 [19] (Supplementary Table S4). Thirteen studies received an overall rating of “Some Concerns,” primarily due to insufficient detail on randomization sequence generation or allocation concealment, industry funding with potential conflicts of interest, and metabolic outcomes reported as secondary endpoints. Two studies were rated “High” risk of bias: Igarashi 2022 [4], due to a supplement preparation error resulting in 52% attrition (only 10 of 21 participants per arm analyzed at 12 weeks), and Elhassan 2019 [9], which was not pre-registered and reported metabolic data only in supplementary tables. Neither high-RoB study contributed data to the quantitative NMA.

### 3.5 Pairwise Meta-Analyses

#### 3.5.1 Glycemic outcomes

Glycemic markers are among the most clinically relevant endpoints for NAD+ precursor supplementation, given that preclinical studies have linked NAD+ repletion to improved insulin signaling through SIRT1-mediated deacetylation of insulin receptor substrate proteins [28,33]. Fasting blood glucose (FBG) showed no significant difference for NMN versus placebo (MD 0.02 mg/dL [95% CI: −4.25, 4.30]; p = 0.99; k = 4; I² = 47%) or NR versus placebo (MD −2.58 [−6.60, 1.43]; p = 0.21; k = 2; I² = 0%). HbA1c was unchanged with both NMN (MD −0.01% [−0.14, 0.12]; p = 0.91; k = 2) and NR (MD −0.20% [−0.48, 0.08]; p = 0.16; k = 1). Given that HbA1c reflects average glycemia over 8 to 12 weeks, the short trial durations may have been insufficient to detect meaningful shifts. HOMA-IR showed a non-significant reduction with NMN in a single study (MD −0.60 [−1.40, 0.20]; p = 0.14; k = 1) and NR (MD −0.19 [−0.90, 0.51]; p = 0.59; k = 1). Neither precursor demonstrated a significant effect on insulin resistance in this analysis, though both HOMA-IR estimates derive from single studies with limited power. Fasting insulin showed no significant effect for either precursor (NMN: MD −3.29 μIU/mL [−8.28, 1.69]; p = 0.20; k = 2; I² = 0%; NR: MD −0.42 [−2.65, 1.82]; p = 0.71; k = 1). Note that 17 of 36 pairwise comparisons across all outcomes are based on a single study (k = 1) and therefore represent individual trial estimates rather than pooled meta-analytic results.

*Individual pairwise forest plots for glycemic outcomes are presented in Supplementary Figure S1*.

#### 3.5.2 Lipid profile

Lipid metabolism is a key area of interest because sirtuin activation, particularly SIRT1 and SIRT3, has been shown to regulate hepatic lipogenesis and fatty acid oxidation in animal models [28,33]. No significant effects were observed for total cholesterol (NMN: MD −4.16 mg/dL [−17.03, 8.71]; p = 0.53; k = 2; I² = 0%; NR: MD −7.92 [−19.87, 4.02]; p = 0.19; k = 3), LDL cholesterol (NMN: MD −1.52 [−11.68, 8.64]; p = 0.77; k = 3; NR: MD −5.07 [−14.71, 4.57]; p = 0.30; k = 3), HDL cholesterol (NMN: MD −0.15 [−5.73, 5.44]; p = 0.96; k = 3; NR: MD −2.19 [−7.59, 3.22]; p = 0.43; k = 3), or triglycerides (NMN: MD −17.78 [−36.53, 0.98]; p = 0.063; k = 4; NR: MD 4.17 [−25.38, 33.71]; p = 0.78; k = 3; I² = 20%). The NMN triglyceride estimate was numerically larger in magnitude than other pairwise effects, but remained non-significant (p = 0.063) and should be interpreted cautiously given imprecision. The wide confidence intervals across all lipid comparisons reflect the small number of contributing studies and limited power to detect the modest effect sizes typical for dietary supplements.

*Individual pairwise forest plots for lipid outcomes are presented in Supplementary Figure S2*.

#### 3.5.3 Hepatic enzymes

Hepatic transaminases serve as surrogate markers for liver inflammation and hepatocellular injury, and are particularly relevant given the role of NAD+ in hepatic fatty acid oxidation and mitochondrial function [1]. Alanine aminotransferase (ALT) was not significantly changed by NMN (MD −1.13 U/L [−5.13, 2.87]; p = 0.58; k = 3; I² = 66%) or NR (MD −3.26 [−7.77, 1.25]; p = 0.16; k = 2). Aspartate aminotransferase (AST) was similarly not significant for either precursor (NMN: MD −0.30 [−1.99, 1.38]; p = 0.72; k = 3; I² = 18%; NR: MD −1.40 [−4.34, 1.54]; p = 0.35; k = 1). The moderate heterogeneity for ALT NMN (I² = 66%) suggests variability in hepatic response across the contributing studies.

*Individual pairwise forest plots for hepatic enzyme outcomes are presented in Supplementary Figure S3*.

#### 3.5.4 Blood pressure

Blood pressure is a well-established modifiable risk factor for cardiovascular disease, and preclinical evidence suggests that NAD+ repletion may improve endothelial function through SIRT1-mediated activation of endothelial nitric oxide synthase (eNOS) [33]. Systolic blood pressure (SBP) showed a non-significant reduction with NMN (MD −3.50 mmHg [−7.85, 0.85]; p = 0.11; k = 4; I² = 0%) but not with NR (MD 2.00 [−5.84, 9.84]; p = 0.62; k = 1). Diastolic blood pressure (DBP) was similarly non-significant for NMN (MD −2.55 [−5.60, 0.51]; p = 0.10; k = 4) and NR (MD 0.80 [−4.47, 6.07]; p = 0.77; k = 1). SBP and DBP point estimates were directionally negative for NMN, but both remained non-significant with confidence intervals crossing the null; these trends should be interpreted as exploratory and require prospective confirmation in adequately powered trials.

*Individual pairwise forest plots for blood pressure outcomes are presented in Supplementary Figure S4*.

#### 3.5.5 Anthropometric measures

Neither BMI (NMN: MD −0.04 kg/m² [−1.29, 1.21]; p = 0.95; k = 3; NR: MD −0.90 [−3.41, 1.61]; p = 0.48; k = 1) nor body weight (NMN: MD 0.61 kg [−3.80, 5.02]; p = 0.79; k = 3; NR: MD −0.30 [−3.22, 2.62]; p = 0.84; k = 1) was significantly altered by either precursor. These null findings are unsurprising given that trial durations of 8 to 12 weeks are generally too short to detect meaningful changes in body composition, and none of the trials were designed with weight loss as a primary endpoint.

*Individual pairwise forest plots for anthropometric outcomes are presented in Supplementary Figure S5*.

#### 3.5.6 Blood NAD+

Blood NAD+ concentration is the most direct pharmacodynamic marker of NAD+ precursor supplementation. However, the two contributing studies measured different constructs and scales: Huang 2022 reported blood cellular NAD+/NADH (pmol/mL), whereas Bandi 2025 reported blood NAD+ (uM). Because these are not directly harmonizable for a valid indirect contrast, NAD+ was retained as separate pairwise evidence within each precursor but excluded from NMN-versus-NR indirect comparison.

### 3.6 Network Meta-Analysis: Indirect Comparisons

The network geometry was star-shaped, with NMN and NR connected only through the common placebo node (Figure 3). Table 3 presents Bucher indirect comparison results for 14 metabolically comparable outcomes (NAD+ excluded for incompatibility of measurement context and units).

**Figure 3.**
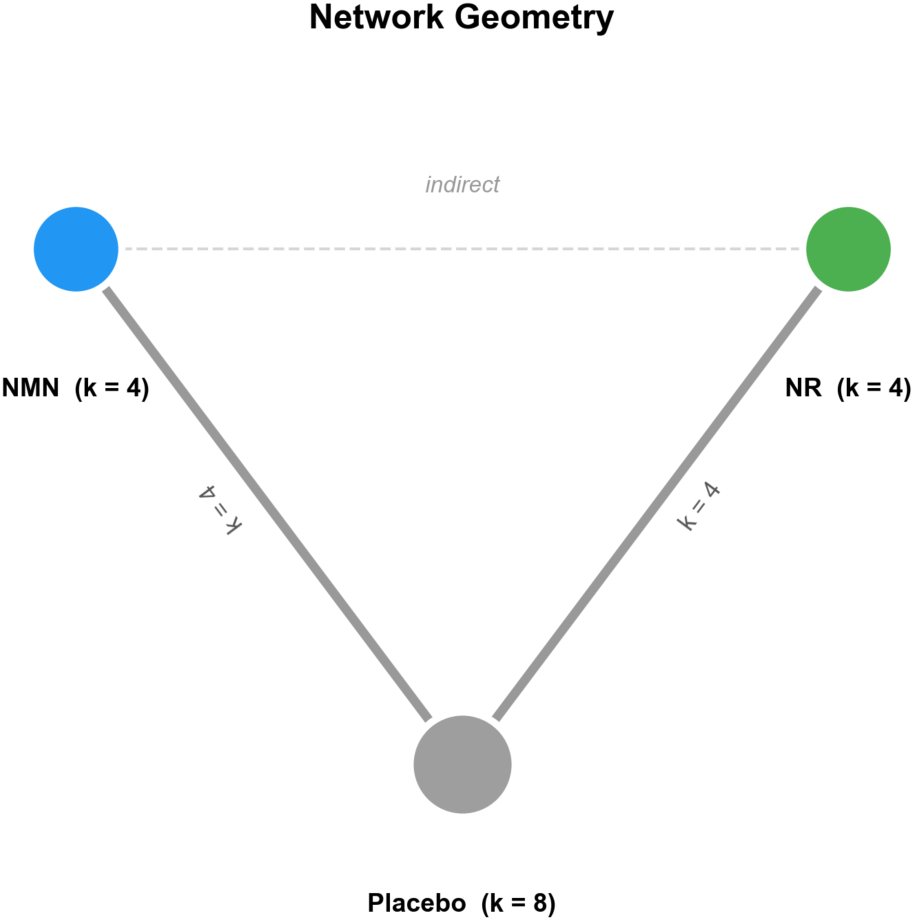
Network diagram showing the star-shaped geometry with NMN, NR, and Placebo nodes. Edge thickness is proportional to the number of studies.

**Table 3.**
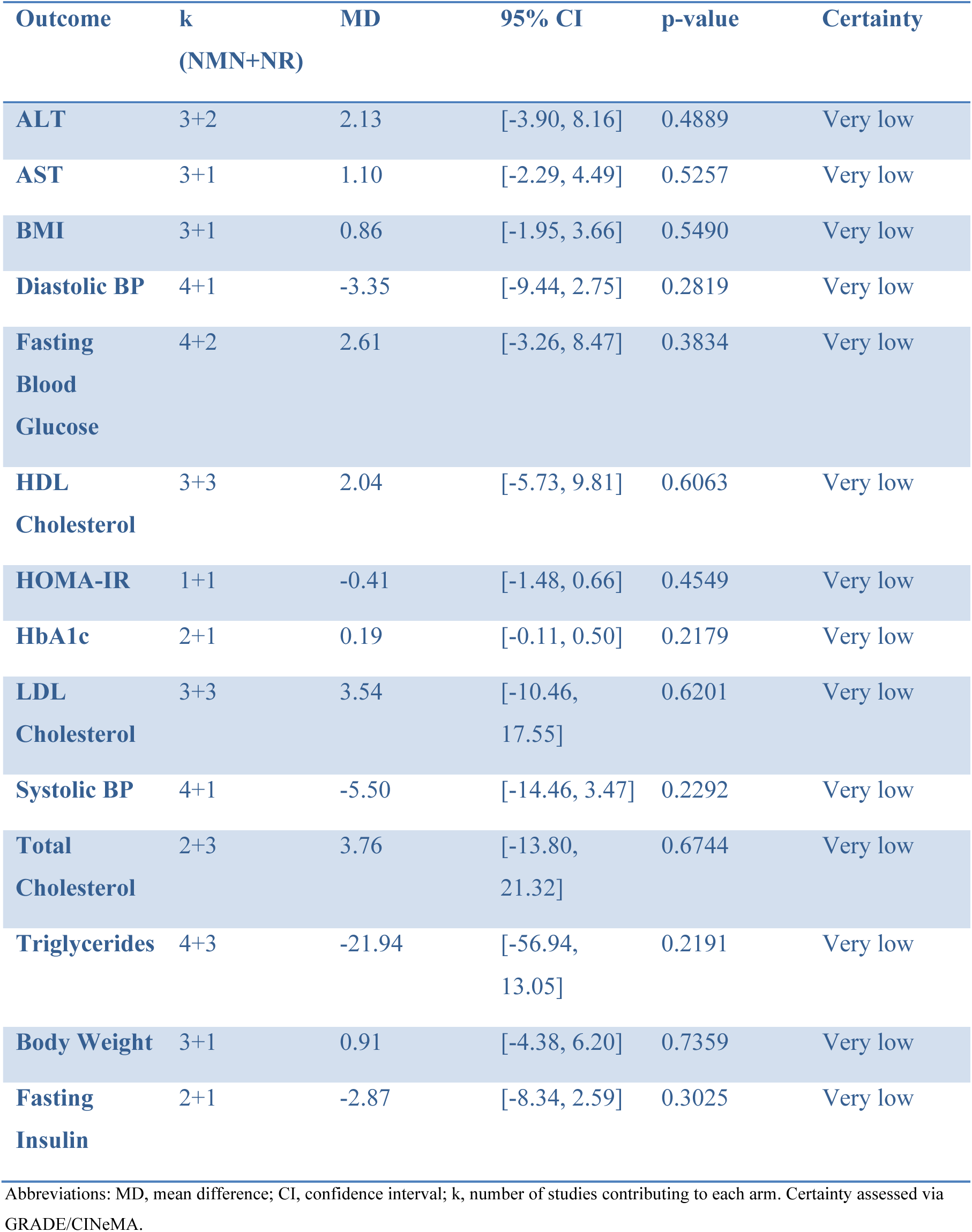
Indirect comparisons of NMN versus NR via Bucher method (14 outcomes; NAD+ excluded as non-comparable).

No indirect comparison showed a statistically significant difference between NMN and NR across the 14 comparable metabolic outcomes (all p > 0.10). Confidence intervals for most outcomes were wide and crossed the null, reflecting substantial imprecision driven by small sample sizes and sparse networks. Given 14 simultaneous indirect comparisons and generally low information size, these estimates should be interpreted as hypothesis-generating rather than confirmatory.

### 3.7 Certainty of Evidence

All 14 indirect comparisons were rated Very Low certainty by GRADE/CINeMA [21] (Figure 5; Supplementary Table S8). Downgrading was driven by within-study bias (Some Concerns in all comparisons, reflecting the preponderance of industry funding and secondary reporting of metabolic outcomes), reporting bias (Some Concerns due to insufficient studies for funnel plot assessment), indirectness (Some Concerns due to population heterogeneity, as NMN trials enrolled predominantly Asian participants while NR trials enrolled predominantly Western participants, and dose ranges differed substantially: 250 mg/day for NMN vs 500 to 2,000 mg/day for NR), imprecision (Some to Major Concerns), and heterogeneity (No to Some Concerns).

**Figure 4.**
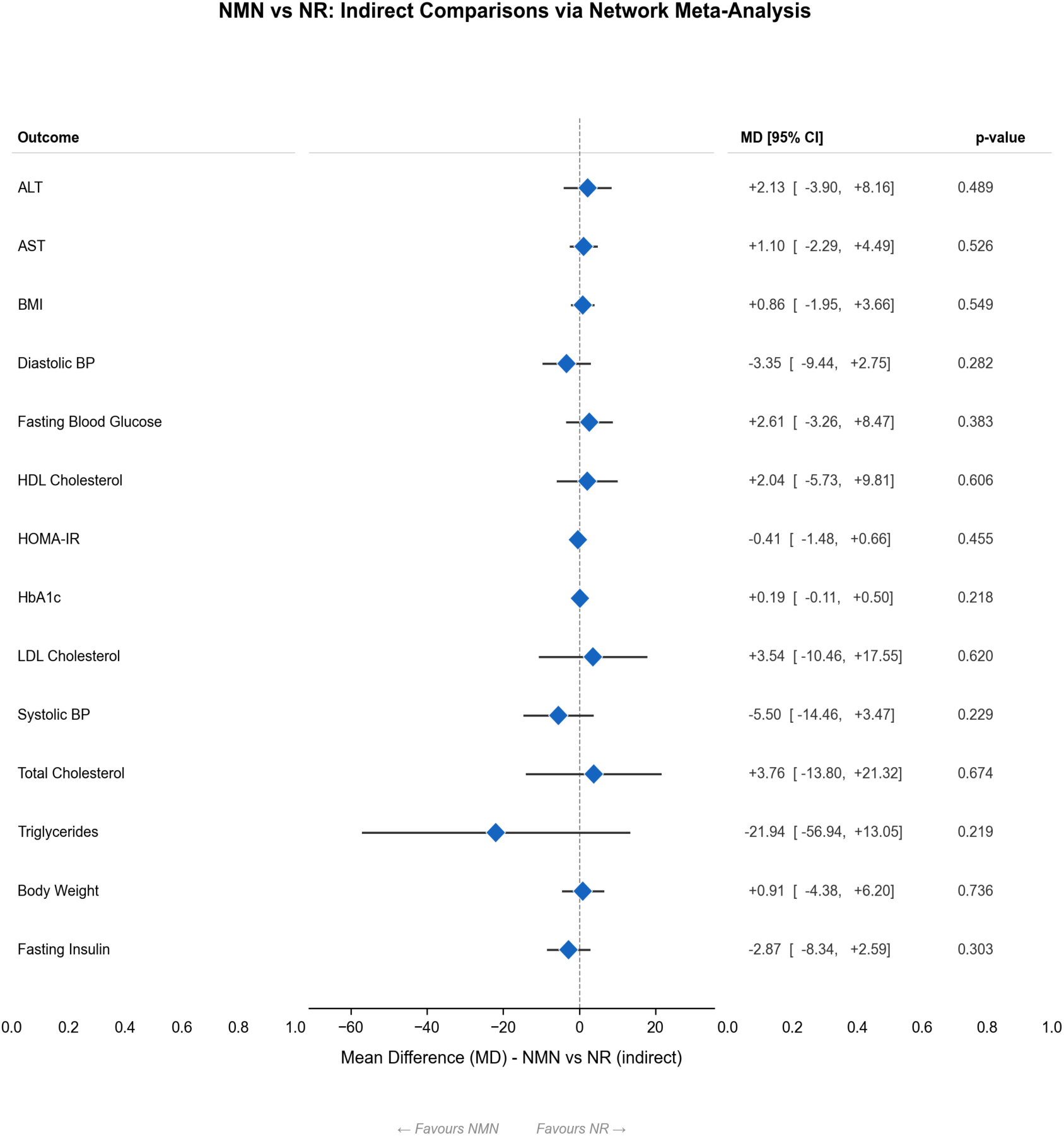
Forest plot of indirect comparisons (NMN vs NR) for 14 metabolically comparable outcomes. No indirect comparison reached statistical significance.

**Figure 5.**
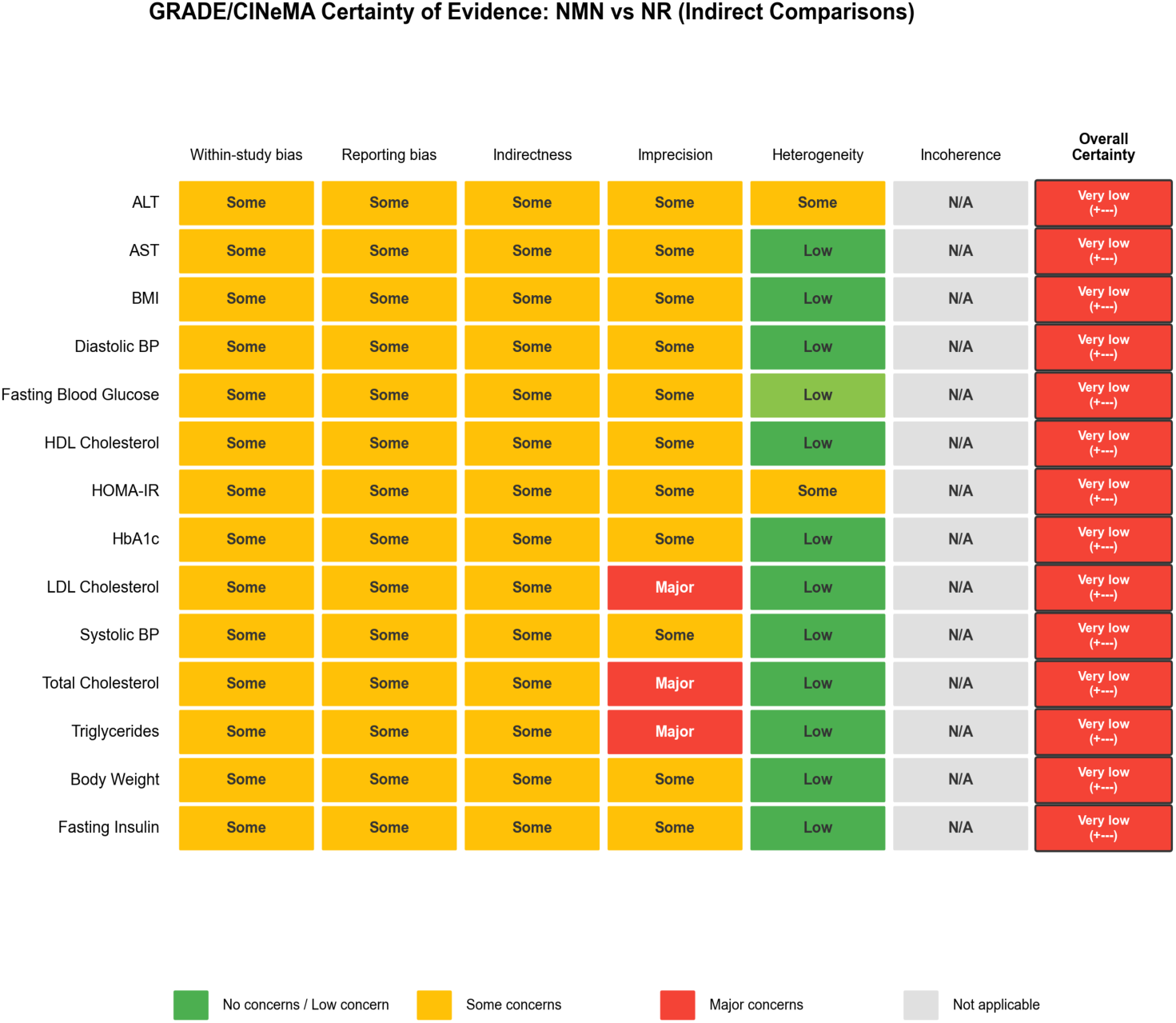
GRADE/CINeMA certainty-of-evidence heatmap for 14 indirectly comparable NMN vs NR outcomes (NAD+ excluded as non-comparable). All assessed comparisons were rated Very Low certainty.

Incoherence was Not Applicable for all outcomes because the star-shaped network contained no closed loops. NAD+ was not rated in indirect certainty because it was excluded from indirect estimation as non-comparable.

### 3.8 Sensitivity Analyses

Leave-one-out analyses were performed for all 56 pairwise iterations across all outcomes and comparisons. No pairwise significance changes were identified in any leave-one-out iteration, indicating that no single study exerts disproportionate influence on any pairwise estimate. Individual leave-one-out forest plots for ALT, SBP, and fasting insulin pairwise comparisons are presented in Supplementary Figure S7 as representative robustness checks.

Leave-one-out analysis of the 13 indirect comparisons amenable to LOO (56 total iterations; HOMA-IR excluded because both arms comprised a single study; Supplementary Table S7, which annotates comparisons where only one arm contributed multiple studies, explaining narrow LOO ranges for outcomes such as HbA1c) revealed one significance change: the triglyceride indirect comparison became nominally significant (p = 0.046, uncorrected) upon exclusion of Conze 2019, though this did not survive Bonferroni correction (threshold p < 0.0036), with the effect driven by the remaining NR studies. All other indirect comparisons remained non-significant regardless of which study was excluded (Figure 6).

**Figure 6.**
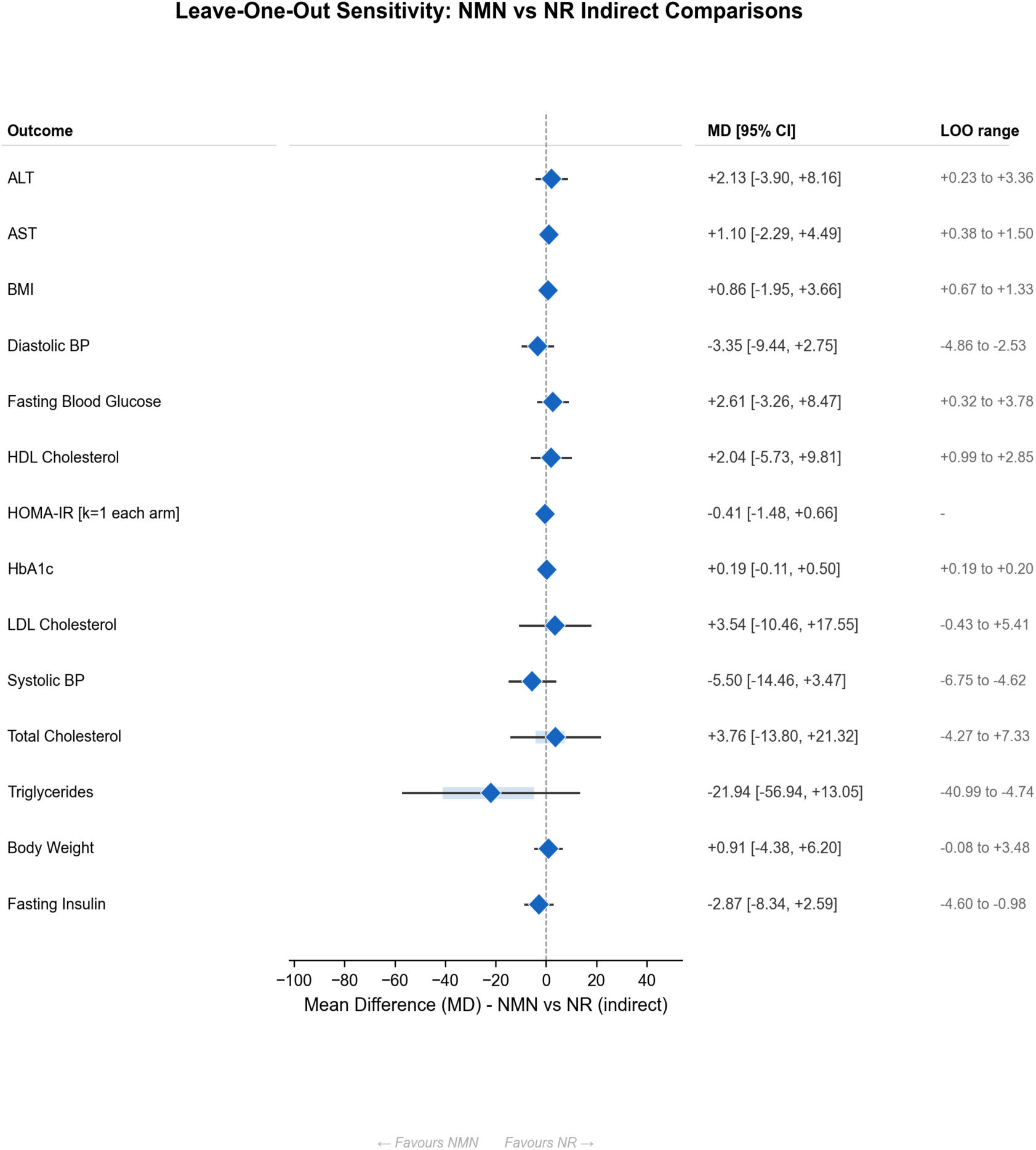
Leave-one-out sensitivity summary for all indirect NMN vs NR comparisons. One study exclusion (Conze 2019) changed triglycerides to nominal significance; all other outcomes remained non-significant.

The high-RoB exclusion analysis had no effect on any result, as neither high-RoB study (Igarashi 2022 [4], Elhassan 2019 [9]) contributed data to the quantitative meta-analysis. Igarashi 2022 [4] was excluded due to 52% attrition from a supplement preparation error, and Elhassan 2019 [9] was excluded due to non-registration and insufficient extractable metabolic data.

### 3.9 Exploratory Subgroup Observations

Although the small number of studies precluded formal subgroup meta-analysis (which requires adequate k within each stratum), a narrative comparison of results across metabolically impaired versus healthy populations provides informative context. Among the NMN trials, Yoshino 2021 [2] enrolled prediabetic obese women (BMI > 30) and showed the largest insulin-sensitivity improvements (hyperinsulinemic clamp) among NMN studies, although results did not reach significance as single-study estimates. The only HOMA-IR estimate available (Huang 2022 [3]; MD −0.60 [−1.40, 0.20]; p = 0.14) was also non-significant. Katayoshi 2023 [5] and Morifuji 2024 [6], which enrolled healthy volunteers, showed no glycemic improvements. Among the NR trials, Dollerup 2018 [7] enrolled obese men and reported numerical but non-significant improvements in hepatic lipid content, while Brakedal 2022 [11] (Parkinson’s disease) and Ahmadi 2023 [12] (chronic kidney disease) studied populations with specific non-metabolic pathologies. These studies were retained in the primary synthesis because eligibility criteria were prespecified to include adult randomized trials reporting metabolic outcomes regardless of underlying disease context; however, disease-specific pathophysiology may limit transitivity and external validity for general metabolic populations.

This pattern suggests that baseline metabolic status may be an important effect modifier: populations with established insulin resistance, hepatic steatosis, or dyslipidemia may benefit more from NAD+ repletion than metabolically healthy individuals. The floor effect in healthy populations, where baseline values are already within normal ranges, may obscure meaningful treatment effects. Future trials should stratify randomization by metabolic status and pre-specify subgroup analyses to test this hypothesis directly.

## 4. Discussion

### 4.1 Summary of Findings

This systematic review represents, to our knowledge, the first formal characterization of the evidence gap between NMN and NR for metabolic outcomes, and the first indirect comparison meta-analysis to systematically demonstrate why that gap cannot yet be reliably closed. The primary finding is not the indirect comparison estimates themselves, which were uniformly non-significant and rated Very Low certainty, but the structural characterization of why reliable comparison is currently impossible: a 1.9-to 9.2-fold molar dose asymmetry, systematic ethnic and geographic separation of trial populations, incompatible NAD+ biomarker definitions, and a sparse evidence network that propagates these asymmetries directly into imprecise, low-certainty indirect estimates. None of the 14 indirect comparisons reached statistical significance. Pairwise analyses showed generally null or imprecise effects across both precursors for most endpoints.

The certainty of evidence was Very Low for all indirect comparisons [21], substantially limiting definitive inference.

### 4.2 Interpretation of the NAD+ Finding

The NAD+ findings require careful interpretation because the available NMN and NR studies did not report directly comparable biomarkers. Huang 2022 reported blood cellular NAD+/NADH in pmol/mL, while Bandi 2025 reported blood NAD+ in uM. This methodological mismatch precludes a valid indirect NMN-versus-NR contrast for NAD+ and reinforces the need for future trials to use harmonized assays, matrix definitions, and unit reporting.

Mechanistically, first-pass metabolism remains relevant as within-arm pharmacodynamic context rather than as evidence for an indirect NMN-versus-NR effect. A substantial fraction of orally ingested NR is cleaved to nicotinamide by purine nucleoside phosphorylase in the gut lumen and liver before reaching the systemic circulation [30]. Despite this hepatic catabolism, several NR trials still showed substantial pairwise NAD+ increases, possibly because higher absolute NR doses (up to 2,000 mg/day) compensated for first-pass losses. NMN may also be subject to gastrointestinal degradation, but the extent of this process in humans remains insufficiently characterized.

From a clinical perspective, future trials should prioritize harmonized NAD+ assay matrices, unit definitions, and reporting standards so that pharmacodynamic findings can be compared across precursors and linked to patient-relevant metabolic outcomes.

### 4.3 Context with Prior Reviews

Our pairwise findings are broadly consistent with recent systematic reviews. Zheng et al. (2024) conducted the most comprehensive NMN-focused meta-analysis to date, pooling 9 RCTs (n = 412) and reporting non-significant effects on fasting glucose, HbA1c, HOMA-IR, lipid profile, and blood pressure [24]. Our NMN pairwise estimates are directionally concordant with theirs, though minor numerical differences exist due to differences in study inclusion and data extraction; notably, four NMN trials included by Zheng et al. did not meet the present review’s eligibility and harmonized extraction criteria for quantitative synthesis. Alegre and Pastore (2024) provided a narrative review of both NMN and NR, concluding that while preclinical evidence was promising, clinical translation remained limited [25]. They specifically highlighted the absence of head-to-head trials as a critical gap. Nascimento and Nogueira-de-Almeida (2024) focused on NR in cardiometabolic disease and found that, despite NR’s consistent ability to raise circulating NAD+, downstream metabolic benefits were not reliably observed [26].

This review extends prior work by formally comparing NMN against NR through Bucher indirect comparison [20] and critically by providing the first systematic diagnosis of why that comparison produces uninformative results. Prior reviews synthesized each precursor in isolation against placebo; none characterized the structural compatibility of the two evidence bases or quantified the barriers to reliable indirect inference. The finding that all 14 indirect comparisons returned Very Low certainty is not a failure of the analysis; it is the informative result. A well-designed indirect comparison applied to a structurally incompatible evidence base should return Very Low certainty, and documenting this formally is the necessary precondition for designing trials that can do better.

### 4.4 Explanations for Null Metabolic Findings

Several factors likely contribute to the absence of detectable metabolic differences between NMN and NR. The included trials had small sample sizes (median n approximately 40), providing insufficient statistical power; a post hoc illustrative calculation (two-sided alpha = 0.05, 80% power, target difference = 5 mg/dL, assumed common SD approximately 16 mg/dL based on observed trial-level FBG variability) indicates that approximately 80 participants per arm would be required in a direct two-arm comparison, far exceeding the effective sample sizes here. Trial durations were short (median 8 to 12 weeks), likely too brief for sirtuin-dependent metabolic reprogramming to produce measurable changes in clinical biomarkers [33]. Most study populations were metabolically healthy or mildly impaired, creating a floor effect. Numerical but non-significant improvements in HOMA-IR were observed with both NMN (MD −0.60; Huang 2022 [3]) and NR (MD −0.19; Dollerup 2018 [7]), suggesting that metabolically at-risk populations may show greater responsiveness to NAD+ repletion. This is a hypothesis that requires prospective testing in adequately powered trials.

The dose asymmetry between NMN and NR trials warrants particular attention. On a mass basis, NMN was tested at 250 to 300 mg/day while NR was tested at 500 to 2,000 mg/day. Our molar dose analysis (see Section 3.2) revealed that NR molar intake was 1.9-to 9.2-fold greater than NMN across included trials. This asymmetry is a major threat to the transitivity assumption and limits interpretability of indirect contrasts. Future head-to-head trials should employ equimolar dosing to enable pharmacologically meaningful comparison.

Finally, circulating NAD+ may be an imperfect surrogate for tissue-level NAD+ in metabolically relevant organs. The distribution of NAD+ to skeletal muscle, liver, and adipose tissue may differ between precursors even when blood levels diverge, and tissue-specific measurements using muscle biopsy or advanced imaging techniques would be needed to resolve this question.

### 4.5 Strengths

This study has several notable strengths. It is the first indirect comparison meta-analysis and formal transitivity assessment to characterize the evidence gap between NMN and NR for metabolic outcomes, addressing a clinically and commercially relevant question that cannot be answered by existing pairwise reviews [24,25,26] or narrative syntheses. We conducted a systematic search across five major databases with broad inclusion criteria to capture the available evidence.

The analytical approach was rigorous and multi-layered. We extracted 21 distinct metabolic outcomes with 73 individual data points, performed GRADE/CINeMA [21] certainty assessment for all 14 indirectly comparable outcomes, and conducted multiple predefined sensitivity analyses, including leave-one-out for both pairwise and indirect estimates and exclusion of high-risk-of-bias studies. All data extraction was independently performed by two reviewers with substantial inter-rater agreement (kappa = 0.75 to 0.92 across phases; Supplementary Table S9), and all extracted values were verified against original publications. Primary Python-based analyses were independently validated using R/metafor with REML estimation, confirming concordance of results.

### 4.6 Limitations

Several important limitations must be acknowledged. The star-shaped network geometry, while appropriate for the Bucher method [20], precludes assessment of incoherence (consistency), a core NMA diagnostic. In addition, 17 of 36 pairwise comparisons (47%) are based on a single study (k = 1), meaning these are single-trial estimates rather than pooled meta-analytic results. NAD+ could not be validly compared indirectly because the available NMN and NR trials reported non-comparable biomarker definitions and units. Crossover and specialized-population studies also introduce additional clinical heterogeneity that may violate transitivity. For crossover trials, between-person variance was used as an approximation in pooled analyses because within-person correlation parameters were unavailable, which may overestimate variance and widen confidence intervals for affected comparisons.

Population heterogeneity between NMN and NR trials represents the most serious threat to the transitivity assumption. NMN trials were predominantly conducted in Asian populations (Japan, India), whereas NR trials were predominantly Western (Denmark, USA, Canada, UK, Netherlands, Norway). Genetic polymorphisms in NAD+ metabolic enzymes, including NAMPT, NRK1, and CD38, differ across ethnic groups and could differentially affect precursor efficacy. Dietary patterns also vary systematically: Japanese diets typically contain higher levels of tryptophan and niacin from fish and fermented foods, potentially affecting baseline NAD+ status and the marginal benefit of supplementation.

Dose asymmetry (250 mg NMN vs 500 to 2,000 mg NR) introduces further heterogeneity that cannot be resolved without pharmacokinetic studies establishing dose equivalence. Our molar dose analysis showed NR was dosed 1.9-to 9.2-fold higher than NMN on a per-molecule basis, yet formal assessment of publication bias via funnel plots or Egger’s test was not possible because no pairwise comparison included more than four studies (minimum recommended: k ≥ 10). Many outcomes relied on a single study per arm (k = 1), making pairwise estimates inherently fragile and unable to account for between-study variability. The predominance of industry funding or supplement provision across included trials raises concerns about reporting bias, particularly for metabolic outcomes that were often secondary endpoints and may have been selectively reported. The very low certainty of all 14 indirectly comparable outcomes, as determined by GRADE/CINeMA, means that the true effects could be substantially different from the estimated effects, and future research is very likely to change these estimates.

### 4.7 Implications for Practice and Research

For clinical practice, current evidence does not support choosing NMN over NR (or vice versa) for any metabolic health indication based on indirect comparative evidence. Marketing claims of metabolic superiority for either precursor are not supported by the present evidence base.

For future research, the structural barriers identified in this review are quantifiable and directly actionable. A definitive head-to-head trial should address each barrier explicitly. First, dosing must be equimolar: using molecular weights of NMN (334.2 g/mol) and NR chloride (290.7 g/mol), 1,000 mg NR provides approximately 3.44 mmol/day, requiring approximately 1,150 mg NMN/day for molar equivalence, substantially higher than the 250–300 mg/day used in all existing NMN trials. Second, NAD+ pharmacodynamic assessment must be harmonized: whole-blood NAD+ measured by enzymatic cycling assay and reported in μmol/L should be the pre-specified primary pharmacodynamic endpoint, enabling valid cross-precursor comparison. Third, the target population should be metabolically at-risk, defined by HOMA-IR ≥ 2.5 or fasting glucose 100–125 mg/dL, to avoid the floor effects that dominate healthy volunteer studies. Fourth, duration should be a minimum of 24 weeks to allow HbA1c-detectable changes in glycemic status. Fifth, sample size should exceed 100 participants per arm based on a two-sided alpha of 0.05, 80% power, and a target difference of 5 mg/dL in fasting glucose with an assumed SD of 16 mg/dL. Sixth, funding should be independent of supplement manufacturers to mitigate the reporting bias documented throughout the existing evidence base. A trial meeting these six criteria would be definitive; none of the 15 trials in this review met more than two.

## 5. Conclusion

In this systematic review and indirect comparison meta-analysis, we provide the first formal characterization of the evidence gap between nicotinamide mononucleotide and nicotinamide riboside for metabolic outcomes. The primary finding is structural: the NMN and NR trial evidence bases are systematically incompatible for reliable indirect comparison, differing by 1.9-to 9.2-fold on a molar dose basis, separated by geography and ethnicity in ways that threaten transitivity, and using incompatible NAD+ biomarker definitions that preclude pharmacodynamic comparison. The consequence of these structural problems is precisely what the analysis found: 14 indirect comparisons across clinically relevant metabolic outcomes, all non-significant, all rated Very Low certainty. These results do not indicate that NMN and NR are equivalent; they indicate that the current evidence base cannot distinguish them. Closing this gap requires head-to-head trials with equimolar dosing (approximately 1,150 mg NMN equivalent to 1,000 mg NR), harmonized NAD+ assay standards, metabolically at-risk populations, and independent funding. Until such trials exist, evidence-based guidance on choosing between NMN and NR for metabolic health remains impossible.

## Declarations

## Data Availability

The complete dataset, analytical code, and all figures are available at Zenodo: https://doi.org/10.5281/zenodo.19403850.

## Author Contributions

ATN: Conceptualization, Data curation, Formal analysis, Investigation, Methodology, Project administration, Software, Validation, Visualization, Writing – original draft, Writing - review & editing. BN: Data curation, Investigation, Validation, Writing - review & editing.

## Ethics Approval

Not applicable (systematic review of published studies).

## Supplementary Materials

**Supplementary Figure S1.**
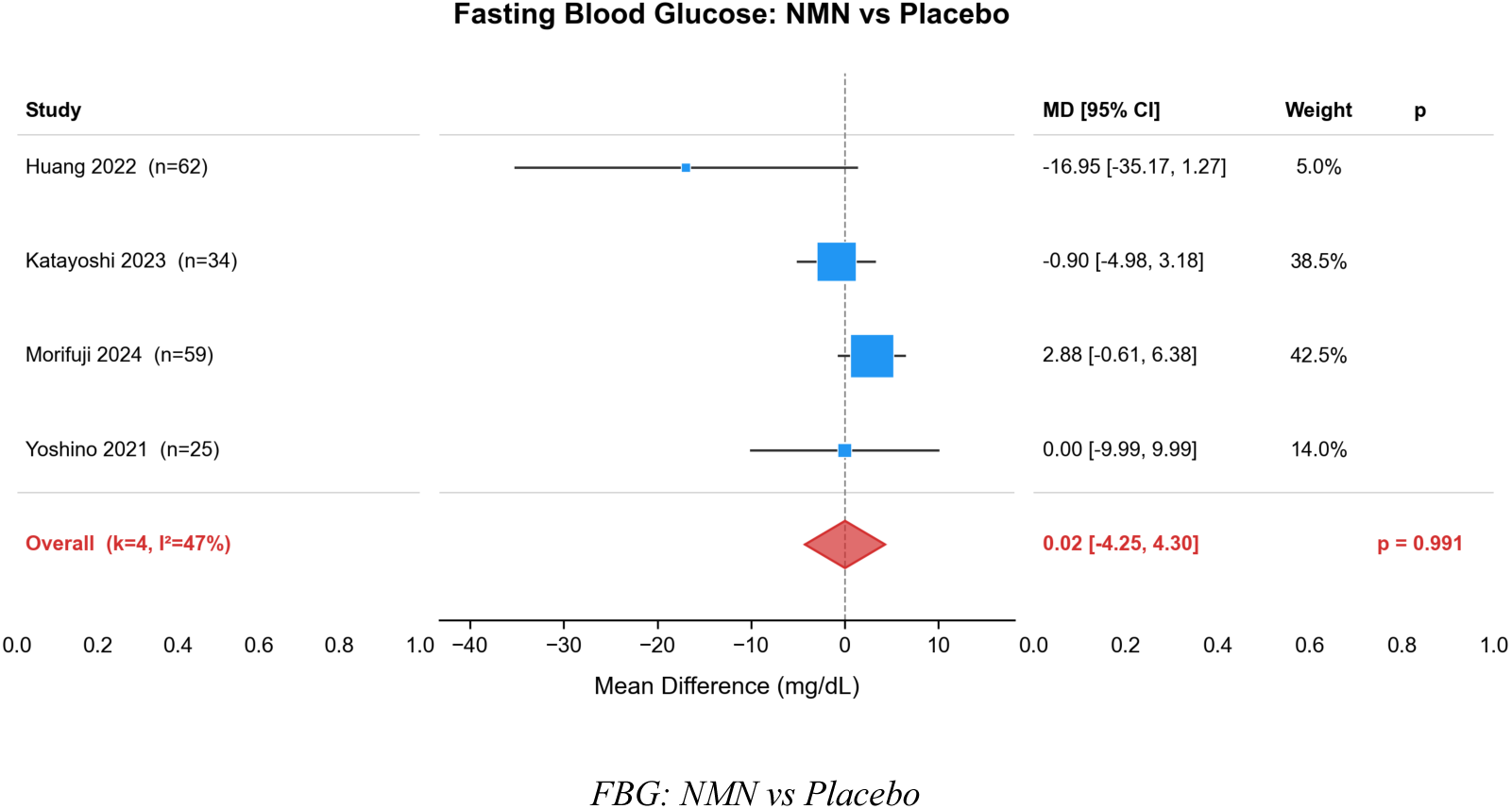
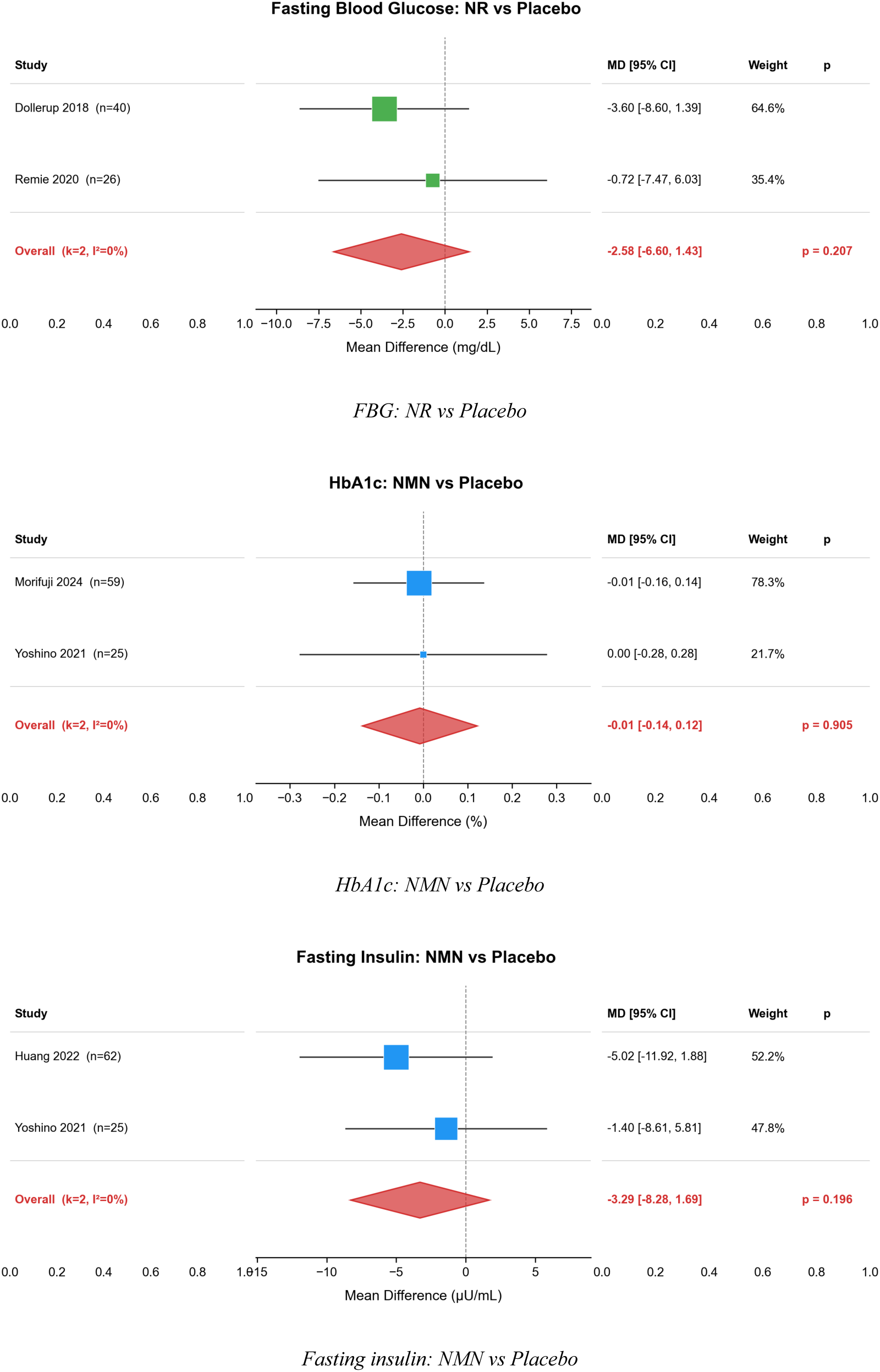
Pairwise forest plots for glycemic outcomes.

**Supplementary Figure S2.**
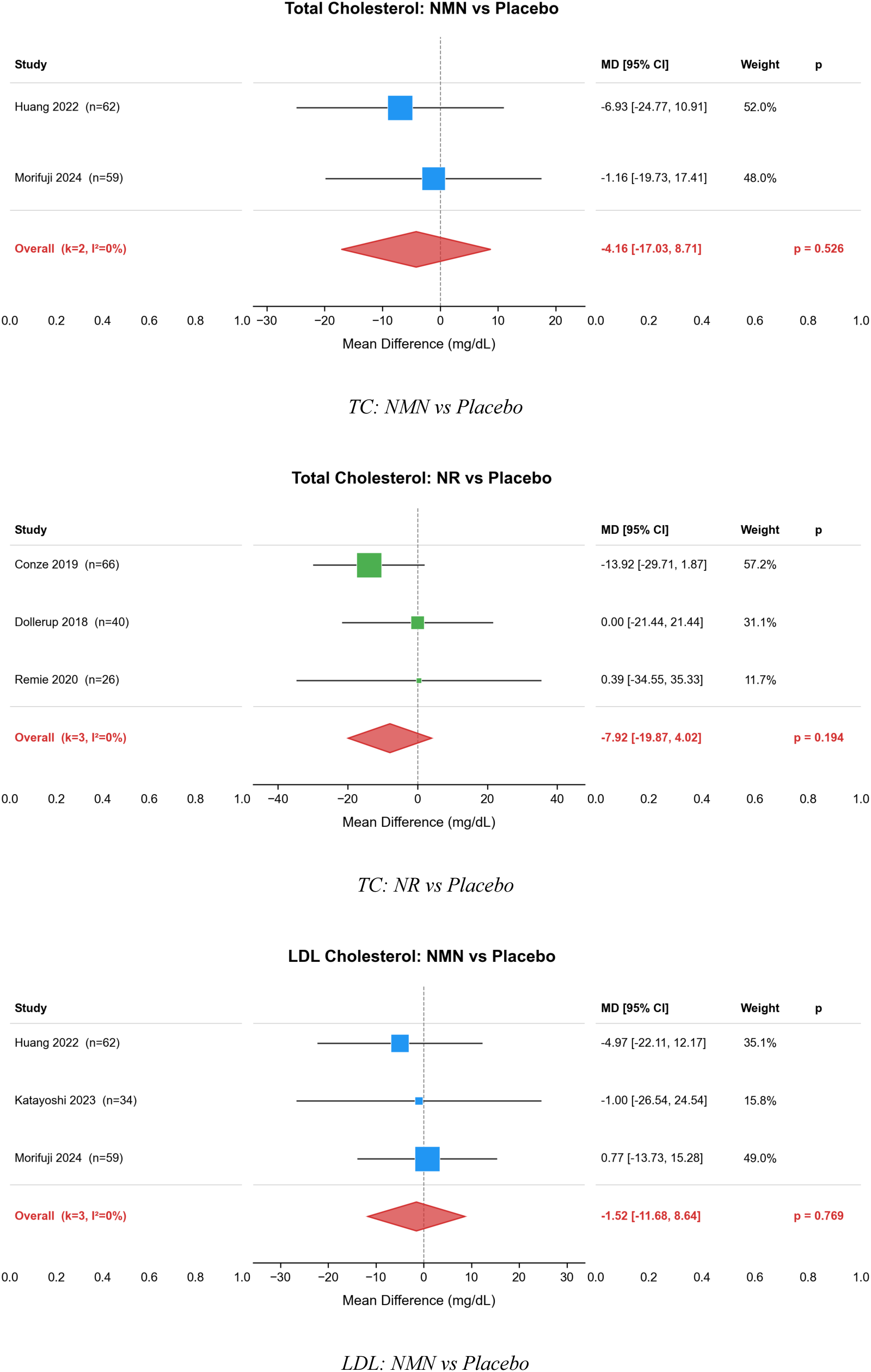
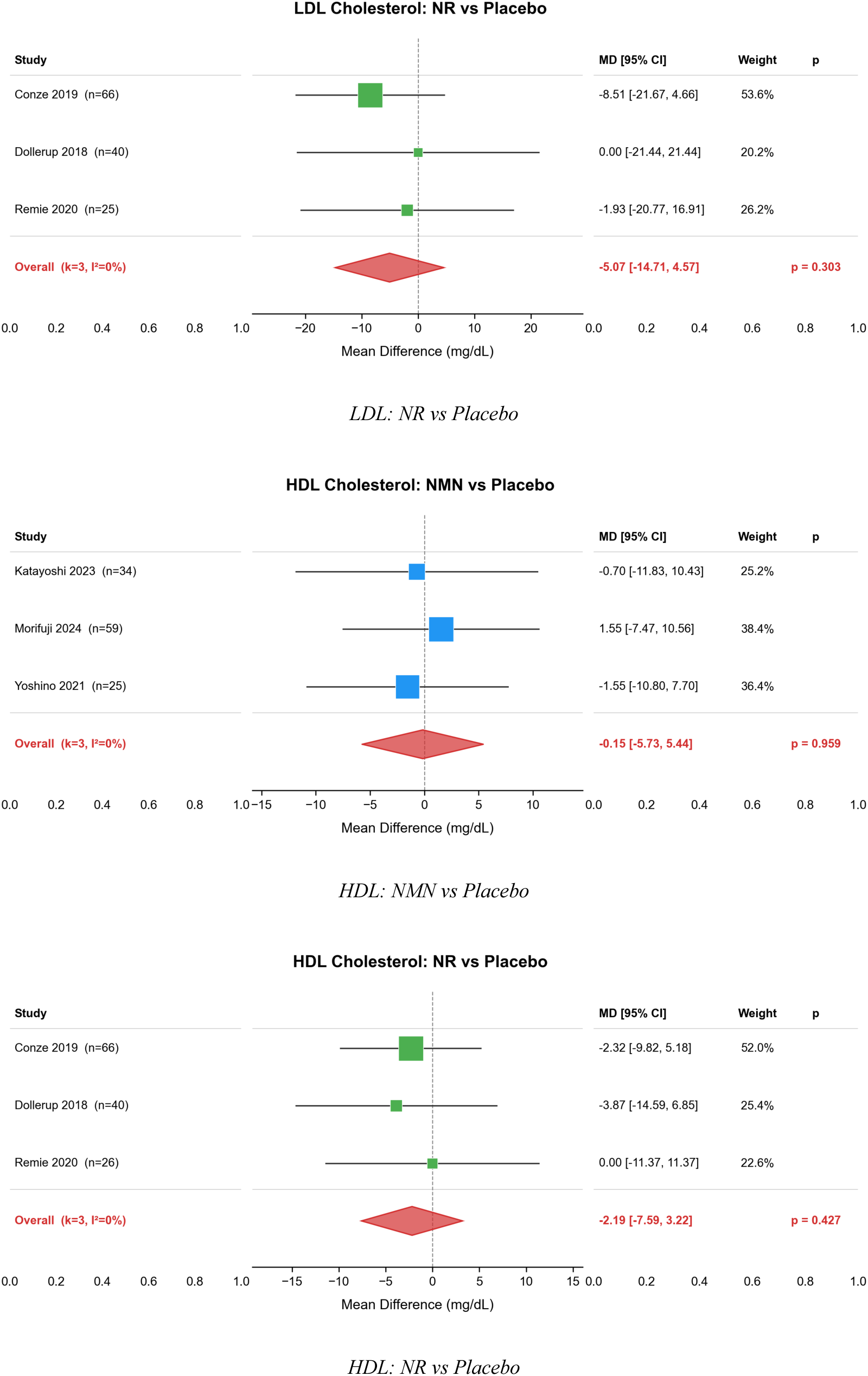
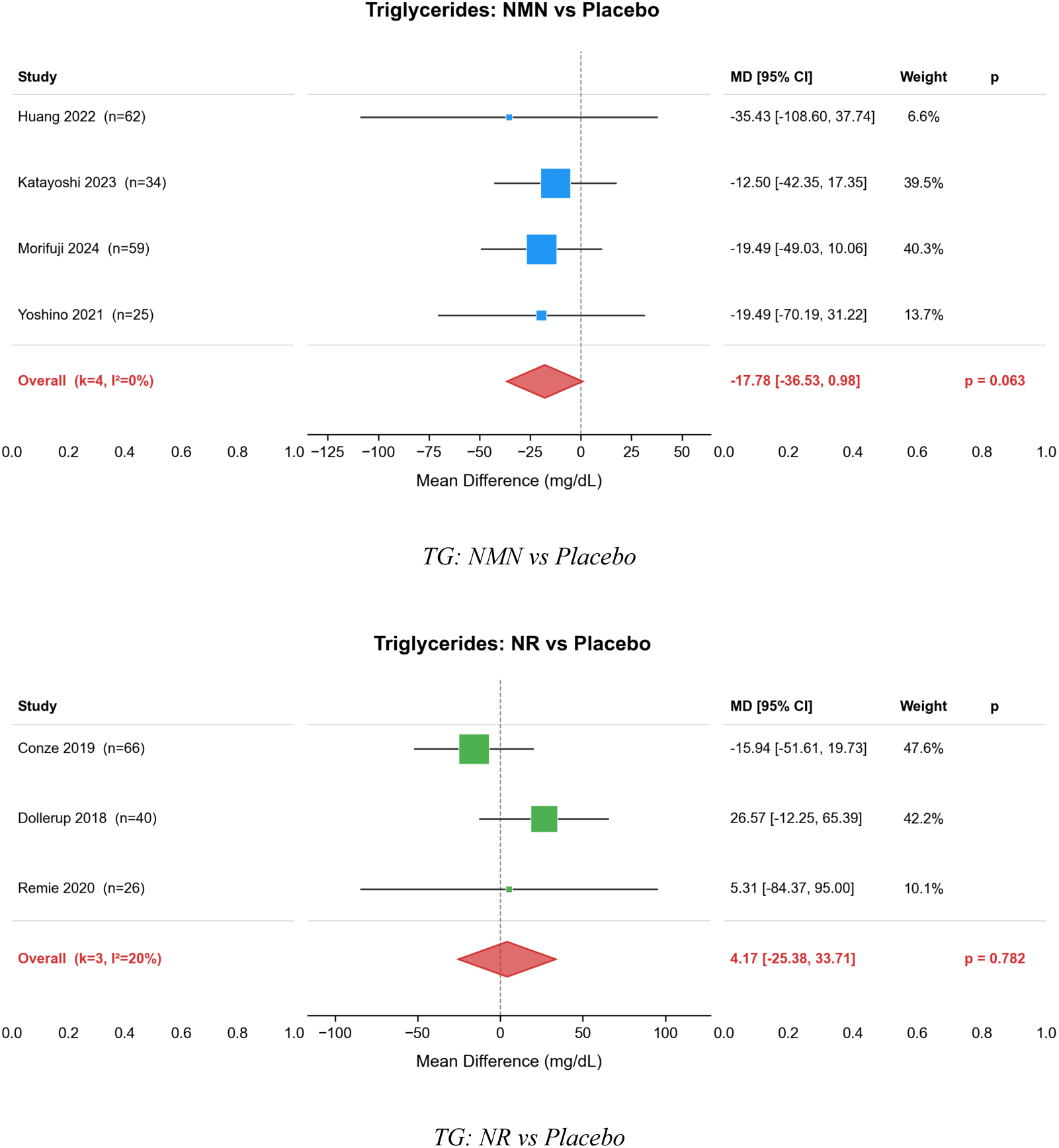
Pairwise forest plots for lipid outcomes.

**Supplementary Figure S3.**
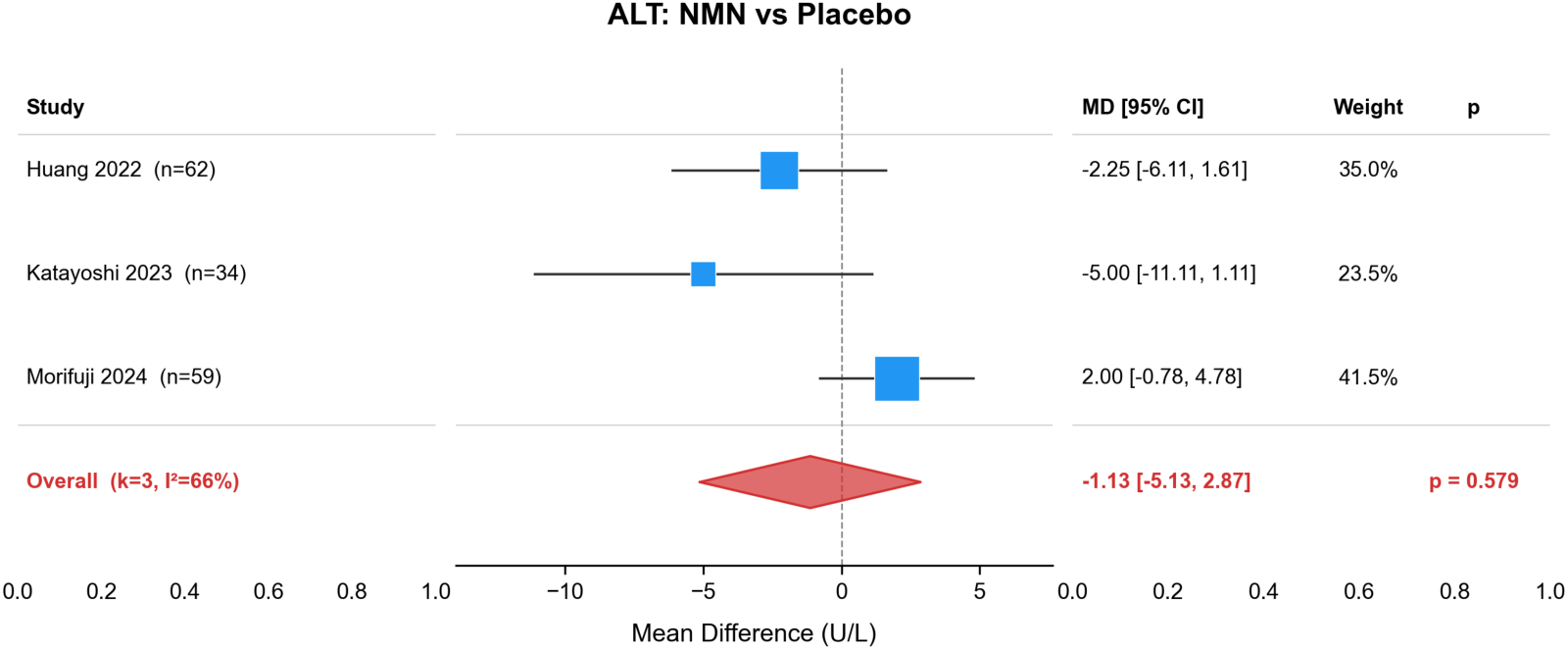
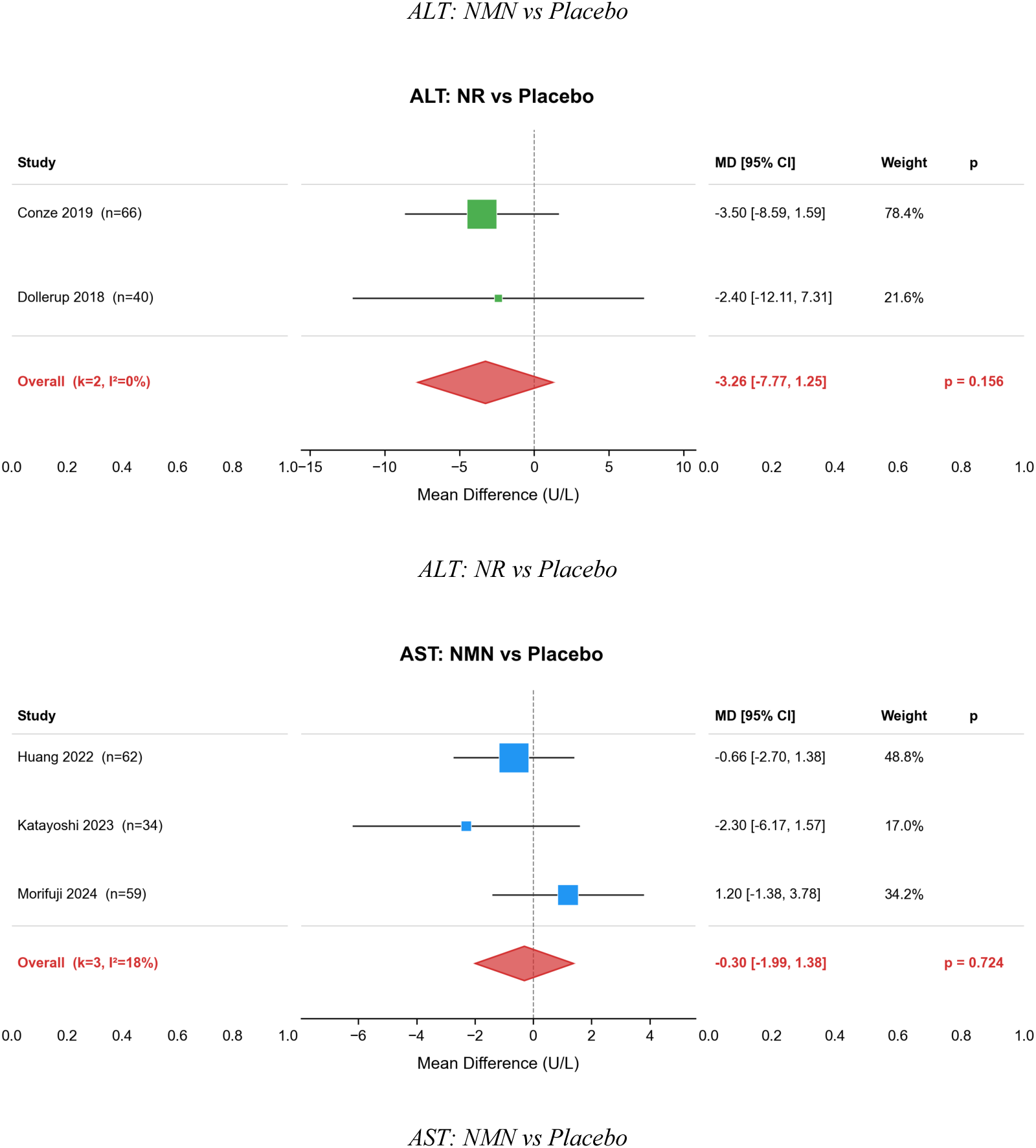
Pairwise forest plots for hepatic enzyme outcomes.

**Supplementary Figure S4.**
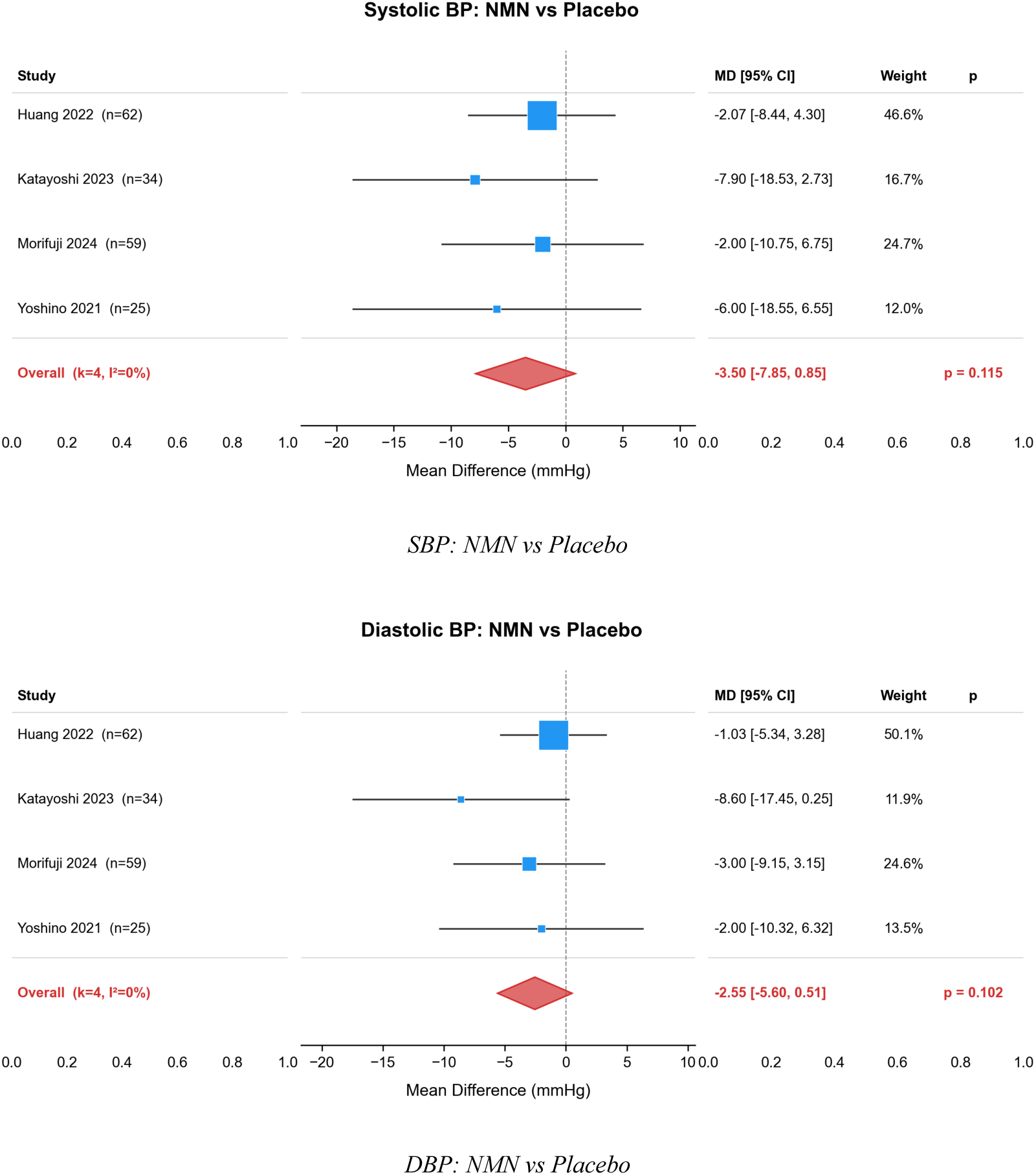
Pairwise forest plots for blood pressure outcomes.

**Supplementary Figure S5.**
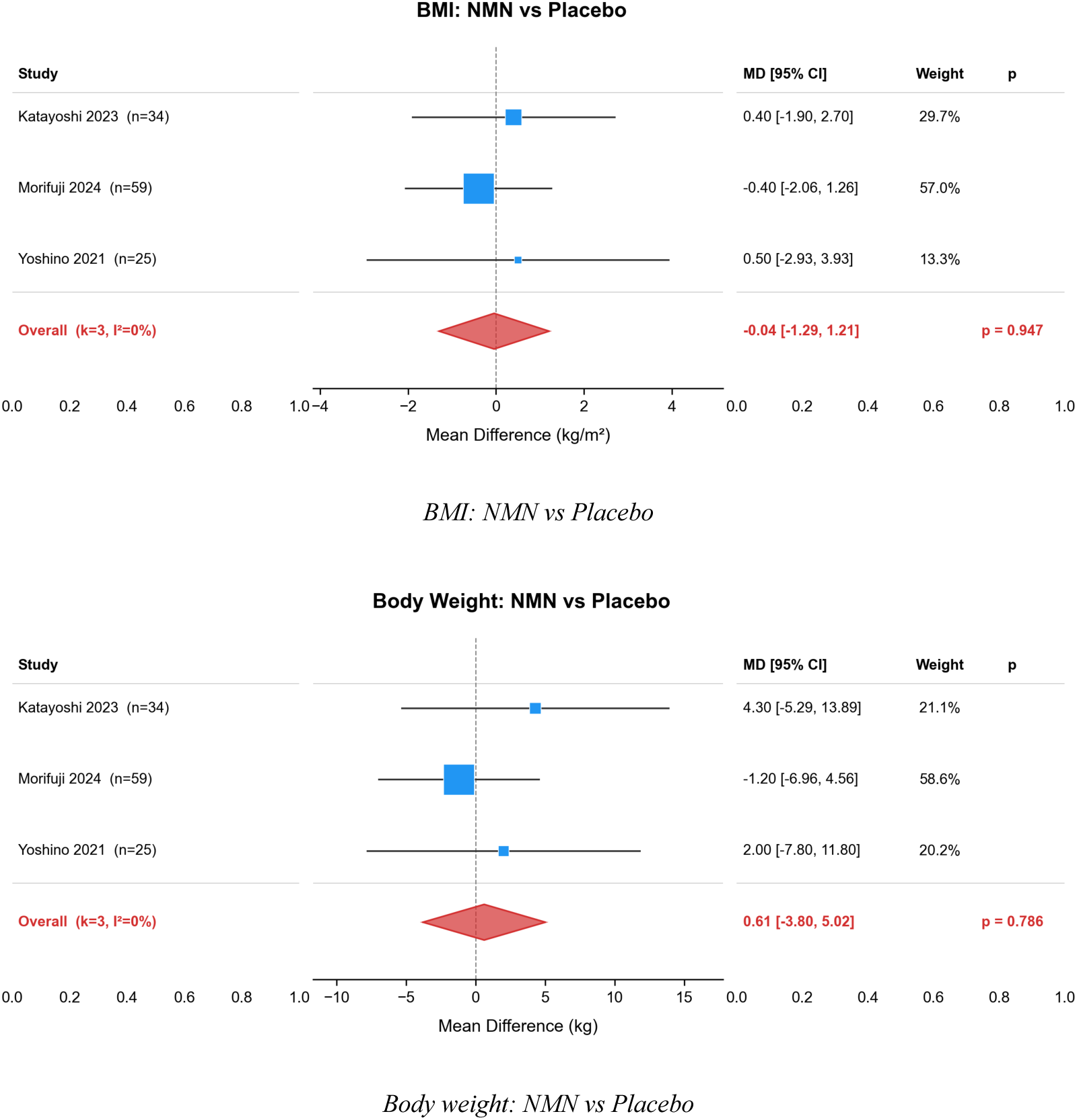
Pairwise forest plots for anthropometric outcomes.

**Supplementary Figure S6.**
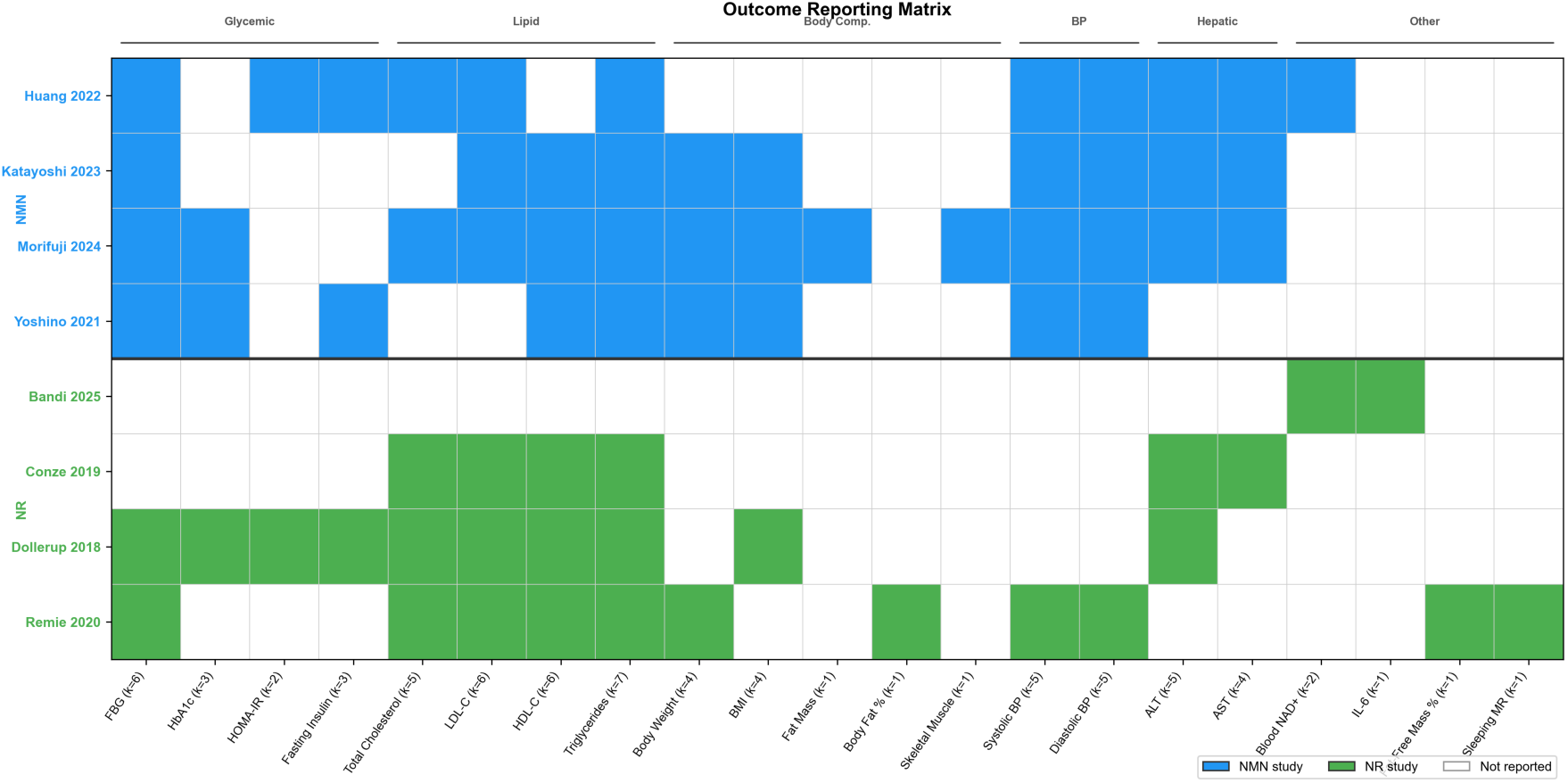
Outcome reporting matrix showing data availability for each metabolic outcome across the 8 studies contributing to quantitative synthesis.

**Supplementary Figure S7.**
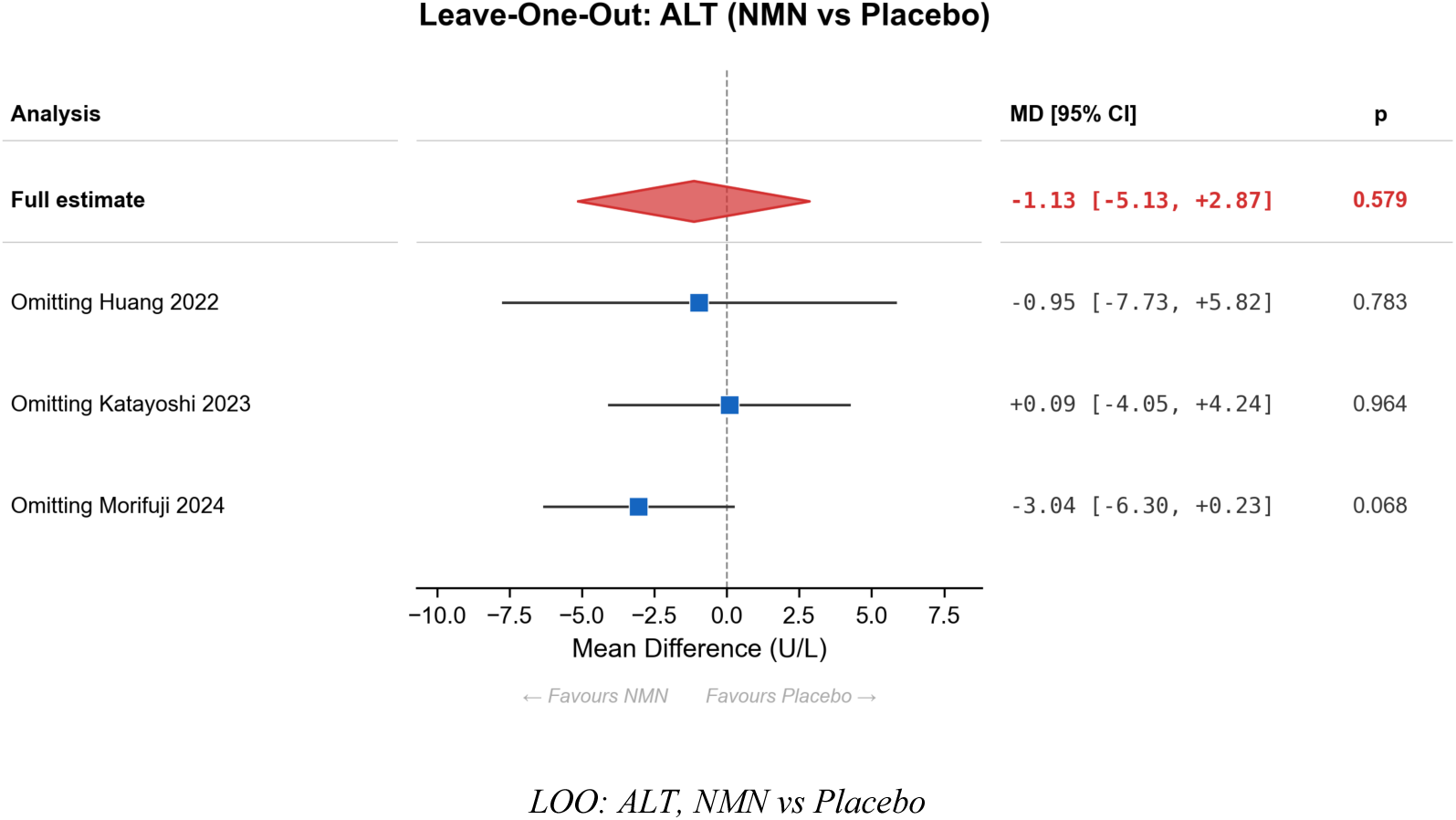
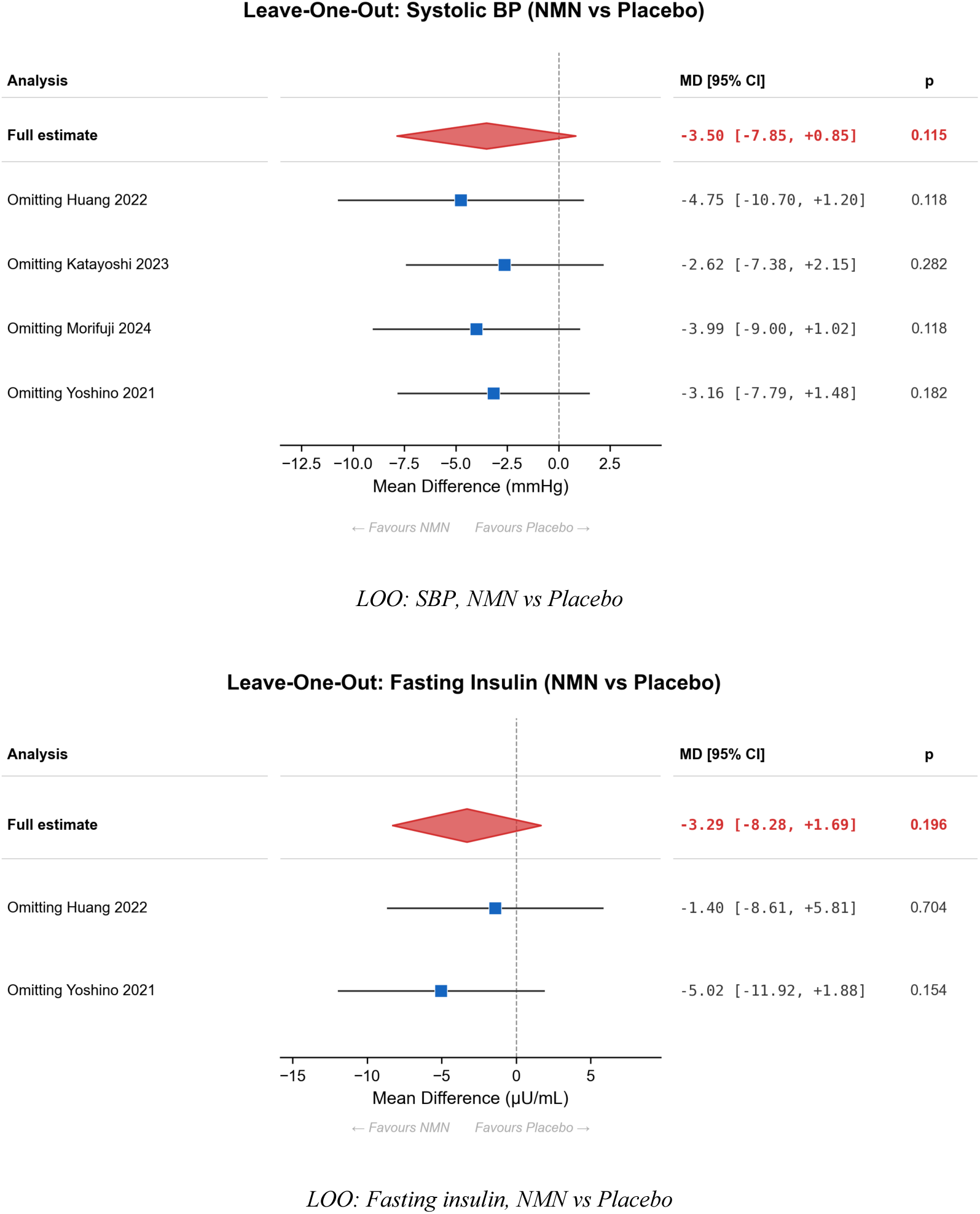
Leave-one-out sensitivity forest plots for selected pairwise comparisons (ALT, SBP, fasting insulin) presented as representative robustness checks; no pairwise significance changes were observed in the current analysis.

**Supplementary Table S1.**
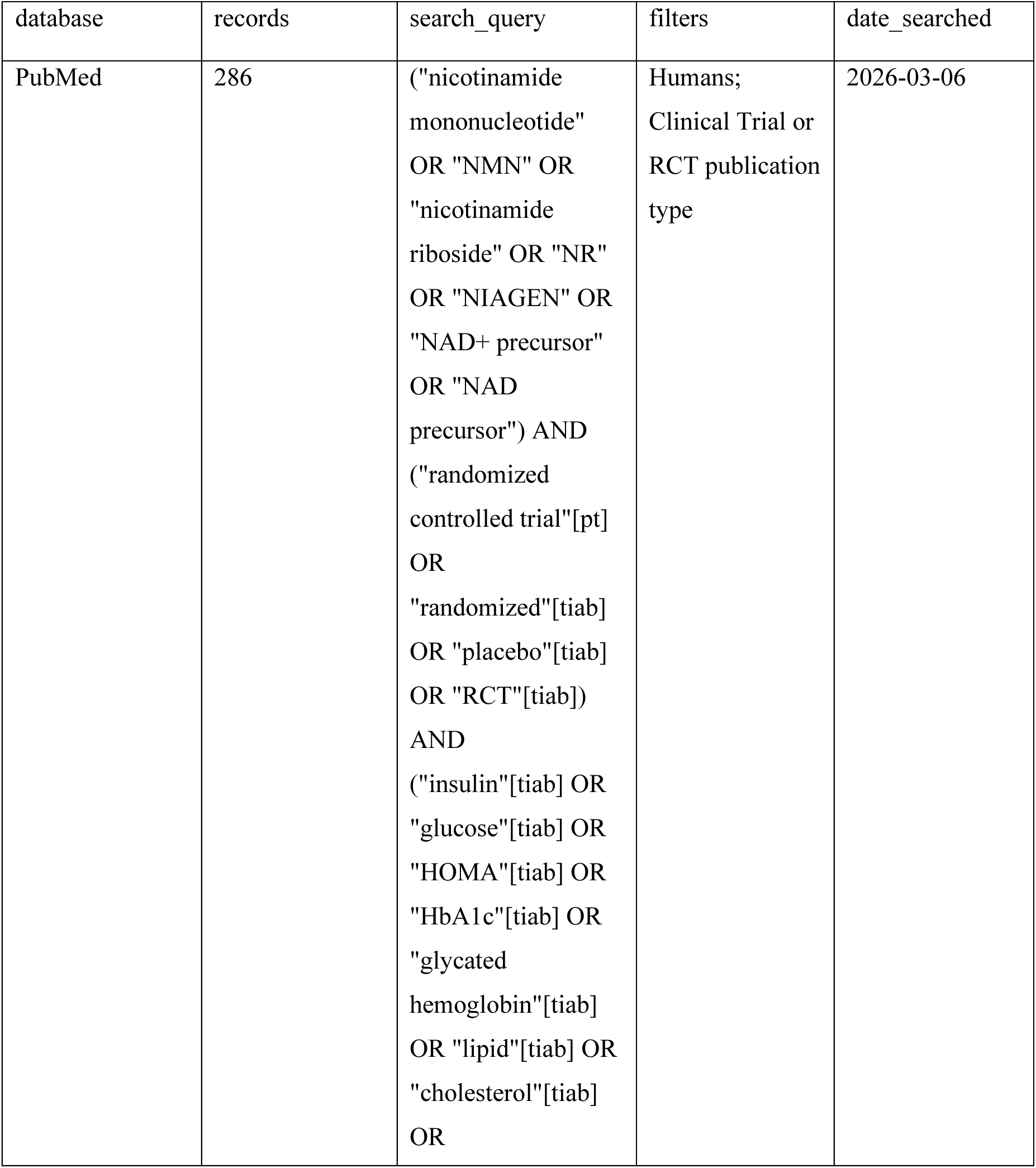
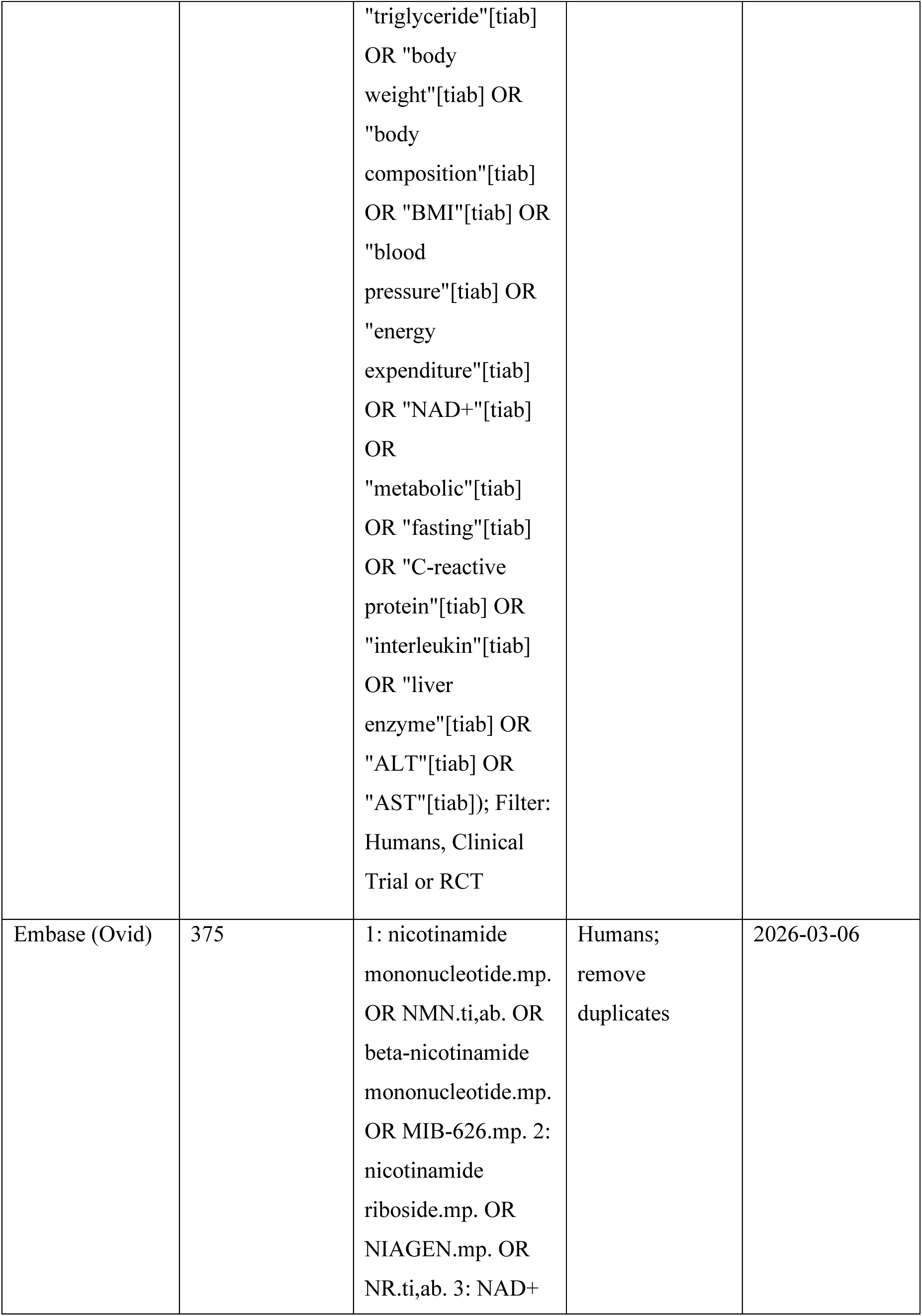
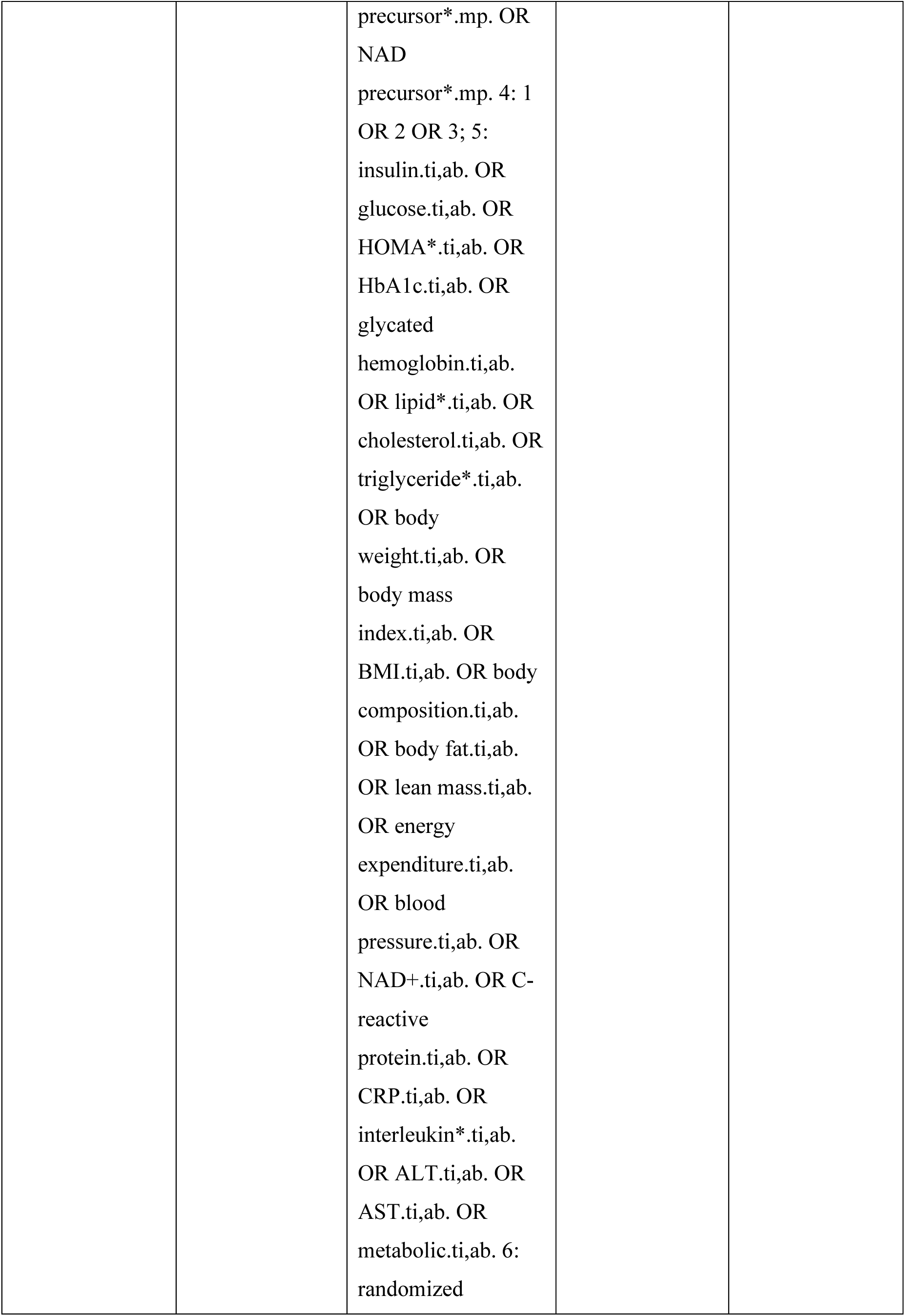
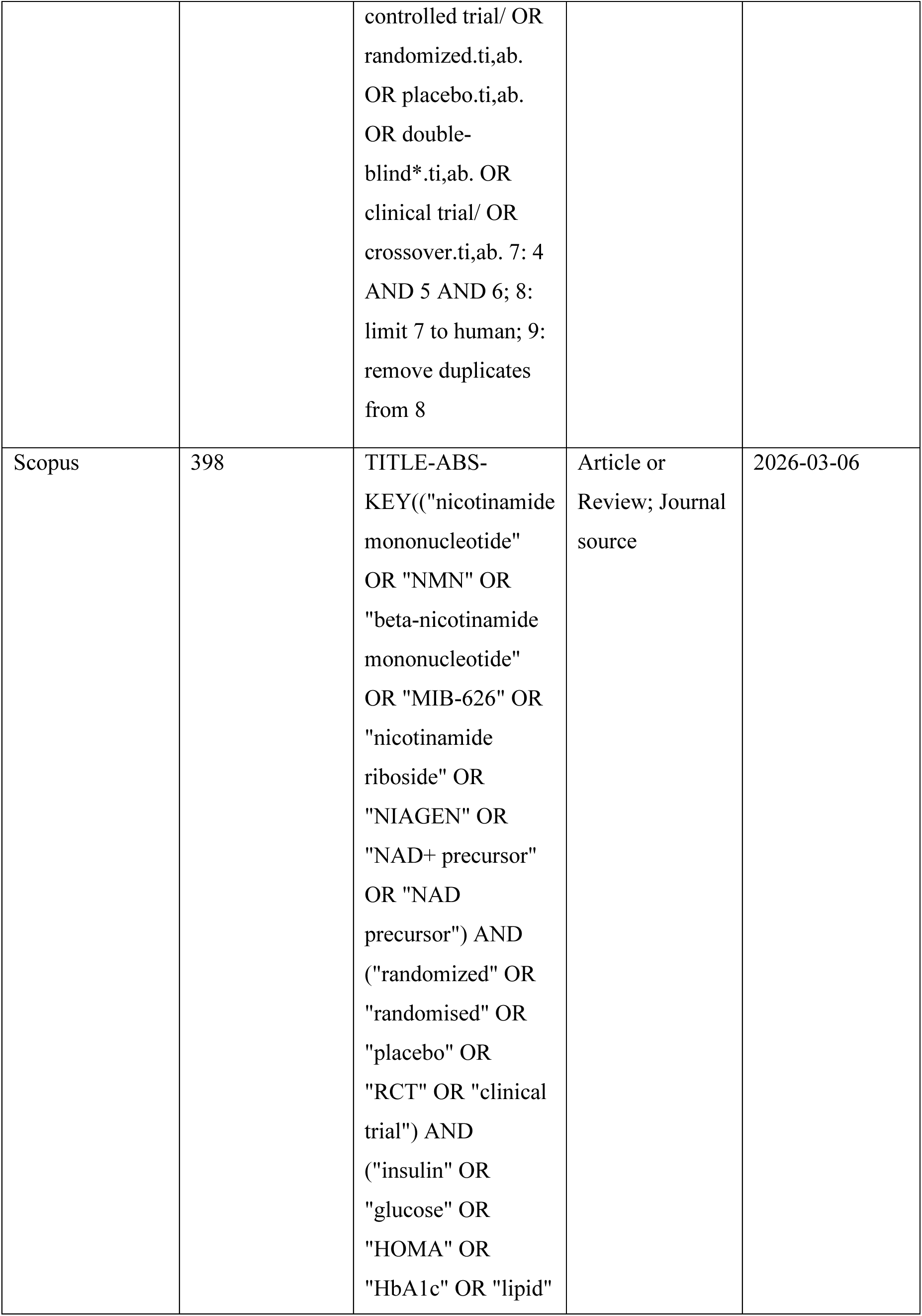
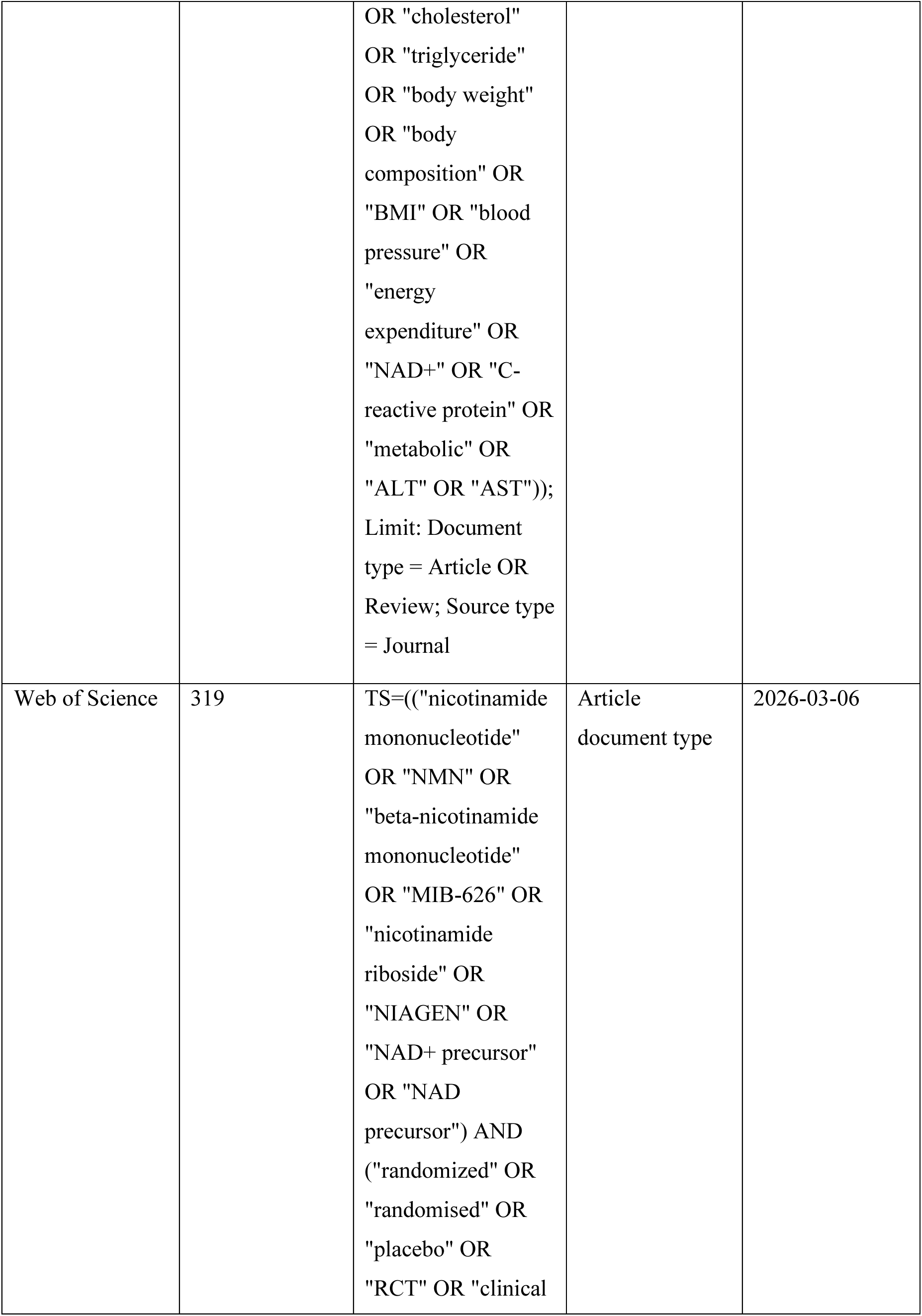
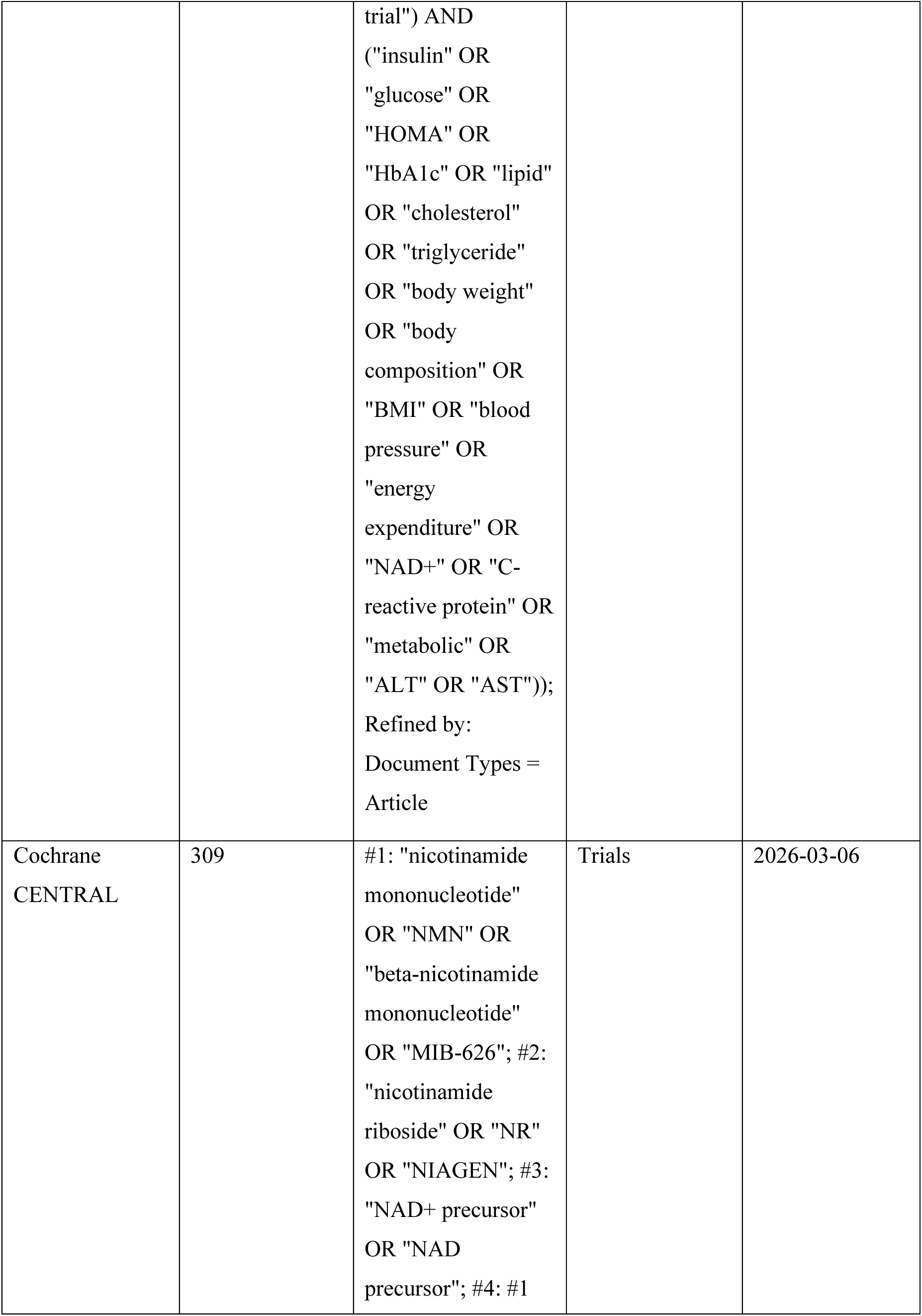
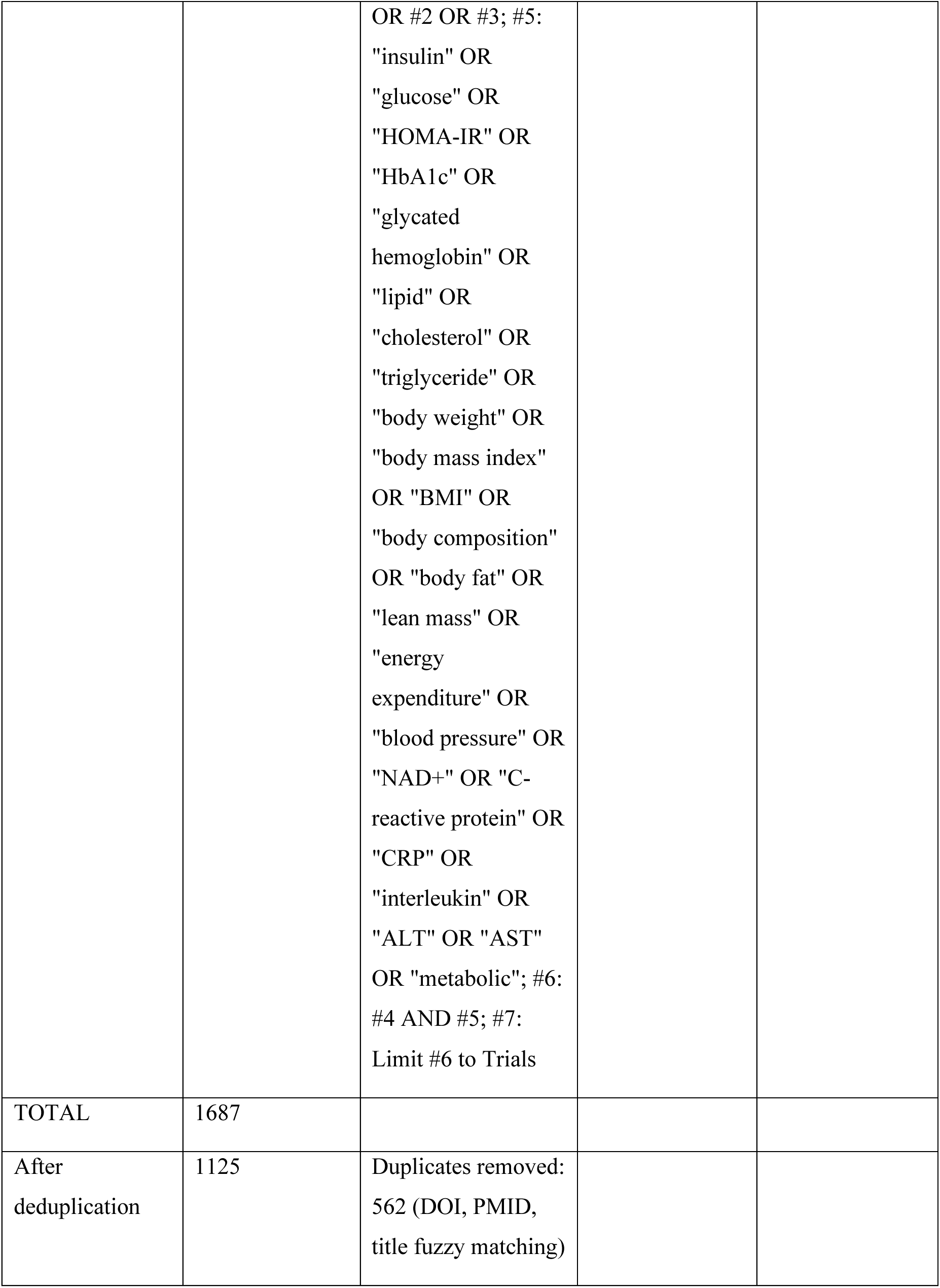
Complete search strategy for all five databases.

**Supplementary Table S2.**
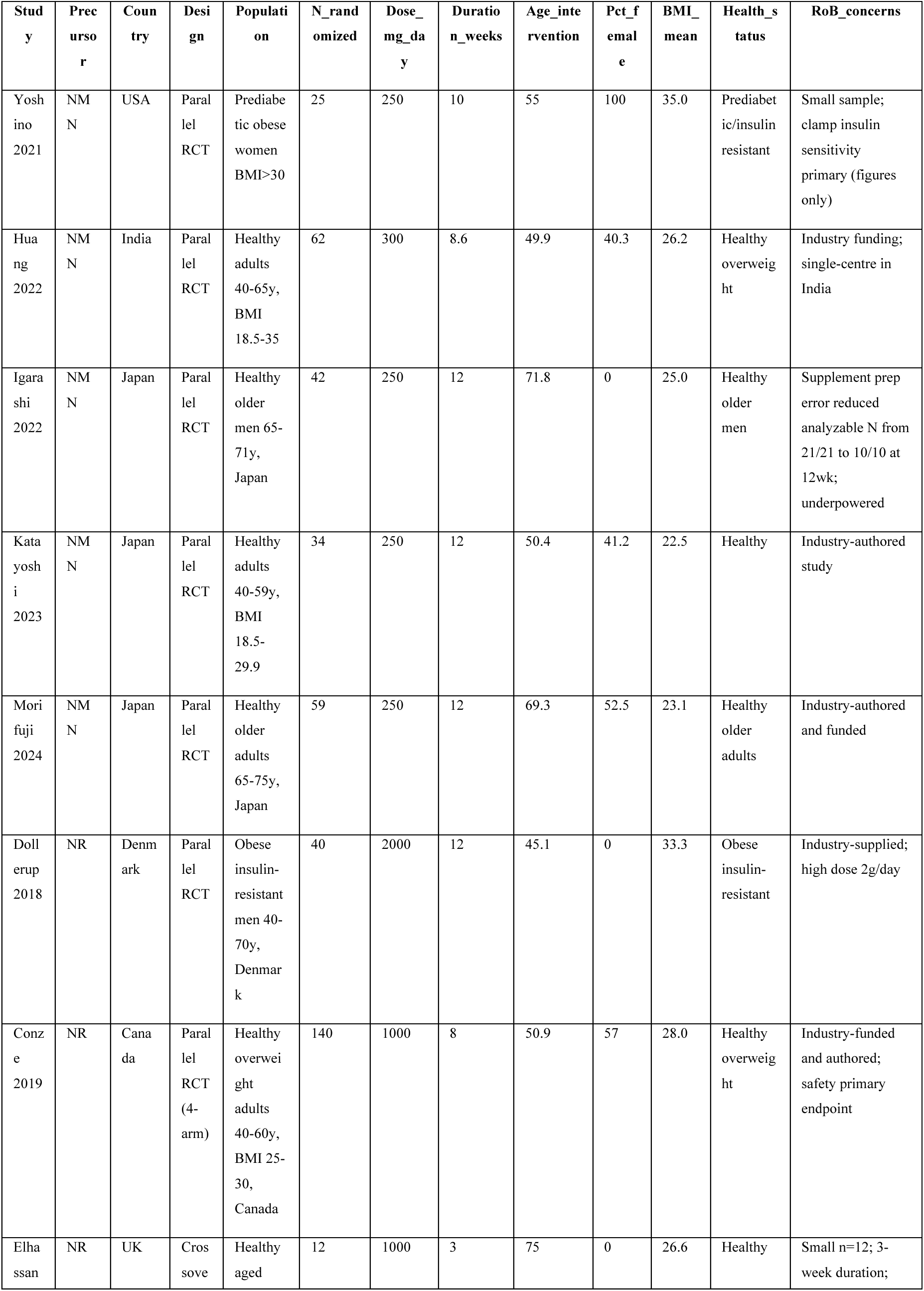
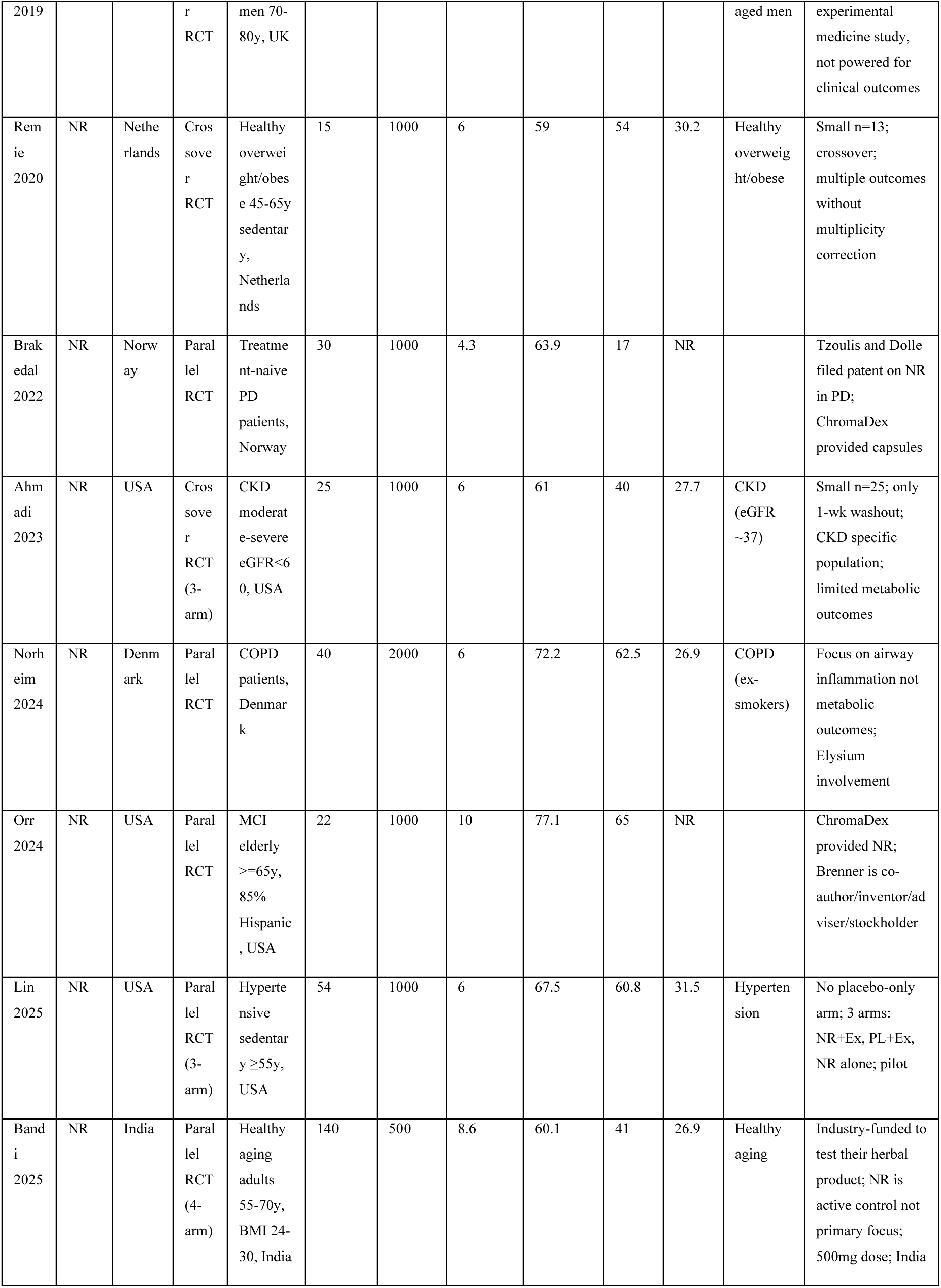
Detailed characteristics of included studies.

**Supplementary Table S3.**
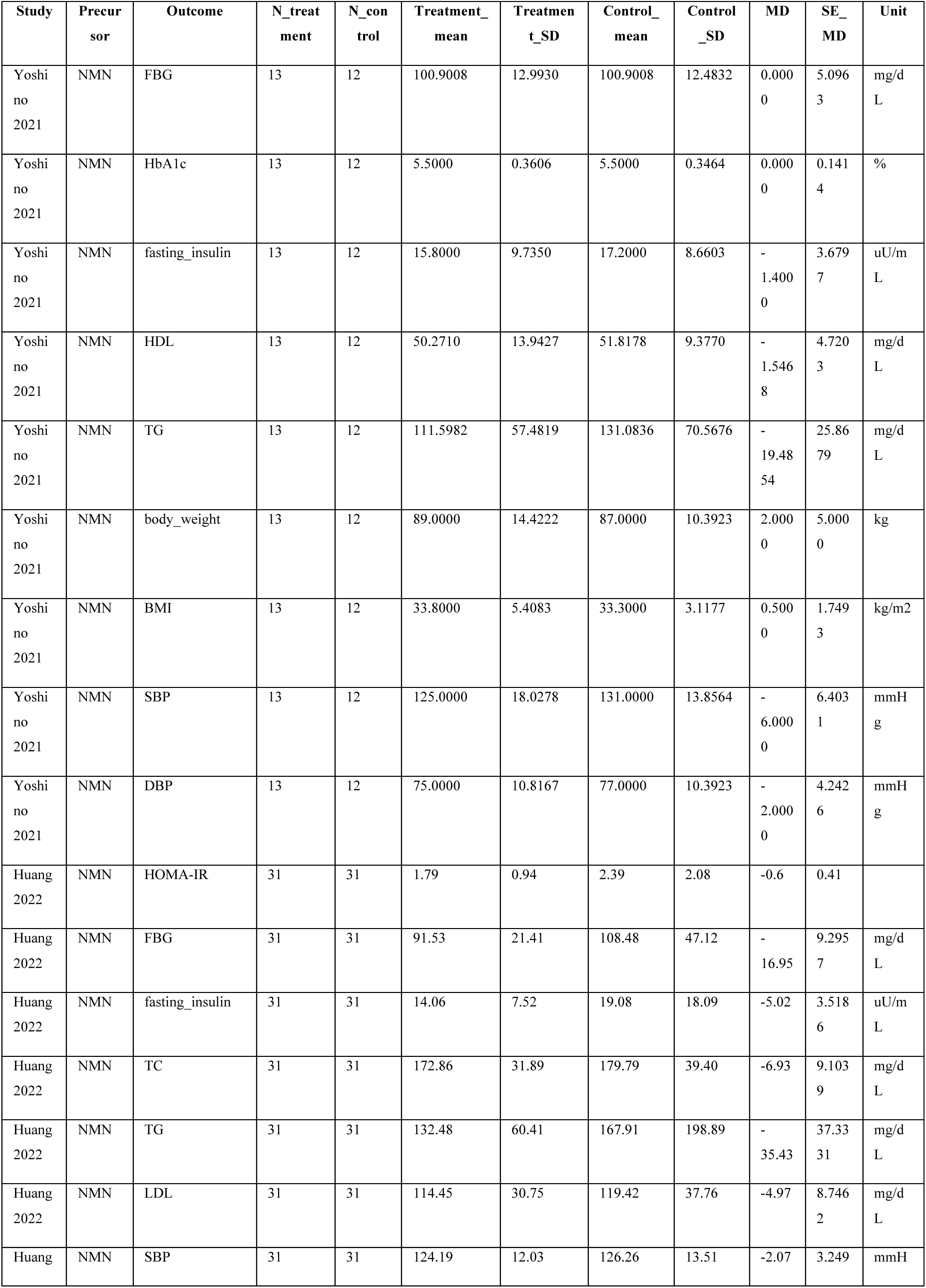
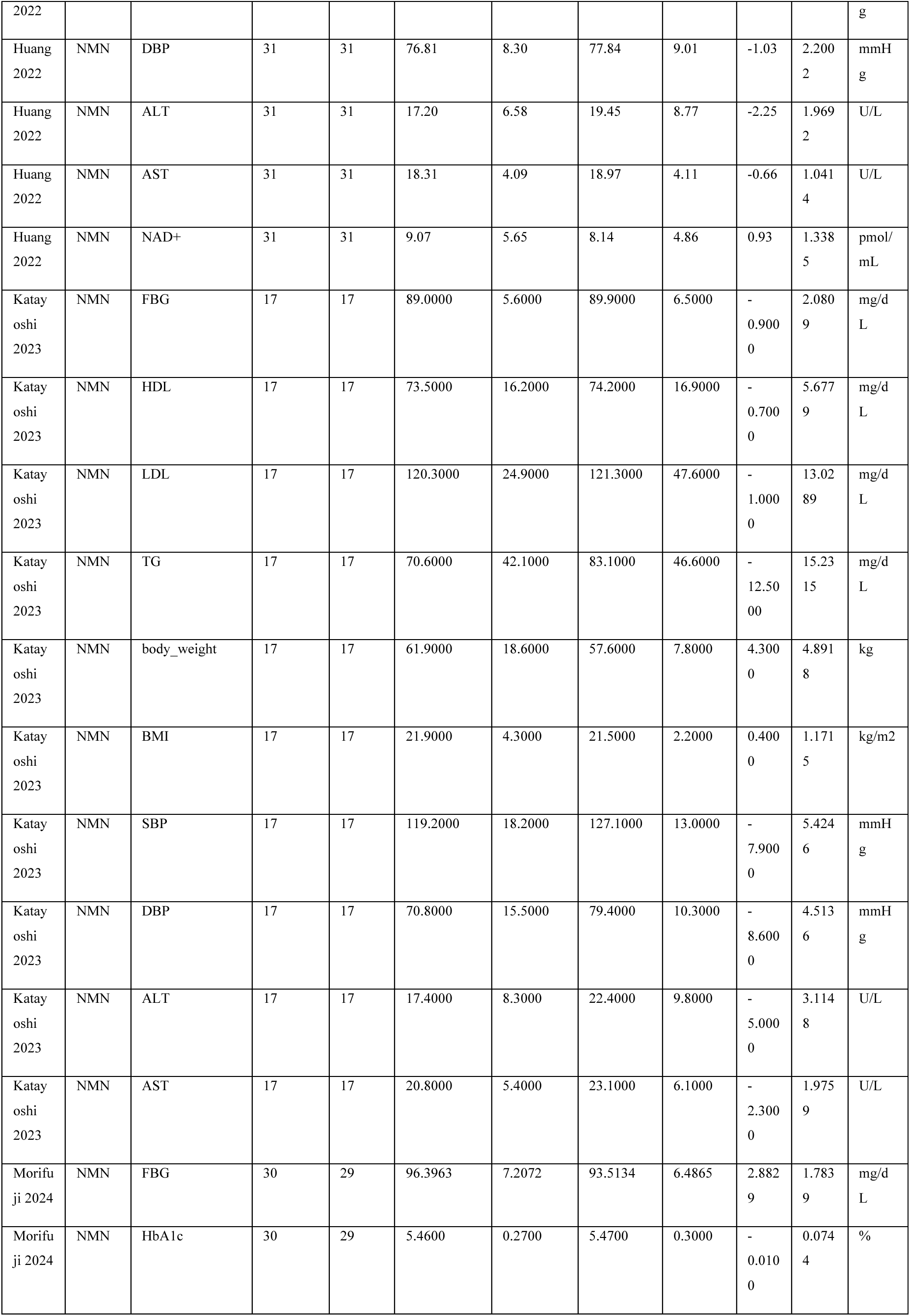
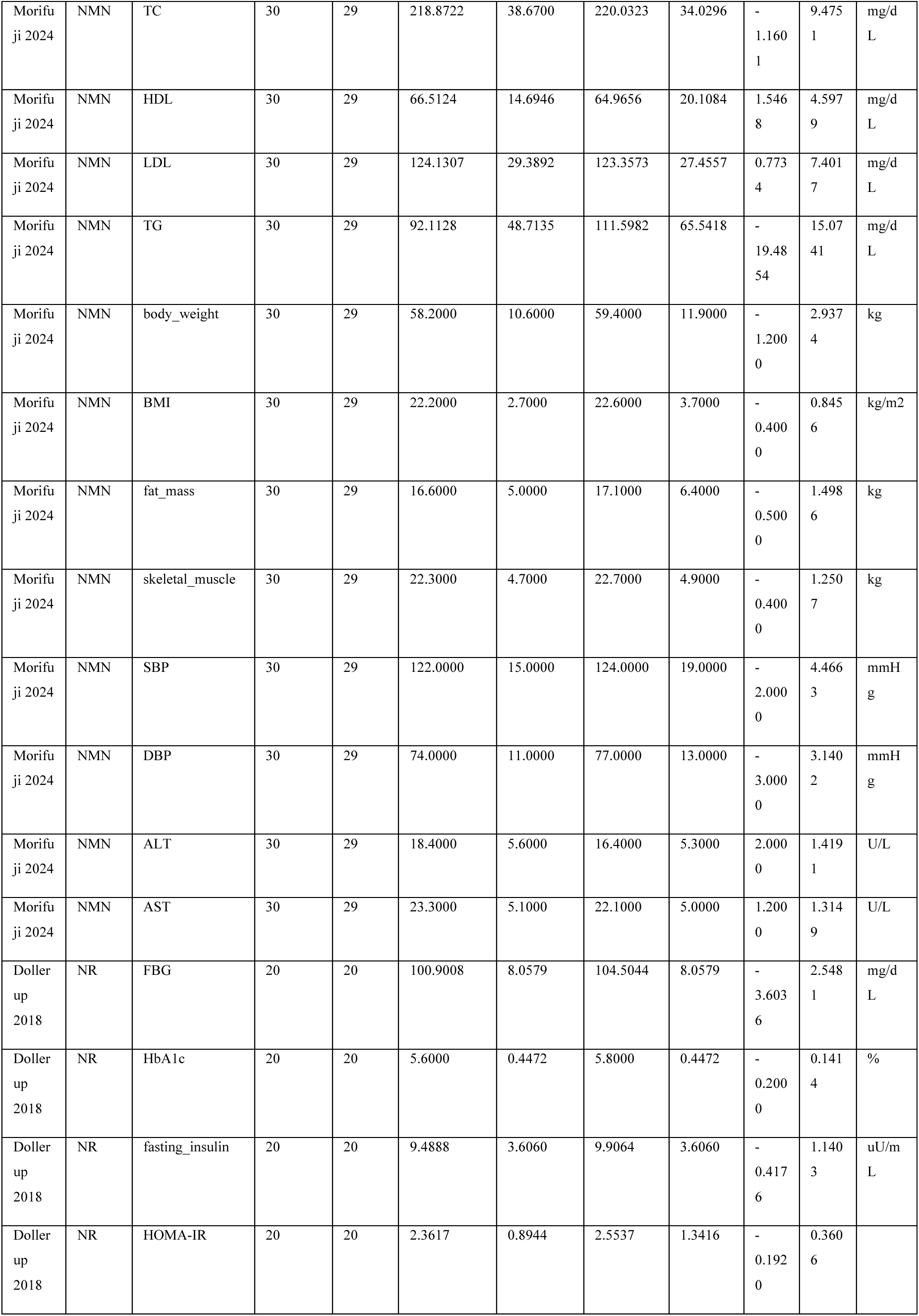
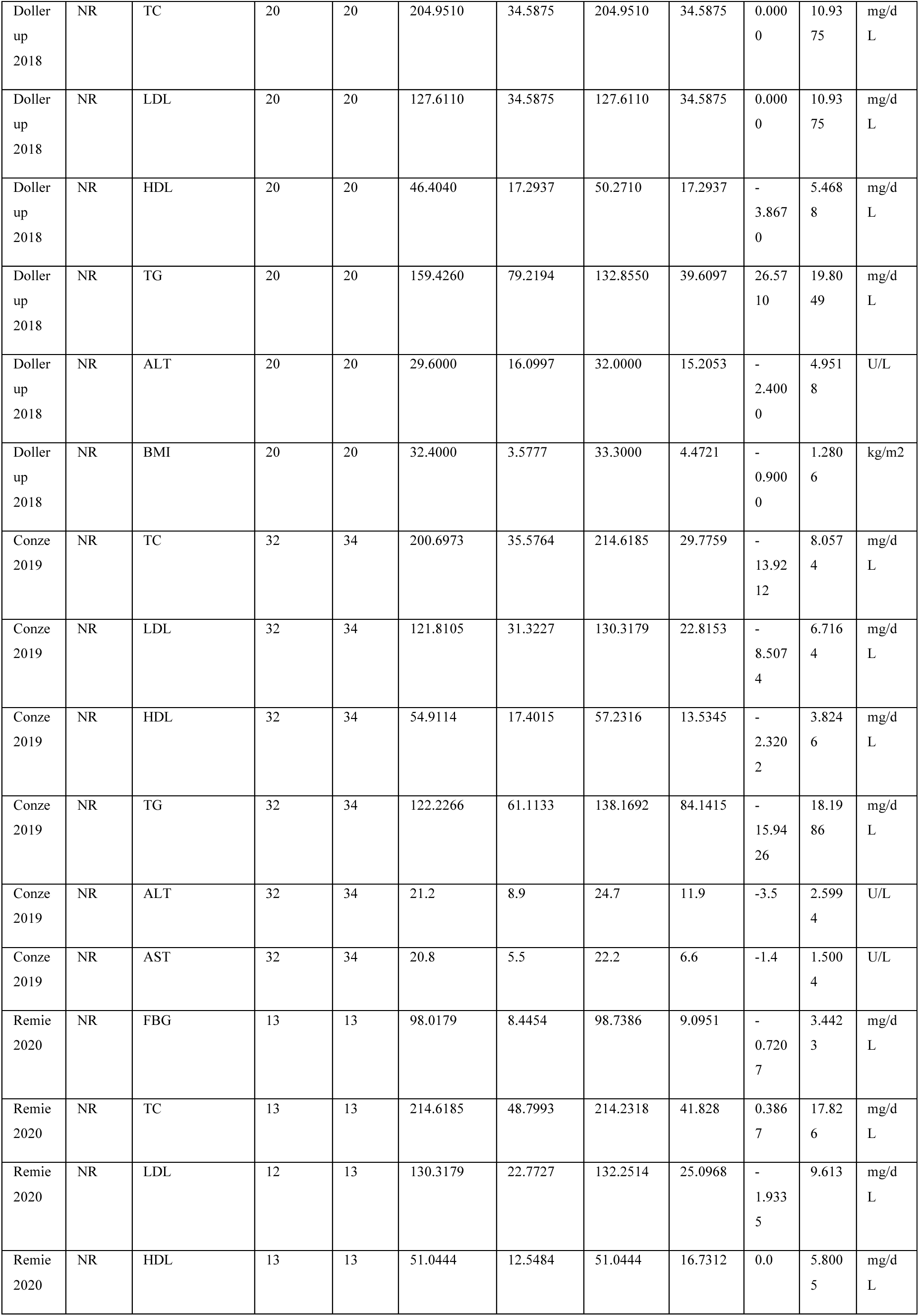
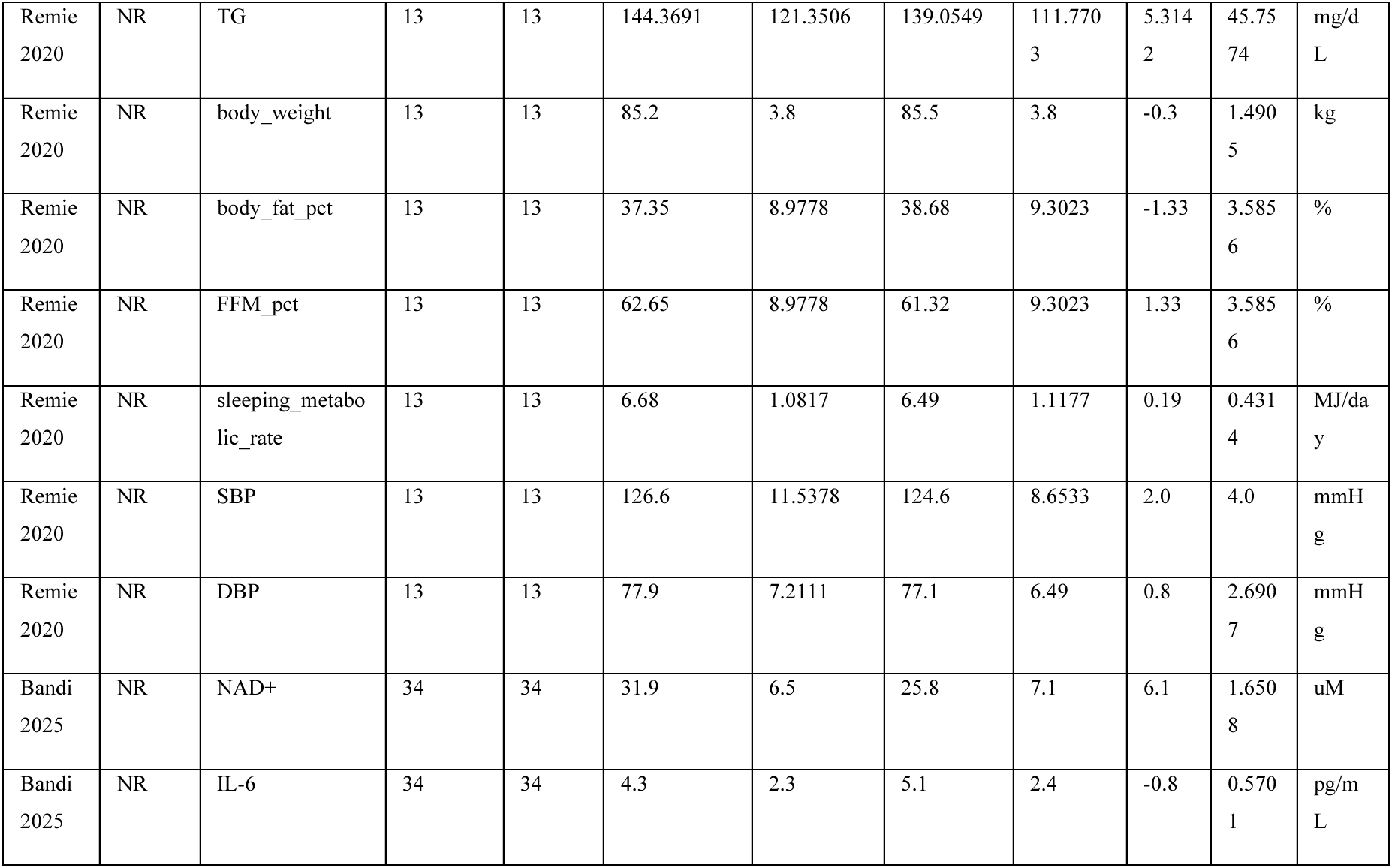
Full data extraction table with all outcome data points.

**Supplementary Table S4.** Risk of bias (RoB 2) domain-level assessments.

| Study | D1_Randomization | D2_Deviations | D3_Missing_data | D4_Measurement | D5_Reporting | Overall |
| --- | --- | --- | --- | --- | --- | --- |
| Yoshino 2021 | Some concerns | Low | Low | Low | Some concerns | Some concerns |
| Huang 2022 | Some concerns | Low | Some concerns | Low | Some concerns | Some concerns |
| Igarashi 2022 | Low | Some concerns | High | Some concerns | Some concerns | High |
| Katayoshi 2023 | Low | Low | Low | Low | Some concerns | Some concerns |
| Morifuji 2024 | Some concerns | Low | Low | Low | Some concerns | Some concerns |
| Dollerup 2018 | Low | Low | Low | Low | Some concerns | Some concerns |
| Conze 2019 | Some concerns | Low | Low | Low | Some concerns | Some concerns |
| Elhassan 2019 | Some concerns | Some concerns | Low | Some concerns | High | High |
| Remie 2020 | Low | Low | Some concerns | Low | Some concerns | Some concerns |
| Brakedal 2022 | Low | Low | Low | Low | Some concerns | Some concerns |
| Ahmadi 2023 | Some concerns | Some concerns | Low | Some concerns | Some concerns | Some concerns |
| Norheim 2024 | Low | Low | Low | Low | Some concerns | Some concerns |
| Orr 2024 | Some concerns | Low | Some concerns | Some concerns | Some concerns | Some concerns |
| Lin 2025 | Some concerns | Low | Low | Low | Some concerns | Some concerns |
| Bandi 2025 | Low | Low | Some concerns | Low | Some concerns | Some concerns |

**Supplementary Table S5.**
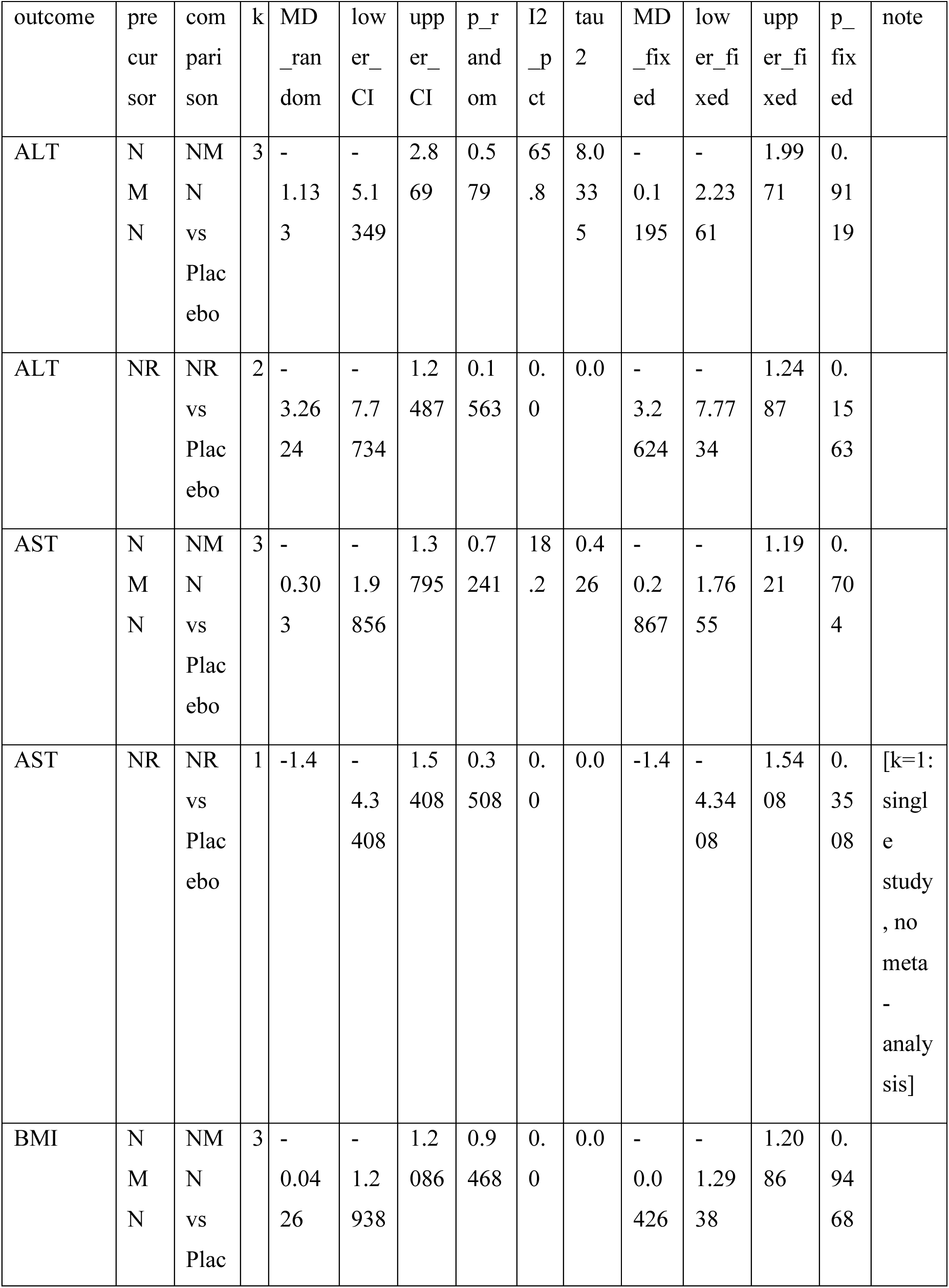
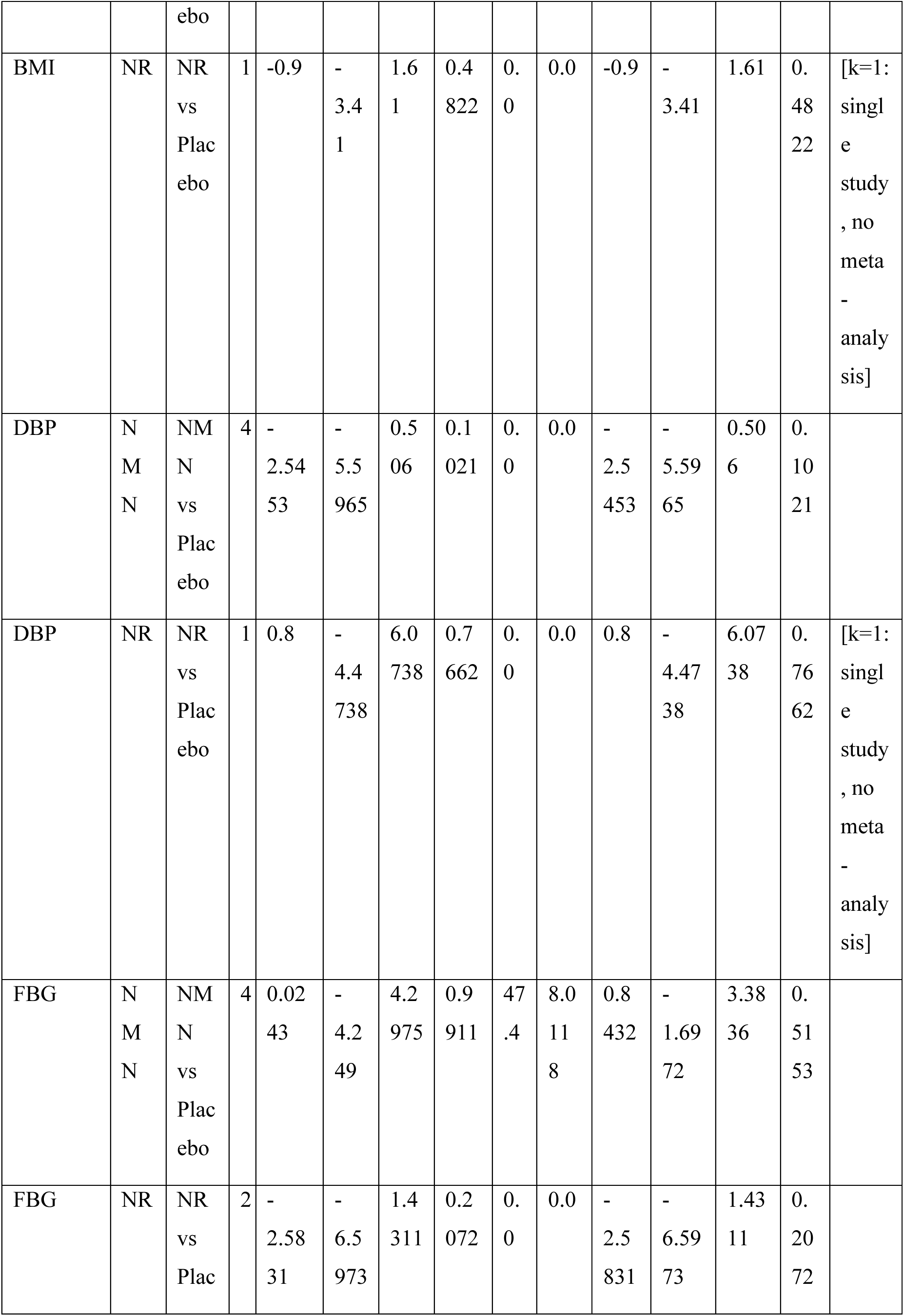
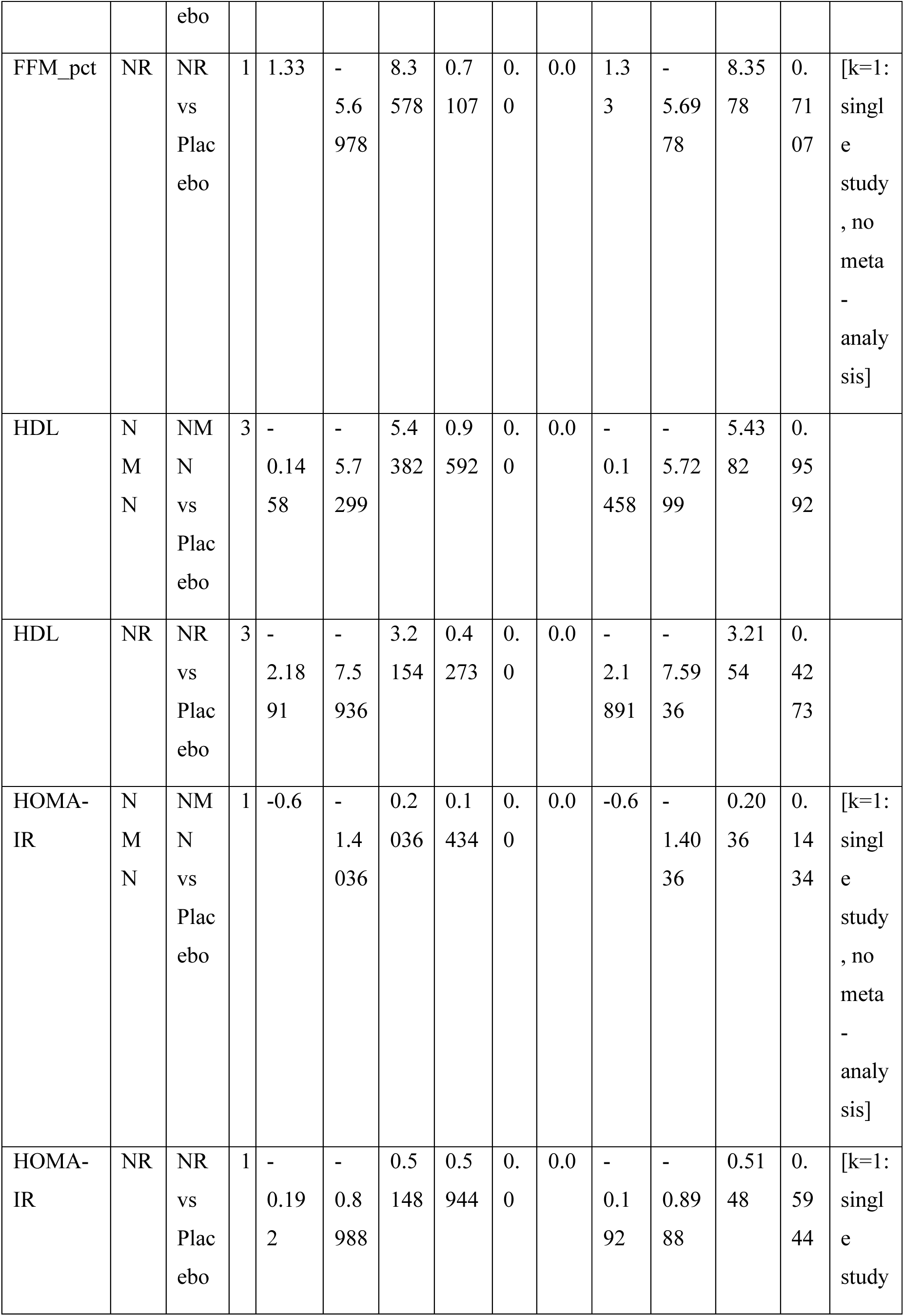
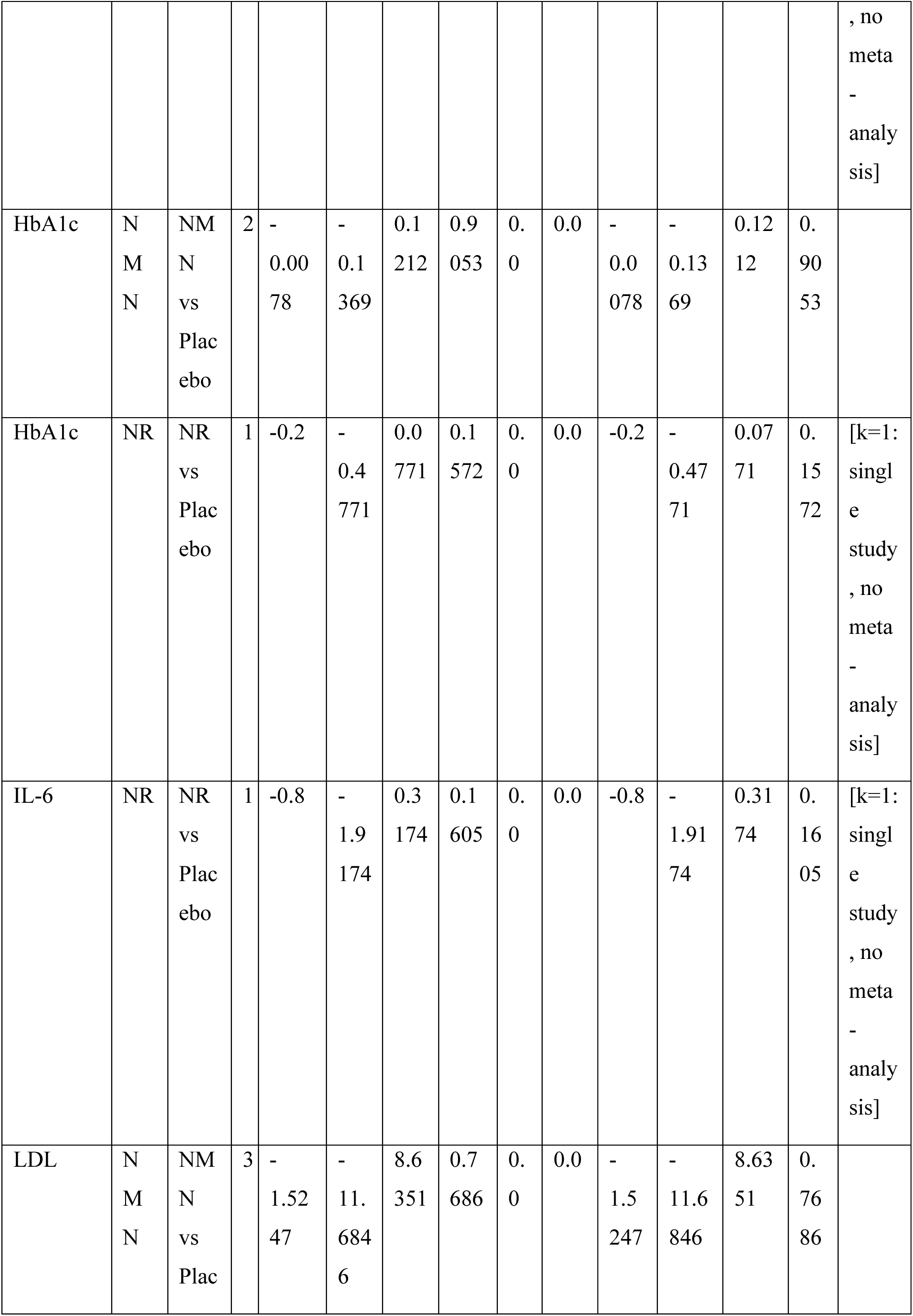
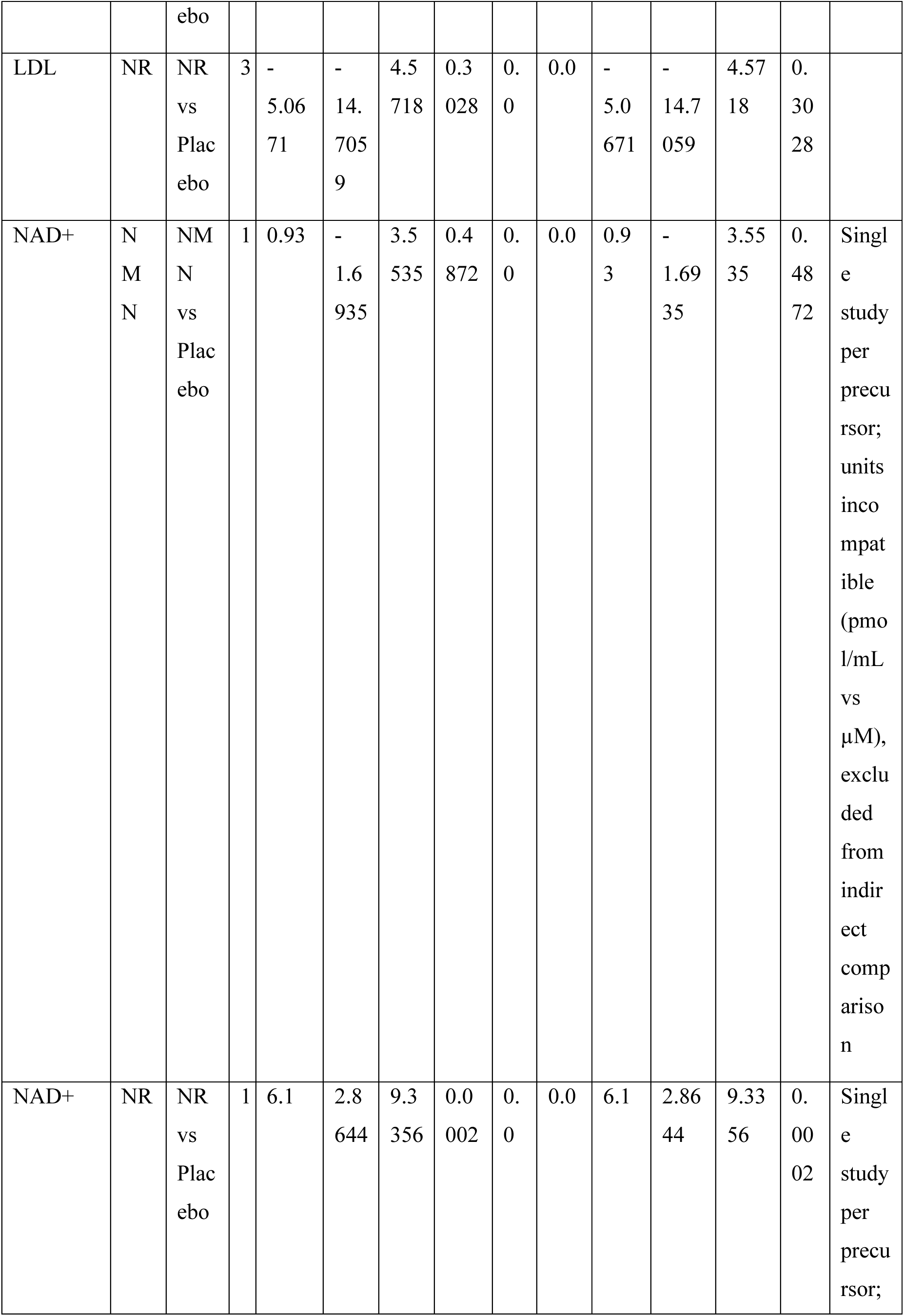
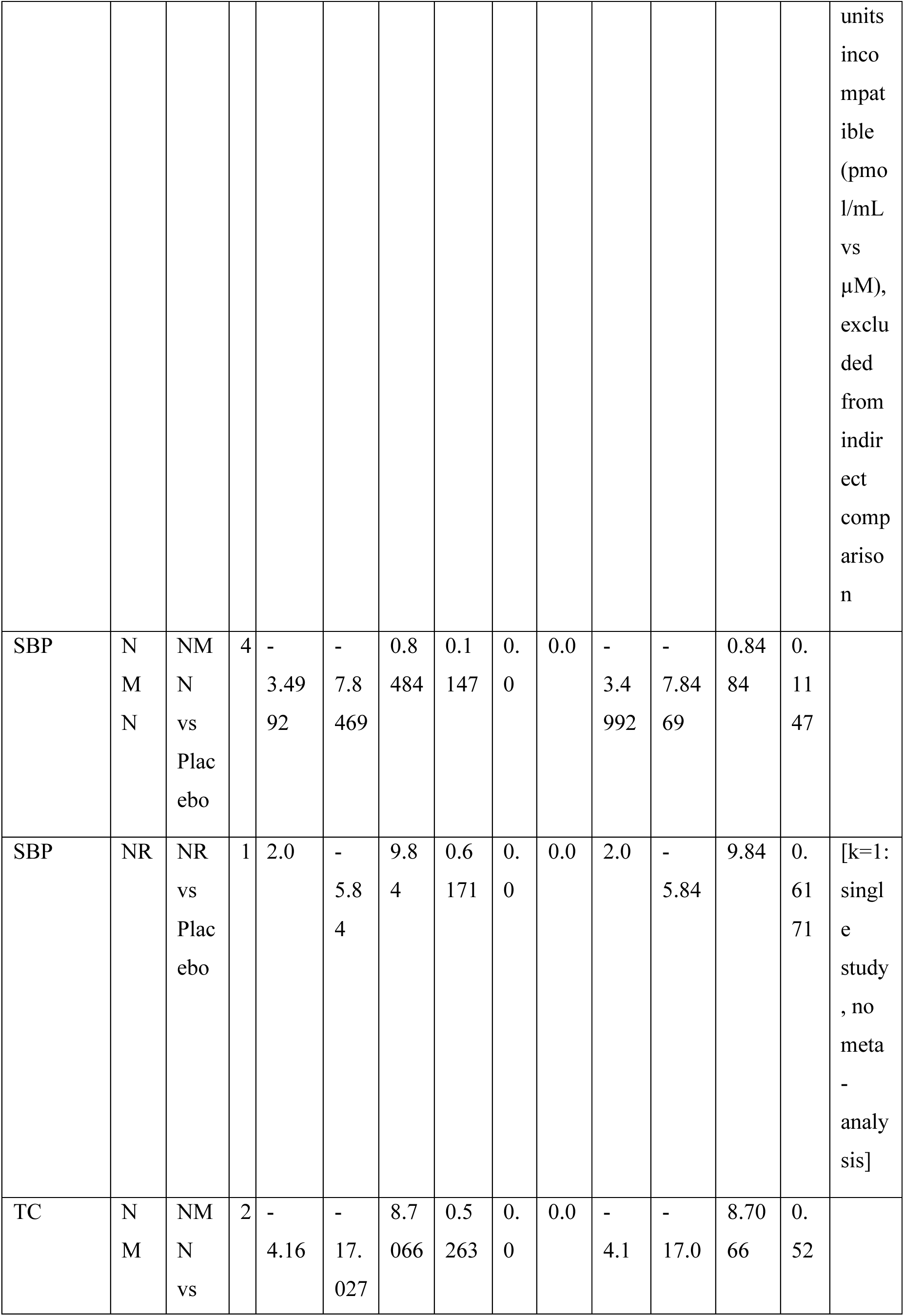
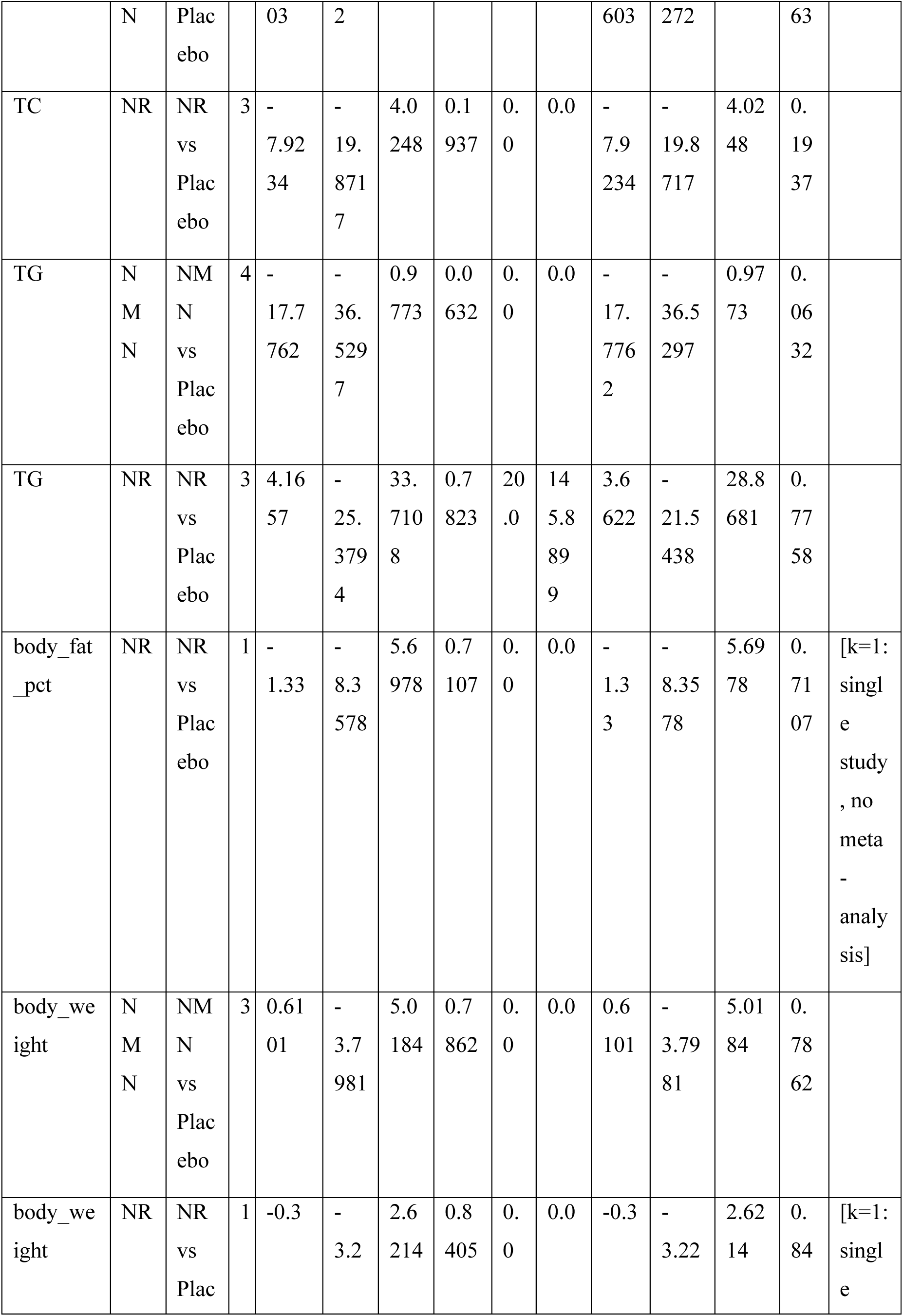
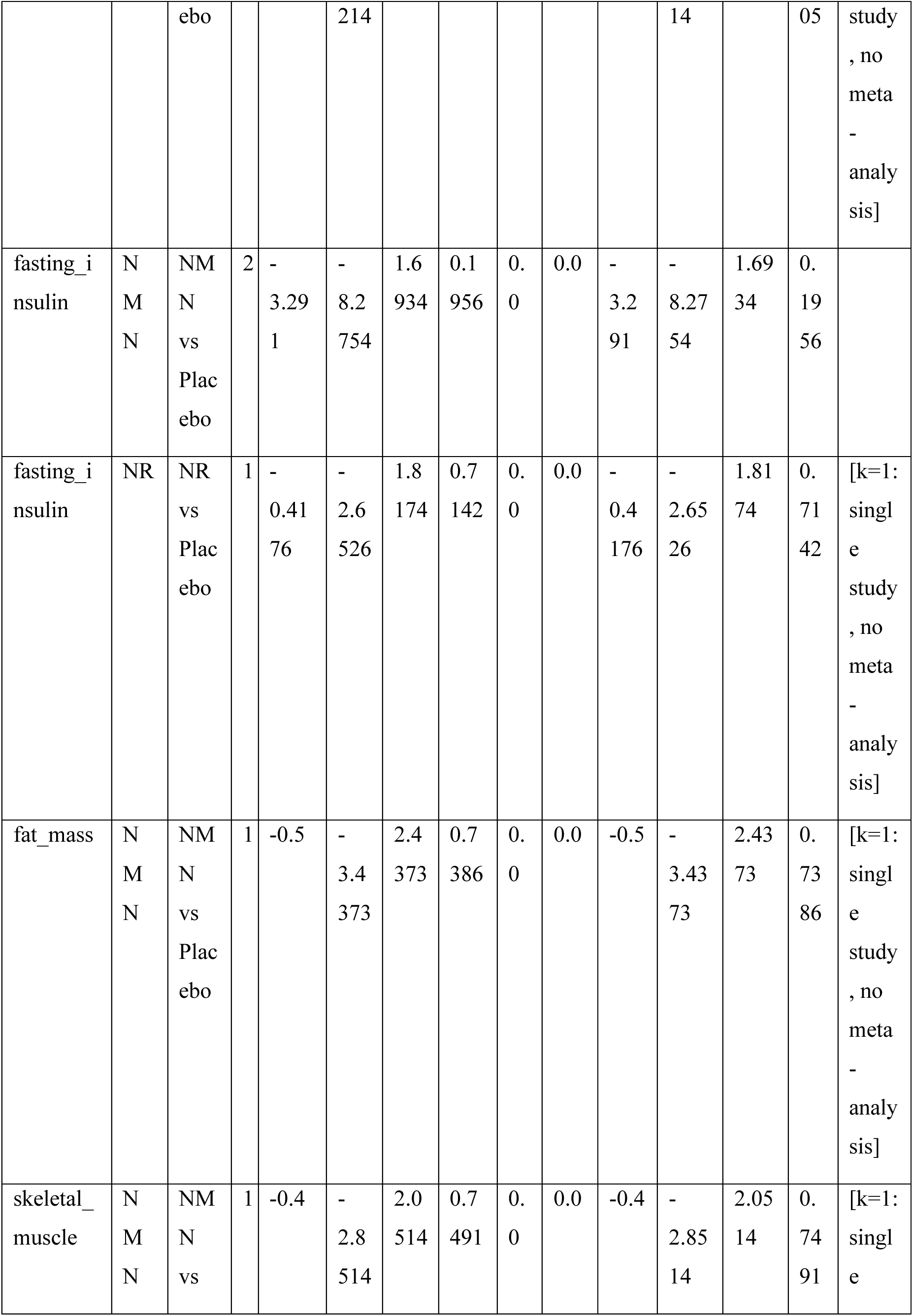
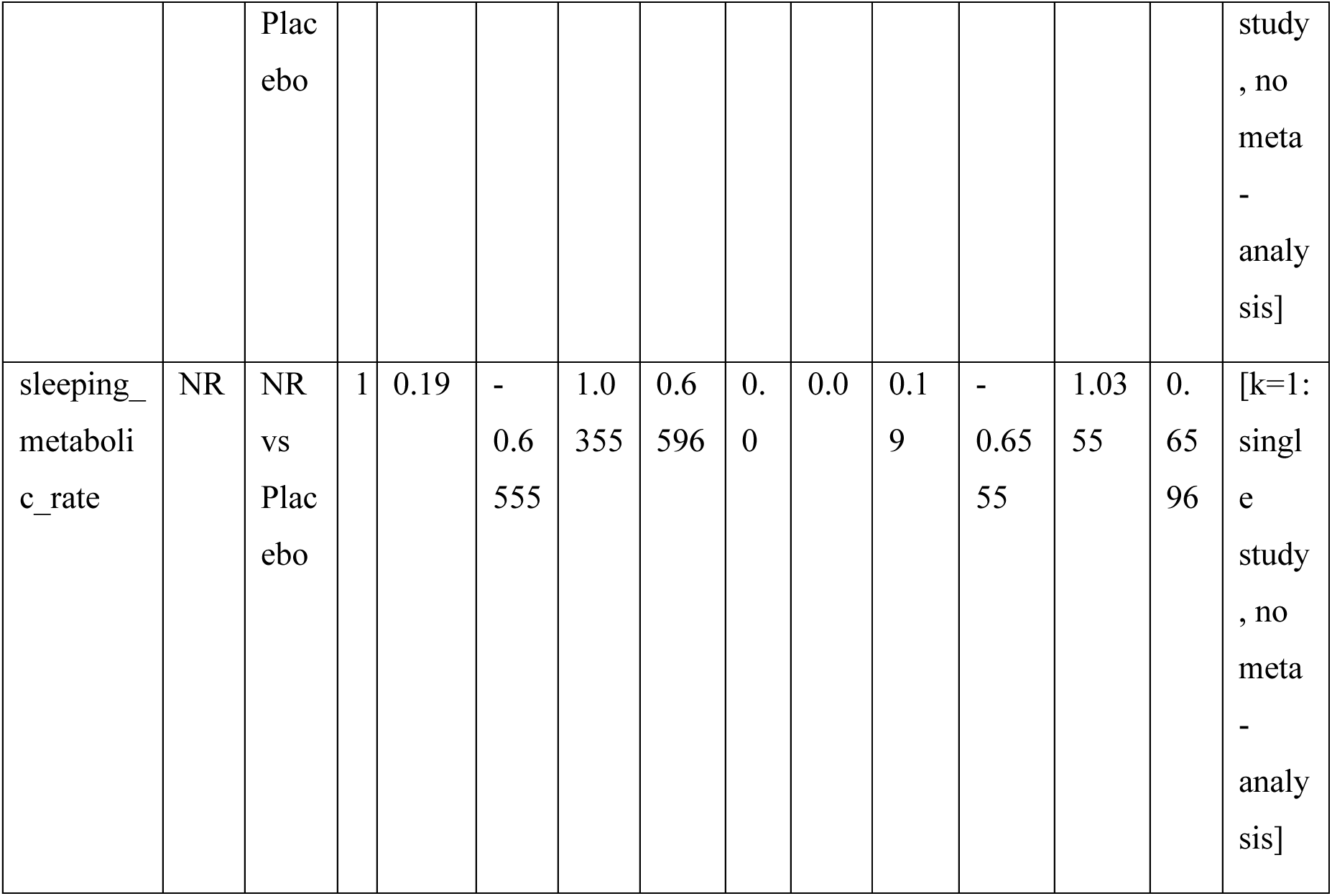
Complete pairwise meta-analysis results for all comparisons.

**Supplementary Table S6.**
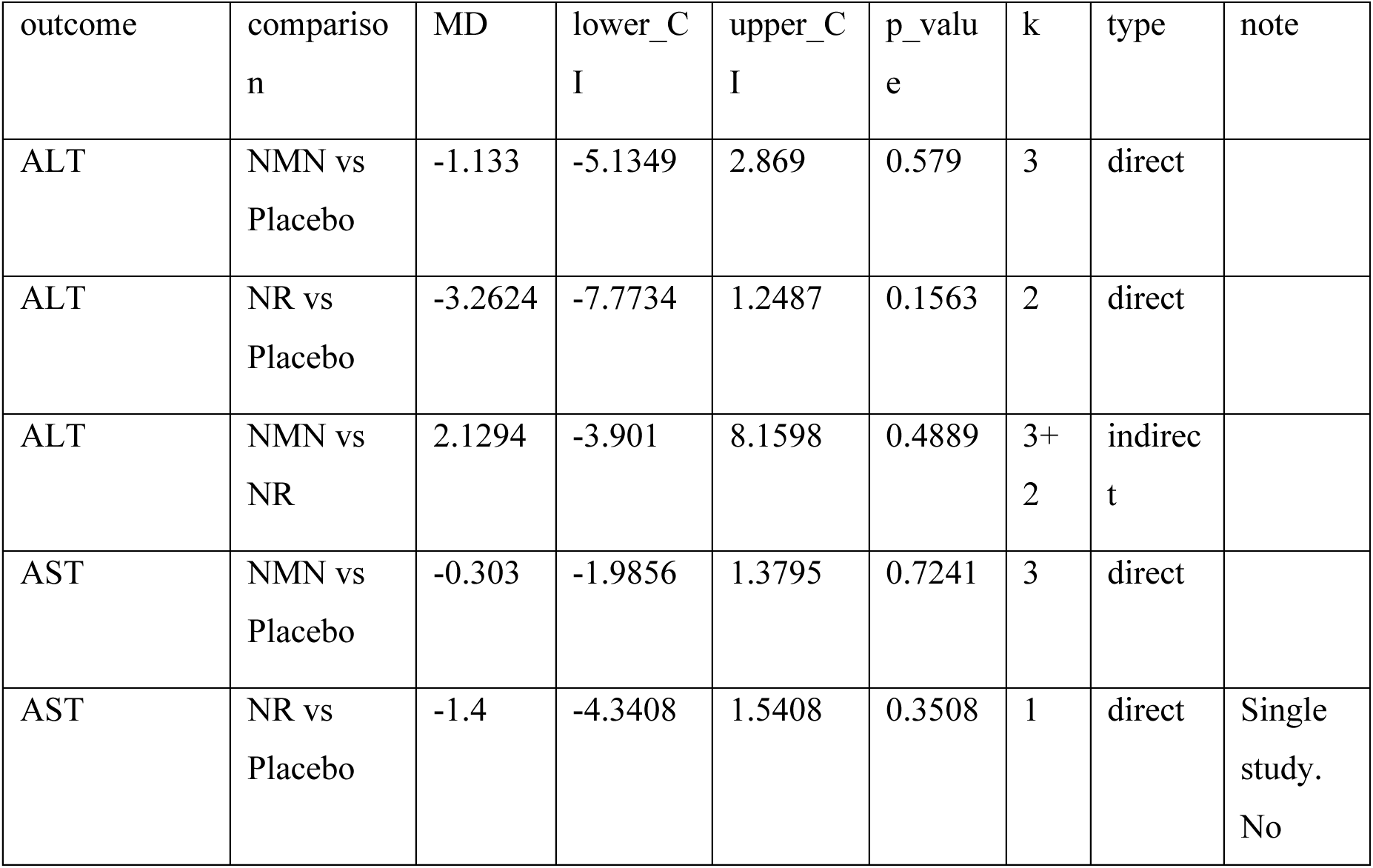
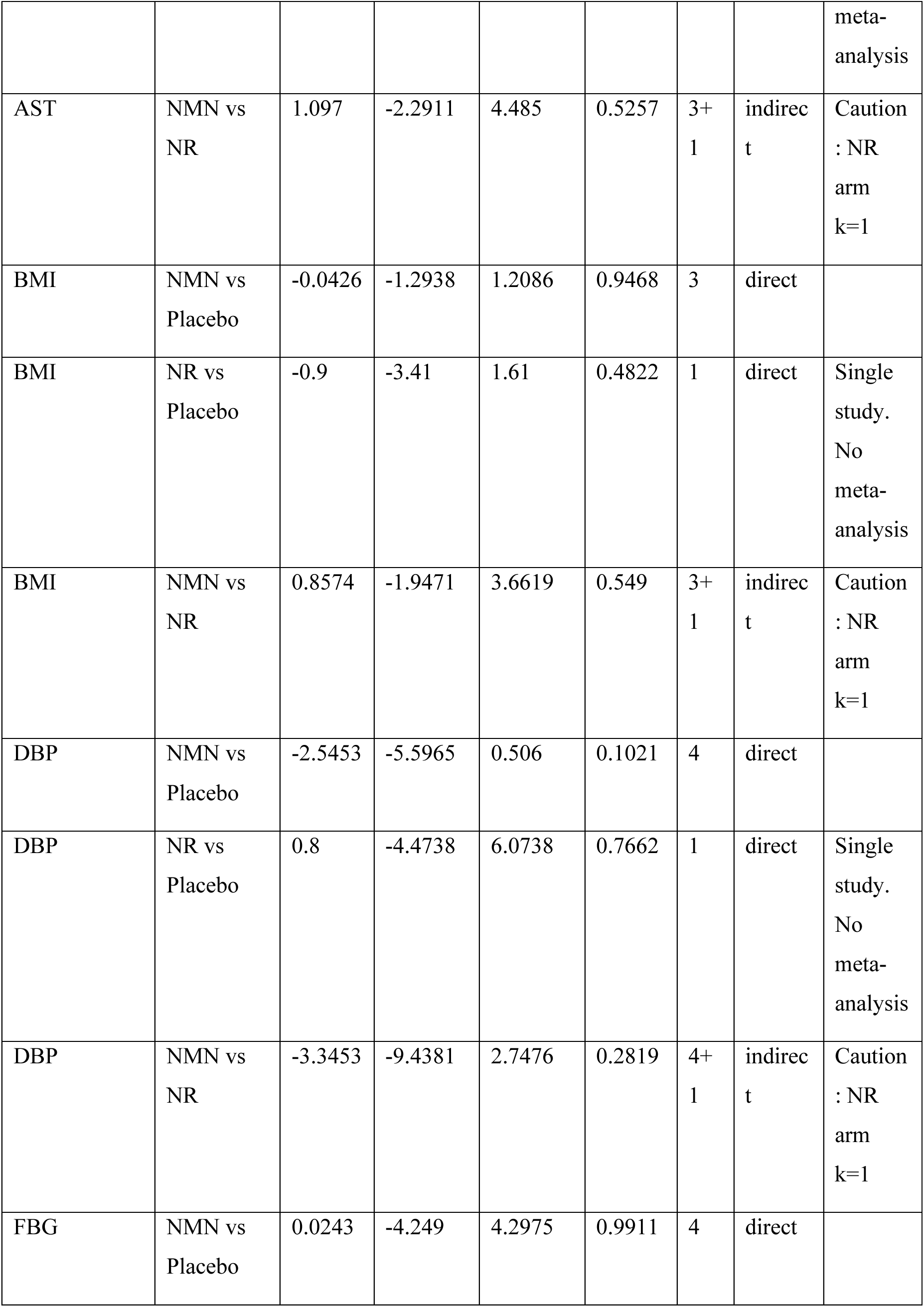
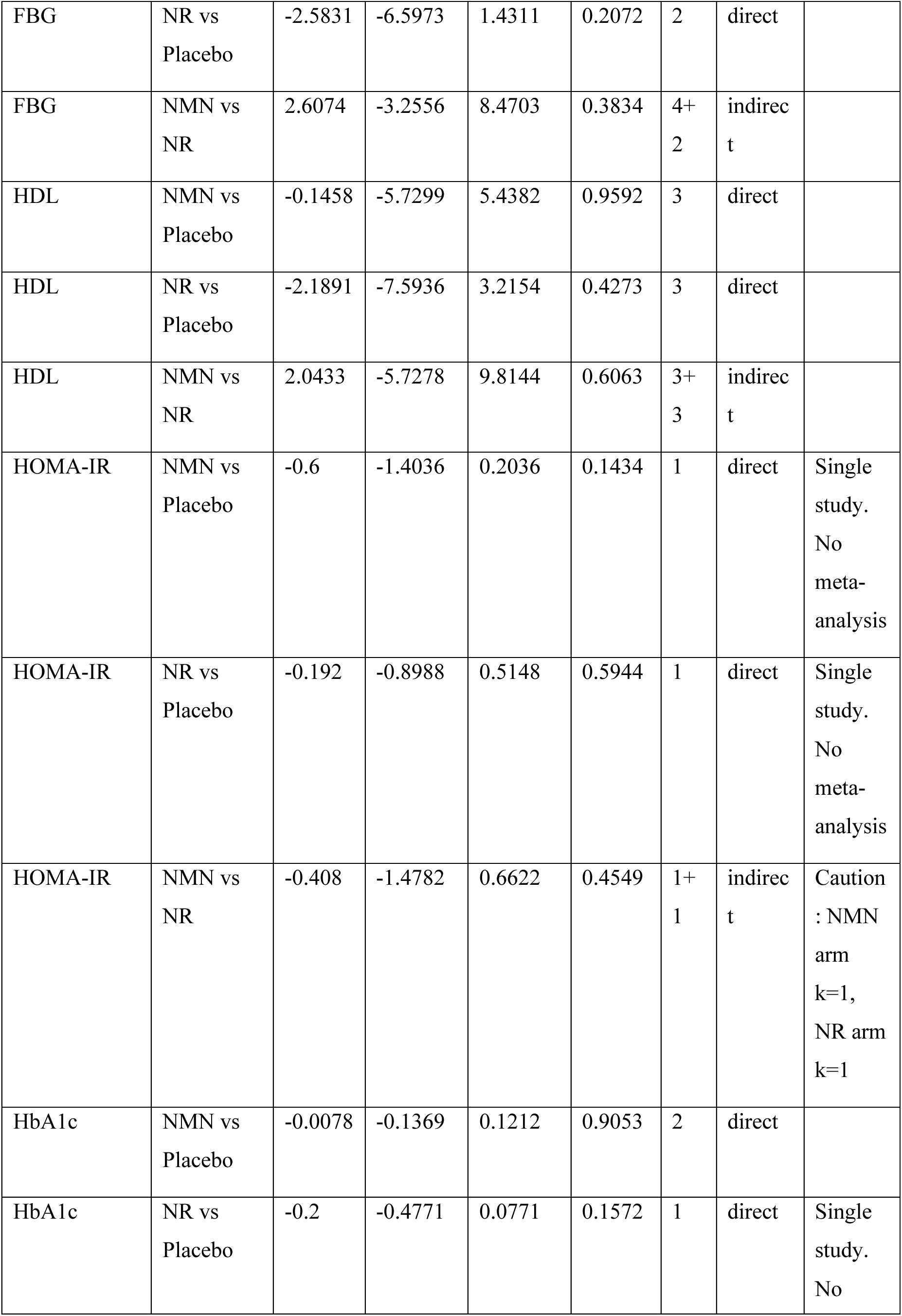
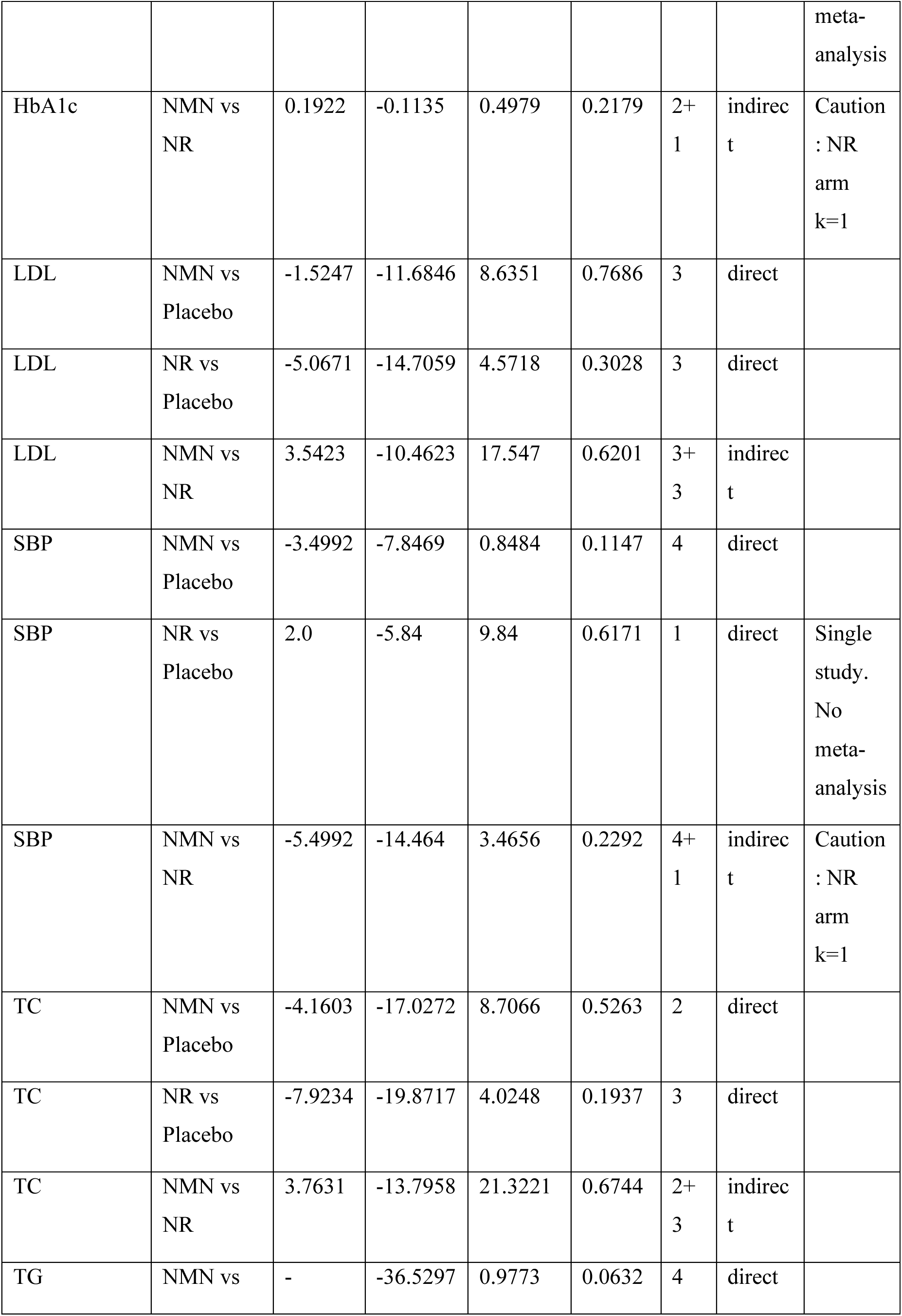
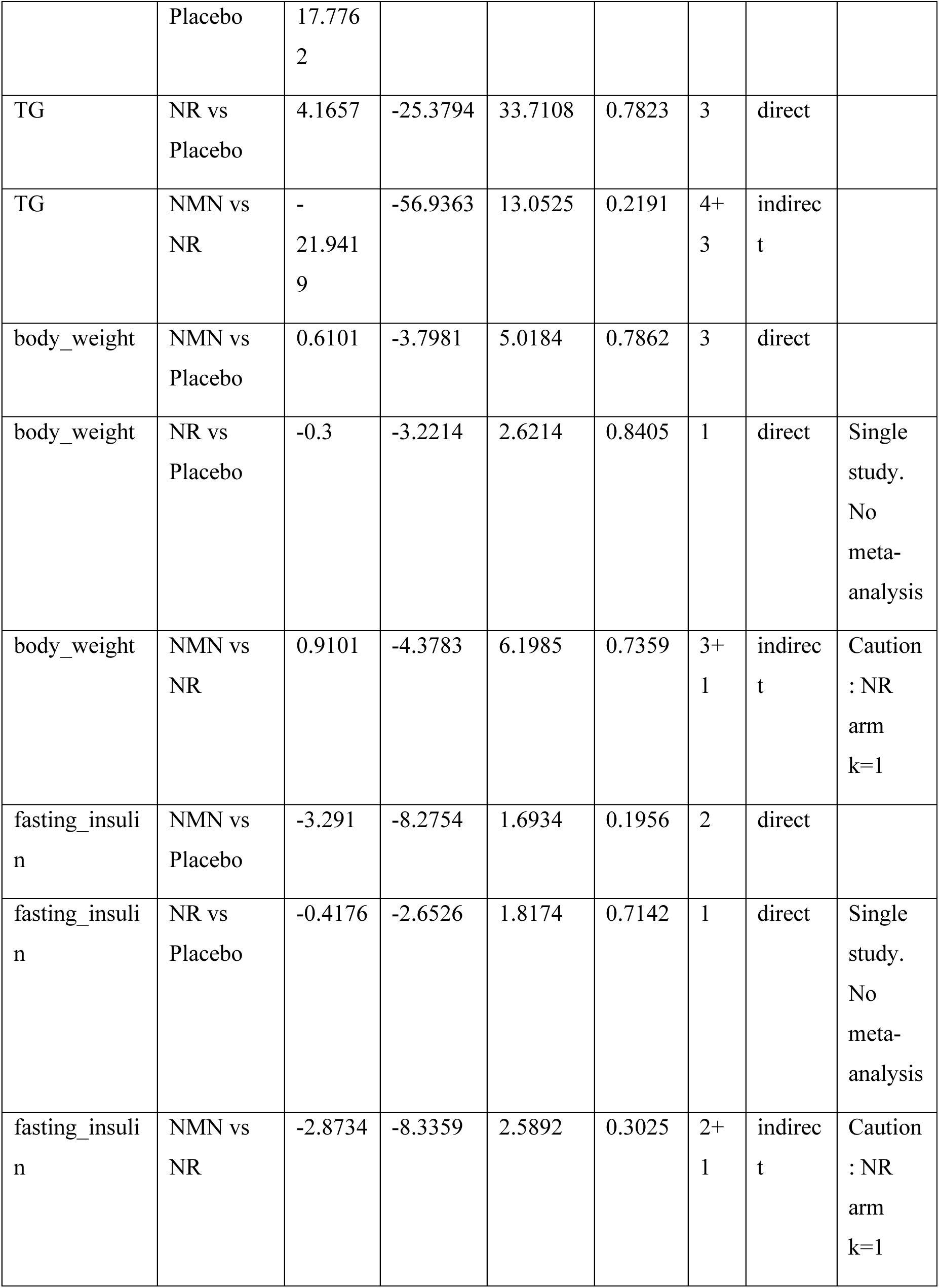
Complete NMA results (direct, indirect, and network estimates).

**Supplementary Table S7.**
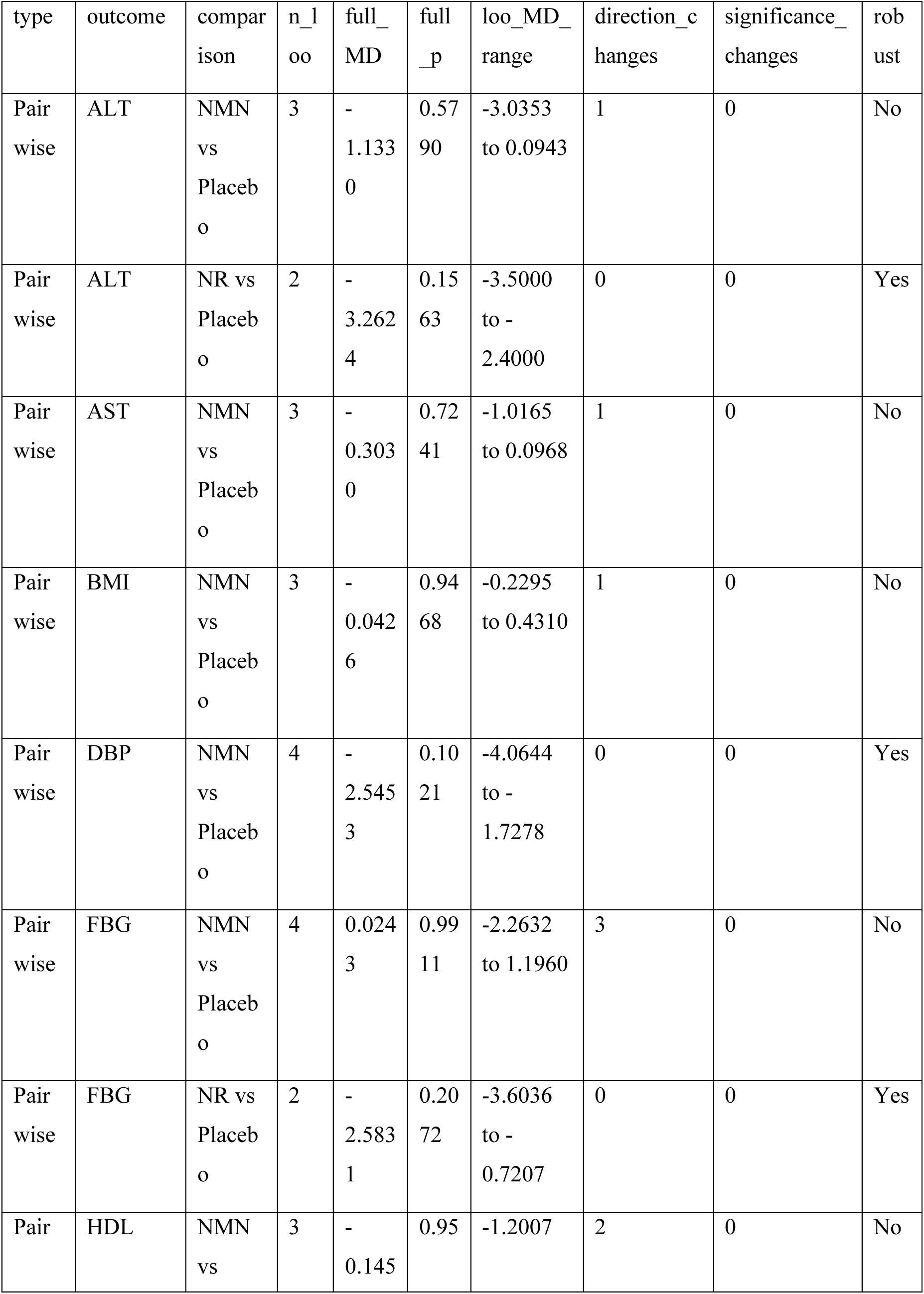
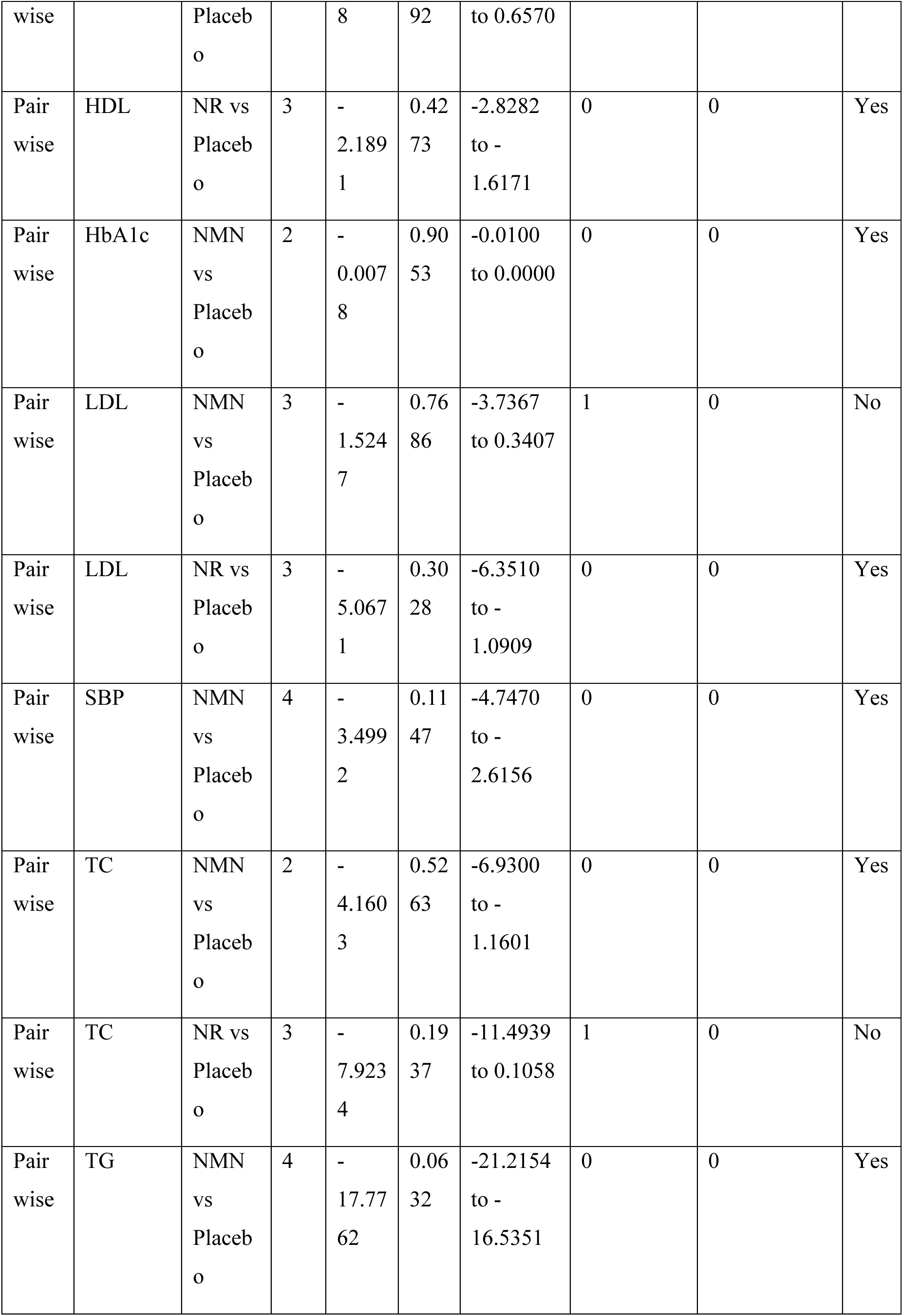
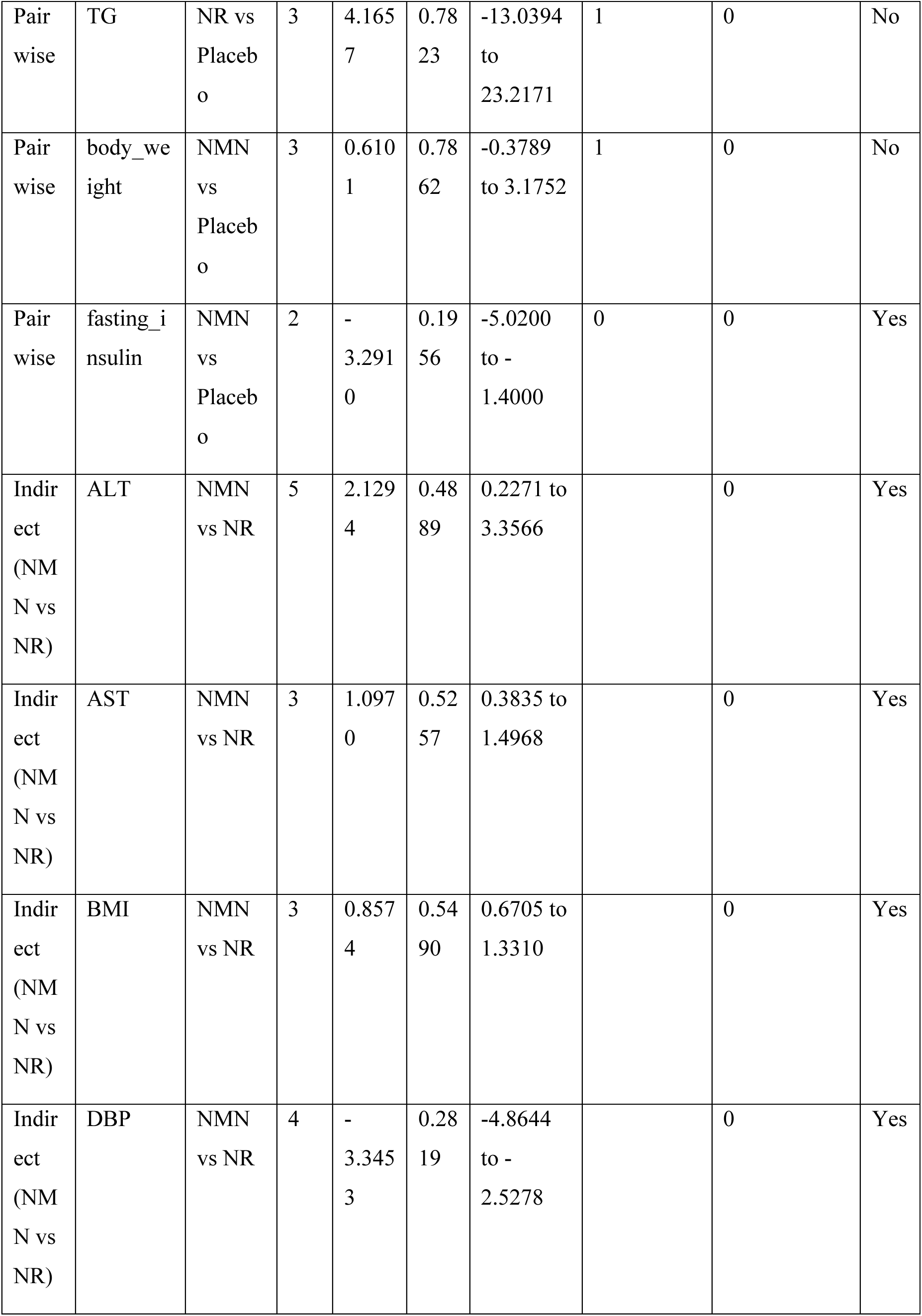
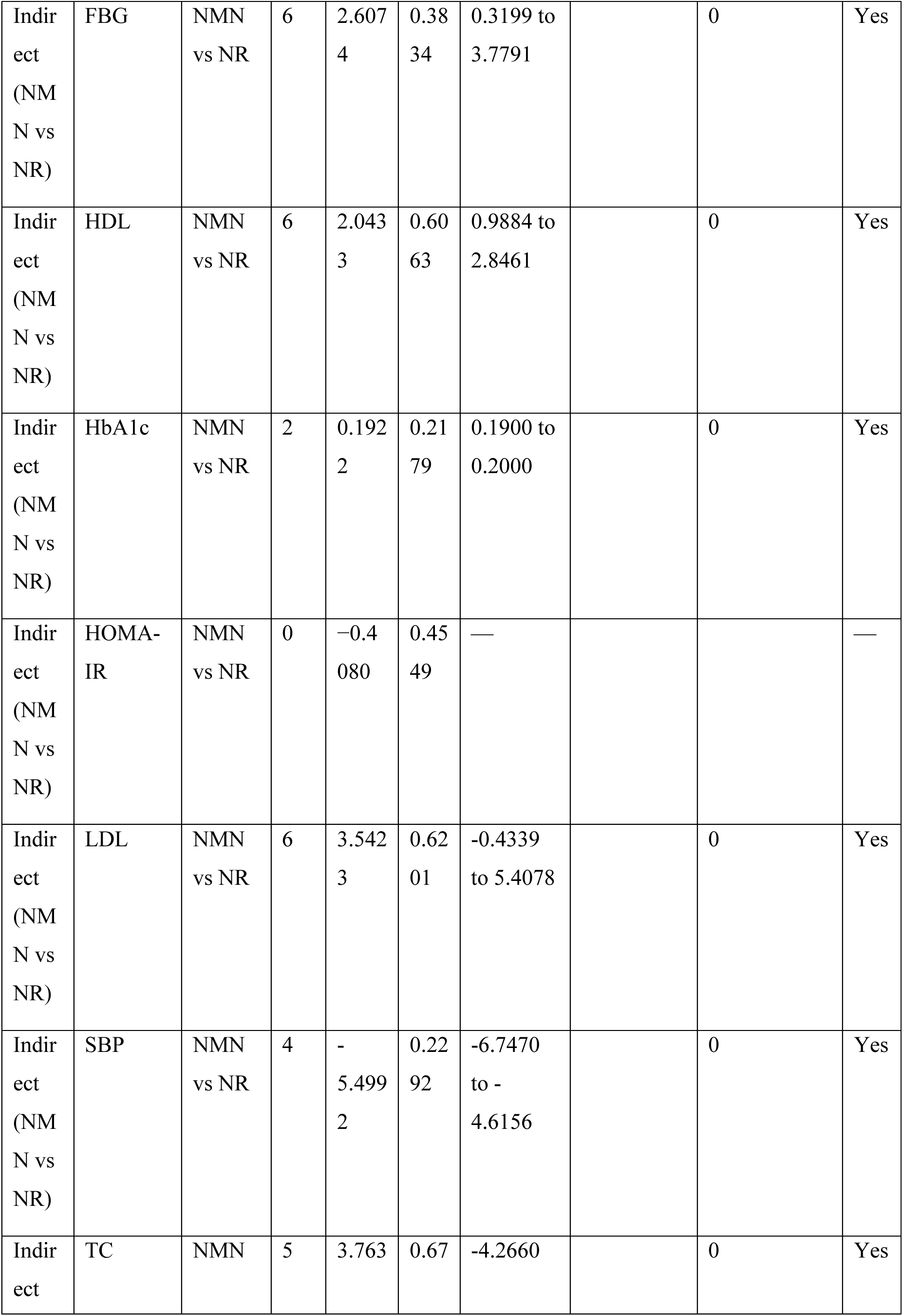
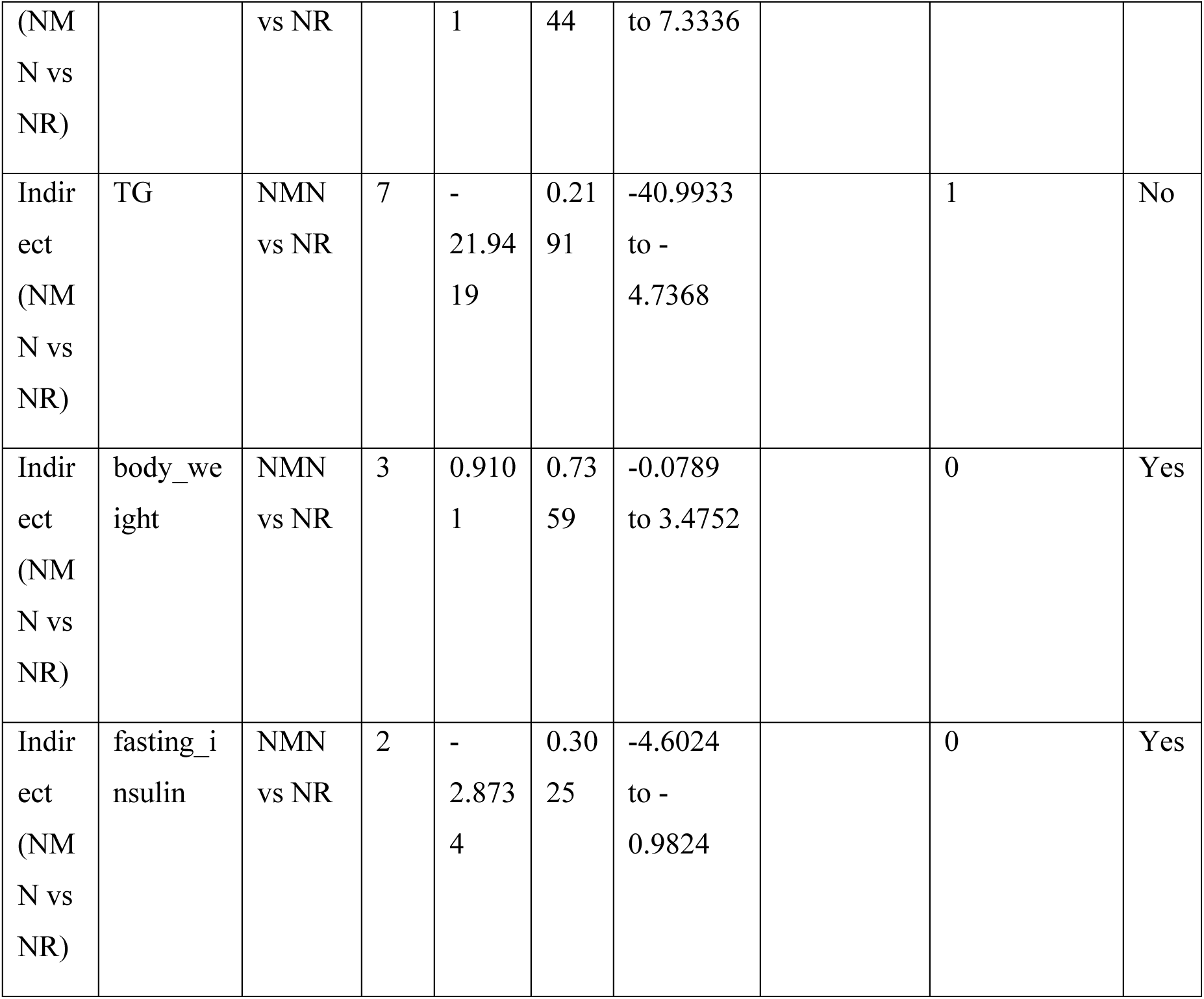
Leave-one-out sensitivity analysis summary.

**Supplementary Table S8.**
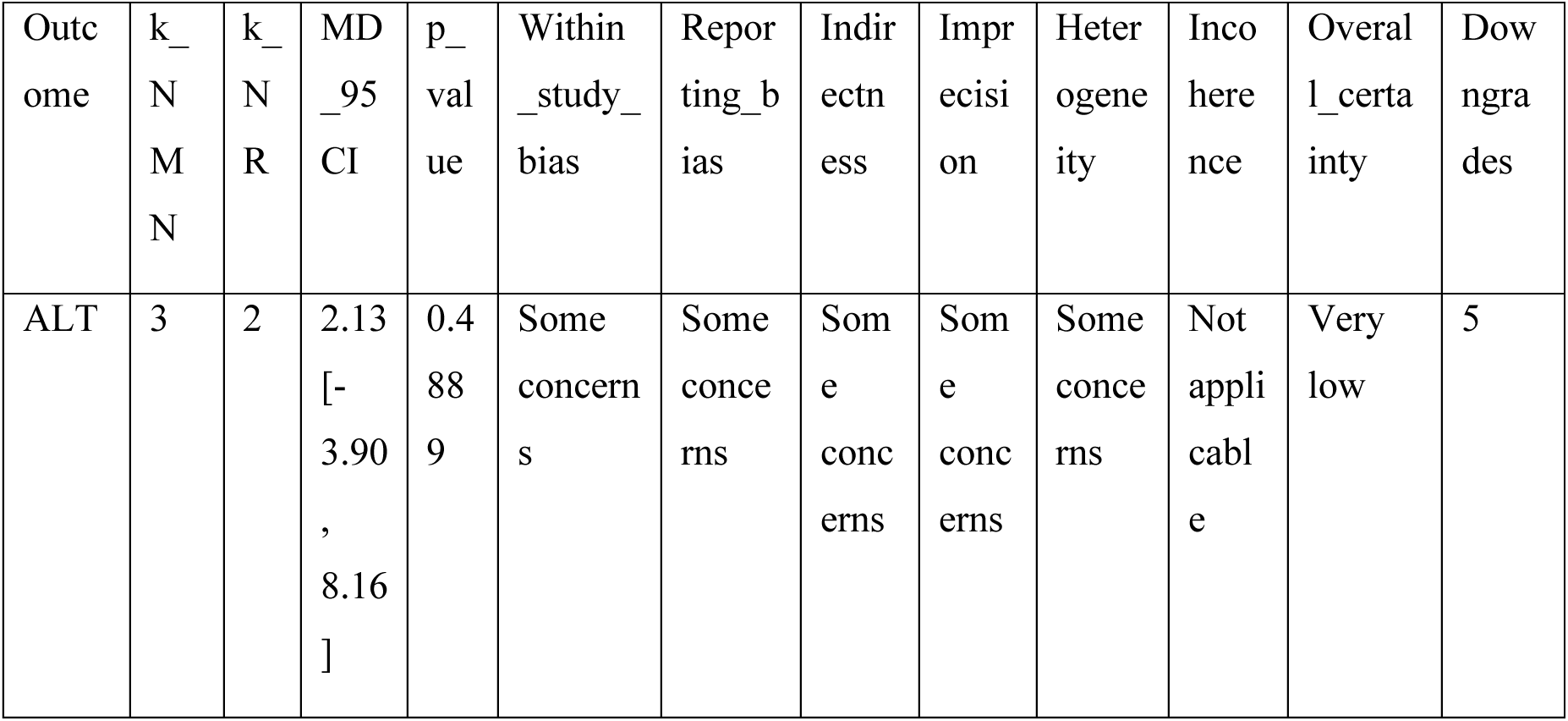
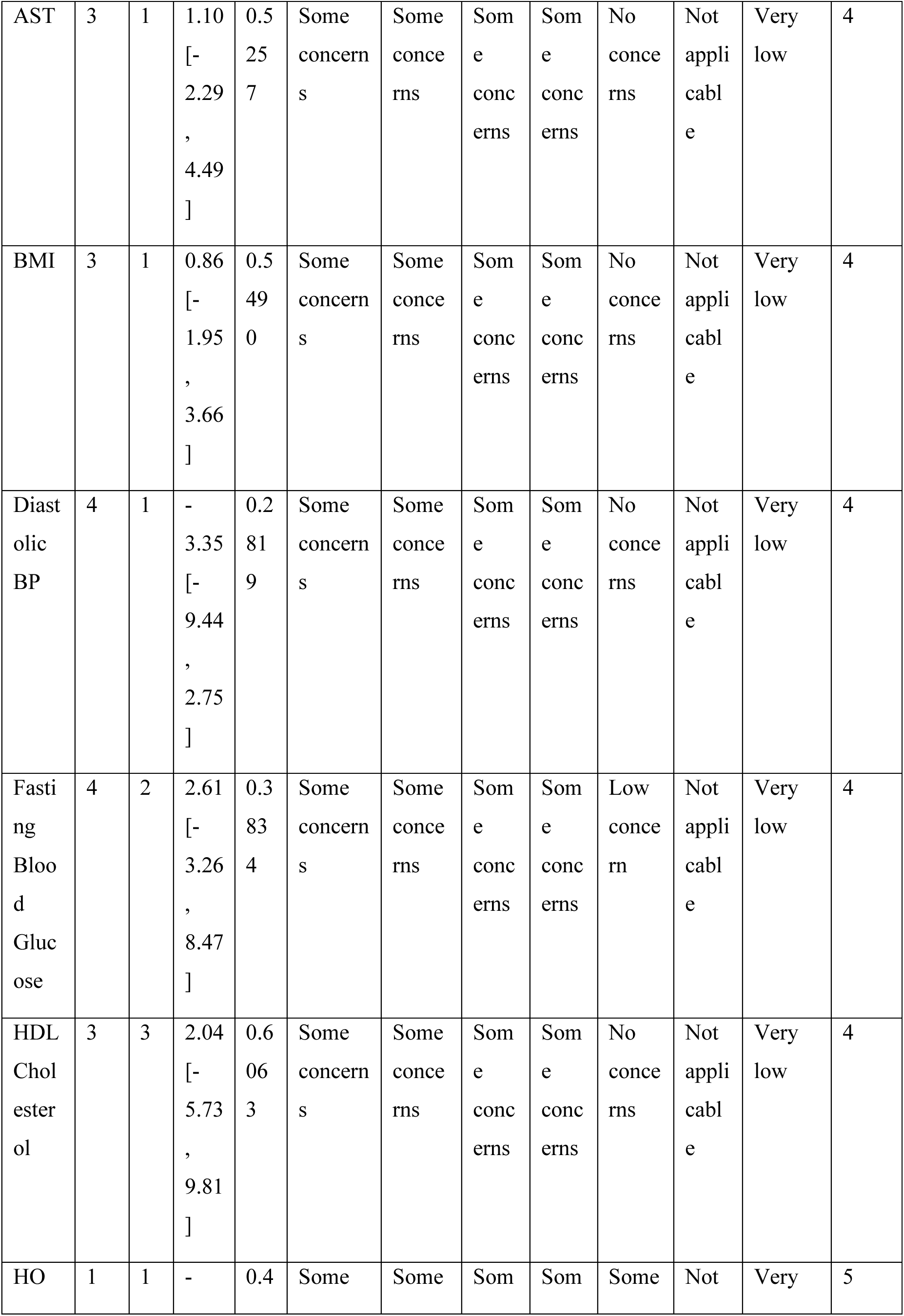
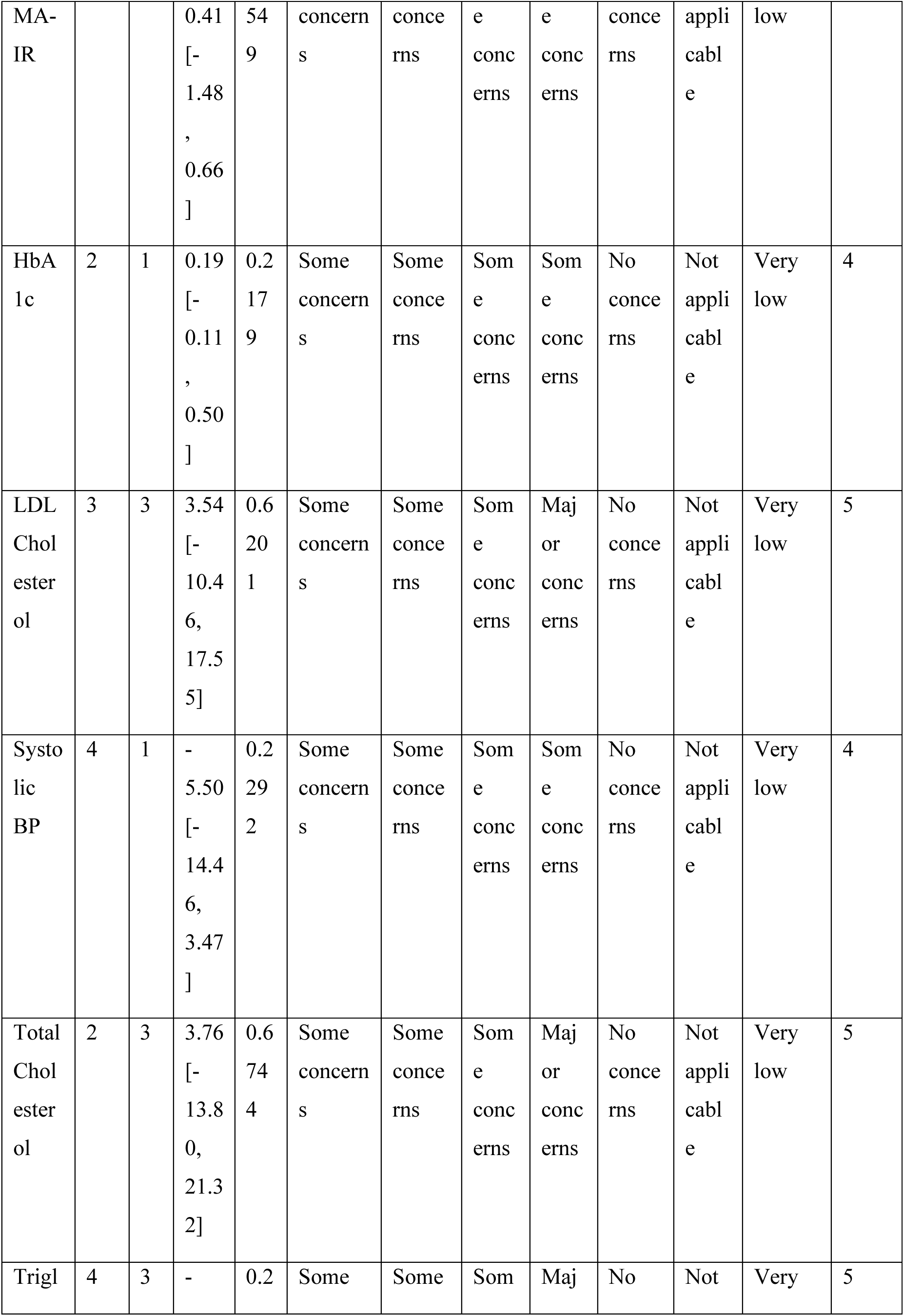
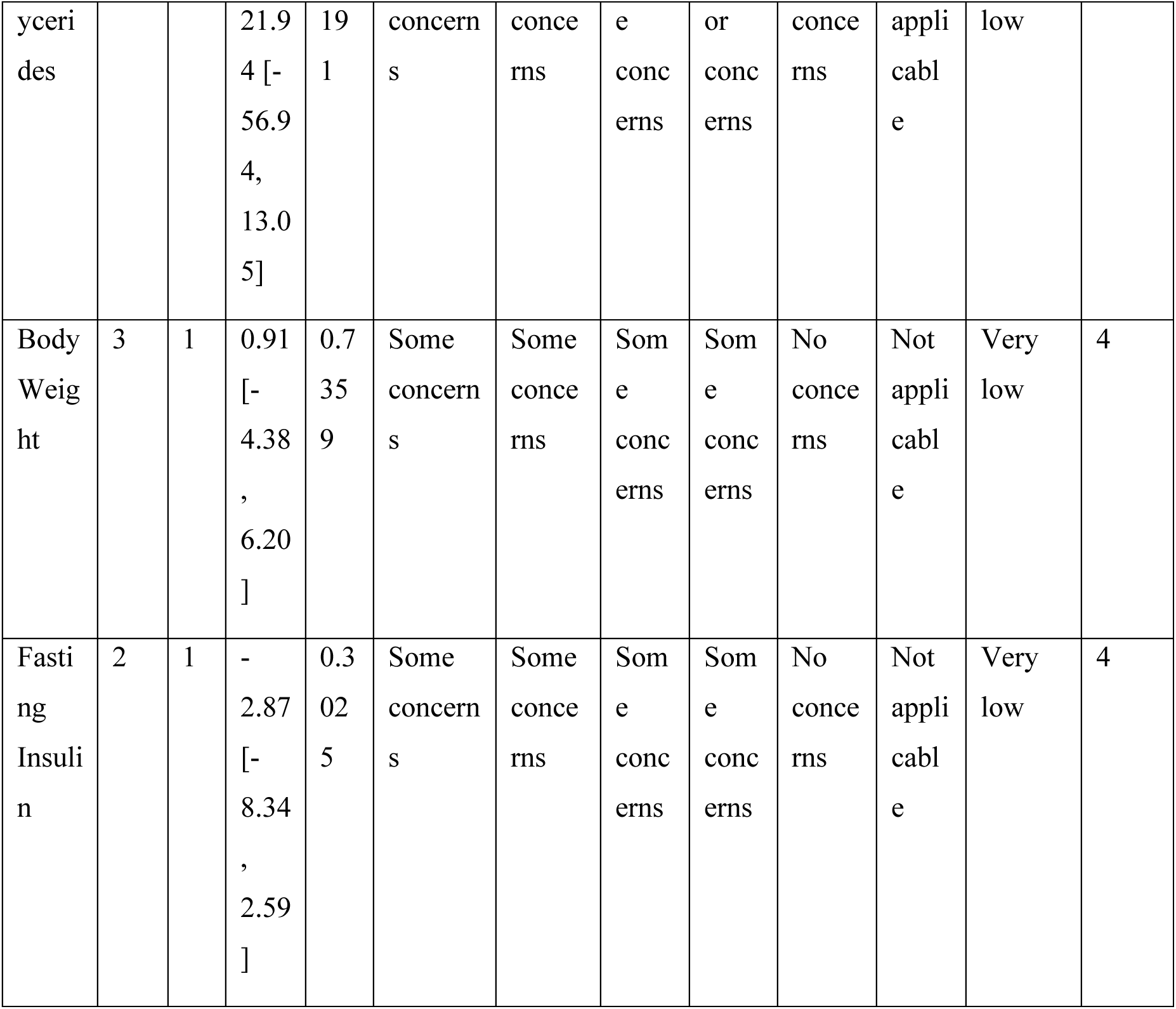
GRADE/CINeMA certainty assessment for 14 indirectly comparable outcomes (NAD+ excluded from indirect estimation).

## References

1. Yoshino J, Baur JA, Imai S-I. NAD+ intermediates: The biology and therapeutic potential of NMN and NR. Cell Metab. 2018;27(3):513–528.

2. Yoshino M, Yoshino J, Kayser BD, et al. Nicotinamide mononucleotide increases muscle insulin sensitivity in prediabetic women. Science. 2021;372(6547):1224–1229.

3. Huang H. A multicentre, randomised, double-blind, parallel design, placebo controlled study to evaluate the efficacy and safety of Uthever (NMN supplement), an orally administered supplementation in middle aged and older adults. Front Aging. 2022;3:851698.

4. Igarashi M, Nakagawa-Nagahama Y, Miura M, et al. Chronic nicotinamide mononucleotide supplementation elevates blood nicotinamide adenine dinucleotide levels and alters muscle function in healthy older men. npj Aging. 2022;8:5.

5. Katayoshi T, Uehata S, Nakashima N, et al. Nicotinamide mononucleotide (NMN) intake increases plasma NMN and insulin levels in healthy subjects. Endocr J. 2023;70(10):1003–1011.

6. Morifuji M, Higashi S, Ebihara S, Nagata K. A randomized, double-blind, placebo-controlled trial of the effects of nicotinamide mononucleotide supplementation on biomarkers of aging in older Japanese adults. Chin Med J. 2024;137(19):2365–2374.

7. Dollerup OL, Christensen B, Svart M, et al. A randomized placebo-controlled clinical trial of nicotinamide riboside in obese men: safety, insulin-sensitivity, and lipid-mobilizing effects. Am J Clin Nutr. 2018;108(2):343–353.

8. Conze D, Brenner C, Kruger CL. Safety and metabolism of long-term administration of NIAGEN (nicotinamide riboside chloride) in a randomized, double-blind, placebo-controlled clinical trial of healthy overweight adults. Sci Rep. 2019;9:9772.

9. Elhassan YS, Kluckova K, Fletcher RS, et al. Nicotinamide riboside augments the aged human skeletal muscle NAD+ metabolome and induces transcriptomic and anti-inflammatory signatures. Cell Rep. 2019;28(7):1717–1728.

10. Remie CME, Roumans KHM, Moonen MPB, et al. Nicotinamide riboside supplementation alters body composition and skeletal muscle acetylcarnitine concentrations in healthy obese humans. Am J Clin Nutr. 2020;112(2):413–426.

11. Brakedal B, Dolle C, Riber F, et al. The NADPARK study: A randomized phase I trial of nicotinamide riboside supplementation in Parkinson’s disease. Cell Metab. 2022;34(3):396–407.

12. Ahmadi A, Begue G, Valencia AP, et al. Randomized crossover clinical trial of niacin and nicotinamide riboside on NAD+ in people with moderate-to-severe CKD. Kidney Int Rep. 2023;8:1055–1068.

13. Norheim KL, Ben Ezra M, Heckenbach I, et al. Effect of nicotinamide riboside on airway inflammation in COPD: a randomized, placebo-controlled trial. Nat Aging. 2024;4:1772–1781.

14. Orr ME, Kotkowski E, Rabinovici GD, et al. Nicotinamide riboside in people with mild cognitive impairment (NR-MCI): A pilot study. Alzheimers Dement (N Y). 2024;10:e12444.

15. Lin J, King AC. Nicotinamide riboside and exercise to augment vascular and brain health in hypertensive older adults: pilot results from the NREX trial. Nutrients. 2025;17(1):112.

16. Bandi R, Gupta R, Koppala D, et al. A randomized, double-blind, placebo-controlled clinical trial assessing efficacy and safety of LN22199 and nicotinamide riboside on biomarkers of aging. J Gerontol A. 2025;80(2):glae260.

17. Page MJ, McKenzie JE, Bossuyt PM, et al. The PRISMA 2020 statement: an updated guideline for reporting systematic reviews. BMJ. 2021;372:n71.

18. Hutton B, Salanti G, Caldwell DM, et al. The PRISMA extension statement for reporting of systematic reviews incorporating network meta-analyses. Ann Intern Med. 2015;162(11):777–784.

19. Sterne JAC, Savovic J, Page MJ, et al. RoB 2: a revised tool for assessing risk of bias in randomised trials. BMJ. 2019;366:l4898.

20. Bucher HC, Guyatt GH, Griffith LE, Walter SD. The results of direct and indirect treatment comparisons in meta-analysis of randomized controlled trials. J Clin Epidemiol. 1997;50(6):683–691.

21. Nikolakopoulou A, Higgins JPT, Papakonstantinou T, et al. CINeMA: An approach for assessing confidence in the results of a network meta-analysis. PLoS Med. 2020;17(4):e1003082.

22. Wan X, Wang W, Liu J, Tong T. Estimating the sample mean and standard deviation from the sample size, median, range and/or interquartile range. BMC Med Res Methodol. 2014;14:135.

23. DerSimonian R, Laird N. Meta-analysis in clinical trials. Control Clin Trials. 1986;7(3):177–188.

24. Zheng Y, Guo Y, Chen Y, et al. Nicotinamide mononucleotide supplementation and cardiometabolic health: a systematic review and meta-analysis of randomized controlled trials. Nutrients. 2024;16(15):2526.

25. Alegre GFS, Pastore GM. NAD+ precursors nicotinamide mononucleotide (NMN) and nicotinamide riboside (NR): Potential dietary contribution to health. Curr Nutr Rep. 2024;13:261–282.

26. Nascimento GP, Nogueira-de-Almeida CA. Nicotinamide riboside supplementation in adults with cardiometabolic disease: a systematic review. Eur J Clin Nutr. 2024;78:815–826.

27. Grozio A, Mills KF, Yoshino J, et al. Slc12a8 is a nicotinamide mononucleotide transporter. Nat Metab. 2019;1(1):47–57.

28. Canto C, Houtkooper RH, Pirinen E, et al. The NAD+ precursor nicotinamide riboside enhances oxidative metabolism and protects against high-fat diet-induced obesity. Cell Metab. 2012;15(6):838–847.

29. Mills KF, Yoshida S, Stein LR, et al. Long-term administration of nicotinamide mononucleotide mitigates age-associated physiological decline in mice. Cell Metab. 2016;24(6):795–806.

30. Trammell SAJ, Schmidt MS, Weidemann BJ, et al. Nicotinamide riboside is uniquely and orally bioavailable in mice and humans. Nat Commun. 2016;7:12948.

31. Verdin E. NAD+ in aging, metabolism, and neurodegeneration. Science. 2015;350(6265):1208–1213.

32. Camacho-Pereira J, Tarrago MG, Chini CCS, et al. CD38 dictates age-related NAD+ decline and mitochondrial dysfunction through an SIRT3-dependent mechanism. Cell Metab. 2016;23(6):1127–1139.

33. Imai S, Guarente L. NAD+ and sirtuins in aging and disease. Trends Cell Biol. 2014;24(8):464–471.

