## Supplementary figures and images for "Mapping Evidence Gap Between NMN and NR for Metabolic Outcomes: A Systematic Review, Transitivity Assessment, and Indirect Comparison Meta-Analysis"

### forest_ALT_NMN_vs_PBO.pdf

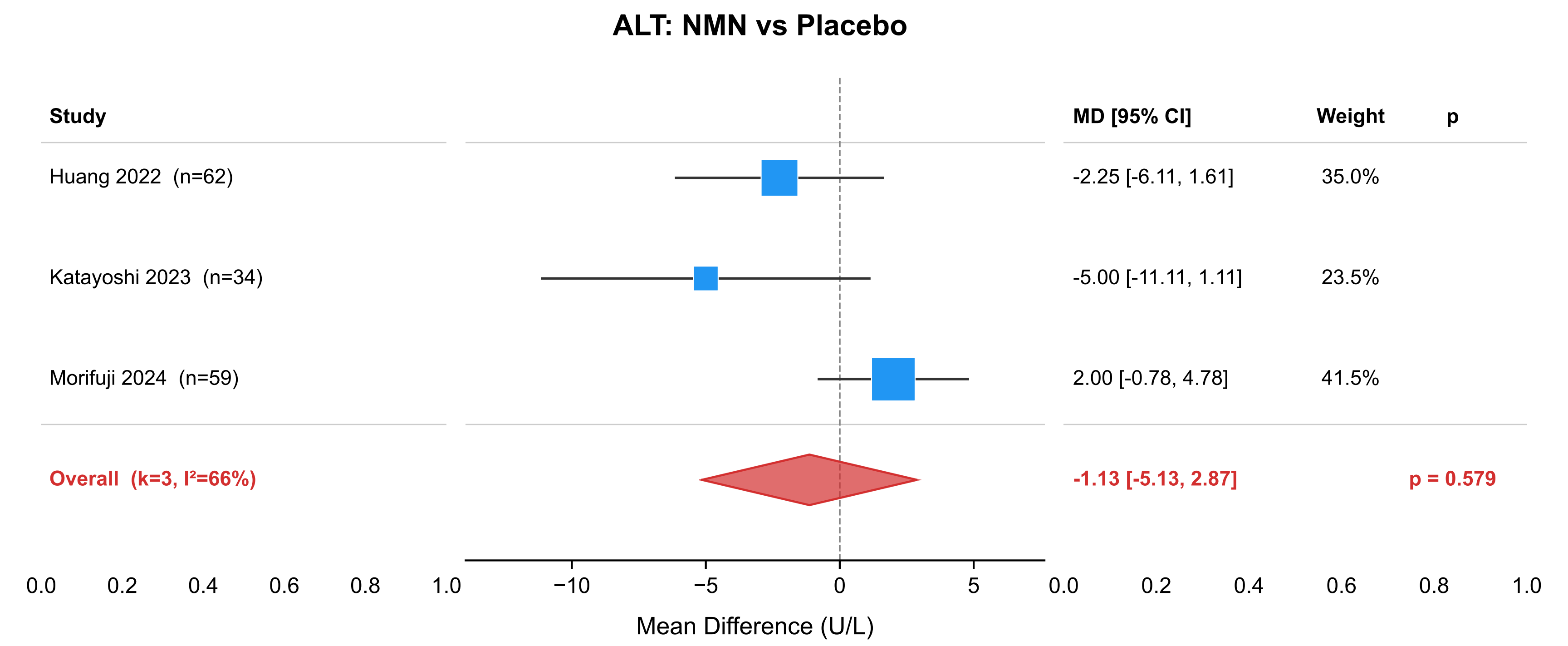

### forest_ALT_NMN_vs_PBO.png

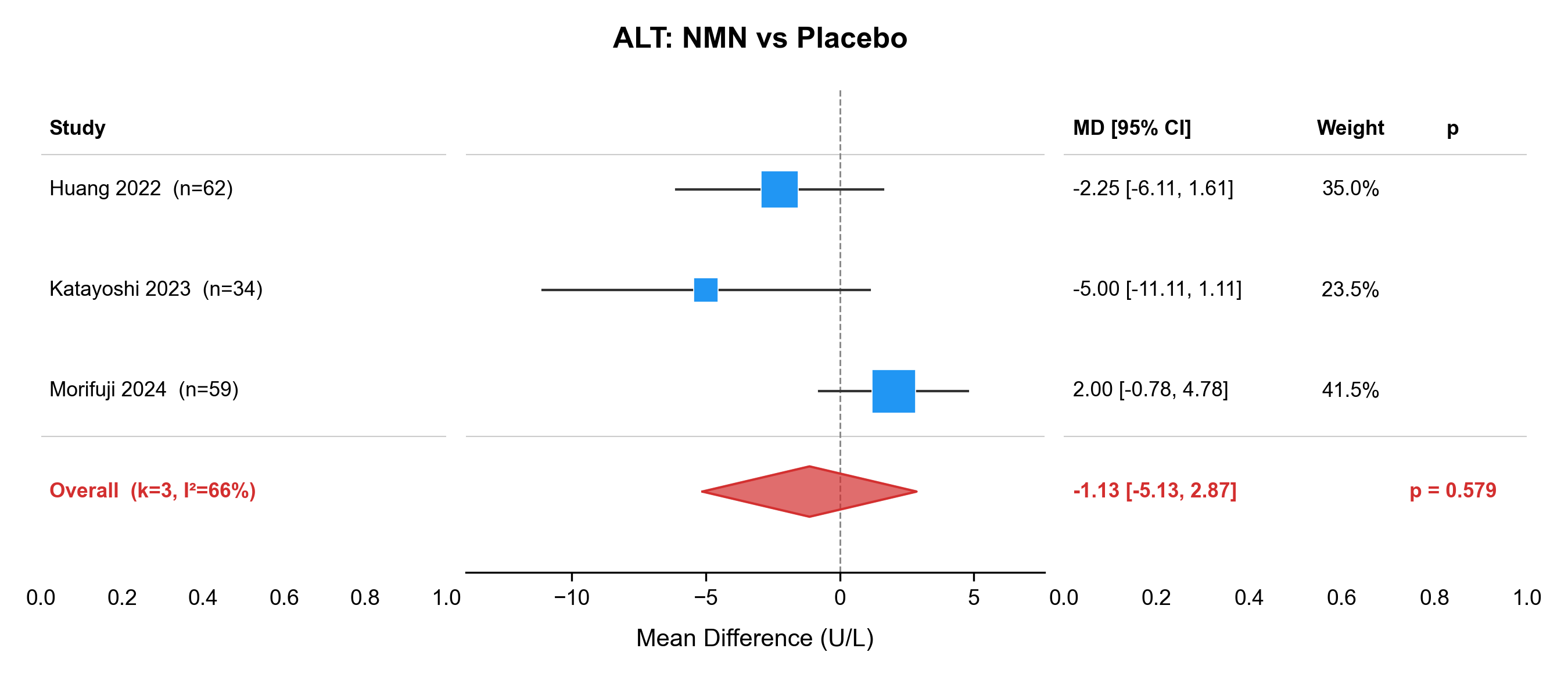

### forest_ALT_NR_vs_PBO.pdf

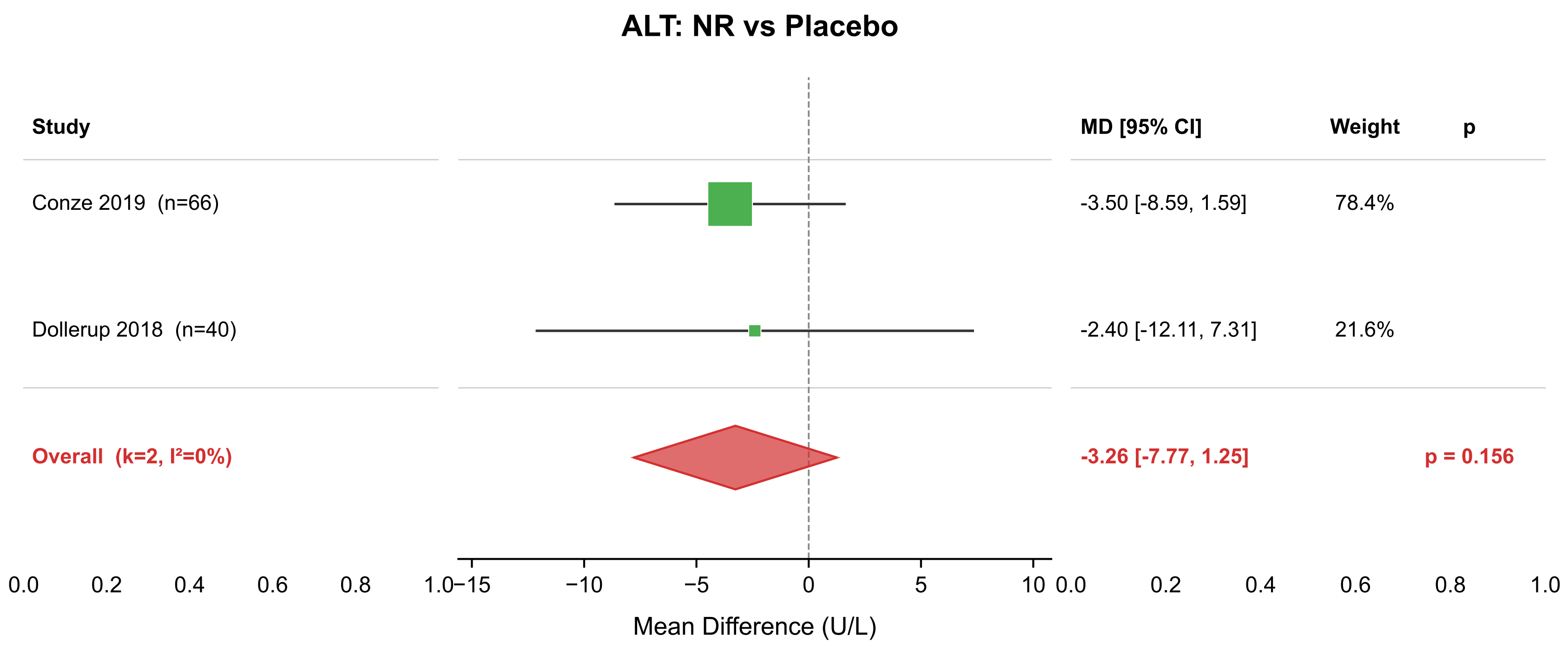

### forest_ALT_NR_vs_PBO.png

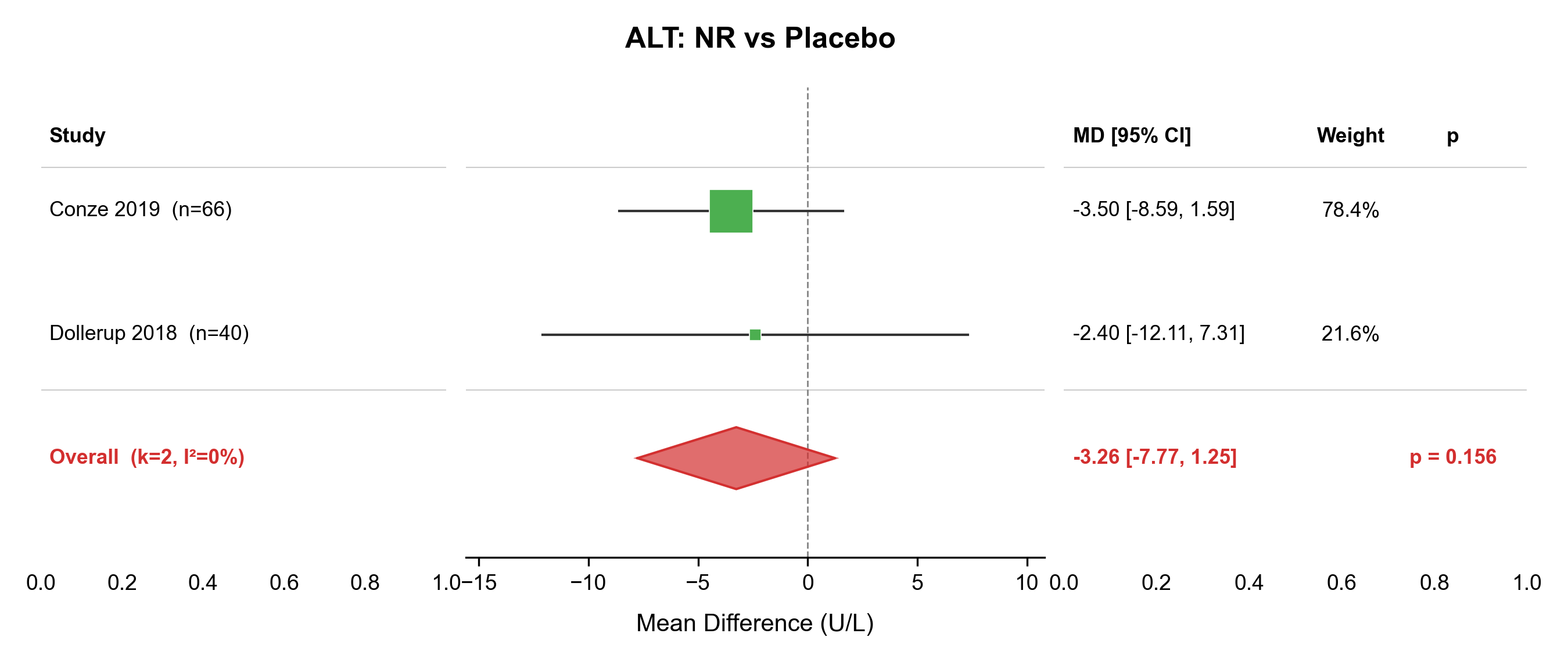

### forest_AST_NMN_vs_PBO.pdf

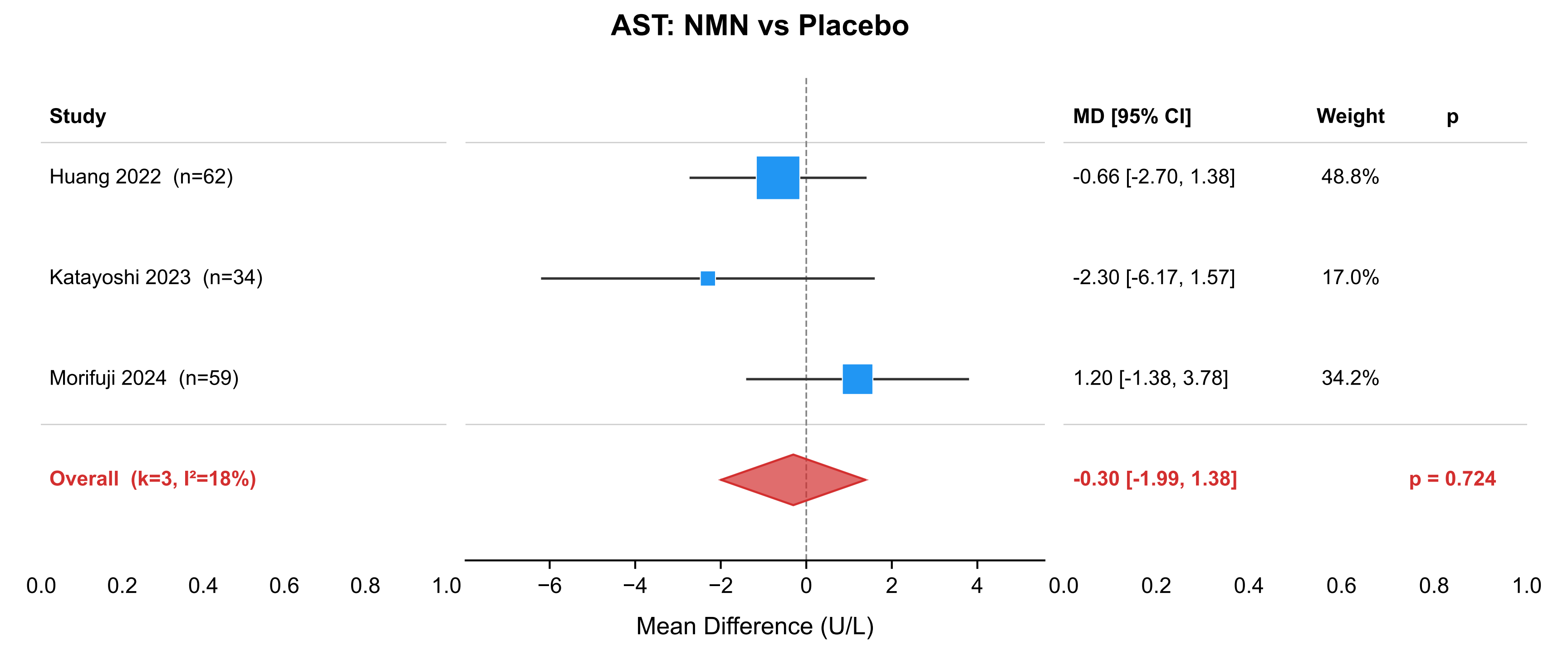

### forest_AST_NMN_vs_PBO.png

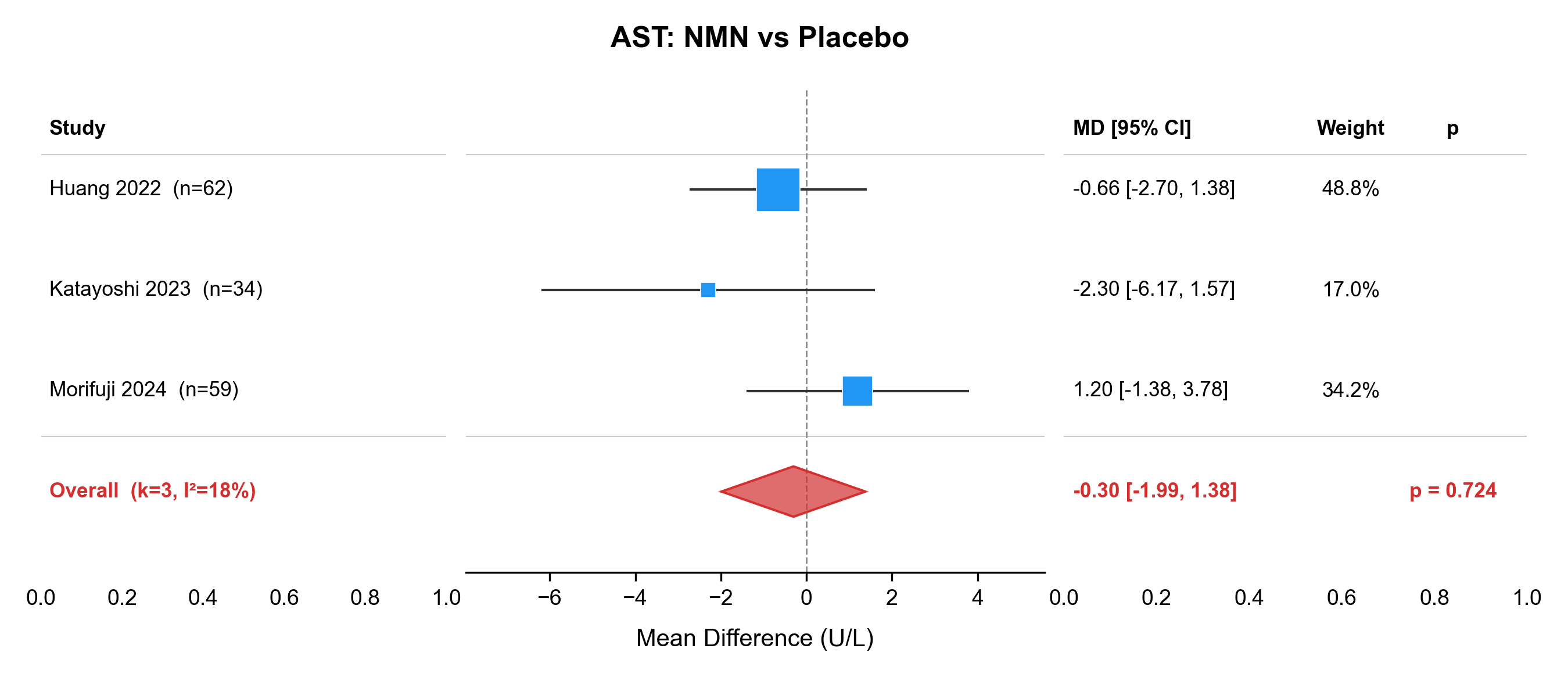

### forest_BMI_NMN_vs_PBO.pdf

#### BMI: NMN vs Placebo

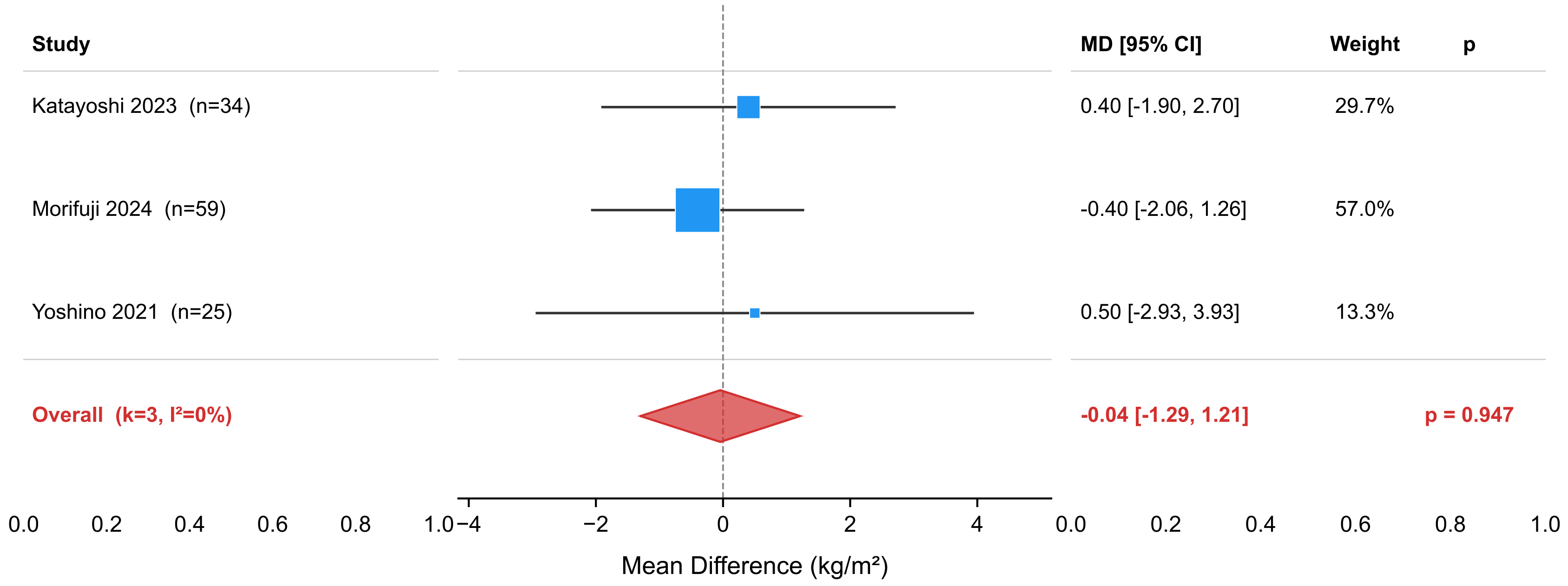

### forest_BMI_NMN_vs_PBO.png

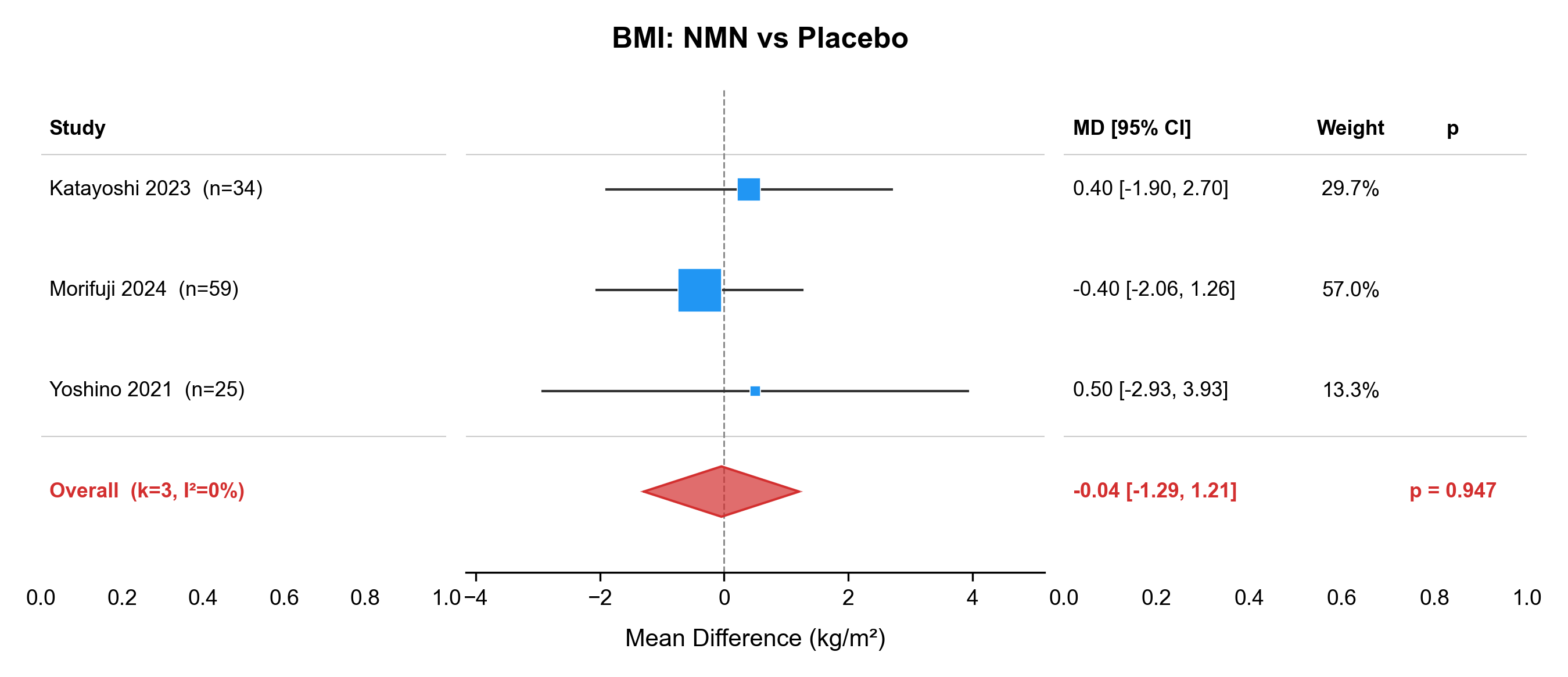

### forest_body_weight_NMN_vs_PBO.pdf

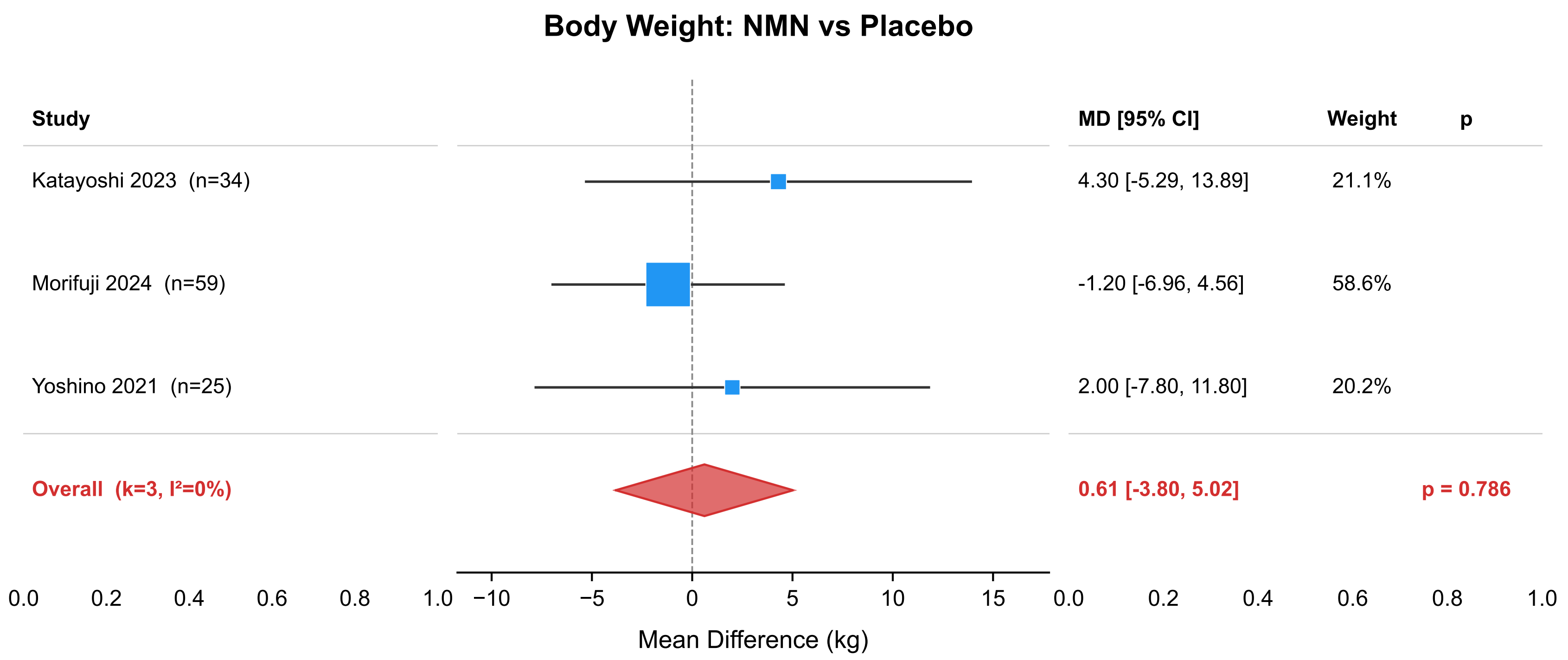

### forest_body_weight_NMN_vs_PBO.png

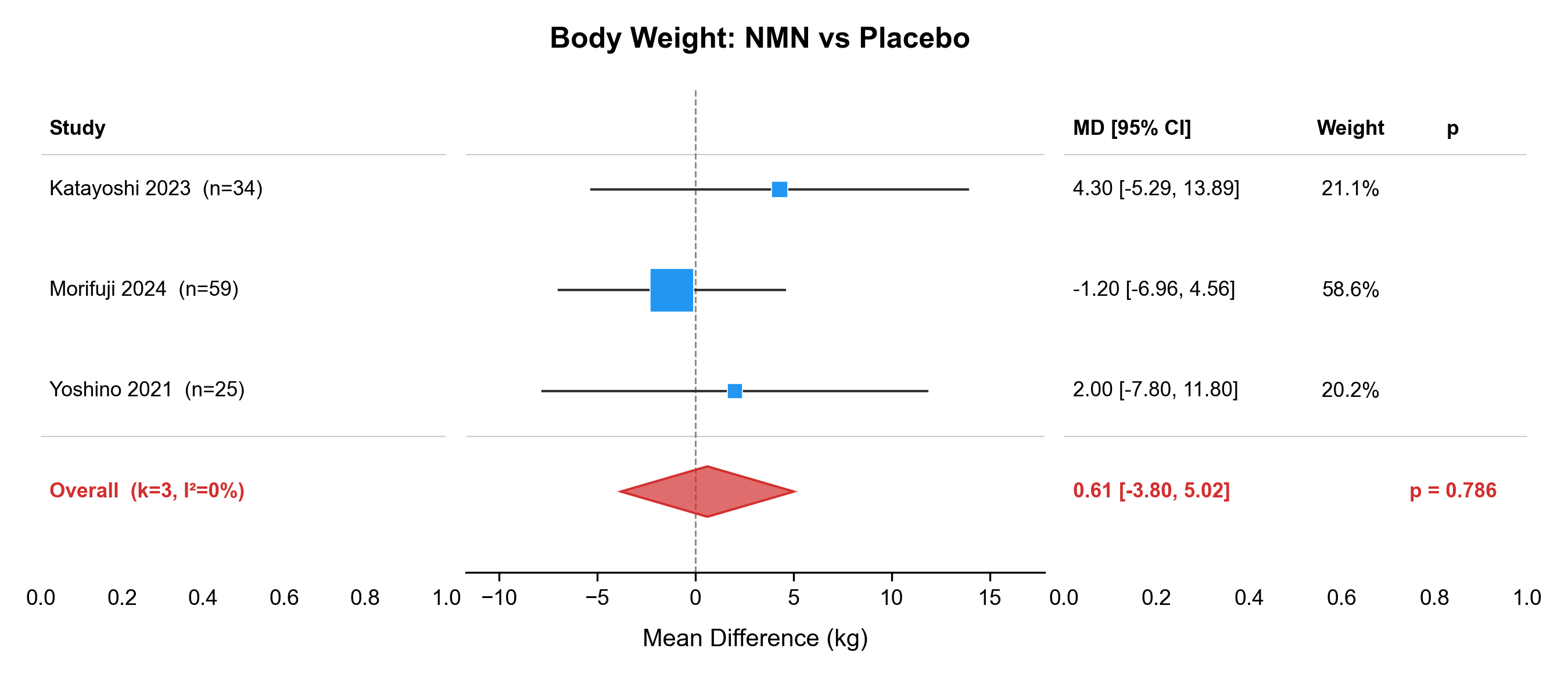

### forest_DBP_NMN_vs_PBO.pdf

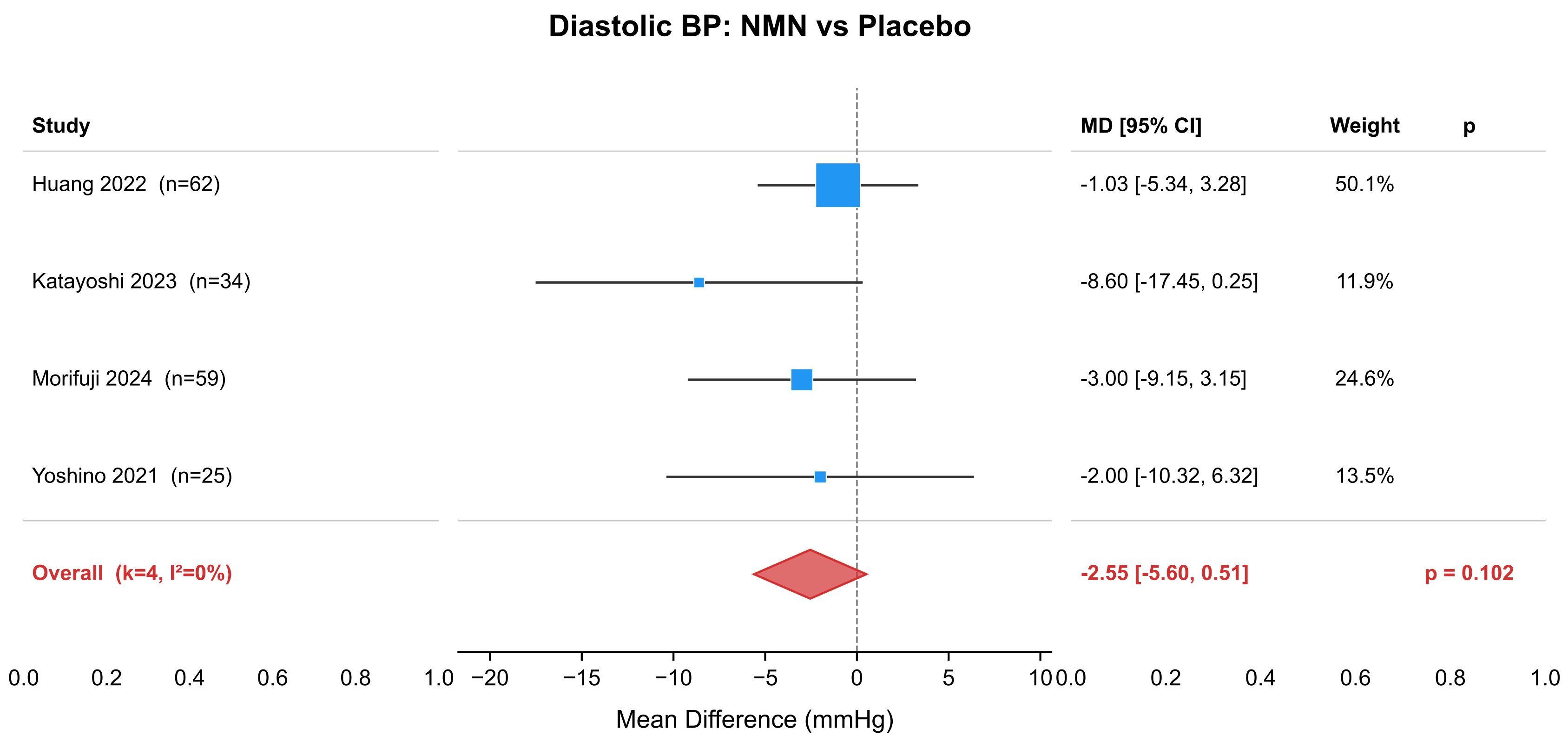

### forest_DBP_NMN_vs_PBO.png

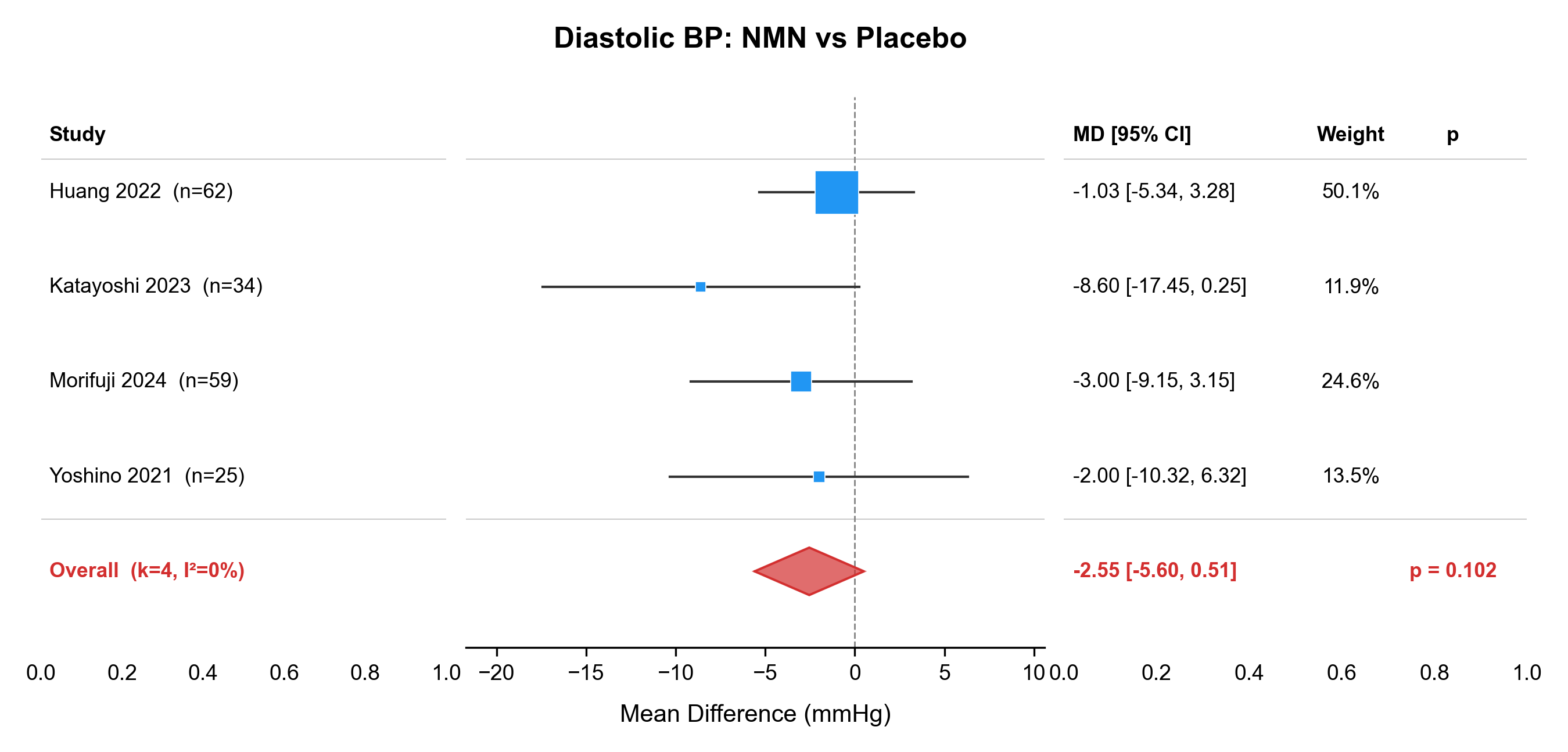

### forest_fasting_insulin_NMN_vs_PBO.pdf

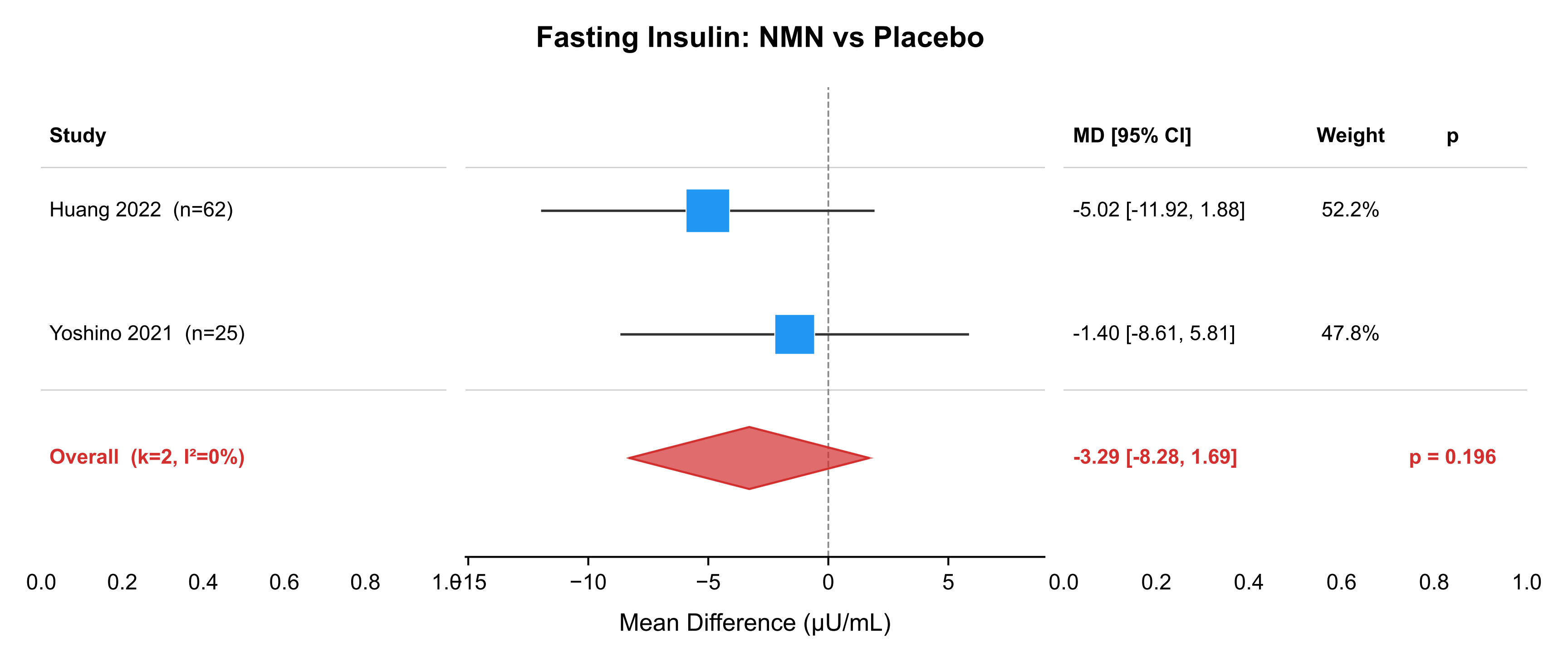

### forest_fasting_insulin_NMN_vs_PBO.png

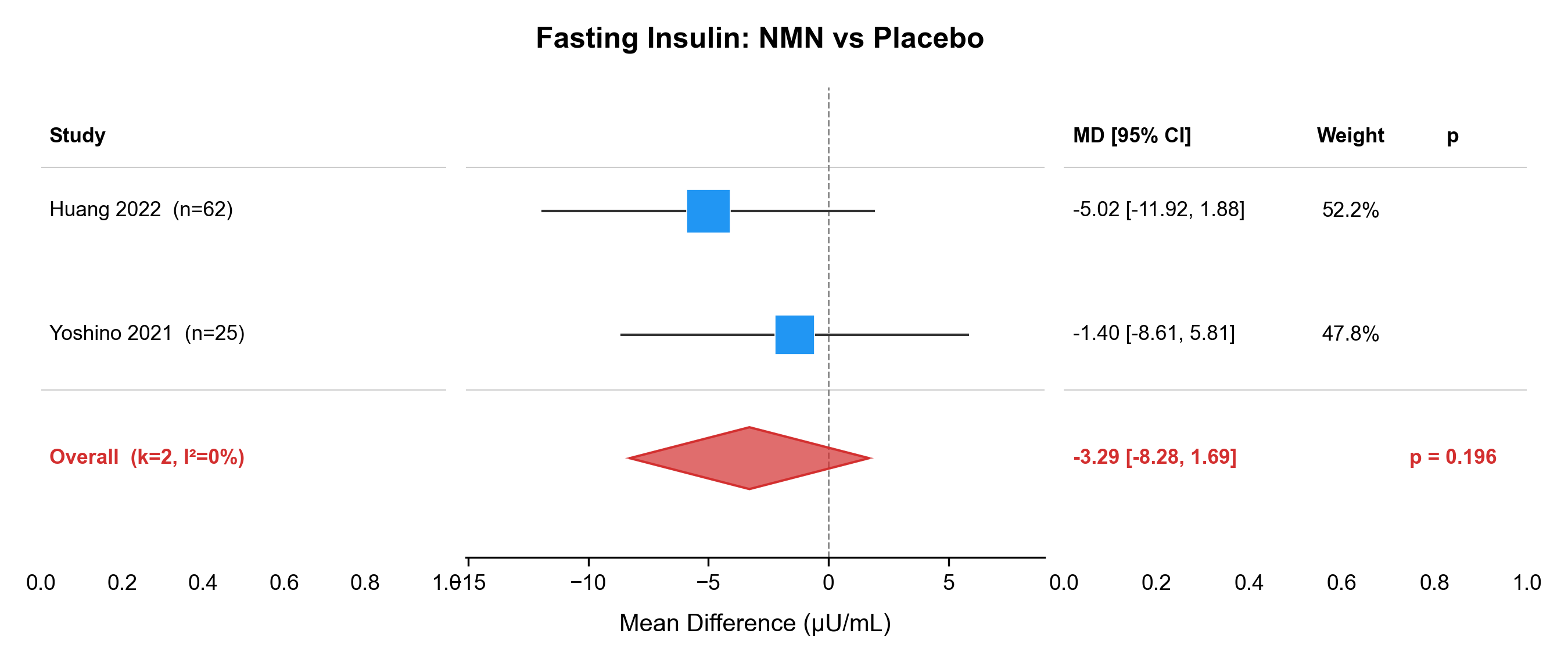

### forest_FBG_NMN_vs_PBO.pdf

#### Fasting Blood Glucose: NMN vs Placebo

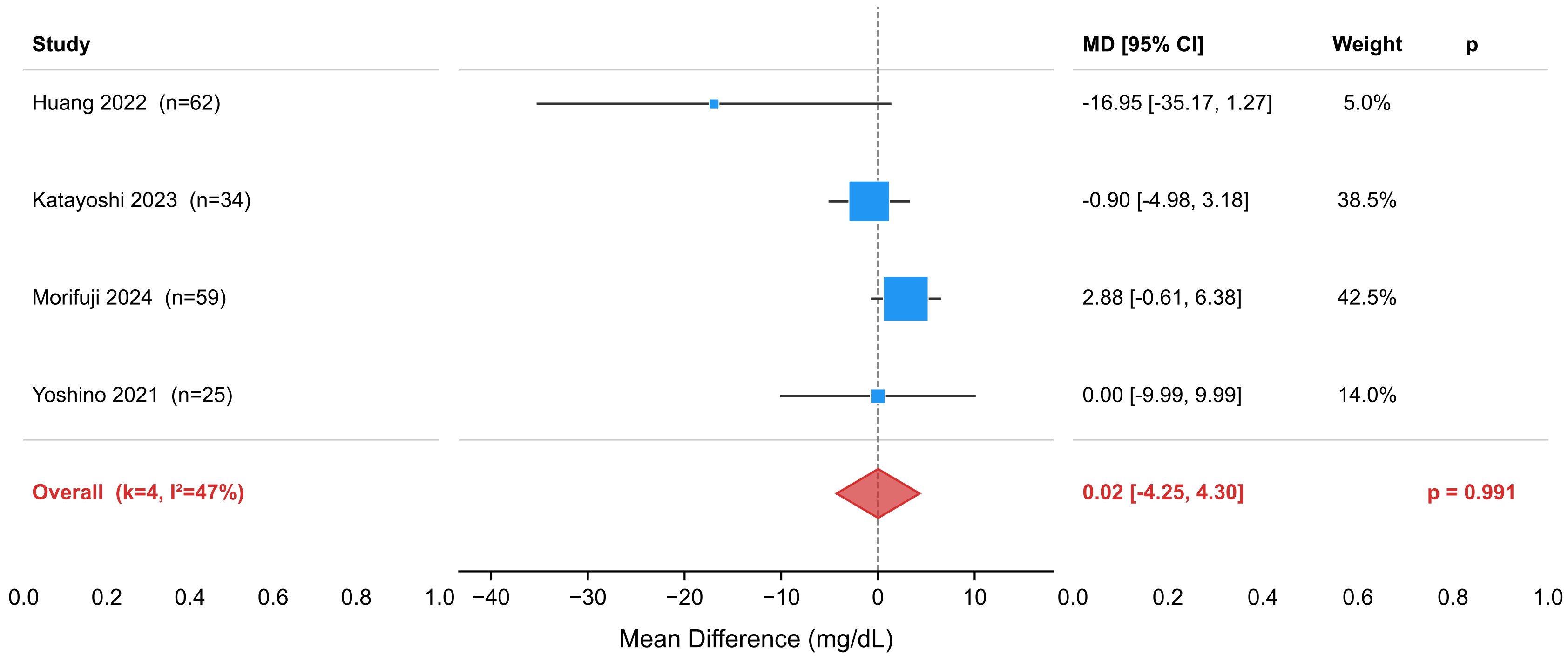

### forest_FBG_NMN_vs_PBO.png

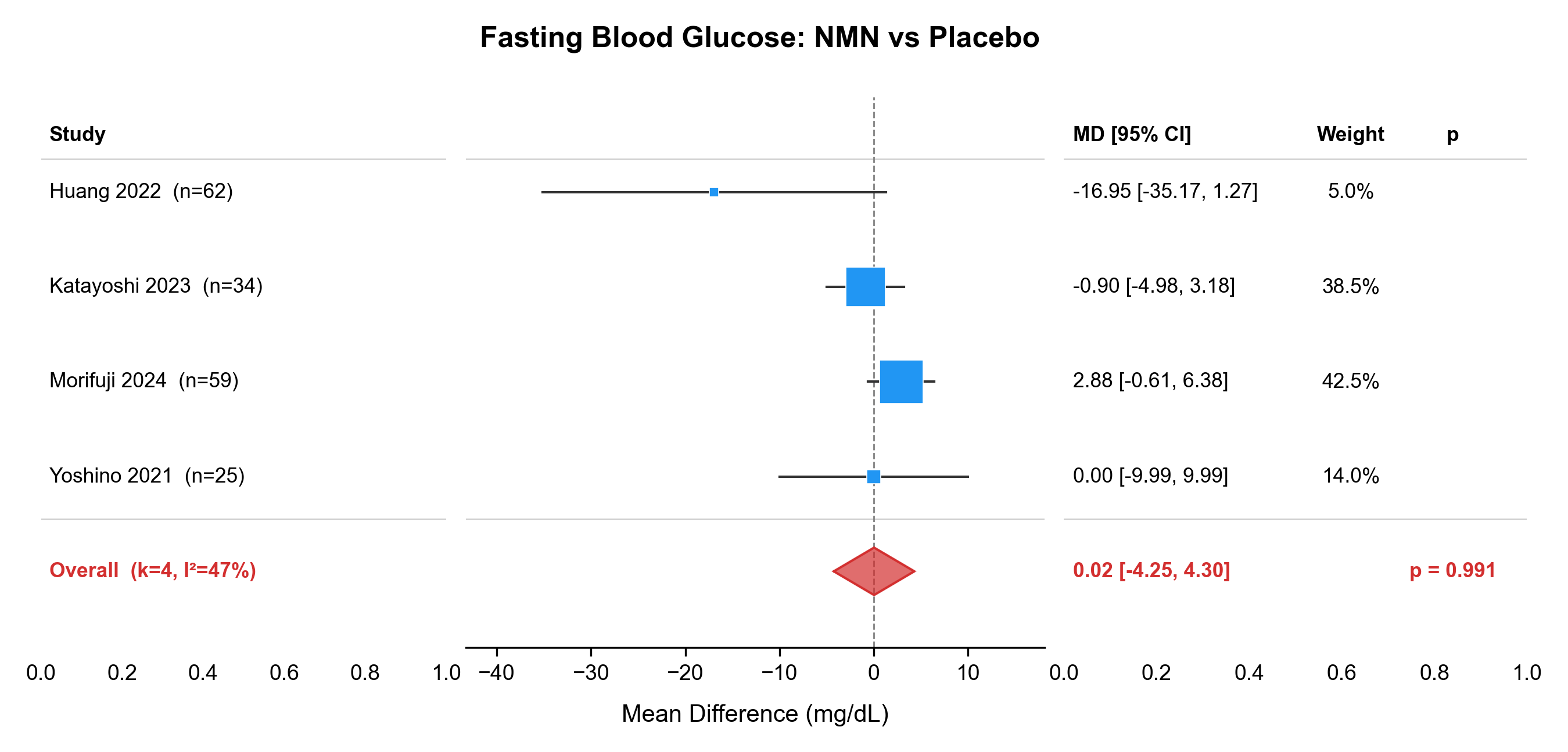

### forest_FBG_NR_vs_PBO.pdf

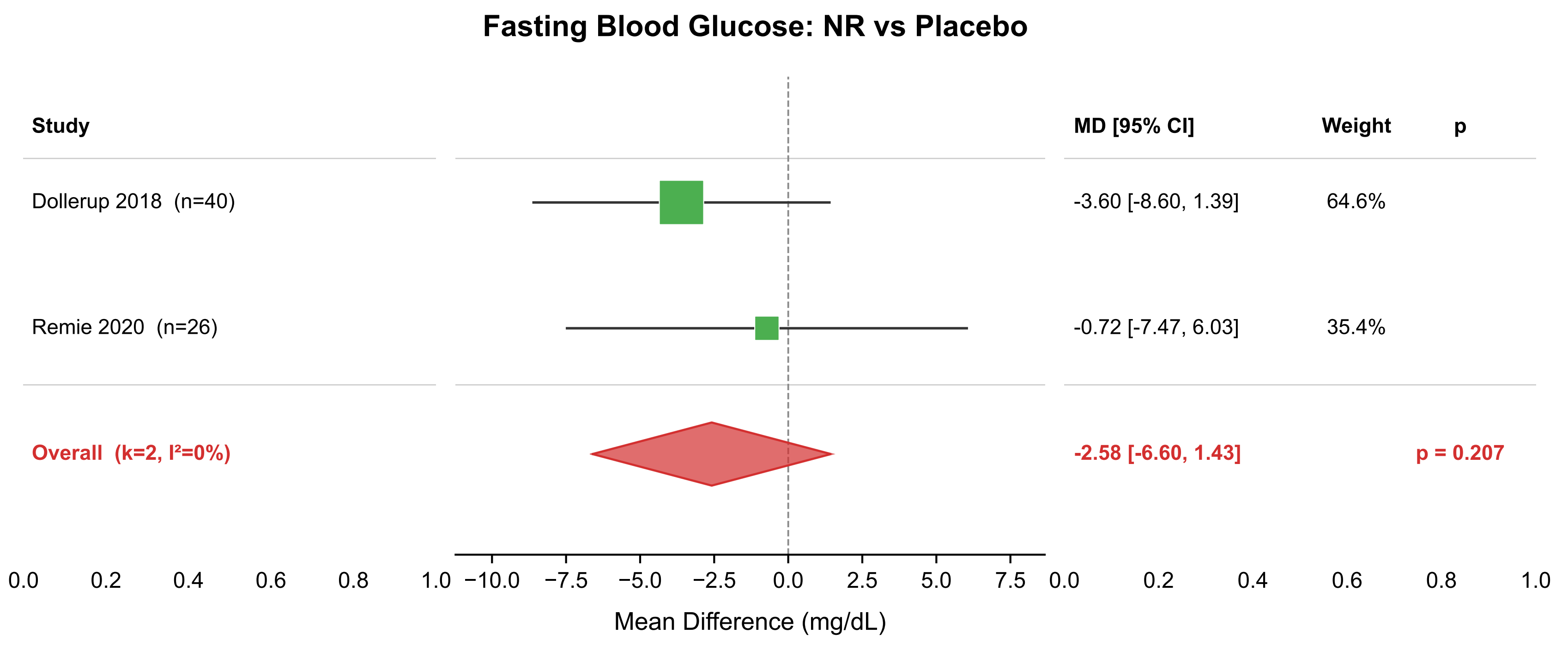

### forest_FBG_NR_vs_PBO.png

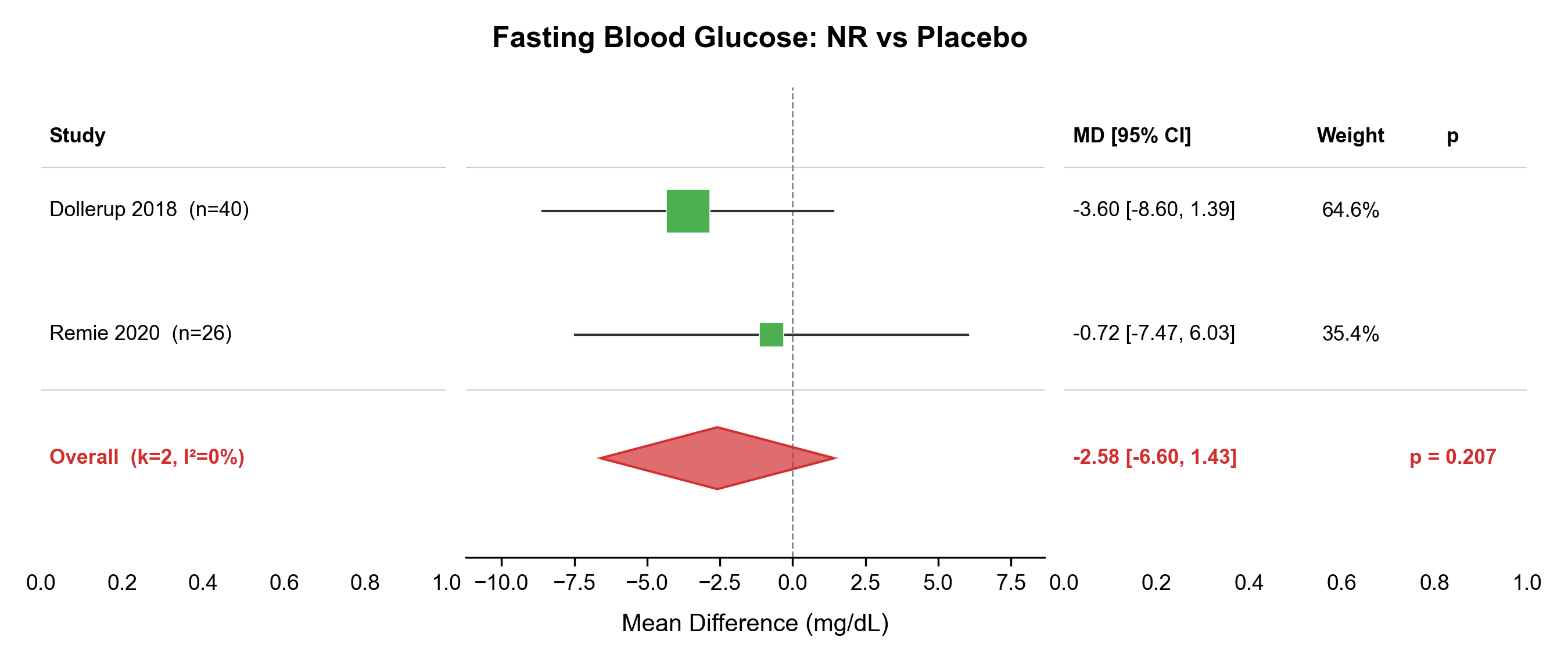

### forest_HbA1c_NMN_vs_PBO.pdf

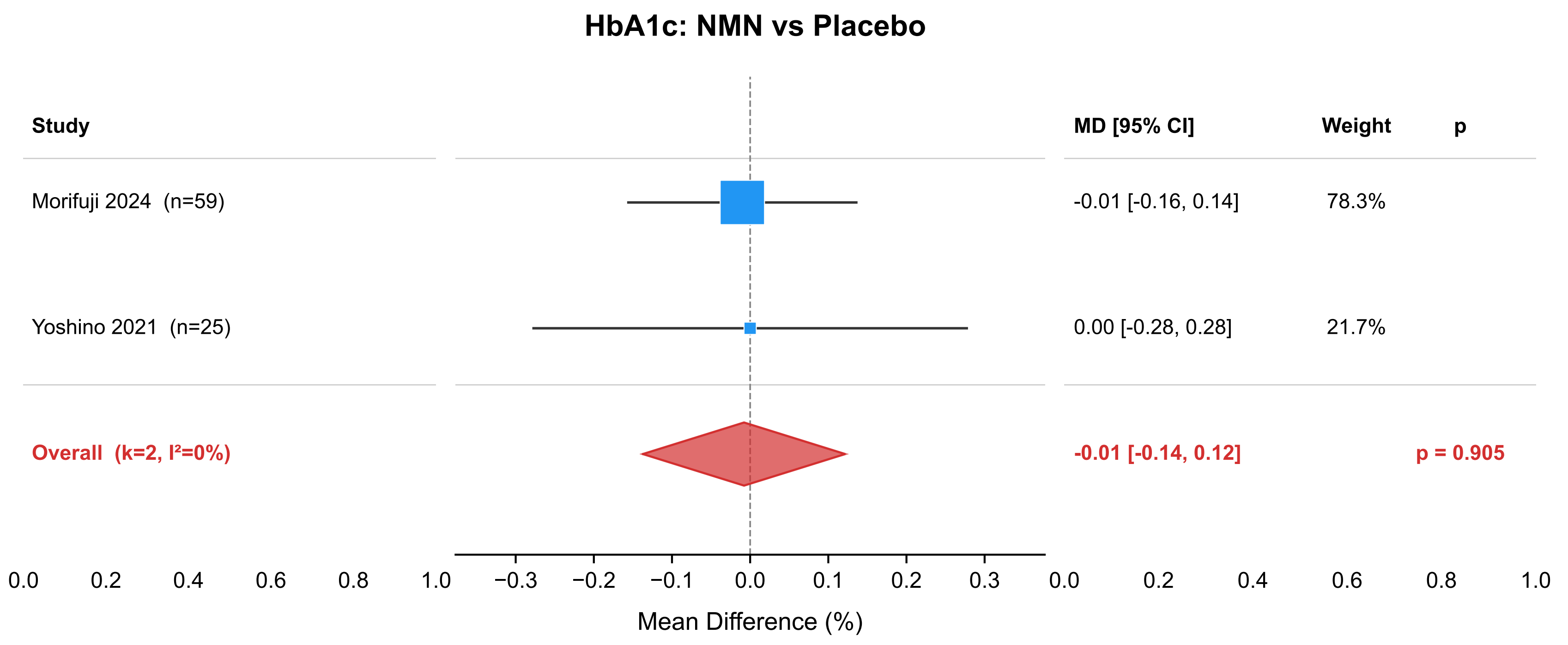

### forest_HbA1c_NMN_vs_PBO.png

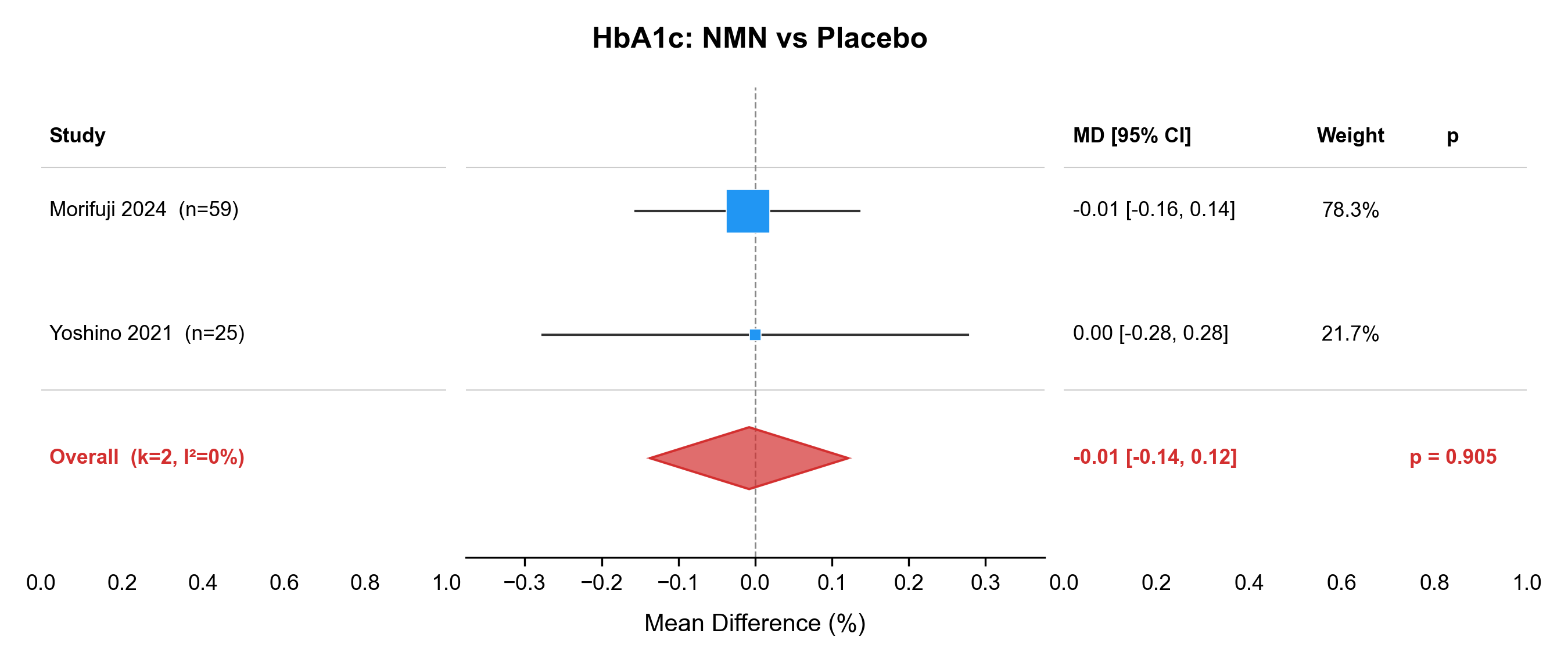

### forest_HDL_NMN_vs_PBO.pdf

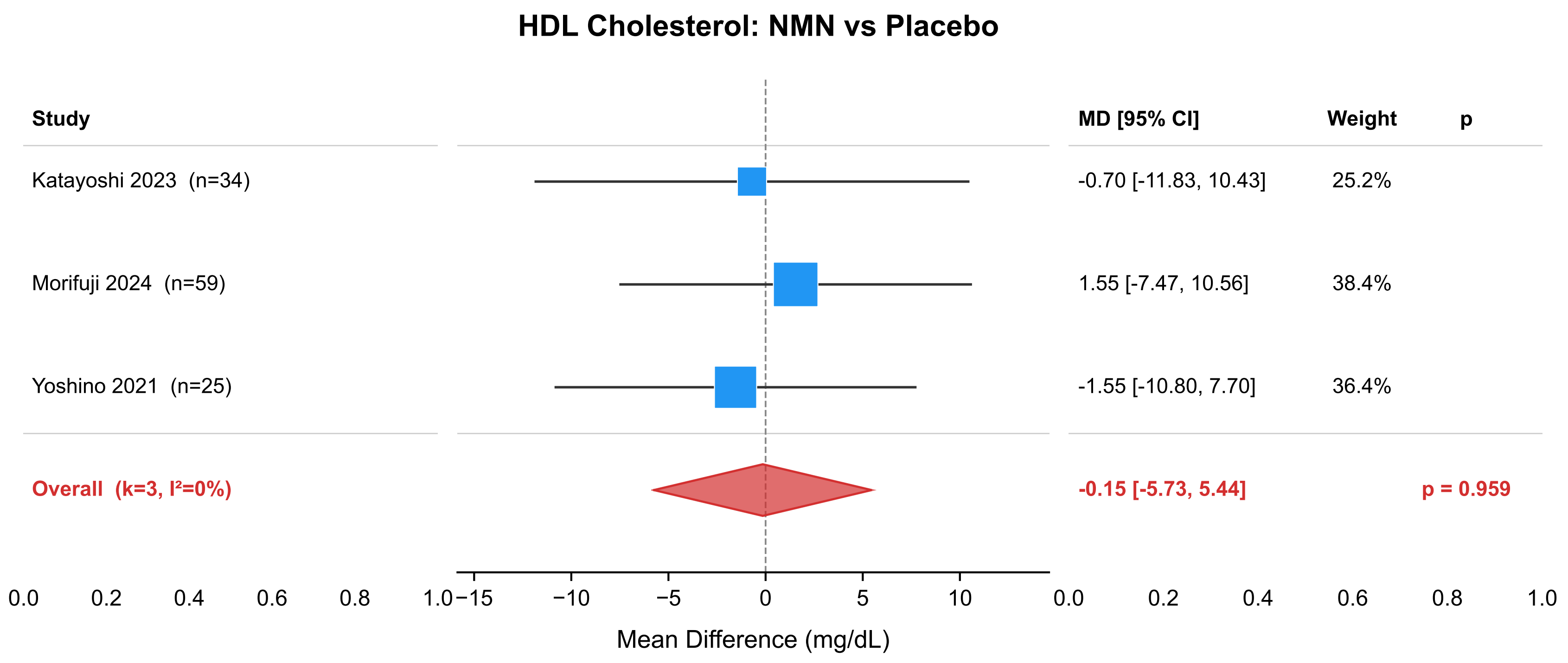

### forest_HDL_NMN_vs_PBO.png

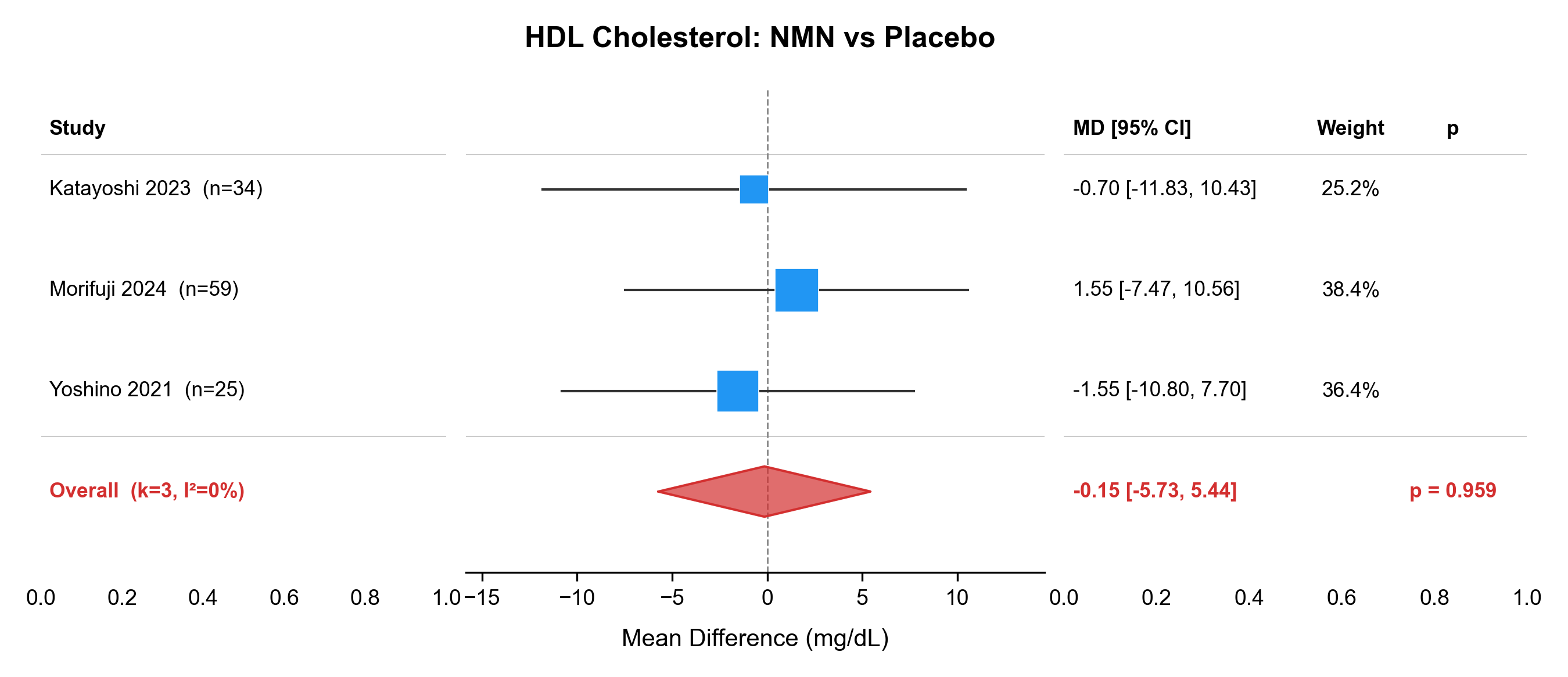

### forest_HDL_NR_vs_PBO.pdf

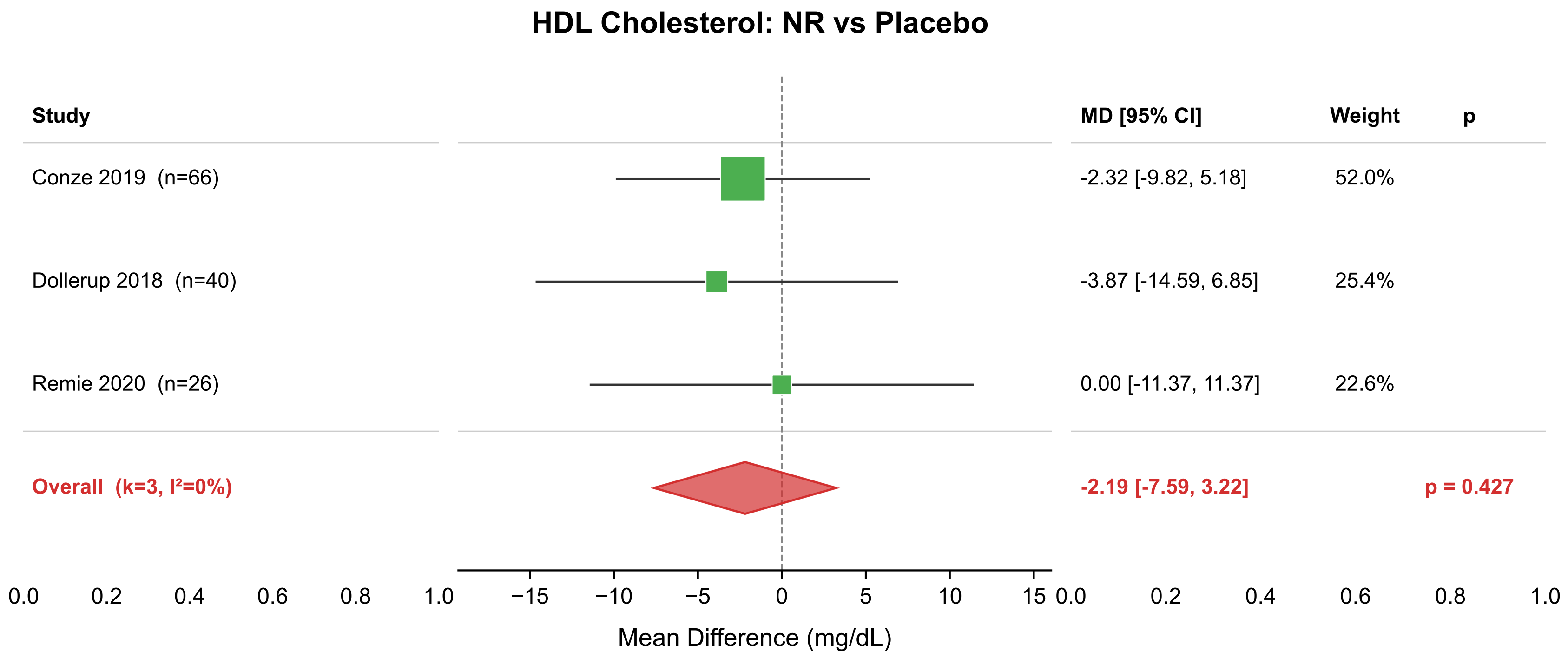

### forest_HDL_NR_vs_PBO.png

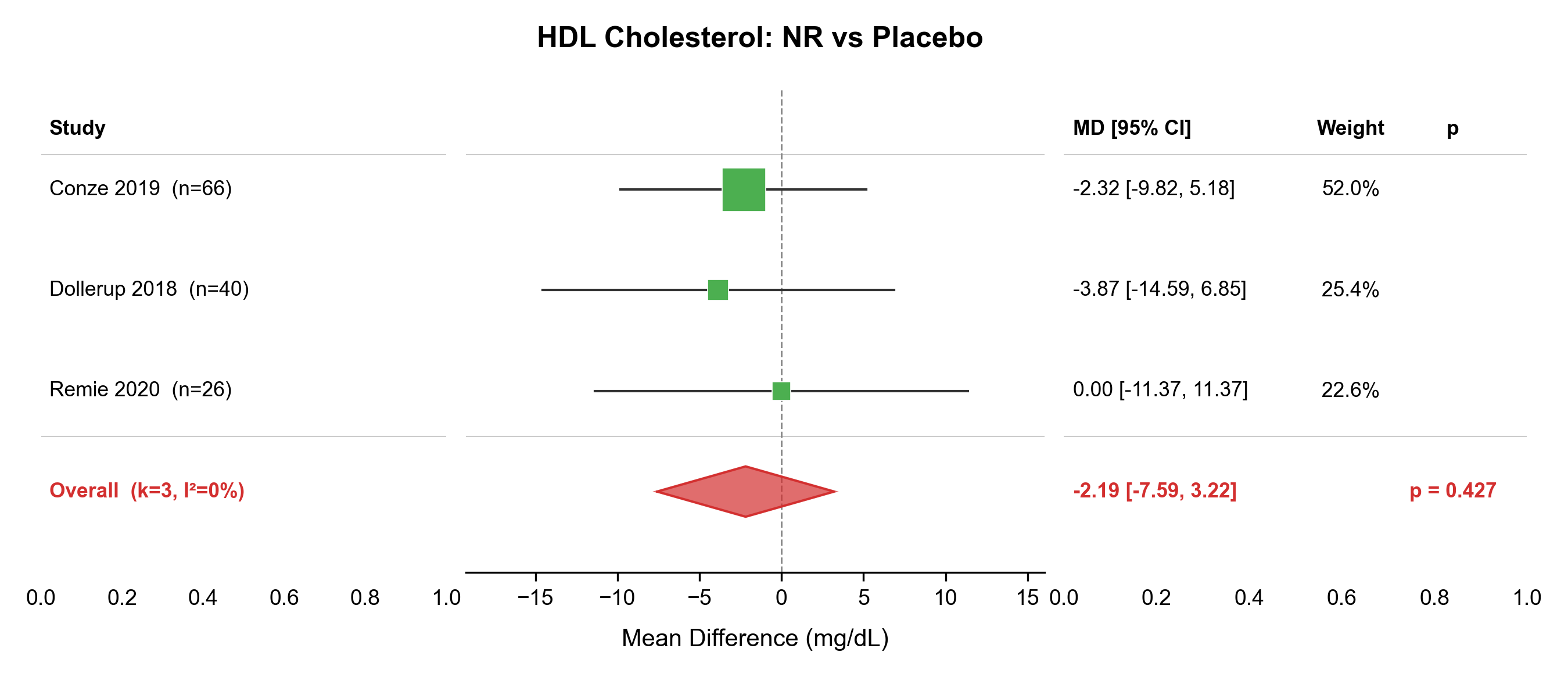

### forest_LDL_NMN_vs_PBO.pdf

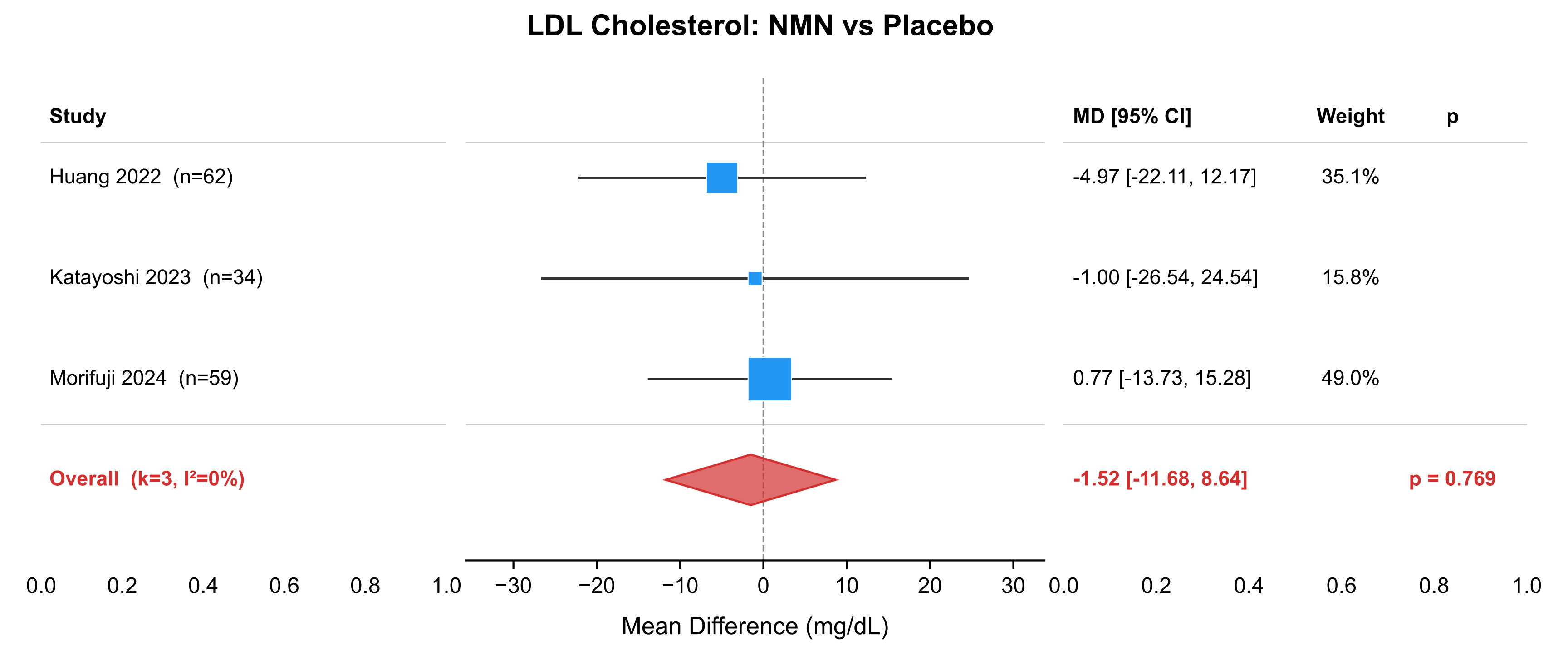

### forest_LDL_NMN_vs_PBO.png

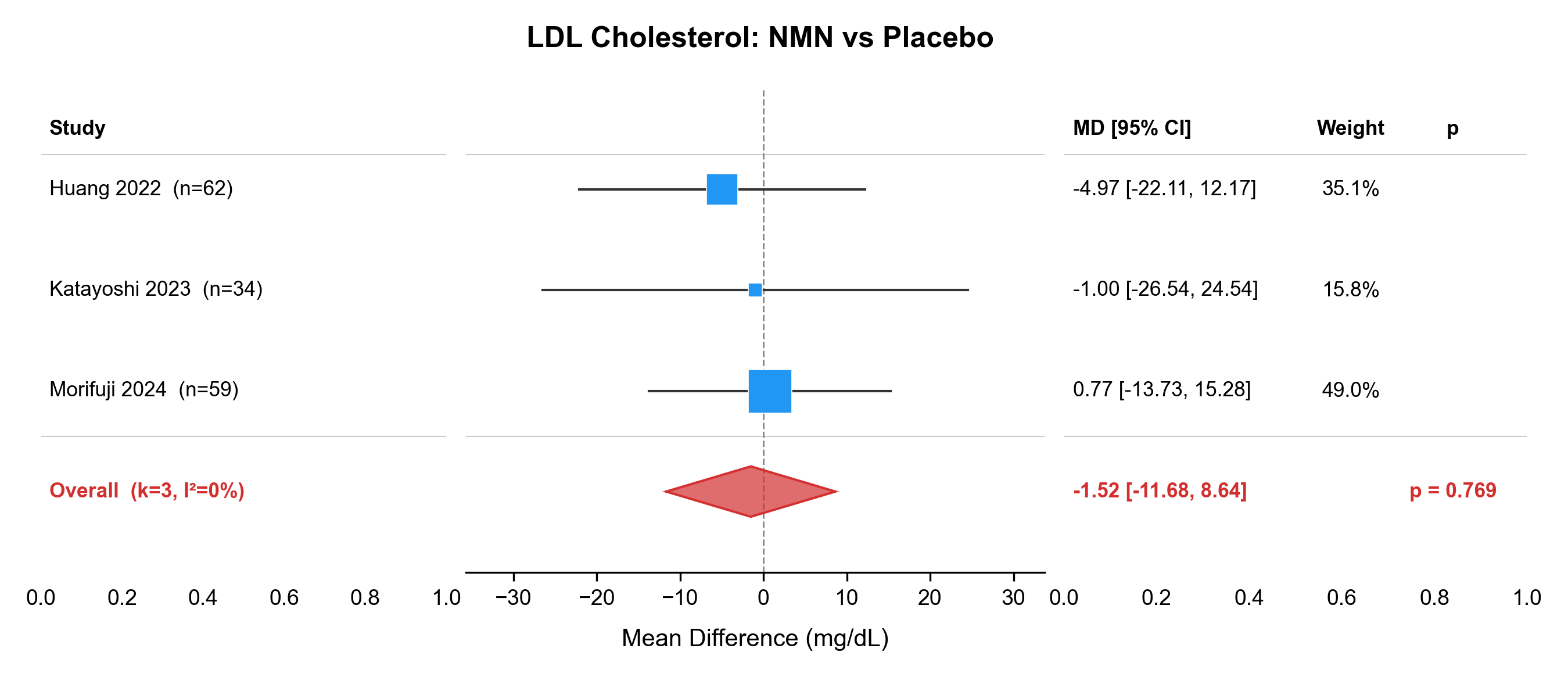

### forest_LDL_NR_vs_PBO.pdf

#### LDL Cholesterol: NR vs Placebo

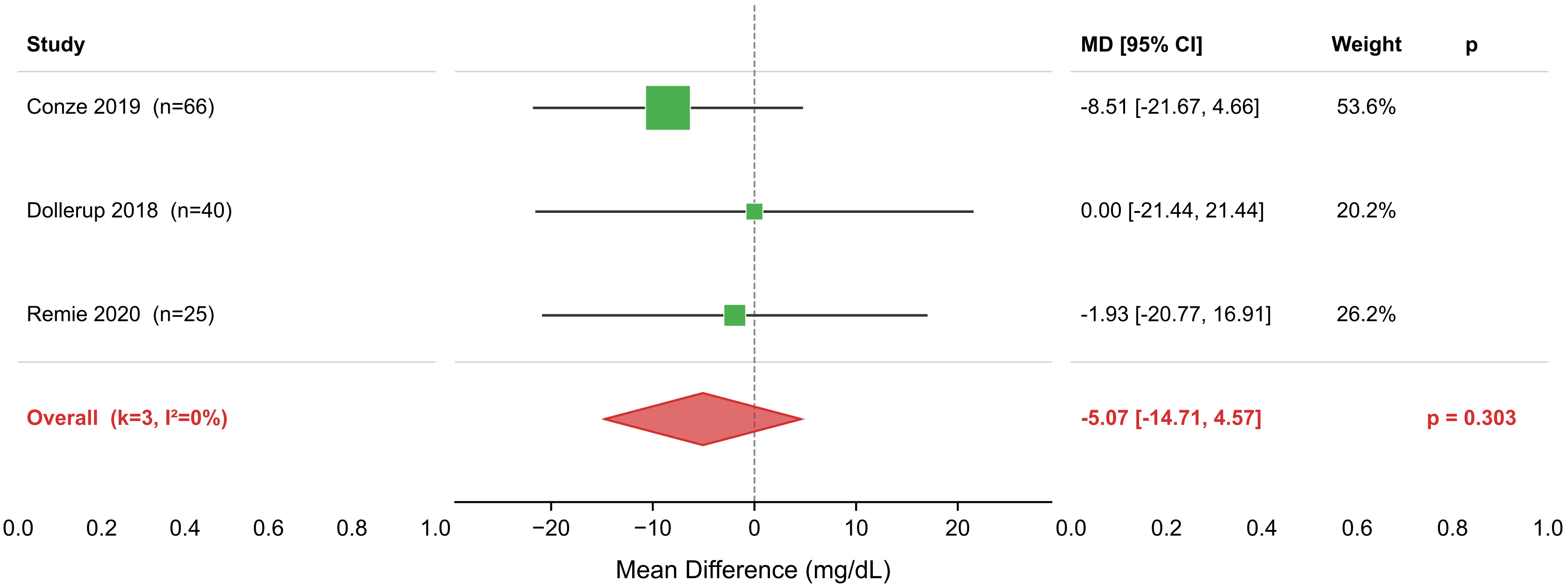

### forest_LDL_NR_vs_PBO.png

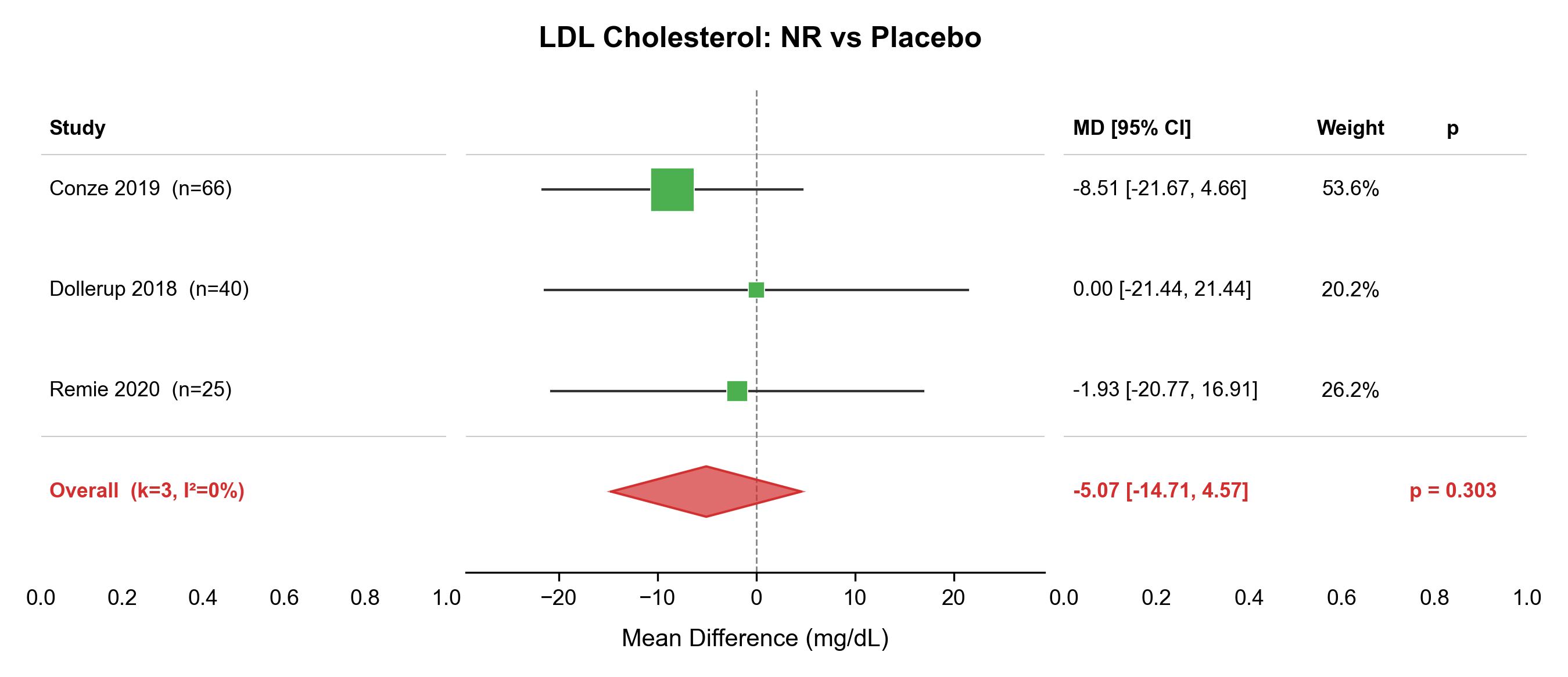

### forest_SBP_NMN_vs_PBO.pdf

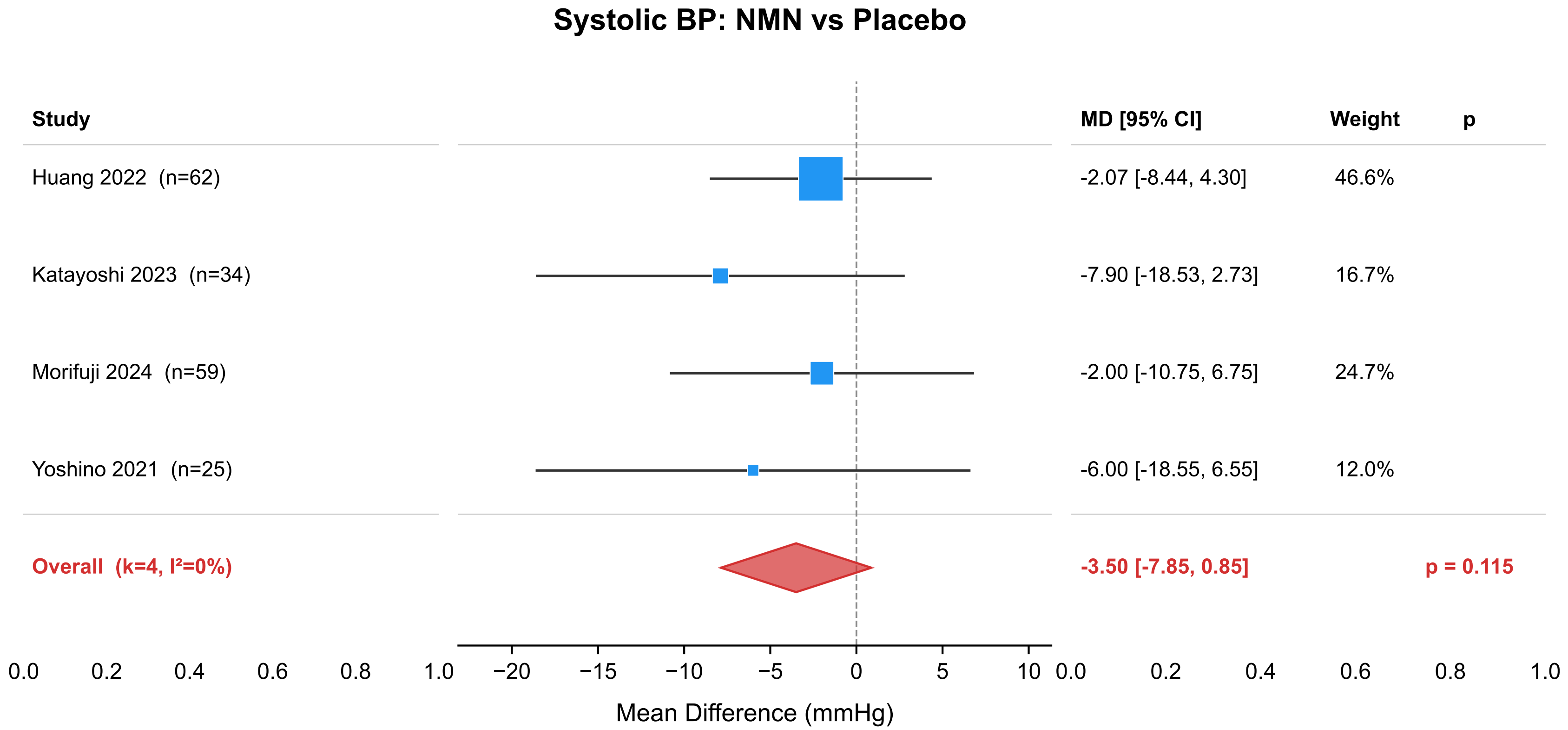

### forest_SBP_NMN_vs_PBO.png

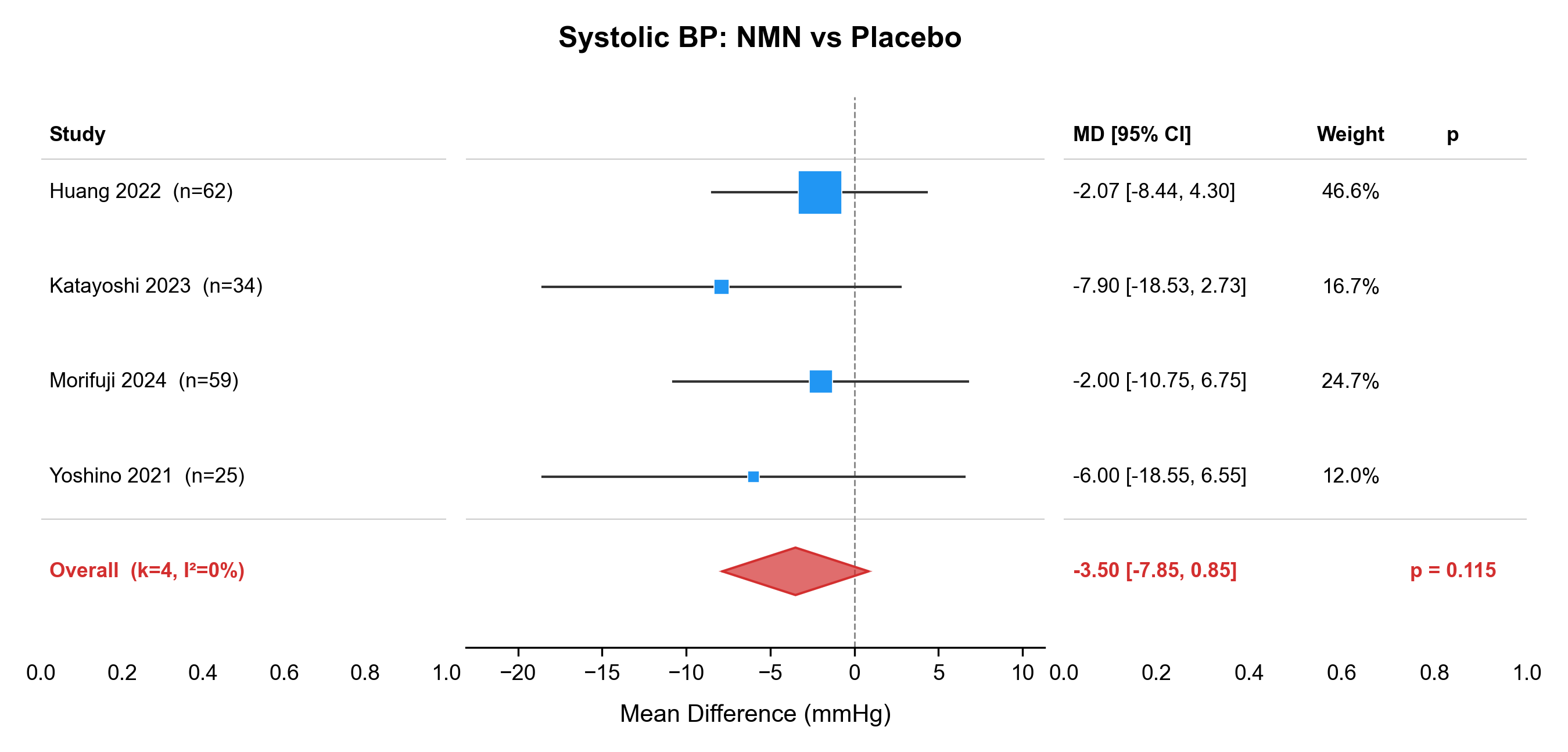

### forest_TC_NMN_vs_PBO.pdf

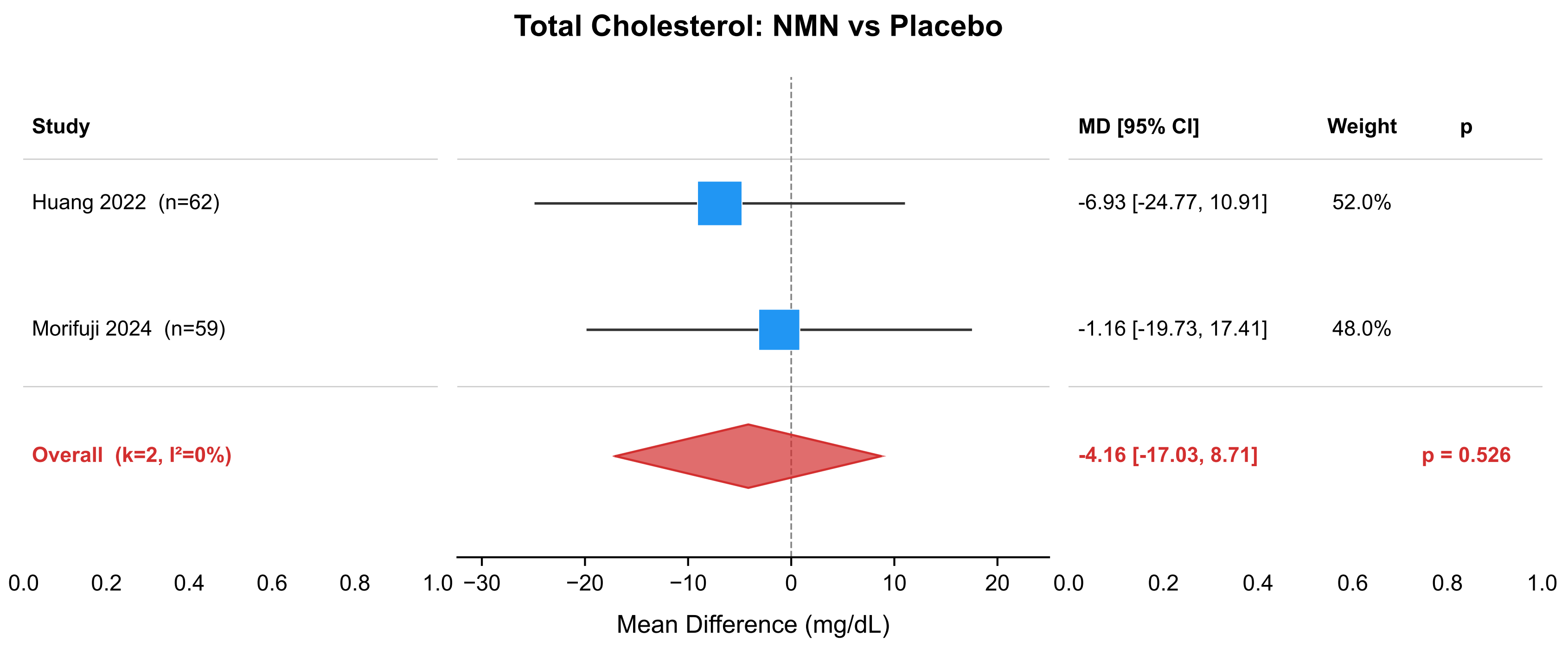

### forest_TC_NMN_vs_PBO.png

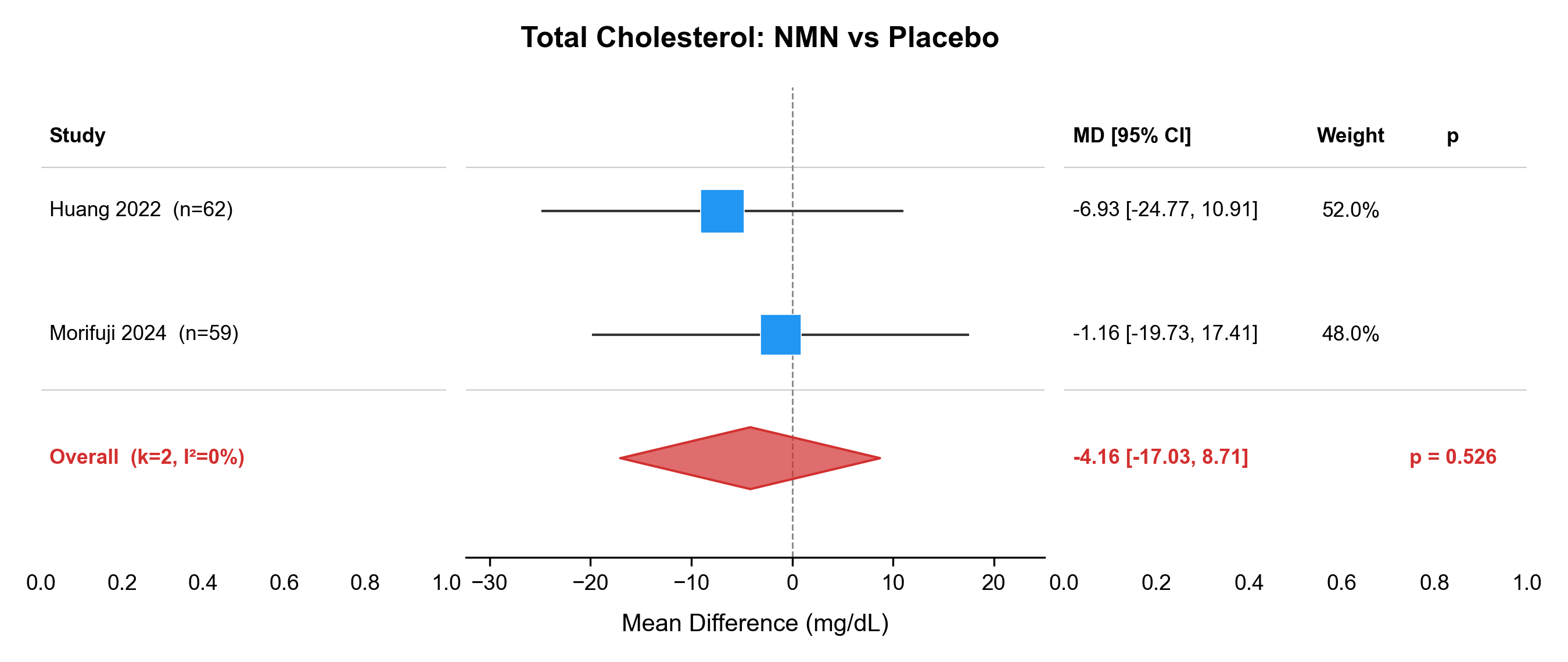

### forest_TC_NR_vs_PBO.pdf

##### Total Cholesterol: NR vs Placebo

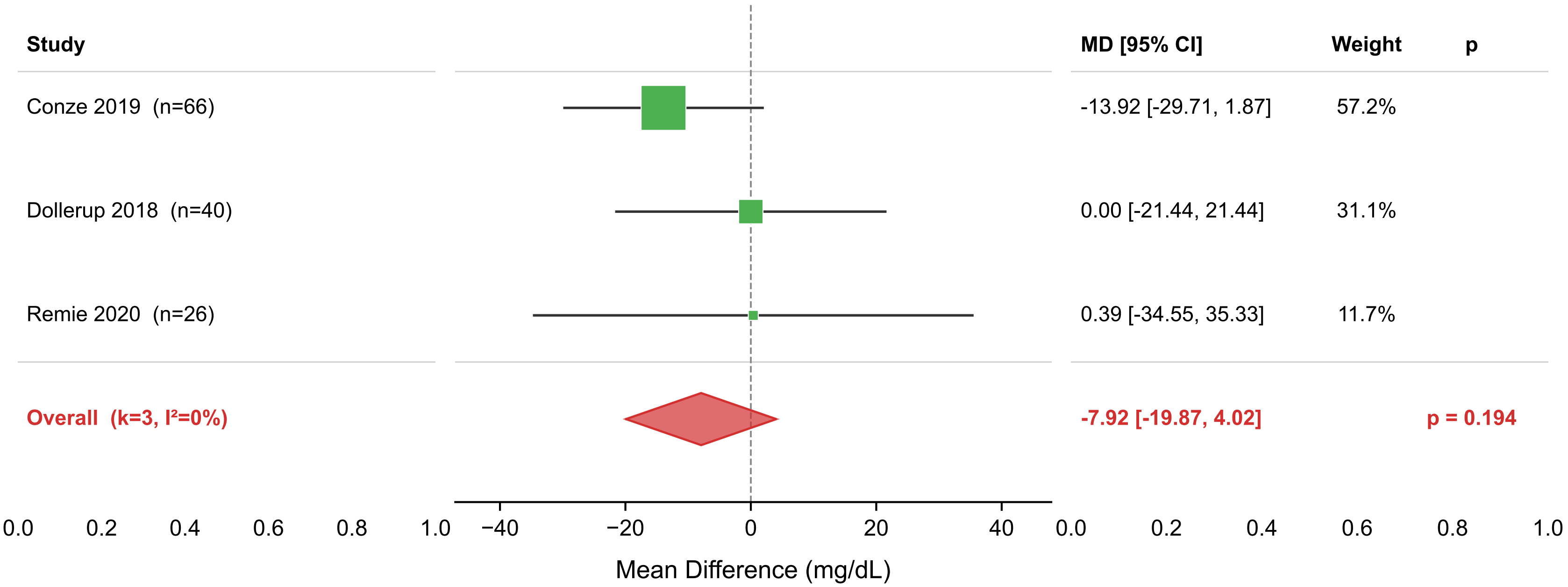

### forest_TC_NR_vs_PBO.png

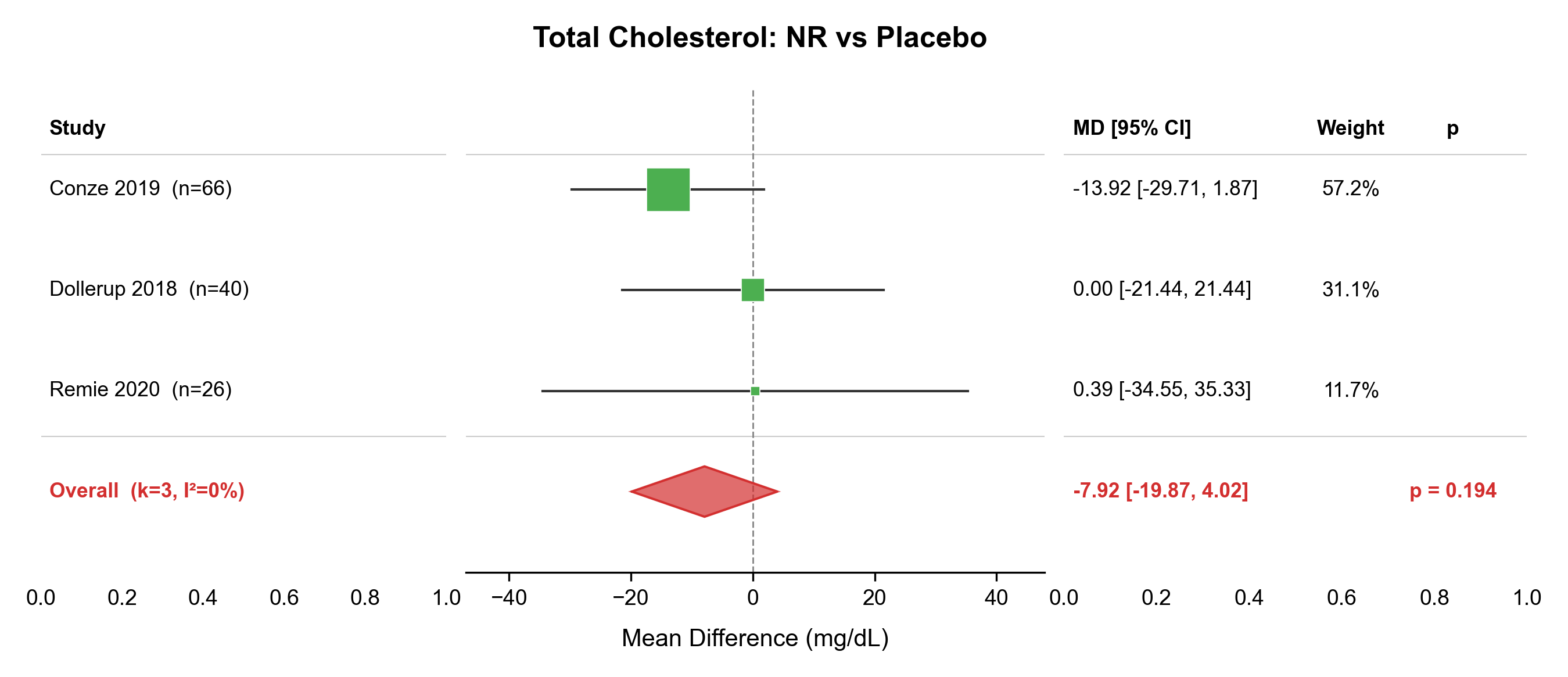

### forest_TG_NMN_vs_PBO.pdf

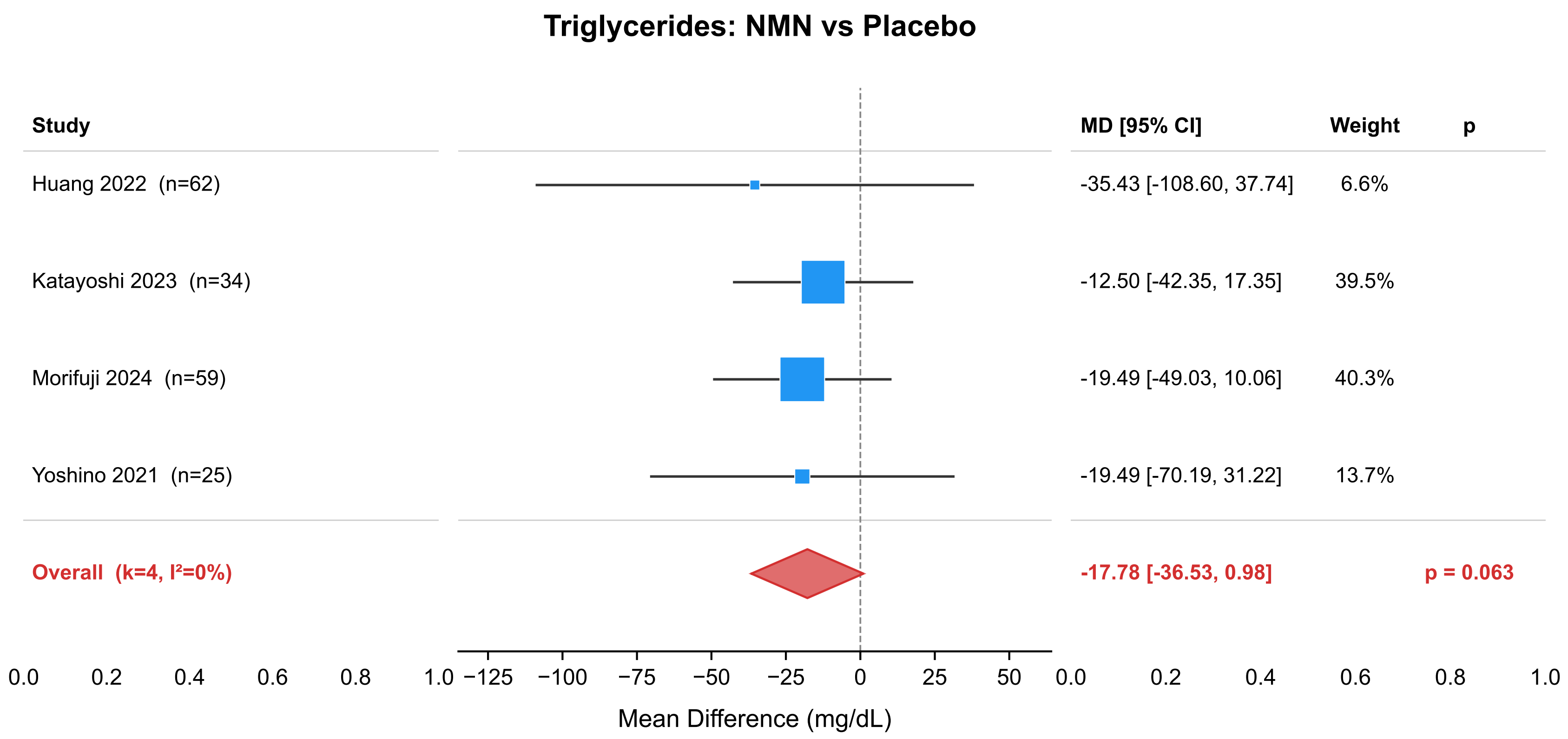

### forest_TG_NMN_vs_PBO.png

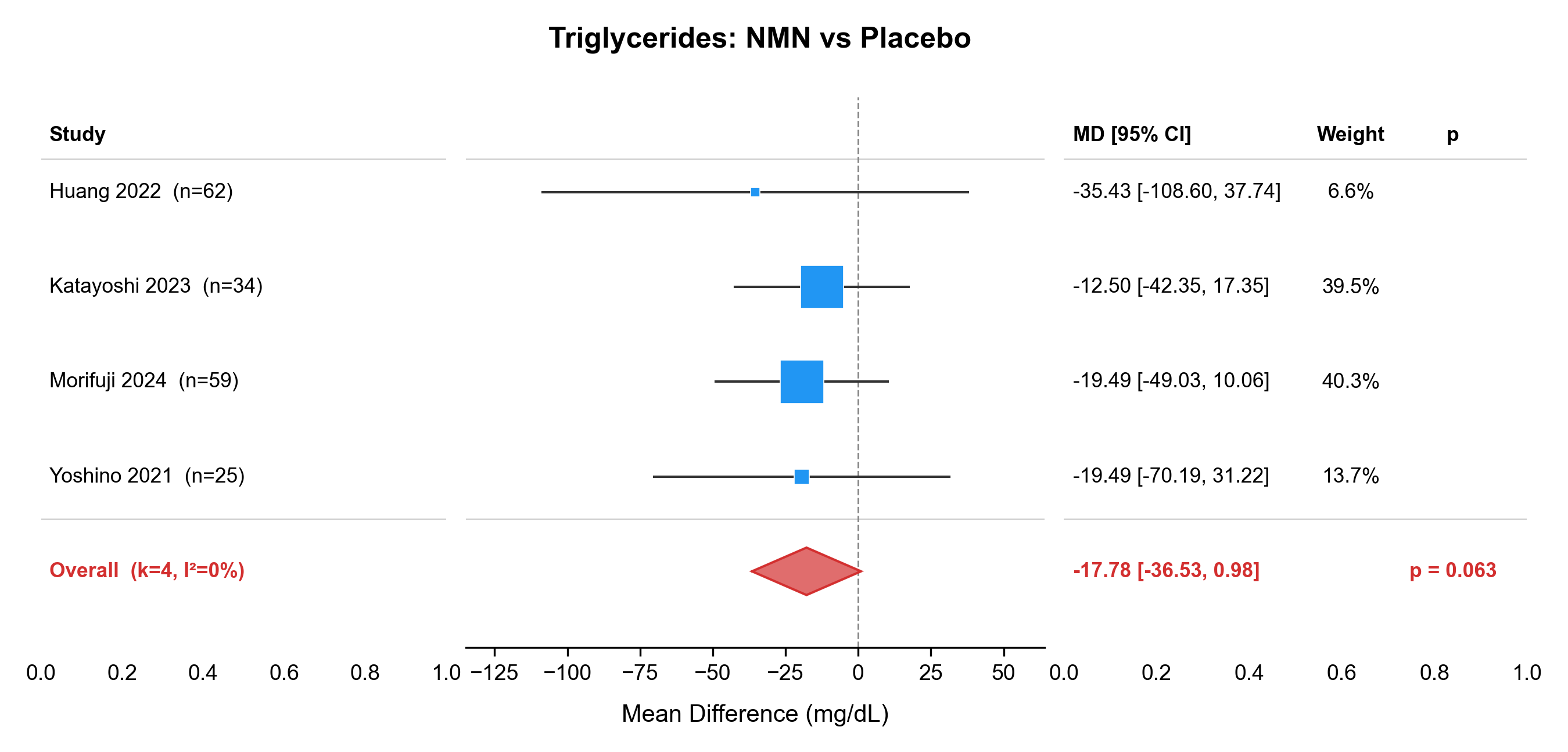

### forest_TG_NR_vs_PBO.pdf

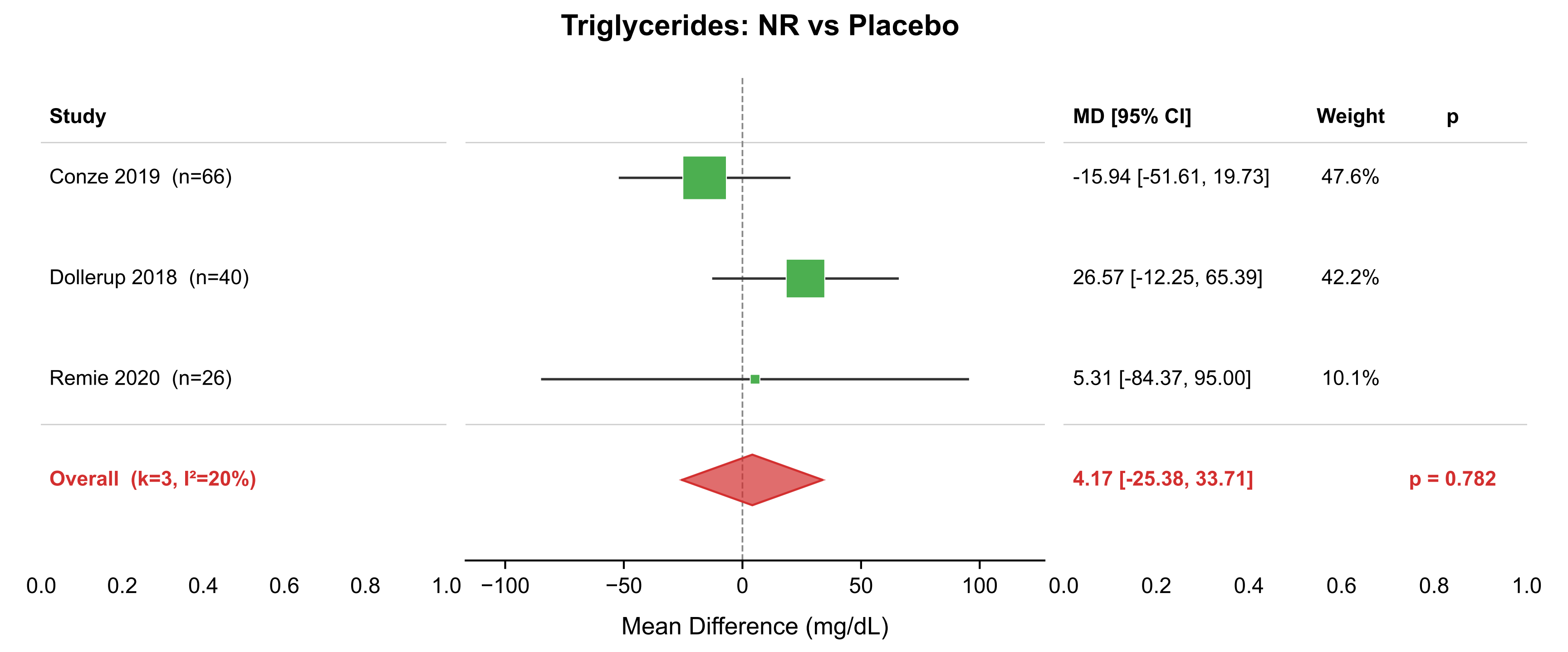

### forest_TG_NR_vs_PBO.png

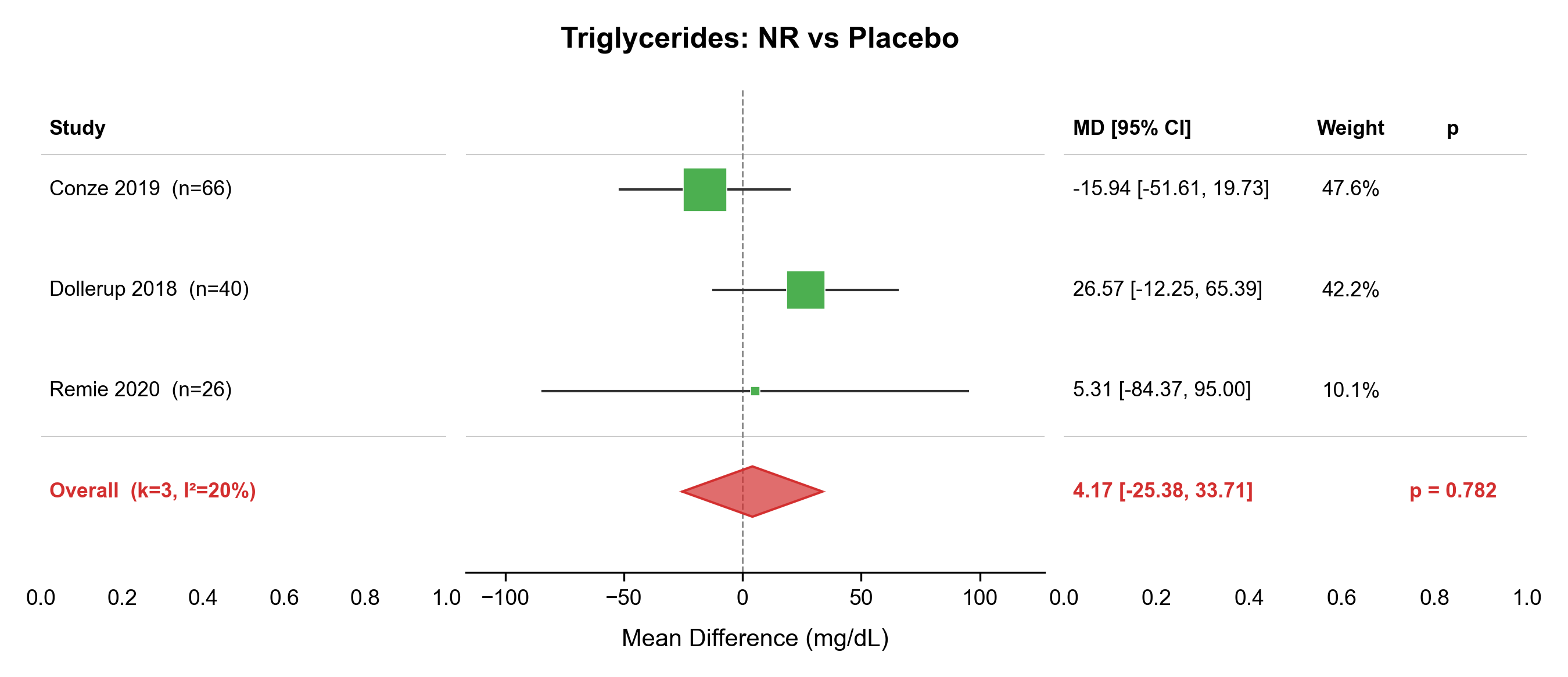

### funnel_ALT_NMN_vs_PBO.png

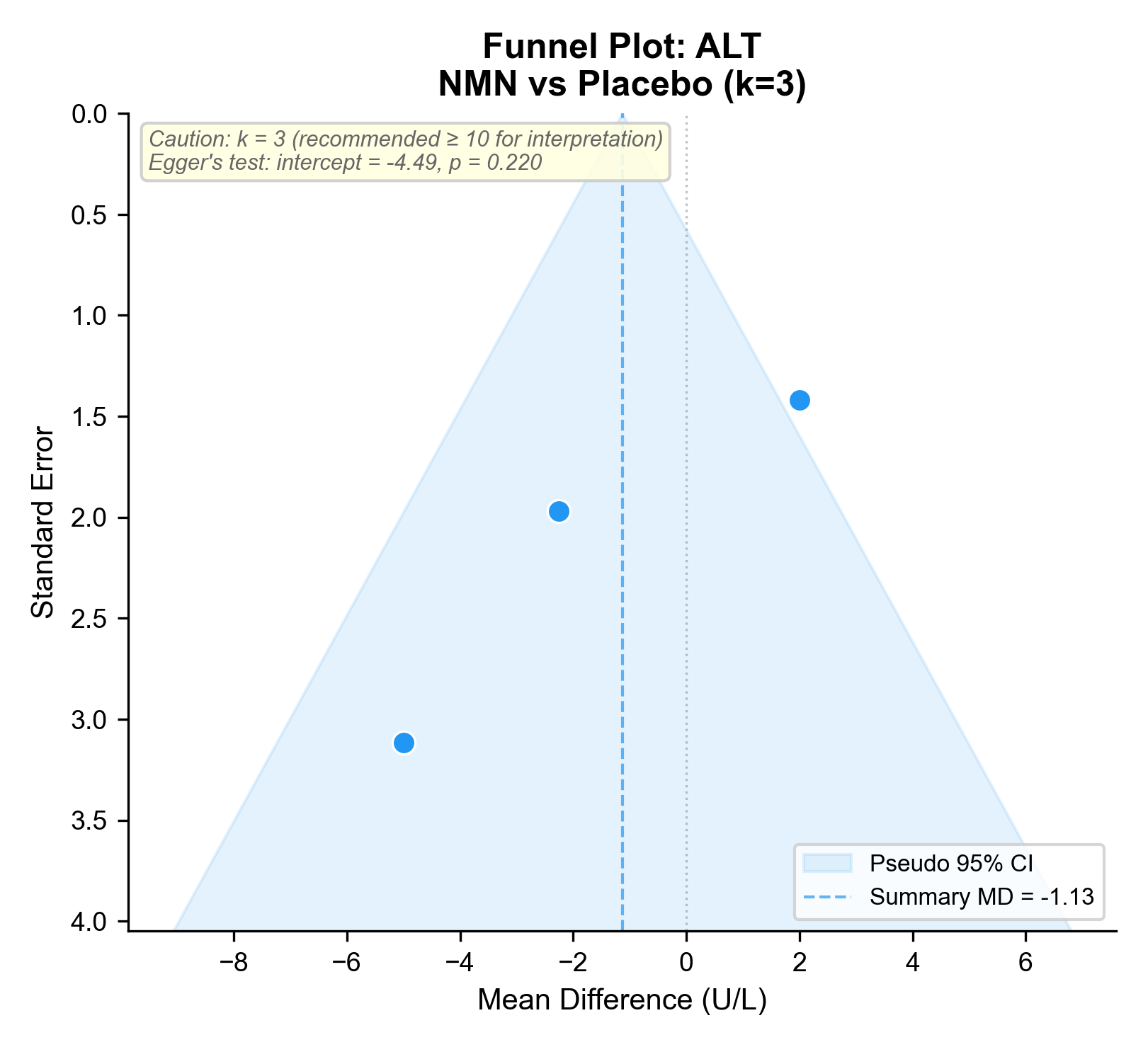

### funnel_AST_NMN_vs_PBO.png

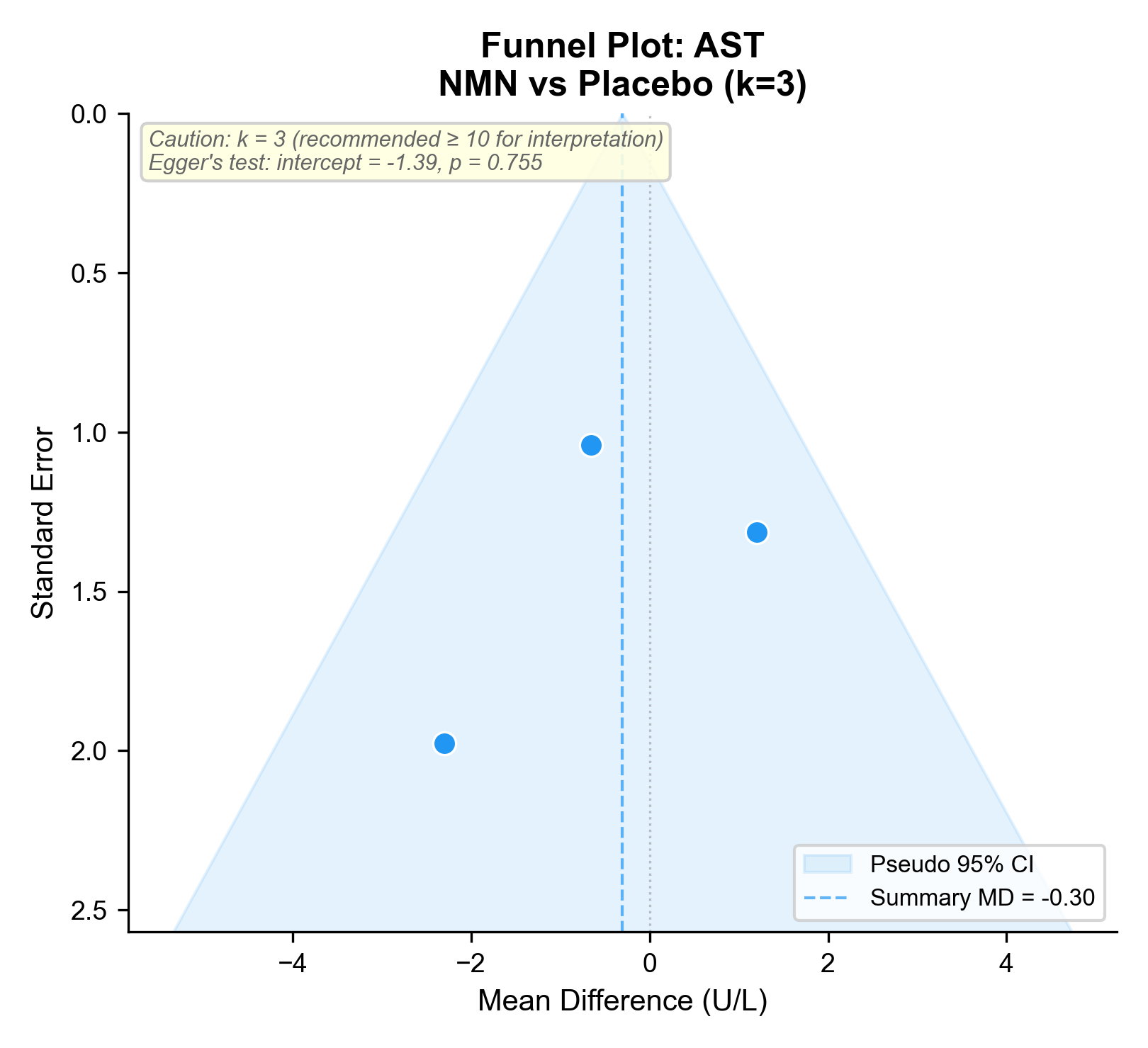

### funnel_BMI_NMN_vs_PBO.png

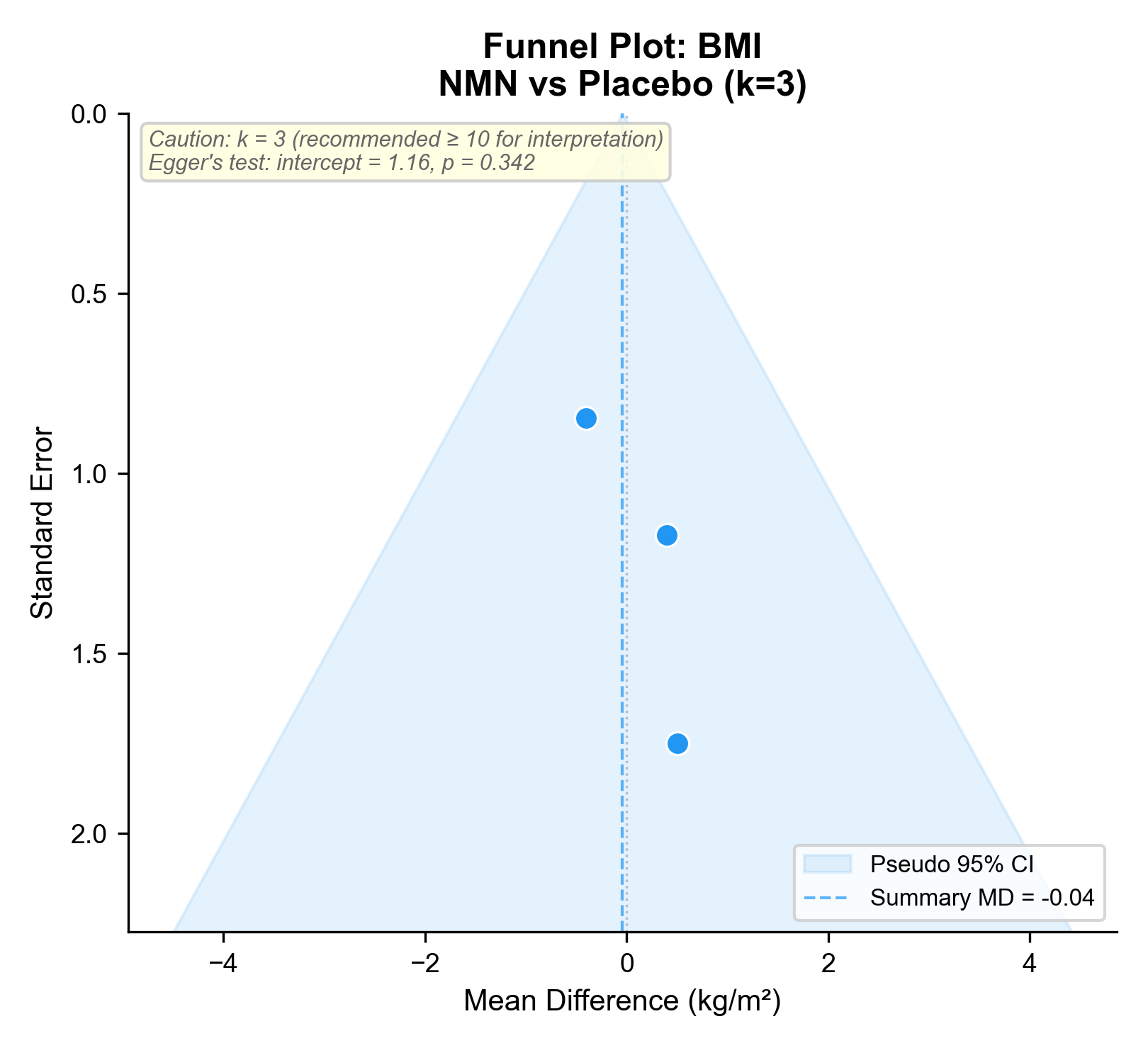

### funnel_body_weight_NMN_vs_PBO.png

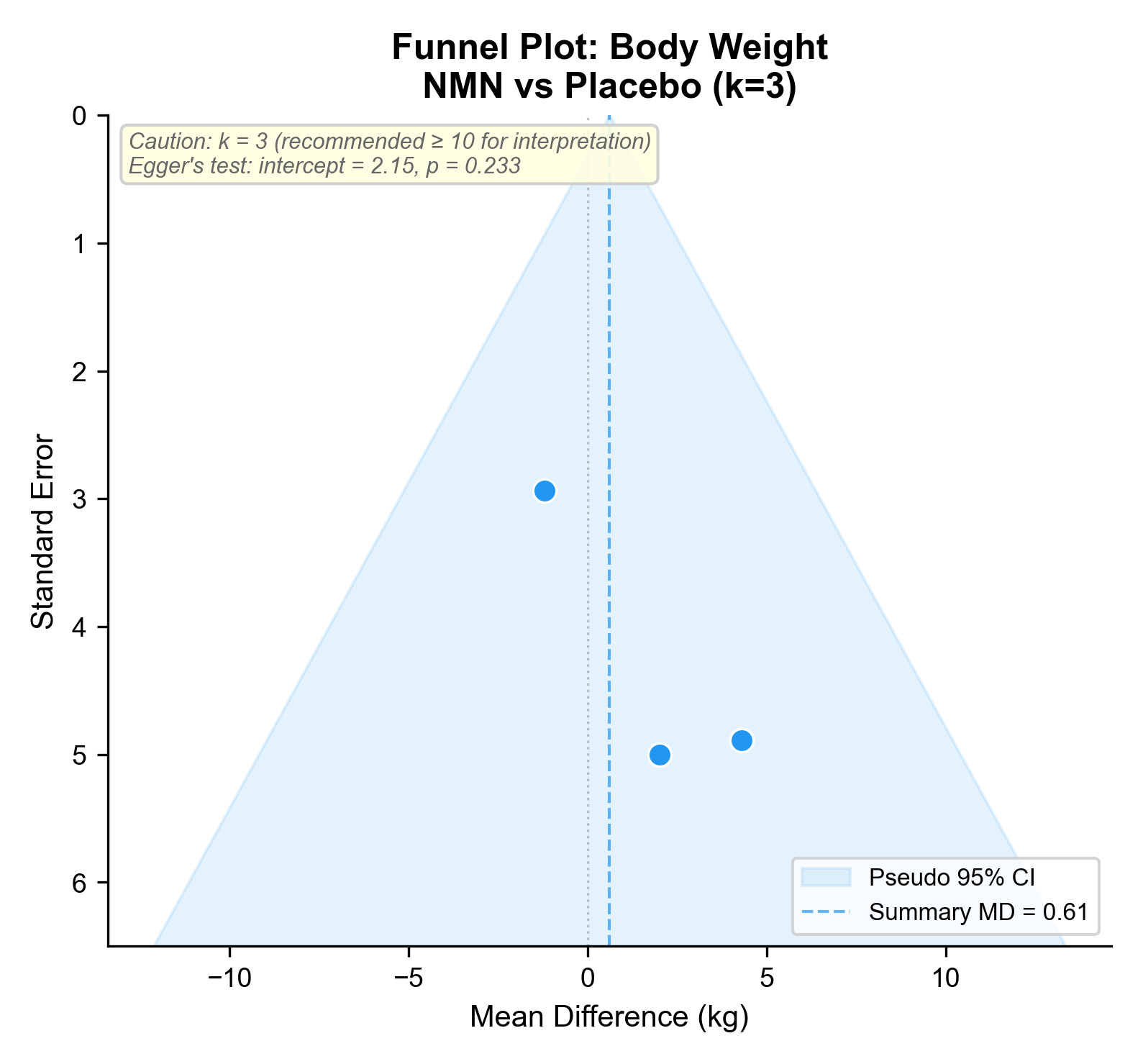

### funnel_DBP_NMN_vs_PBO.png

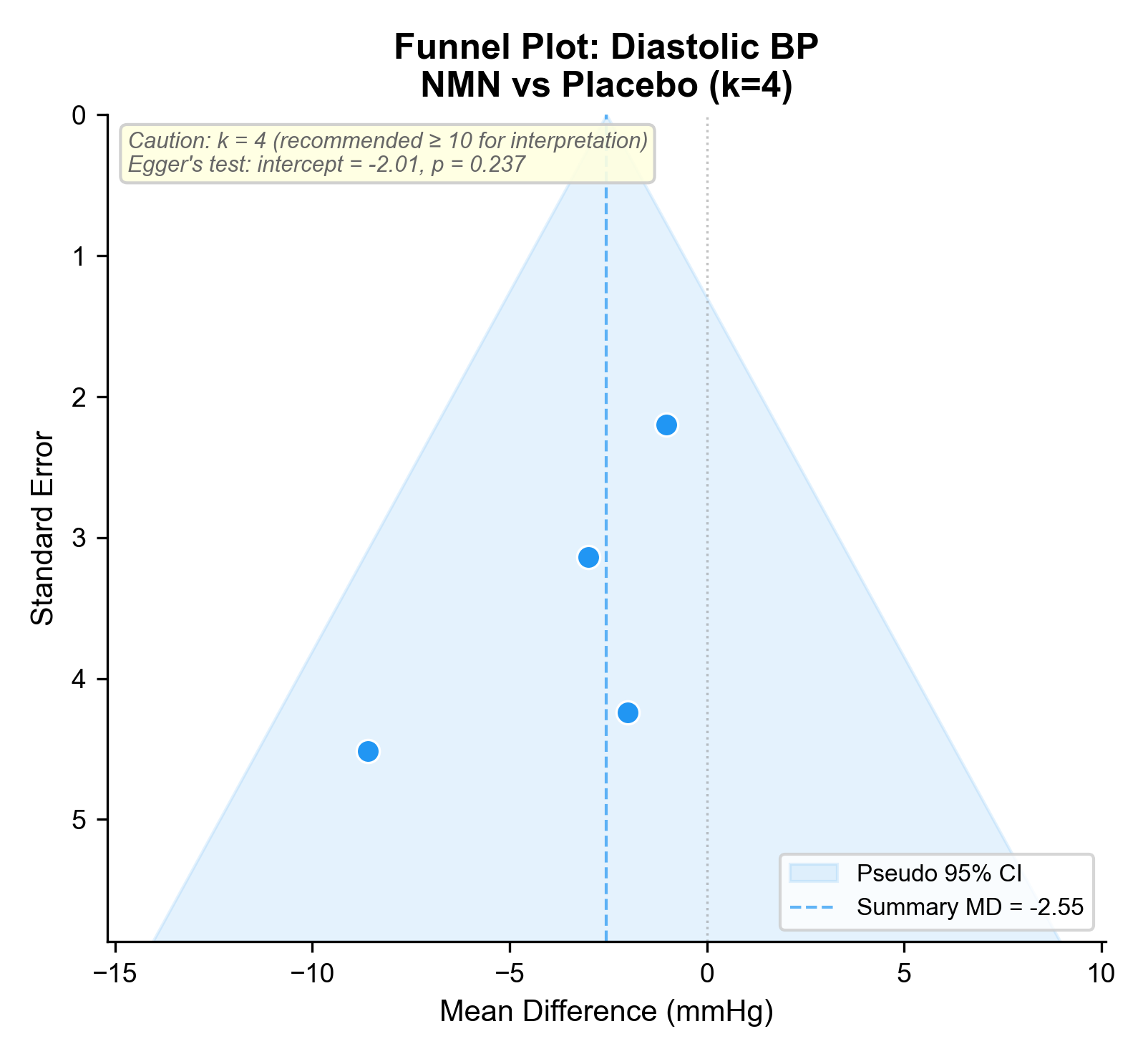

### funnel_FBG_NMN_vs_PBO.png

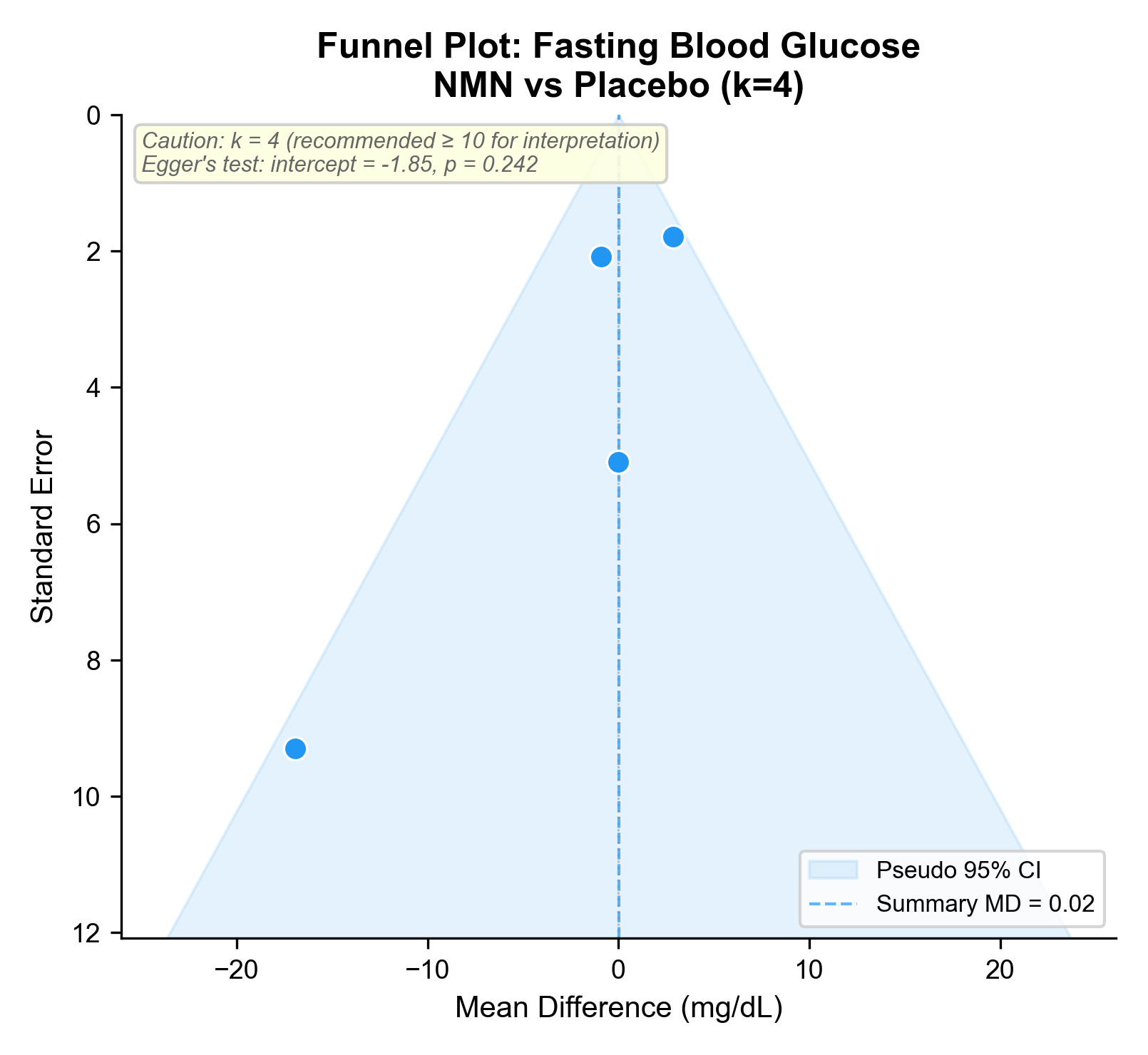

### funnel_HDL_NMN_vs_PBO.png

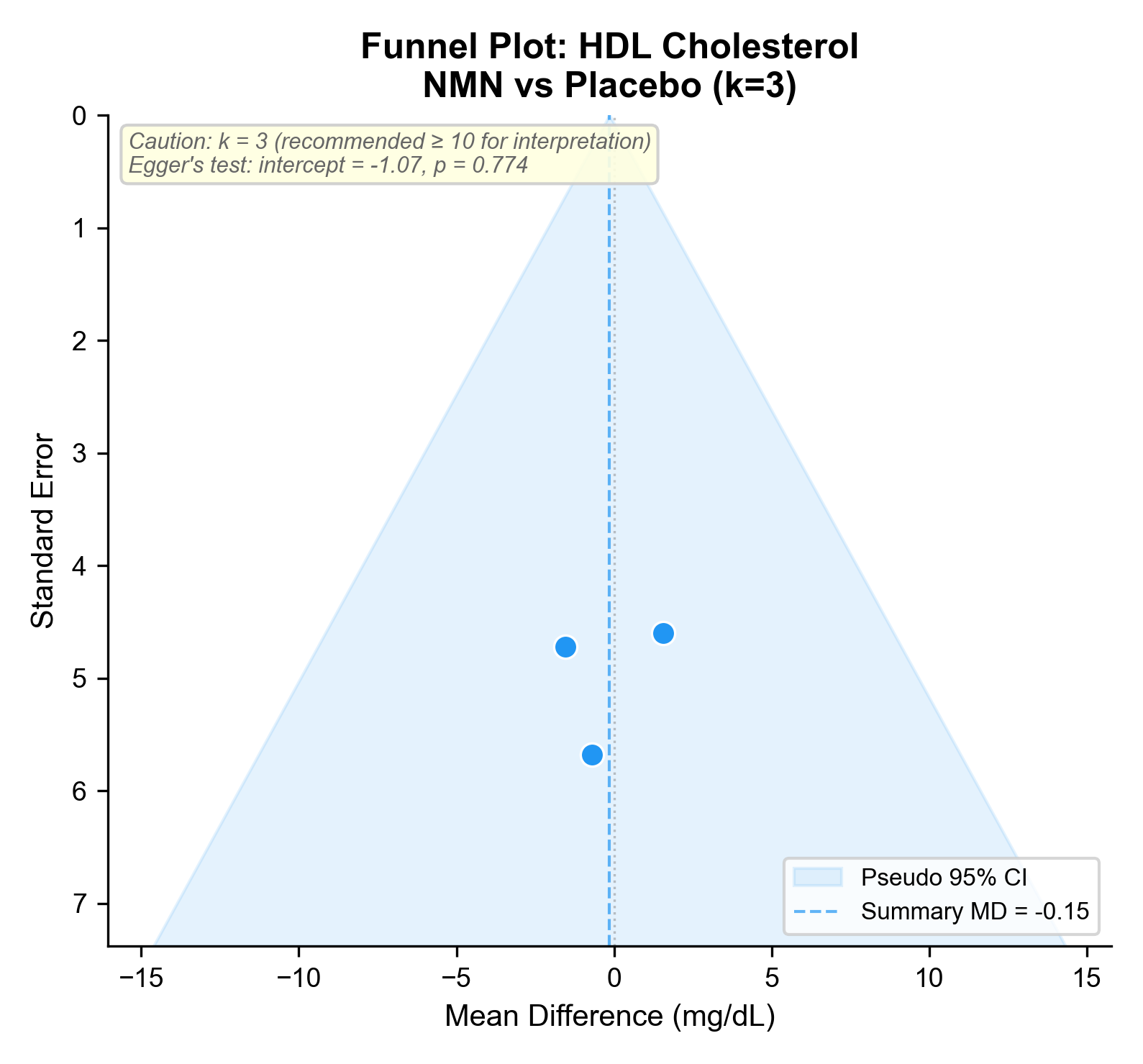

### funnel_HDL_NR_vs_PBO.png

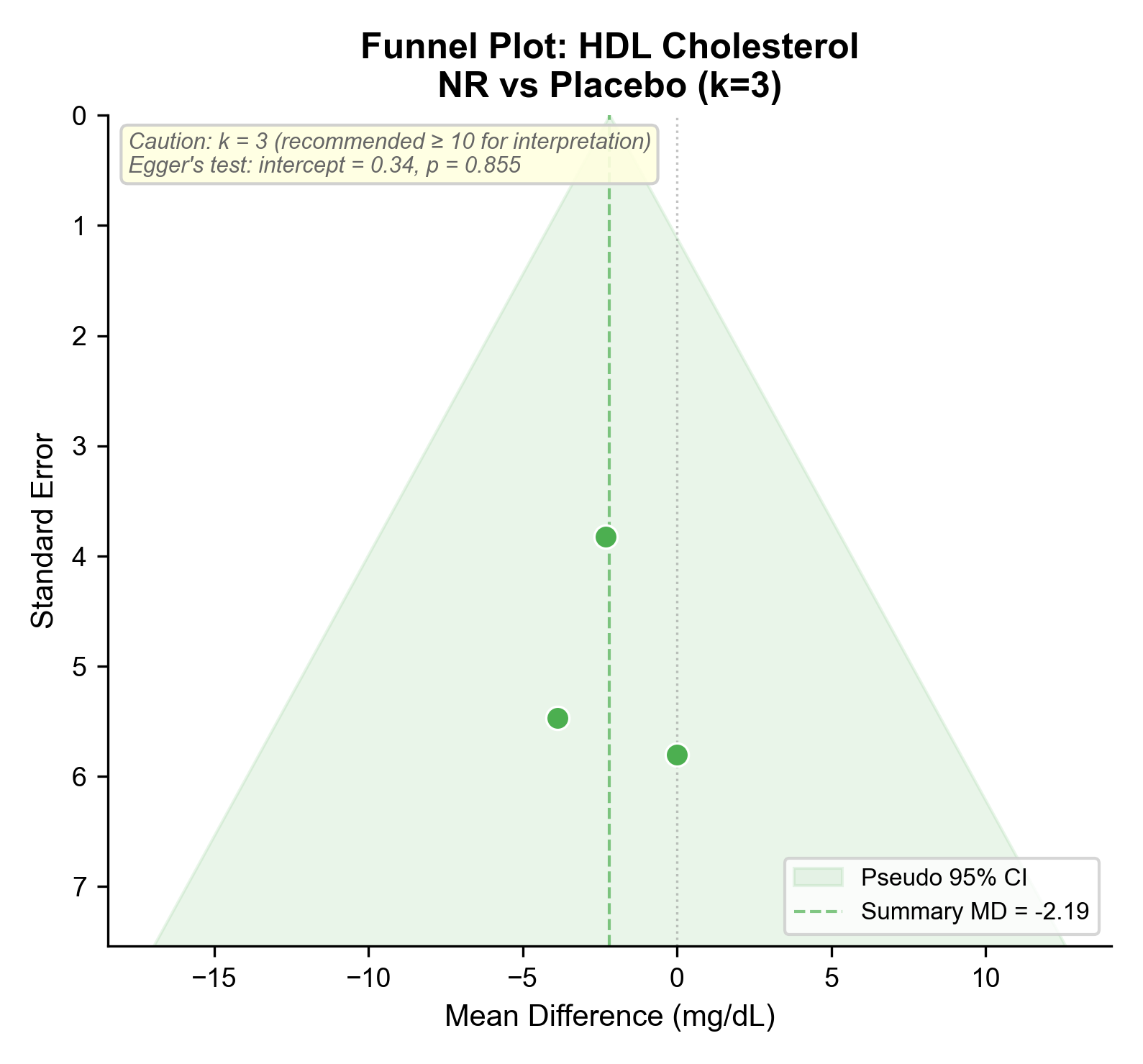

### funnel_LDL_NMN_vs_PBO.png

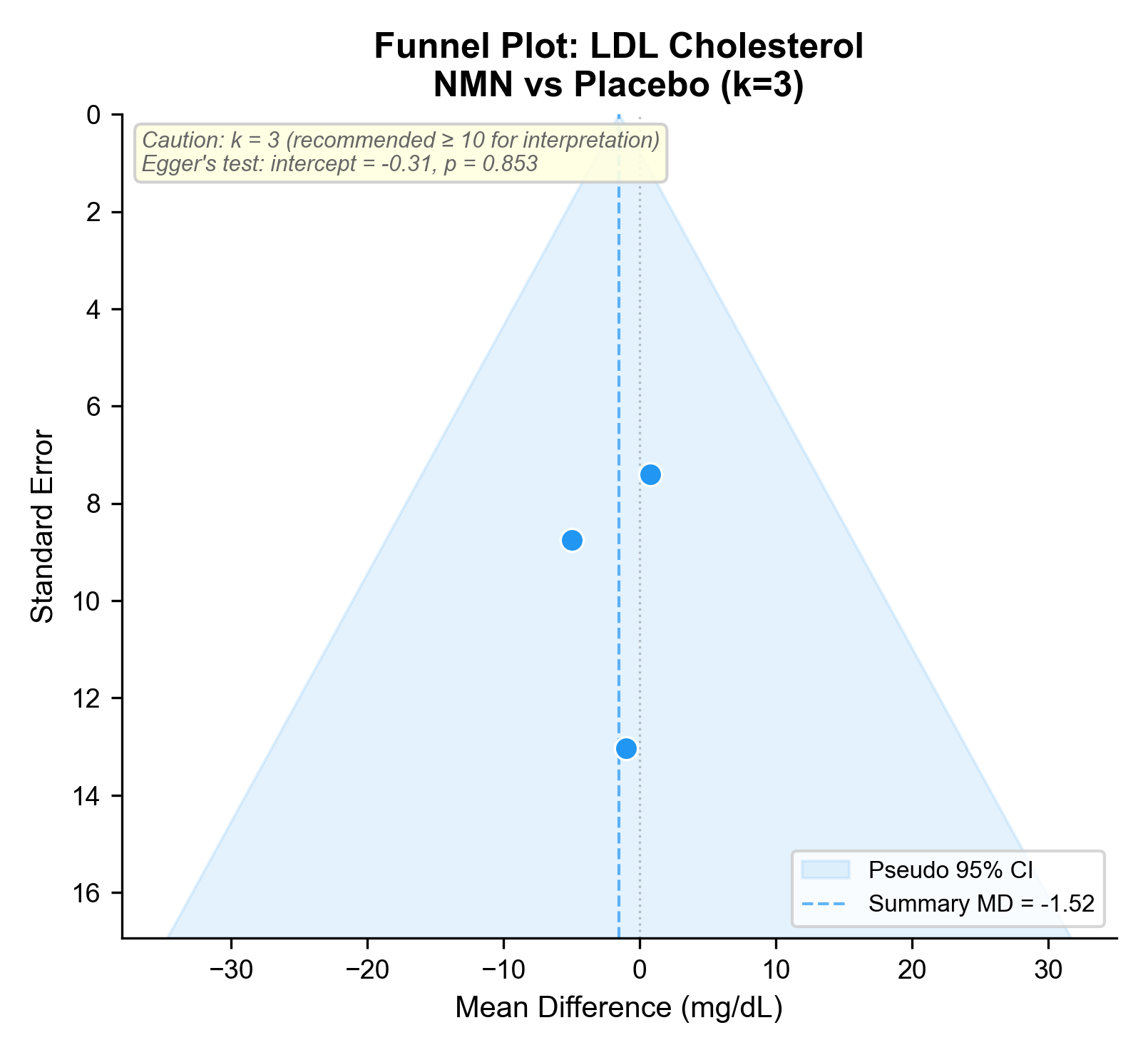

### funnel_LDL_NR_vs_PBO.png

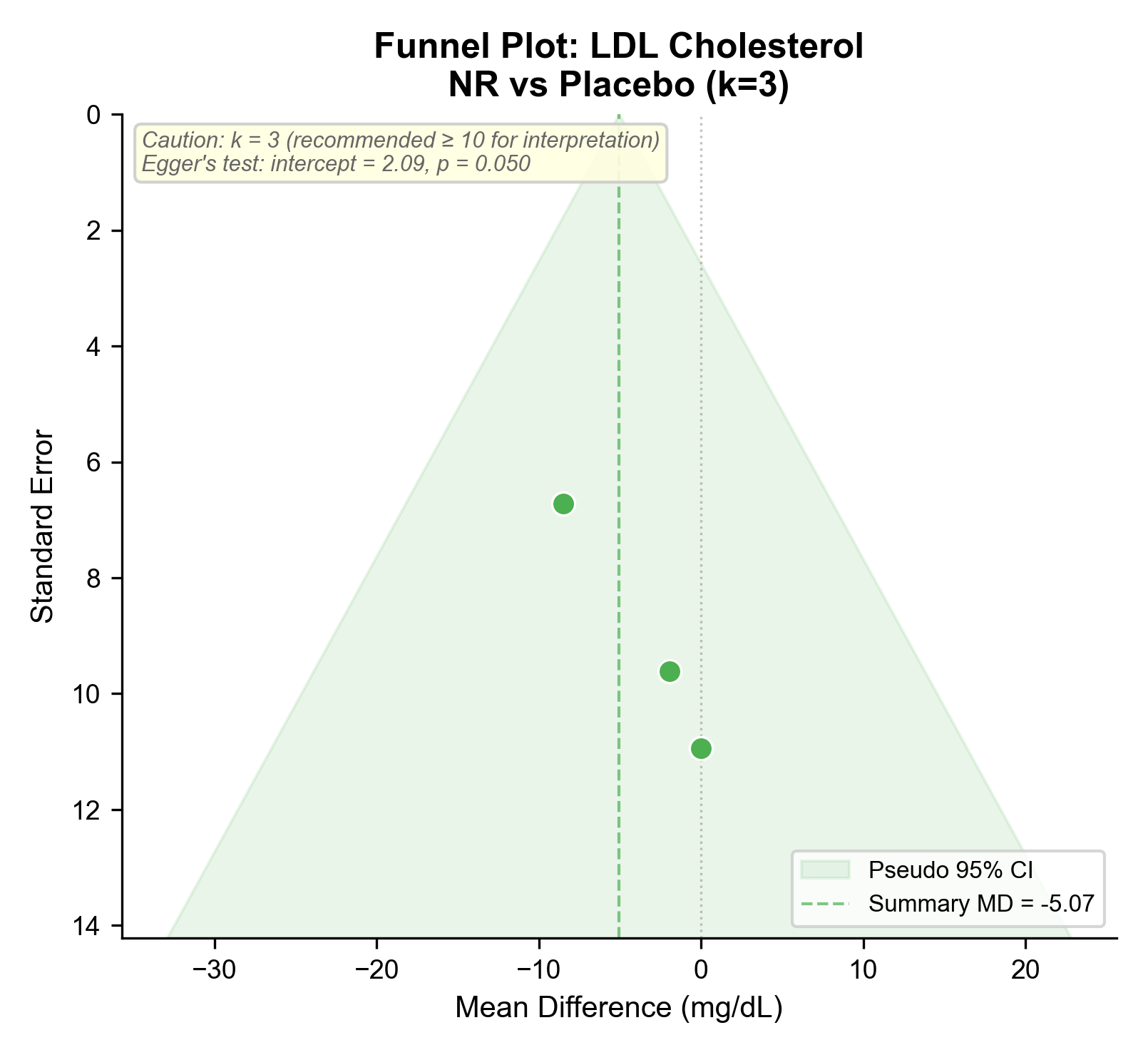

### funnel_SBP_NMN_vs_PBO.png

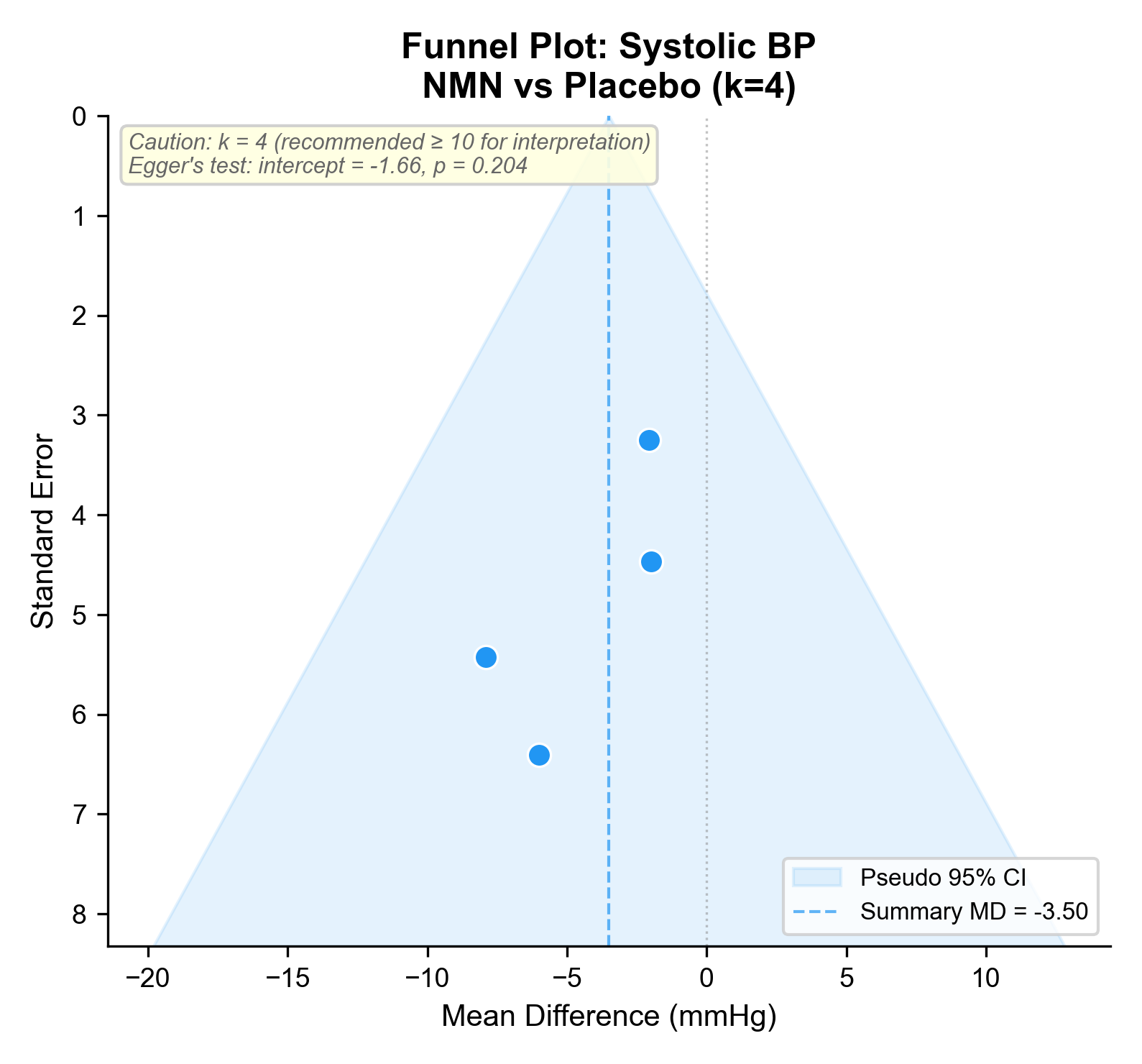

### funnel_TC_NR_vs_PBO.png

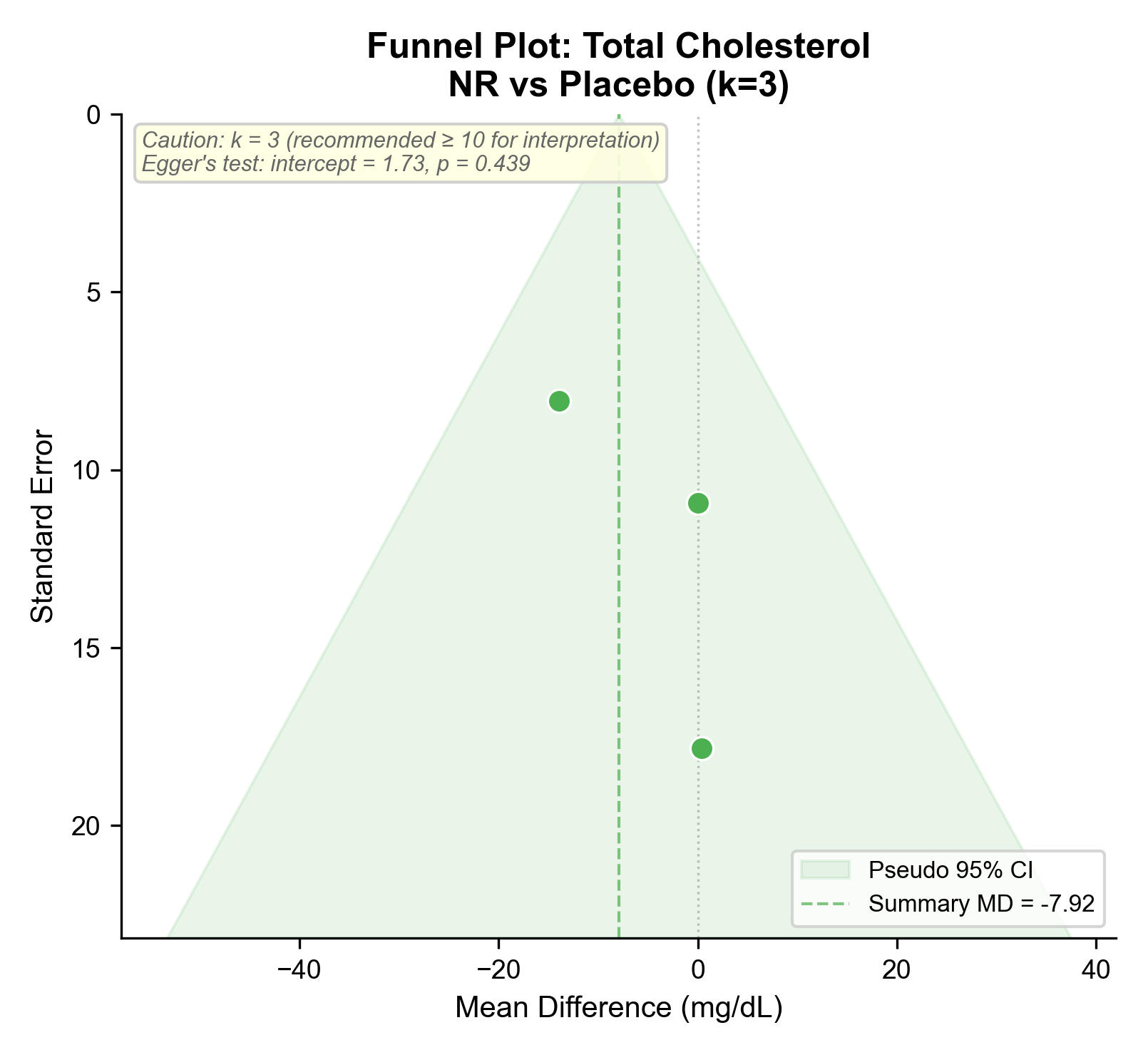

### funnel_TG_NMN_vs_PBO.png

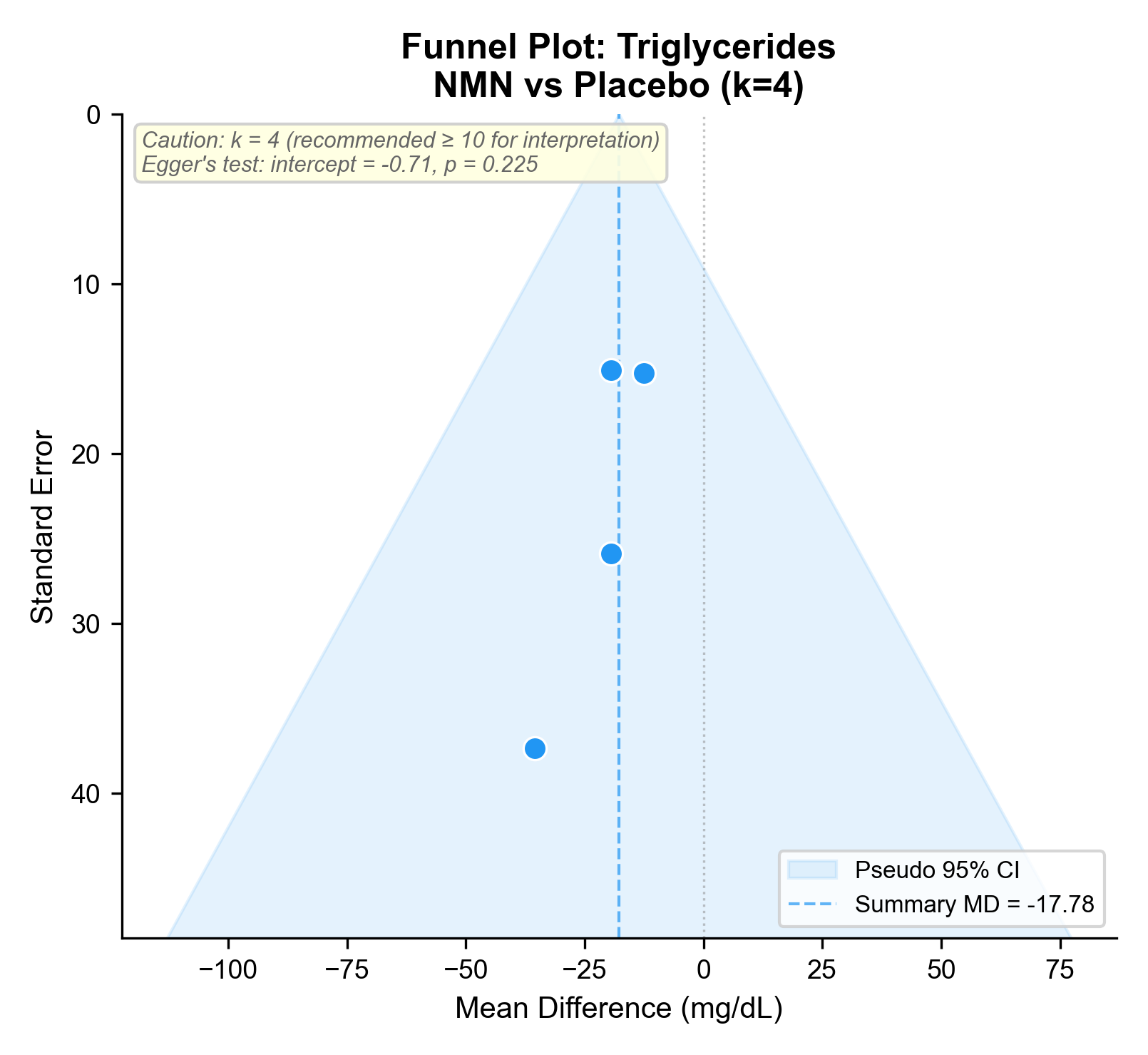

### funnel_TG_NR_vs_PBO.png

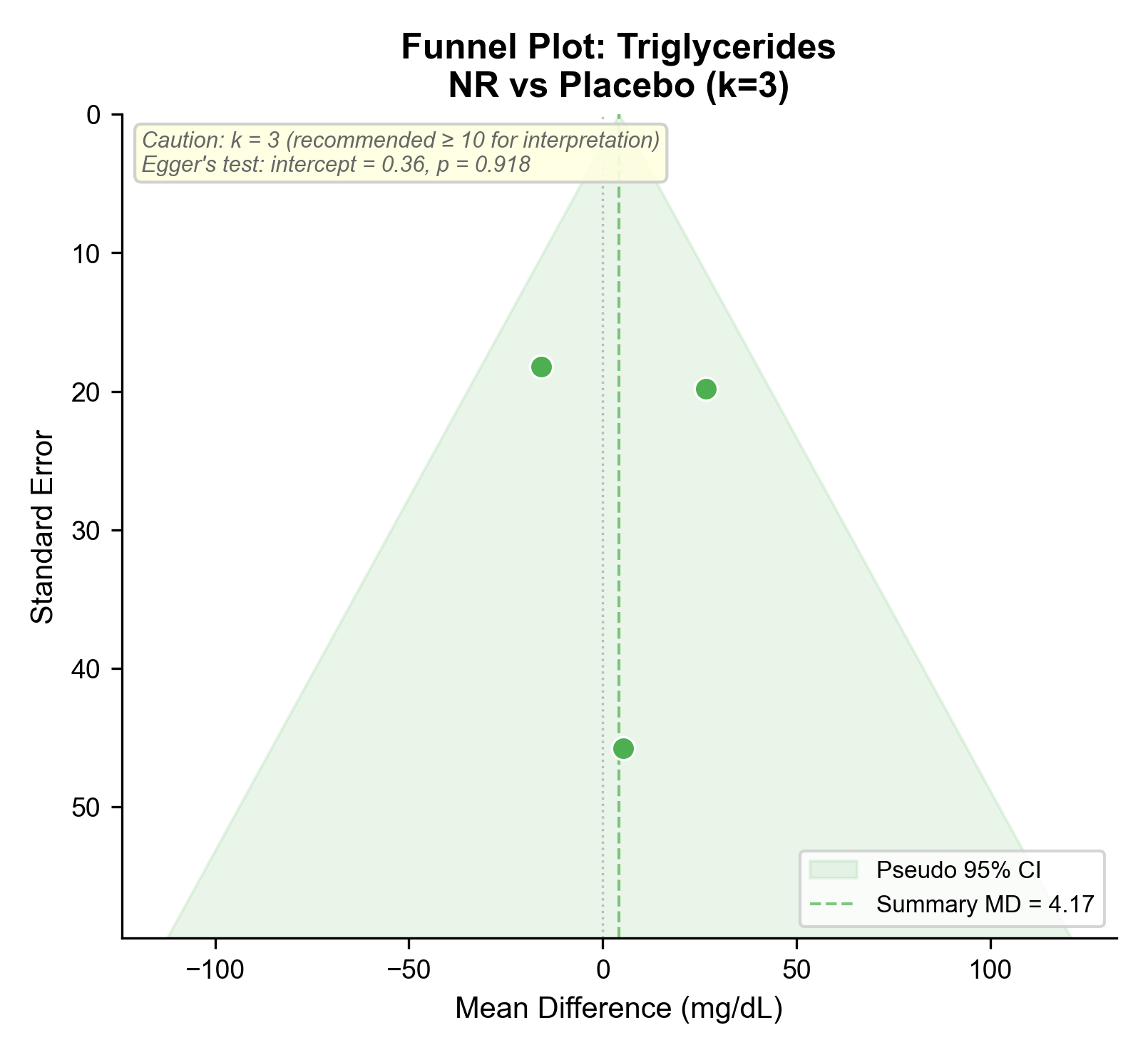

### outcome_reporting_matrix.png

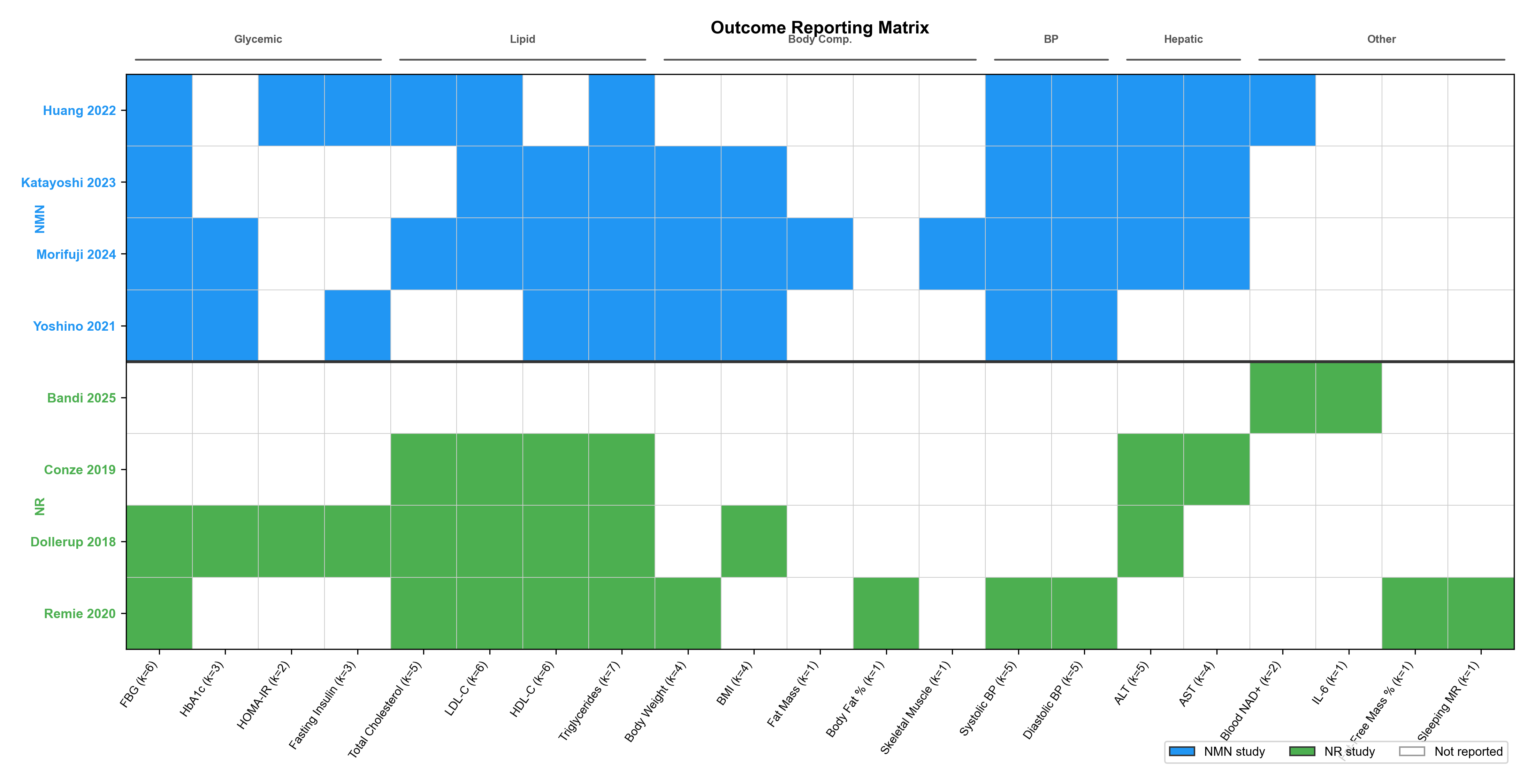

### sensitivity_loo_forest_ALT_NMN_pairwise.pdf

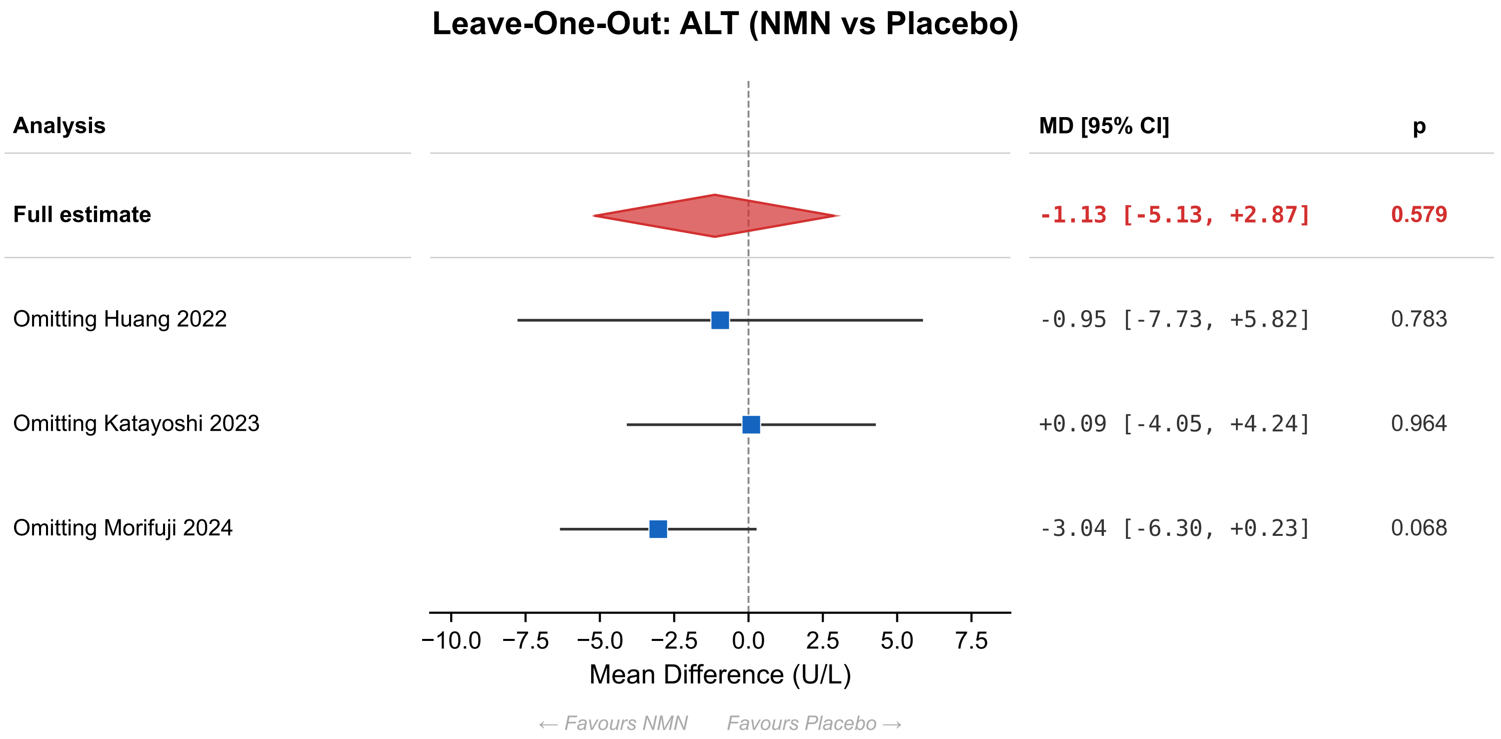

### sensitivity_loo_forest_ALT_NMN_pairwise.png

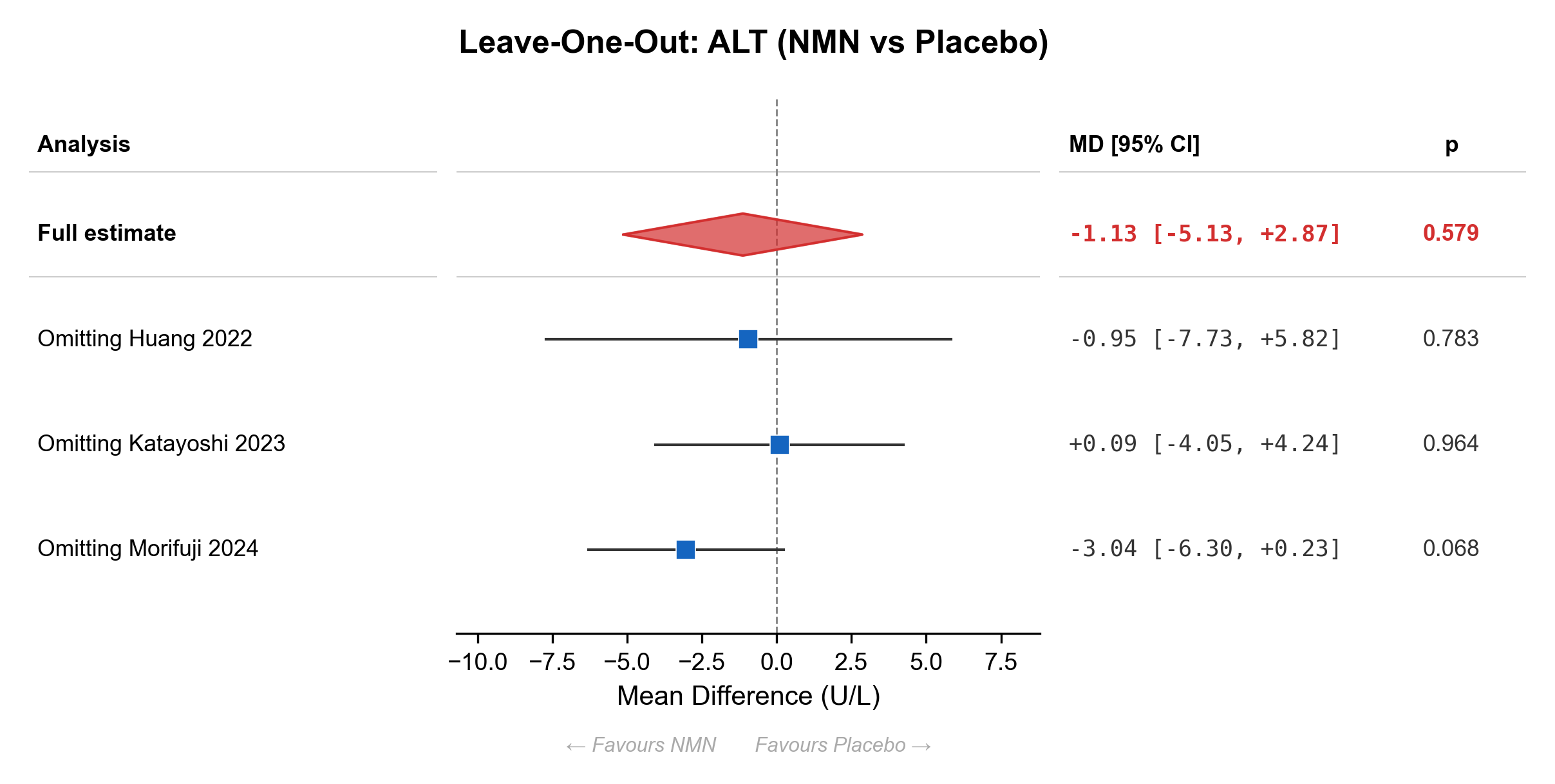

### sensitivity_loo_forest_fasting_insulin_NMN_pairwise.pdf

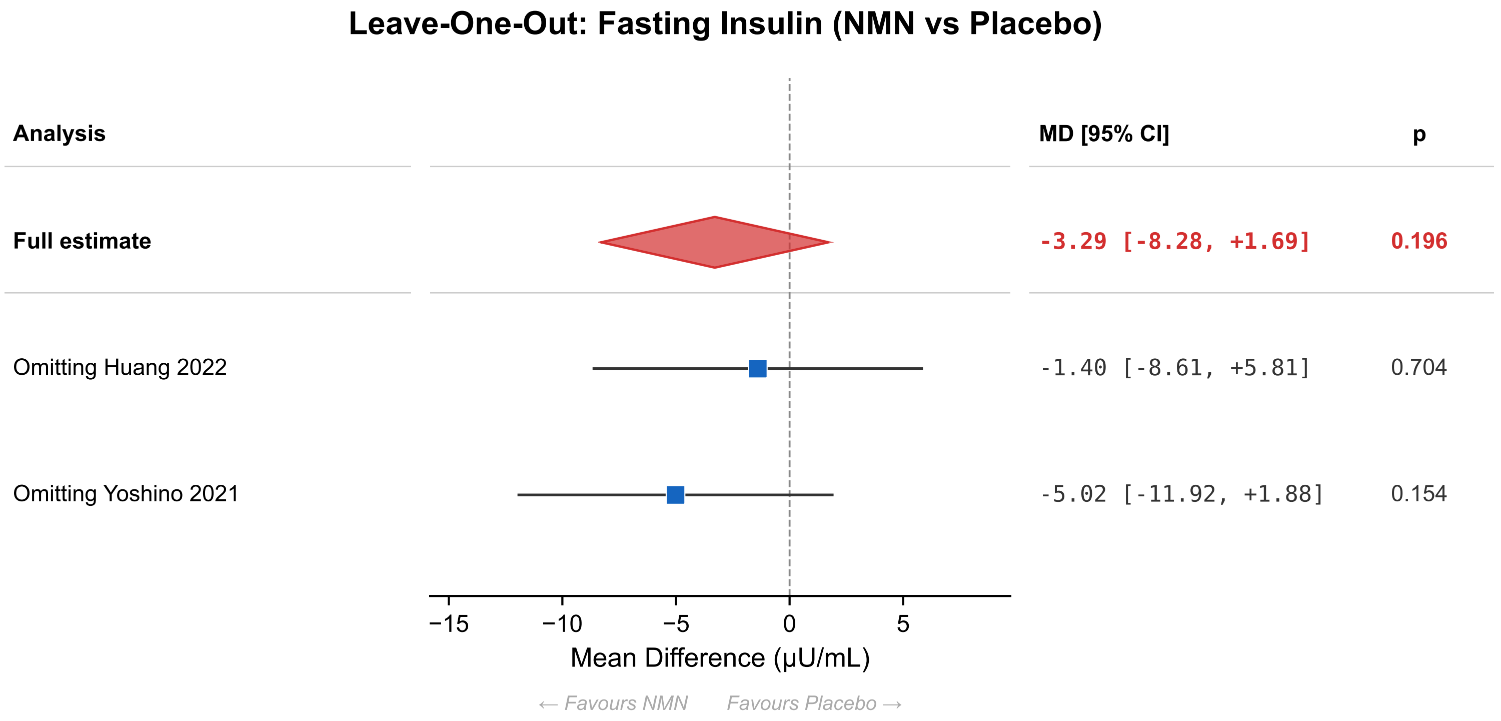

### sensitivity_loo_forest_fasting_insulin_NMN_pairwise.png

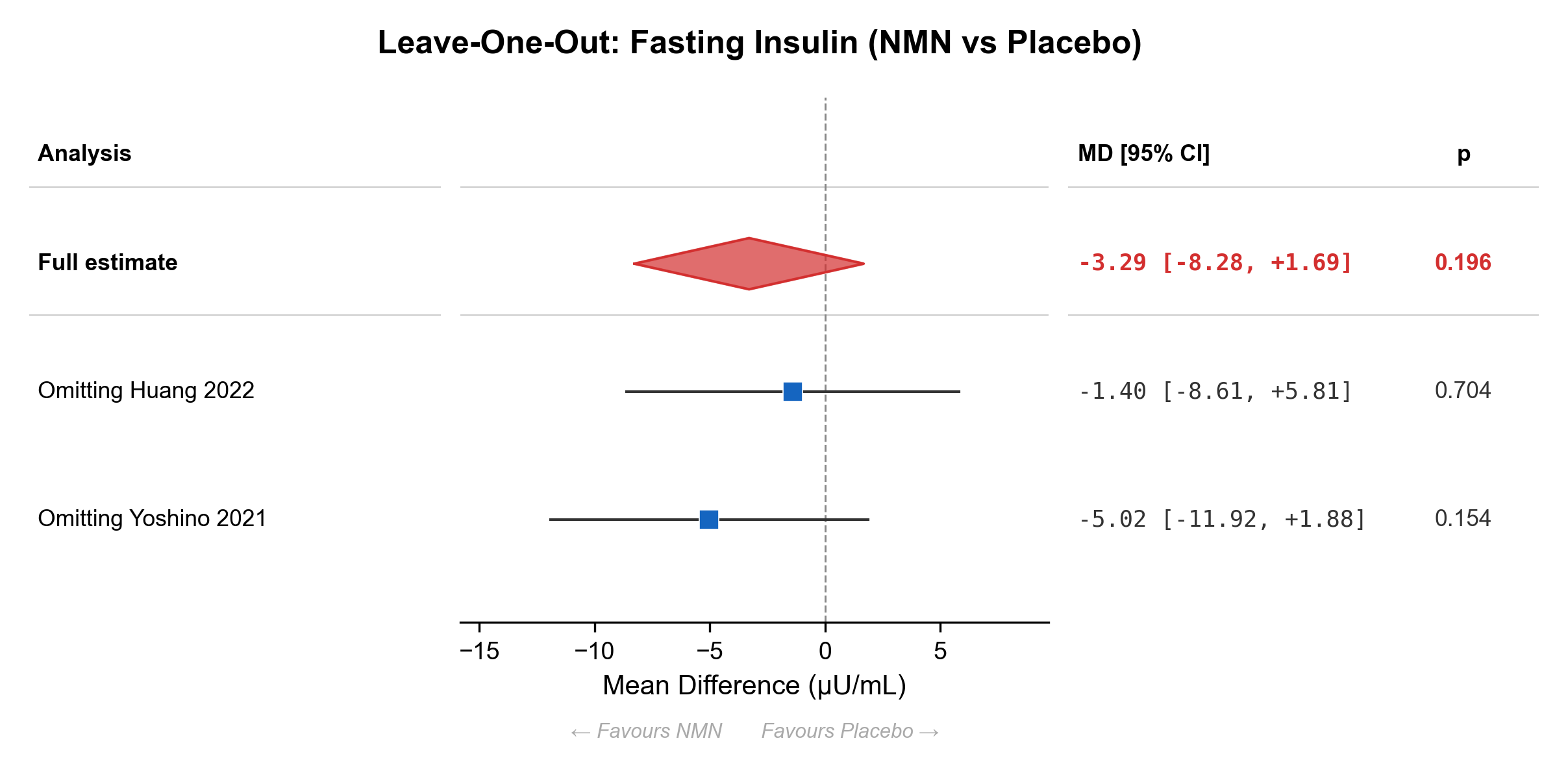

### sensitivity_loo_forest_SBP_NMN_pairwise.pdf

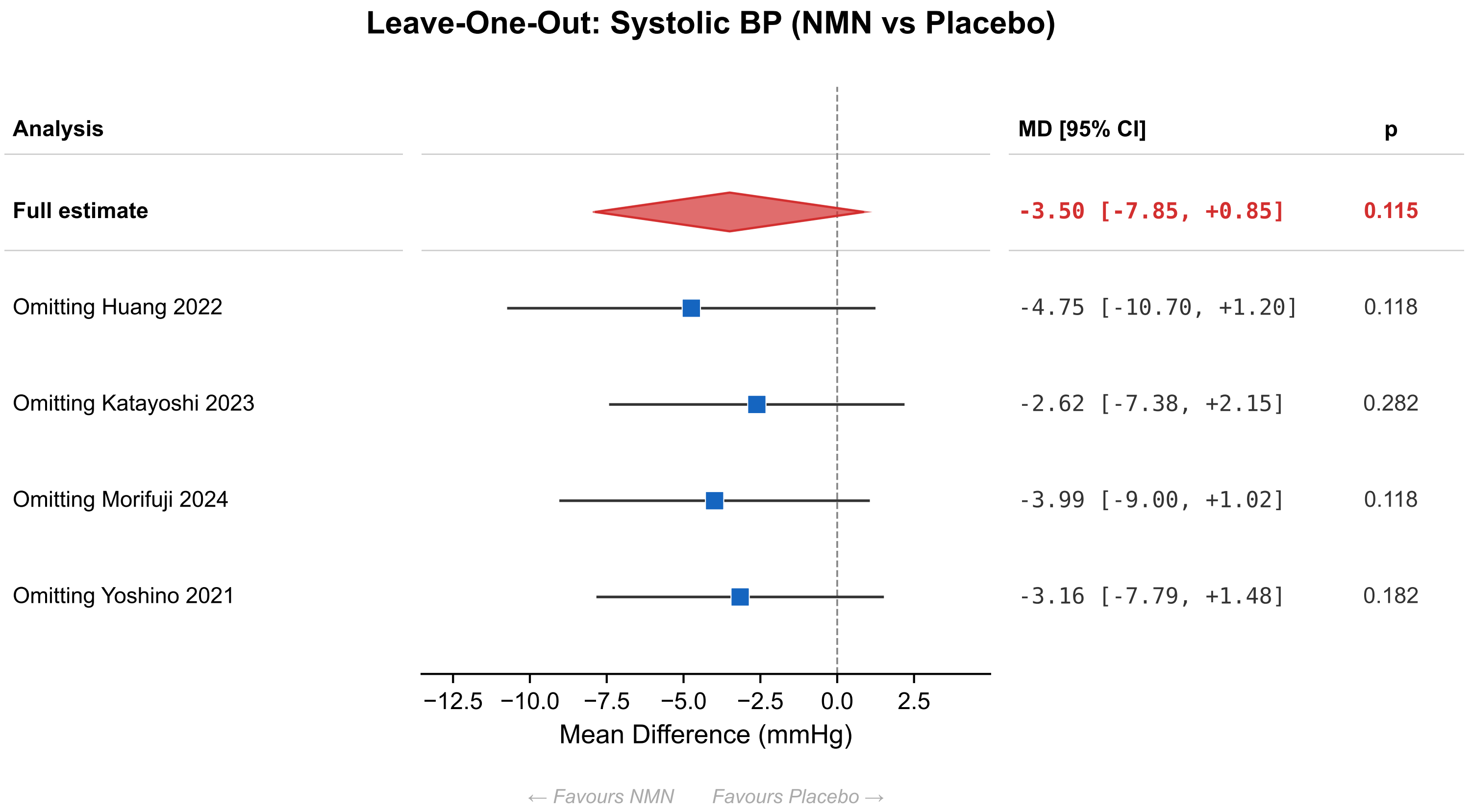

### sensitivity_loo_forest_SBP_NMN_pairwise.png

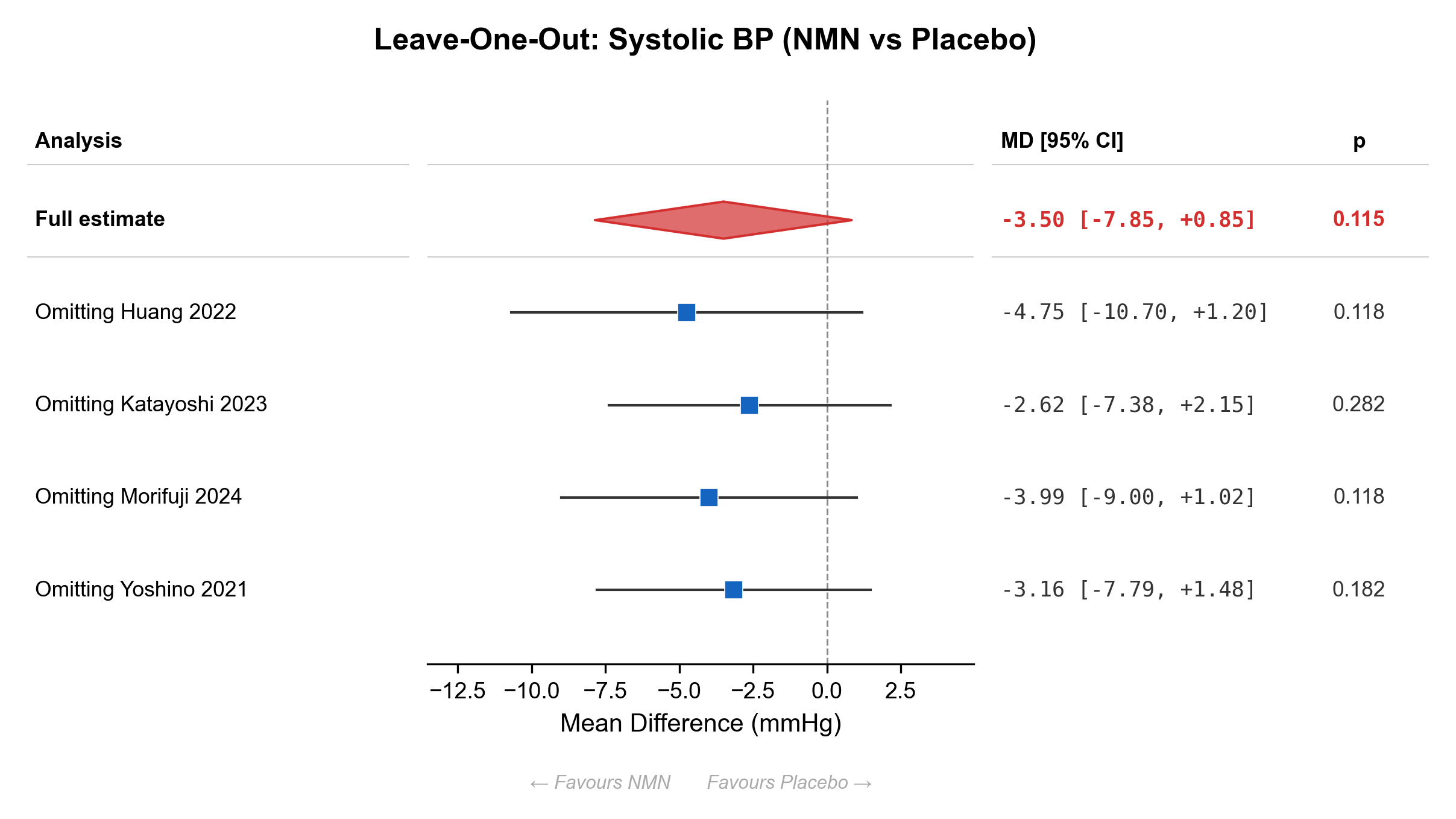
